# Evolution of lineage-specific trafficking proteins and a novel post-Golgi trafficking pathway in Apicomplexa

**DOI:** 10.1101/2022.12.12.520010

**Authors:** Christen M. Klinger, Elena Jimenez-Ruiz, Tobias Mourier, Andreas Klingl, Leandro Lemgruber, Arnab Pain, Joel B. Dacks, Markus Meissner

## Abstract

The Organelle Paralogy Hypothesis (OPH) posits a mechanism to explain the evolution of non-endosymbiotically derived organelles, predicting that lineage-specific pathways organelles should result when identity-encoding membrane trafficking components duplicate and co-evolve. Here we investigate the presence of such lineage-specific membrane-trafficking machinery paralogs in the globally important lineage of parasites, the Apicomplexa. Using a new phylogenetic workflow, we are able to identify 18 novel paralogs of known membrane-trafficking machinery, the emergence of several of which correlate with the presence of new endomembrane organelles in apicomplexans or their larger lineage. Gene coregulation analysis of a large set of membrane-trafficking proteins in Toxoplasma both corroborate known molecular cell biological interactions between characterized machinery and suggest involvement of many of these new components into established pathways for biogenesis of or trafficking to the microneme and rhoptry invasion organelles. Moreover, focused molecular parasitological analysis of the apicomplexan Arf-like small GTPases, and the ArlX3 protein specifically, revealed a novel post-Golgi trafficking pathways involved in delivery of proteins to micronemes and rhoptries, with knock down demonstrating reduced invasion capacity. The totality of our data has identified an unforeseen post-Golgi trafficking pathway in apicomplexans and is consistent with the OPH mechanism acting to produce novel endomembrane pathways or organelles at various evolutionary stages across the Alveolate lineage.

**Significance statement:** The mechanism of non-endosymbiotic organelle evolution has been relatively poorly explored and yet is relevant to many eukaryotic compartments, including the endomembrane system. The Organelle Paralogy Hypothesis predicts novel lineage-specific paralogs evolutionarily concurrent with emergence of new endomembrane organelles or pathways. By investigating this phenomenon in the apicomplexan parasites and their relatives, we identify and profile over a dozen new trafficking factors, several correlating with emergence of lineage-specific organelles. Cell biological study of one such factor demonstrates the existence of a novel post-Golgi trafficking pathway for components to the invasion organelles in the parasite *Toxoplasma gondii*. This work reveals how non-endosymbiotic organelle evolution has shaped cellular novelty in this lineage, relevant both to global health and fundamental evolutionary biology.

## Introduction

Eukaryotes are defined in part by the extensive presence of intracellular membrane-bound organelles. In addition to being a key steppingstone in eukaryotic evolution (1), the advent and expansion of organelles necessitated the development of protein machinery to facilitate the movement of protein and lipid components between organelles. The organelles and the machinery that mediates this movement together comprise the membrane-trafficking system (MTS), which is a key feature of all eukaryotes(2).

The MTS machinery is responsible for cargo loading and vesicle formation, vesicle scission, vesicle transport from donor to acceptor compartment, and eventual vesicle tethering and fusion(2, 3). Notably, a large amount of MTS machinery comprises paralogous gene families, wherein distinct paralogs perform the same basic function at a distinct cellular location. This observation led to the proposal of the organelle paralogy hypothesis (OPH), which posits that the duplication and co-evolution of MTS machinery encoding organelle identity facilitated the diversification of an ancestral organelle(s) to give rise to the diversity of organelles found in modern eukaryotes(1, 4).

Molecular evolutionary studies have reconstructed the presence of an extensive MTS machinery complement in the last eukaryotic common ancestor (LECA).. Cell biological studies have confirmed a general conservation of organelles themselves, with most eukaryotes possessing identifiable homologs of an endoplasmic reticulum (ER), Golgi apparatus, early and late endolysosomal compartments, mitochondria, and peroxisomes(2). Despite evidence of clear precursor homologs for some of these MTS components, orthologs of the organelle or pathway-specific protein machinery are absent, with the overall conclusion of these data that ancestral MTS machinery present in the first eukaryotic common ancestor (FECA) expanded and diversified into a complex set of machinery in the LECA, complete with an extensive organelle complement(1,4, 10).

Despite the clear presence of pan-eukaryotic MTS orthologs, indicating an ancient origin and widespread retention, not all modern eukaryotes retain a LECA complement. The parasitic organism *Giardia*, causative agent of giardiasis, possess only an ER, Golgi-like encystation vesicles, and “peripheral vacuoles” (more recently termed peripheral endocytic compartments (PECs)(11),; this reduced organelle complement correlates with a similar reduction in retained MTS machinery. On the other side of the spectrum, organisms like *Arabidopsis thaliana* have massively expanded some MTS families, such as Rab GTPases (12).

We have previously argued that, while the OPH was initially proposed to provide a mechanistic framework to explain the FECA-LECA MTS transition, it is also likely to have continued functioning in eukaryotes post-LECA(2, 10). This is in part motivated by the presence of additional organelles in many eukaryotes, including secretory granules and melanosomes in human cells, as well as other enigmatic organelles such as the contractile vacuole present in diverse eukaryotes(12, 13). However, the extent and ways in which the OPH have shaped the diversity of lineage-specific endomembrane organelles is largely unexplored. Such an exploration requires a eukaryotic lineage with clear novel organelles, available genomic data, and, ideally, the capacity for molecular biological study.

The Apicomplexa are a phylum of unicellular eukaryotic parasites that invade the cells of a variety of hosts, including humans. Although almost universally parasitic, apicomplexans possess free-living relatives including chromerid (representing colpedellids) and dinoflagellate algae, many members of which are photo- or mixotrophic(14–16); together, these taxa form a group known as the Myzozoa. Molecular and ultrastructual data support the presence of numerous unique organelles in apicomplexans, including the endo-lysosomal micronemes and rhoptries, dense granules, and the innermembrane complex (IMC, a series of connected membranous sacs subtending the plasma membrane)(17). Furthermore, the apicoplast, a relict non-photosynthetic plastid present in apicomplexans apart from *Cryptosporidium spp*., is unique among secondary (or higher levels) red plastids in that it is present outside of the ER and is surrounded by four membranes (18, 19). Curiously, some of these structures have presumed homologs in closely related taxa. Structures resembling micronemes and rhoptries are present in other myzozoans(20), and the IMC is considered homologous to the alveoli of ciliates (which as a basal group to the Myzozoa form the alveolate clade) (21,22). In addition to the clear presence of unique organelles in apicomplexans and their close relatives, genomic data exists for all the main groups of alveolates, allowing for comparative genomic analyses. Within the Apicomplexa, *Toxoplasma gondii* provides a useful system for conducting molecular biology due to its ease of culture, haploid genome, and available range of genetic tools (23, 24).

Hence, we set out to analyze the possibility of an OPH-like mechanism giving rise to the novel organelles present in the Apicomplexa and their close relatives. Through large-scale comparative genomic and phylogenetic analyses of paralogous MTS families, we were able to identify 18 paralogs found in Apicomplexa but not found across eukaryotes. Of these paralogs, we chose three, all from the ARF-related (Arl) family, for further investigation. Although we localize all three novel Arls in asexual *T. gondii* parasites, only one proved essential following genetic disruption. This protein, termed ArlX3, localizes primarily to the *trans-*Golgi network (TGN), and results in the mislocalization of microneme, rhoptry, and apicoplast cargo proteins when knocked down as well as a general fragmentation of the Golgi itself. Overall, our results identify the presence of novel trafficking paralogs in an enigmatic group of eukaryotic parasites and provide insight into the possibility of modern examples of the OPH in shaping eukaryotic evolution.

## Results

### A bioinformatics screen to identify LSPs in Apicomplexa and related taxa

In order to identify paralogs with a restricted phylogenetic distribution, hereafter referred to as lineage-specific paralogs (LSPs), we first carried out both HMMer and BLAST searches to identify all homologs for each family in our dataset (Table S1). Next, we carried out phylogenetic analyses using proteins of known identity to classify any pan-eukaryotic paralogs from a series of smaller taxon-specific datasets. Finally, we collected all previously unclassified sequences for each family and performed additional phylogenetic analyses to identify those that formed supported monophyletic clades; these clades were then run with previously classified pan-eukaryotic clades to establish their possible origins (Figure 1A, Materials and methods, Supplementary text).

**Figure 1.**
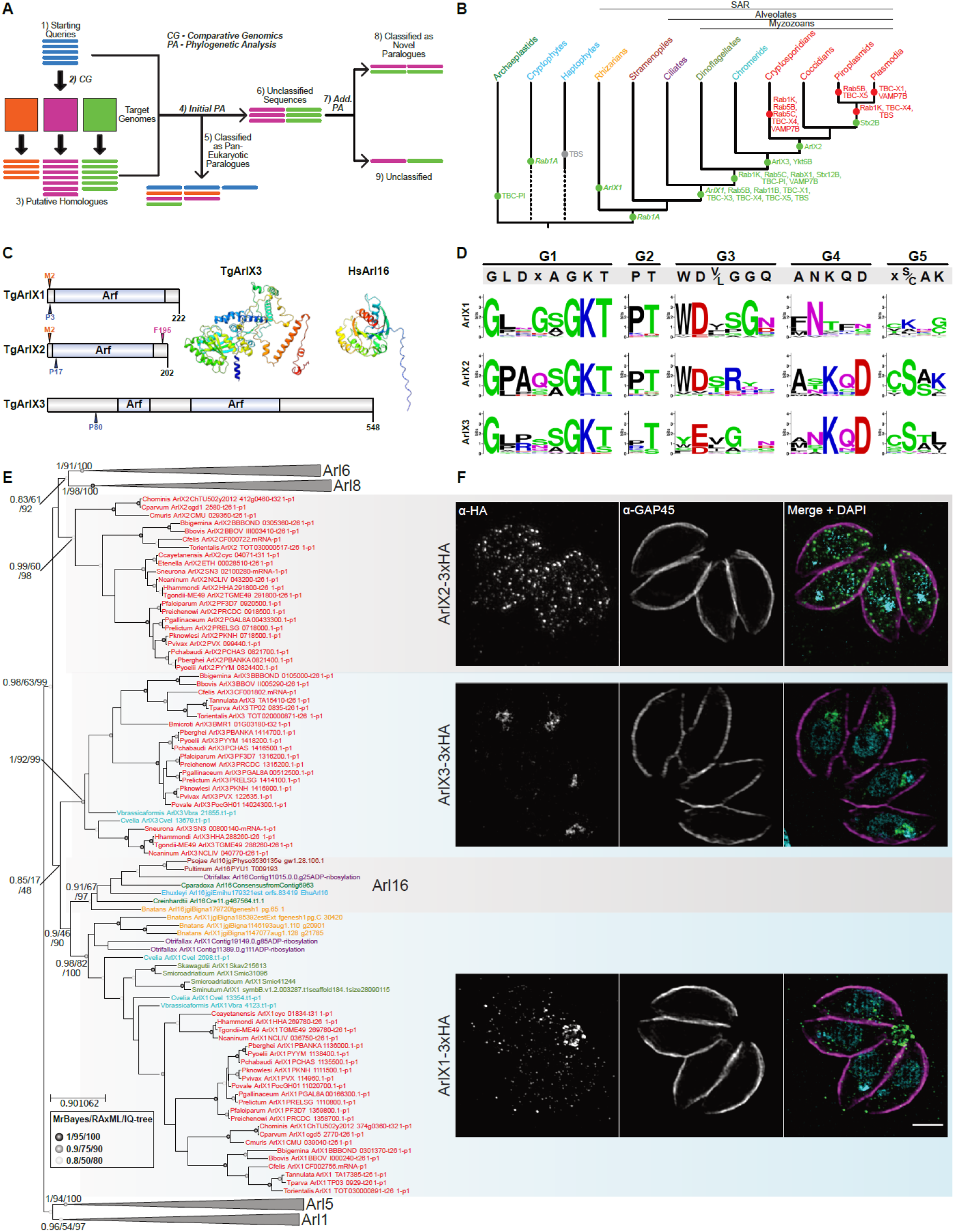
A phylogenetic screen identifies LSPs in Apicomplexa and their close relatives. This figure summarizes the methodology behind and results of a phylogenetic screen to identify lineage-specific paralogs (LSPs) in apicomplexans. A) Overview of the screen. Homologs of each trafficking family were identified (1–3), and all pan-eukaryotic orthologs were identified via phylogenetic analysis with known marker sequences (4–5). Any unclassified sequences (6) were then ran in additional phylogenetic analyses (7); some formed monophyletic clades (LSPs; 8) while others remained unclassified (9). B) Summary of presumed patterns of gain and loss of LSPs in study taxa. Green dots represent gain while red dots represent loss; bold italics represent uncertain provenance (between independent gain or ancient gain and subsequent loss). The single grey dot represents the presence of an analogous TBS protein in *E. huxleyi*. Taxa are colour-coded: Apicomplexa, red; chromerids, teal; dinoflagellates, light green; ciliates, purple; stramenopiles, brown; Rhizaria, orange; cryptophytes/haptophytes, blue; archaeplastids, dark green. Larger taxonomic groups are summarized above. C) Schematic overview of Arl LSPs in *T. gondii*. Each sequence is represented as a grey bar (length shown below each), with Pfam Arf (PF00025) domains in light blue. The highest scoring predicted site for each type of lipid modification (F, farnesylation; M, myristoylation; P, palmitylation) is shown. The 3-D structure of TgArlX3 is shown, coloured from N-(red) to C-terminus (blue). D) Overview of G-box motifs in Arl LSPs, with canonical Arf family residues listed above. Note the overall lack of conservation at most positions. E) Phylogenetic analysis of Arl LSPs. The best Bayesian topology is shown with RAxML bootstrap (RB) and IQ-Tree rapid bootstrap (IB) support mapped. Important node support is shown in the order (Bayesian posterior probability/RB/IB) while internal node support is denoted by symbols as per figure legend. Scale bar is number of substitutions per site. F) Arl LSP localization in *T. gondii*. Intracellular parasites, each line with a different Arl LSP C-terminally tagged with a 3xHA tag, were stained with α-HA antibodies (coloured green in the merge); parasites were outlined using α-Gap45 (an IMC marker, shown in magenta). Parasite nuclei were stained with DAPI (blue in the merge). Note the distinct localization pattern of each Arl LSP.

We analyzed the following paralogous families: SM proteins, SNARE proteins, Rab GTPases, the TBC family of Rab GTPase-activating proteins (RabGAPs), ARF family G proteins, and their GAP and GTPase effector protein (GEF) regulators (Figure S1, Table S1). The results here highlight notable results from several families, focussing on the presence of LSPs; full details for each family can be found in the supplementary text. Overall, we identified 18 LSPs in Apicomplexa and their relatives; the presumed patterns of gain and loss for these LSPs is summarized in Figure 1B.

### SNAREs

Soluble *N*-ethylmaleimide sensitive factor (NSF) attachment receptor proteins (SNAREs) are coiled-coil proteins, often membrane-associated, that function in vesicle membrane fusion(25). As expected, we identified homologs of the Qa, Qb, Qc, and R SNAREs across our study taxa (Figure S1, Supplementary text). There was a clear duplication of Stx12, giving rise to a myzozoan Stx12B paralog (Figures 1B, S1, S2C). Likewise, we identified duplications of the R SNAREs VAMP7 (in Myzozoa) and Ykt6 (in the apicomplexan-chromerid ancestor, Figures 1B, S1, S2K, S2L). The presence of Qbc SNAREs in Apicomplexa and sparsely across our study dataset (Figure S1, Table S1) prompted us to analyze the origins of the Qb and Qc domains from these proteins. The Qb domain is most closely related to NPSN11 (Figure S2H) and our data raises the intriguing possibility of a close evolutionary link between the Qc domain and SYP71 or Use1 (Figure S2I). Although it did not meet our criteria for a LSP, there was also a clear duplication of the Qa SNARE Stx2 in hematozoans (Figures 1B, S2B).

### Rab GTPases

Rabs are small (~200 amino acid) GTPases from the Ras superfamily that function in membrane trafficking and a variety of other cellular functions (26). Overall, our results confirm the previously reported relationships between Rabs, including the overall division into endo/exocytic clades (Figures S2M, S2N, Supplementary text)(5). We confirm the presence and taxonomic distribution of several previously reported Rab LSPs (Figure 1B): Rab1A, present across SAR as well as the cryptophyte *Guillardia theta* (Figures S1,S2O)(27); Rab11B, an alveolate-specific Rab11 paralog (Figures S1, S2R)(28); Rab5C, a myzozoan-specific Rab5 paralog (Figures S1, S2Q); and Rab5B, which is known as an atypical Rab5-like protein that we confirm here is conserved across alveolates (Figures S1, S2Q)(29). Additionally, we report the presence of two previously unreported Rab LSPs. Rab1K is a further duplication of Rab1 that is restricted to chromerids, dinoflagellates, and coccidian apicomplexans closely related to *T. gondii* (Figures 1B, S1, S2O). RabX1 is a myzozoan duplication of RabL2/RTW (Figures 1B, S1, S2S).

### TBC RabGAPs

The Tre-2/Bub-2/Cdc 16 (TBC) proteins are a diverse family of RabGAPs unified by the presence of a TBC domain. Overall, our results confirm the previously reported relationships between TBCs (Figures S2T, U, Supplementary text)(8). We identify five TBC LSPs, simply termed TBC-X1 through TBC-X5 (Figures 1B, S1). TBC-X2 groups with previously reported archaeplastid-specific TBC-PI proteins (Figure S2W), suggesting a more complex origin for this group. TBC-X1, 3, 4, and 5 clades instead represent alveolate-specific duplications of TBC-Q (Figure S2X). Finally, a group of previously reported proteins containing both an ArfGEF Sec7 and TBC domain (TBS proteins)(30) are confirmed here to be restricted to alveolates, with analogous proteins present in the haptophyte *Emiliania huxleyi* (Figures 1B, S2V, S3L, SY)(9).

### ARF family G proteins

Like Rabs, ADP-ribsosylation factor (ARF) family proteins, including ARF, ARF-like (Arl), and Sar proteins, are members of the Ras superfamily(31, 32). We identified three Arl LSPs, simply termed ArlX1 (conserved in alveolates and the rhizarian *Bigelowiella natans*), ArlX2 (found only in apicomplexans), and ArlX3 (present in chromerids and apicomplexans but absent from cryptosporidians, Figure 1B, Table S1). While TgArlX1 and TgArlX2 are similar in size to other canonical Arl homologs (~200 amino acids), TgArlX3 is almost three times the length with the canonical Arf domain split into two (Figure 1C). Molecular modeling (materials and methods) suggests the presence of additional N-terminal helices, an internal loop between the canonical Arf folds, and an unstructured C-terminus (Figure 1C). Analysis of conserved G-box motifs among ArlX homologs suggests extensive variability among orthologs and some notable divergence from the eukaryotic ARF consensus sequence (Figure 1D). Despite extensive analyses (Supplementary text, Figures S3A-S3J), we were unable to determine a clear origin for all three ArlX proteins. ArlX1 is most likely a duplication of Arl16 while ArlX2 appears similar to Arl6; ArlX3, sequences of which are larger and more divergent than most Arl proteins, could not be resolved as related to any particular pan-eukaryotic Arl sub-family (Figure 1E). Its origins remain unclear. Despite the presence of three novel Arl proteins, there were no LSPs present for either ArfGEF or ArfGAP families (Figures S1, S3K-M, Supplementary text); other regulators may govern these Arls.

### Analysis of LSP co-expression identifies blocks of co-regulated trafficking components

Assuming that functionally related LSPs will show correlated gene expression profiles, gene functions can potentially be linked using transcriptome data (33). We therefore looked for patterns of gene co-expression of LSPs in publicly available RNA-seq datasets from a range of *T. gondii* life stages (Table S2). We compared the LSPs with a known suite of existing paralog families. We further looked for co-regulation between the LSPs and known proteins of the Mic and Rop machinery to assess whether any might be involved in the biogenesis or secretion of proteins targeted to the micronemes or rhoptries respectively. Three blocks of co-regulated machinery of note were observed (Figure 2).

**Figure 2.**
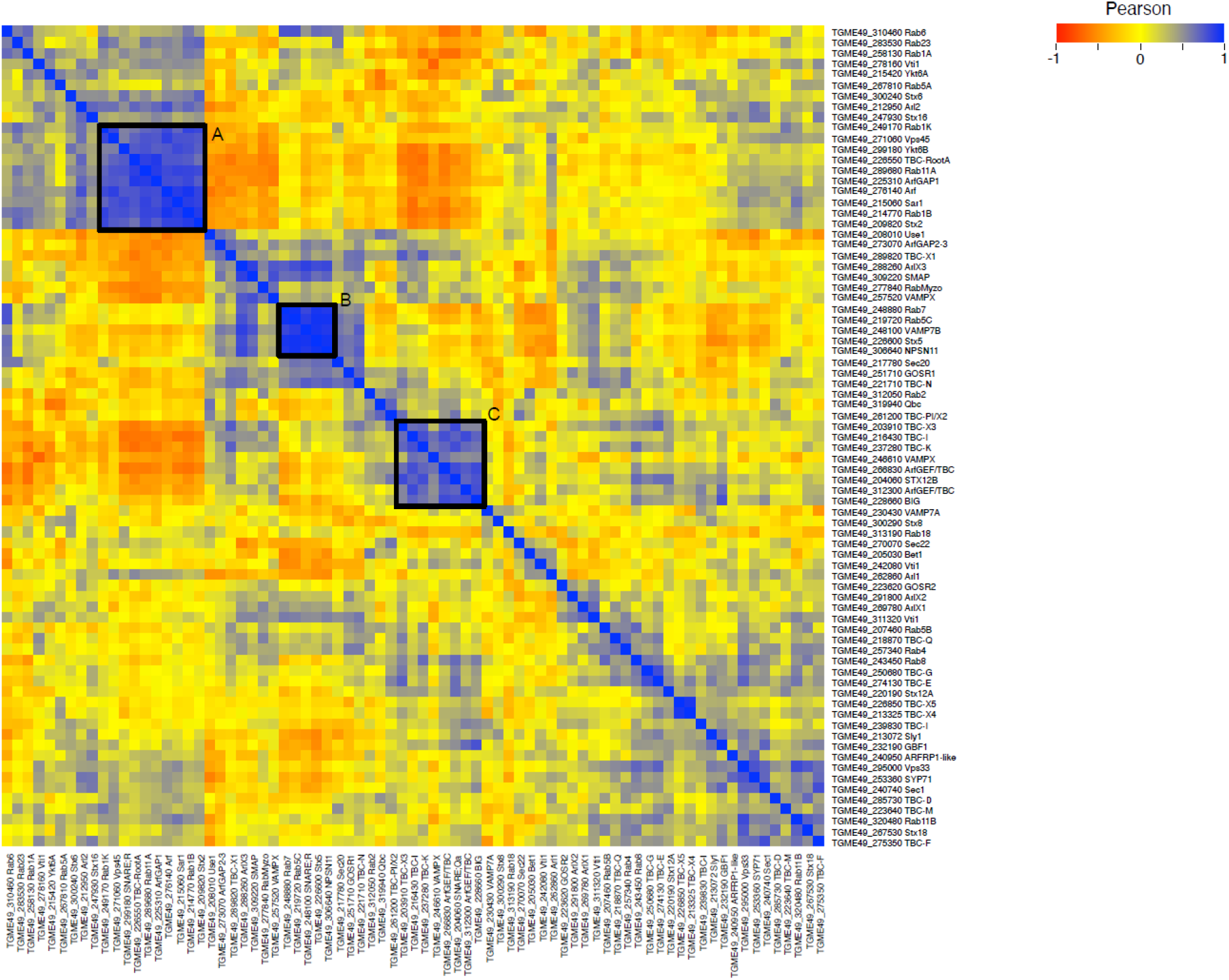
Co-regulation heat map of LSP genes and their known paralogs. Heat map showing Pearson correlation coefficients between gene expression profiles obtained from publicly available RNA-Seq data from various *T. gondii* developmental stages. Correlations are shown for LSP genes and their known paralogs, and colors correspond to levels of correlation as indicated to the right of the figure. Three clusters of co-regulated genes (A, B, and C) are outlined in black and described in the main text.

The first block (Figure 2-Block A) comprises genes most known in other systems and *T. gondii* to be involved in Golgi or endosome to TGN trafficking (29, 34, 35). Notably, in the LSP and MIC/ROP comparison (Figure S4-Block 1), Rab1K was also seen to be co-regulated with several ROP proteins. A second small, co-regulated block (Figure 2-Block B) was identified, whose members suggested involvement in secretory or post-Golgi trafficking pathways. A final co-regulated block (Figure 2-Block c, Table 2) was observed, which included several LSPs. Based on either direct characterization in *T. gondii* or relationships to known paralogs, the genes in this block are inferred to be involved in endolysosomal trafficking. Included in this cluster are the LSP of the endosomal Qa-SNARE Stx12B and both TBS proteins, fusions of cytohesin and TBC-N, both with endosomal trafficking functions in other systems. Notably, all three of these proteins are also found co-regulated with a large cluster of MIC and ROP proteins (Figure S4-Block2), raising clear associations for further study.

### Molecular characterization of Arl LSPs in Toxoplasma gondii

Molecular characterization of some SNARE and Rab proteins has been performed in *Toxoplasma* (35–37). However, to date there has been no systematic analysis of Arls in Apicomplexa. Therefore, we sought to characterize these three Arl LSPs using the model apicomplexan *T. gondii*. Using LIC tagging, we created cell lines expressing each Arl LSP C-terminally tagged with a 3x-HA epitope tag (Figures 1F, S5, S6,).

ArlX1 localization varied but inevitably appeared as a single punctum at the extreme apical end of intracellular tachyzoites, as well as dotted throughout the cell periphery. Signal was frequently observed in the basal body and throughout the intravacuolar network (Figure S6A). Given the potential relationship of ArlX1 with Arl16, and its recent implication in flagellar biogenesis, this could indicate a role for ArlX1 in trafficking to the basal body and/or apical complex (38). ArlX2 had no clear localization pattern, appearing as a series of punctate dots throughout the cell. Lastly, ArlX3 appeared concentrated apical to the nucleus, in the region known to be occupied by the Golgi (39). To further explore ArlX3 localization, we transiently transfected plasmids encoding fluorescently tagged markers: P30-GFP-HDEL, which labels the ER(40), and ERD-GFP (cis-Golgi), GRASP-RFP (cis-Golgi), and GalNAc-YFP (trans-Golgi), which primarily label the Golgi(39, 41–43). There was little overlap with the ER marker, but varying degrees of overlap with each of the three Golgi markers (Figure S6B), suggesting that ArlX3 localizes to the trans Golgi.

In order to rapidly identify candidates for further examination, we targeted each LSP using an inducible CRISPR-Cas9 system and examined the localization of example cargo proteins (Supplementary text, Figure S7). Although ArlX3 appeared to have the most significant impact, we could not rule out the possibility of ArlX1 or ArlX2 proteins having an appreciable impact on asexual growth. Hence, we carried out disruptions of both ArlX1 and ArlX2 (Supplementary text, Figure S8). Overall, neither ArlX1 or ArlX2 disruption had a significant impact on the lytic cycle or the growth of asexual stage *T. gondii* (Figure S8).

### Creation of an inducible ArlX3 knockdown line

As our previous attempts to create a straight knockout line for ArlX3 failed, we attempted an inducible knockdown approach using the TATi system(44). To this end, we constructed a TetO7-myc-pSag1-ArlX3 line, hereafter referred to as ArlX3-iKD, in which ArlX3 transcription is switched off by the addition of anhydrotetracycline (ATc; Figure S9). Quantitative IFA and western blot analysis demonstrate tight ATc-mediated regulation of ArlX3, with significant downregulation as soon as 6 hours and only background expression levels 24 hours post induction (Figures 3A-C, S9D).

**Figure 3.**
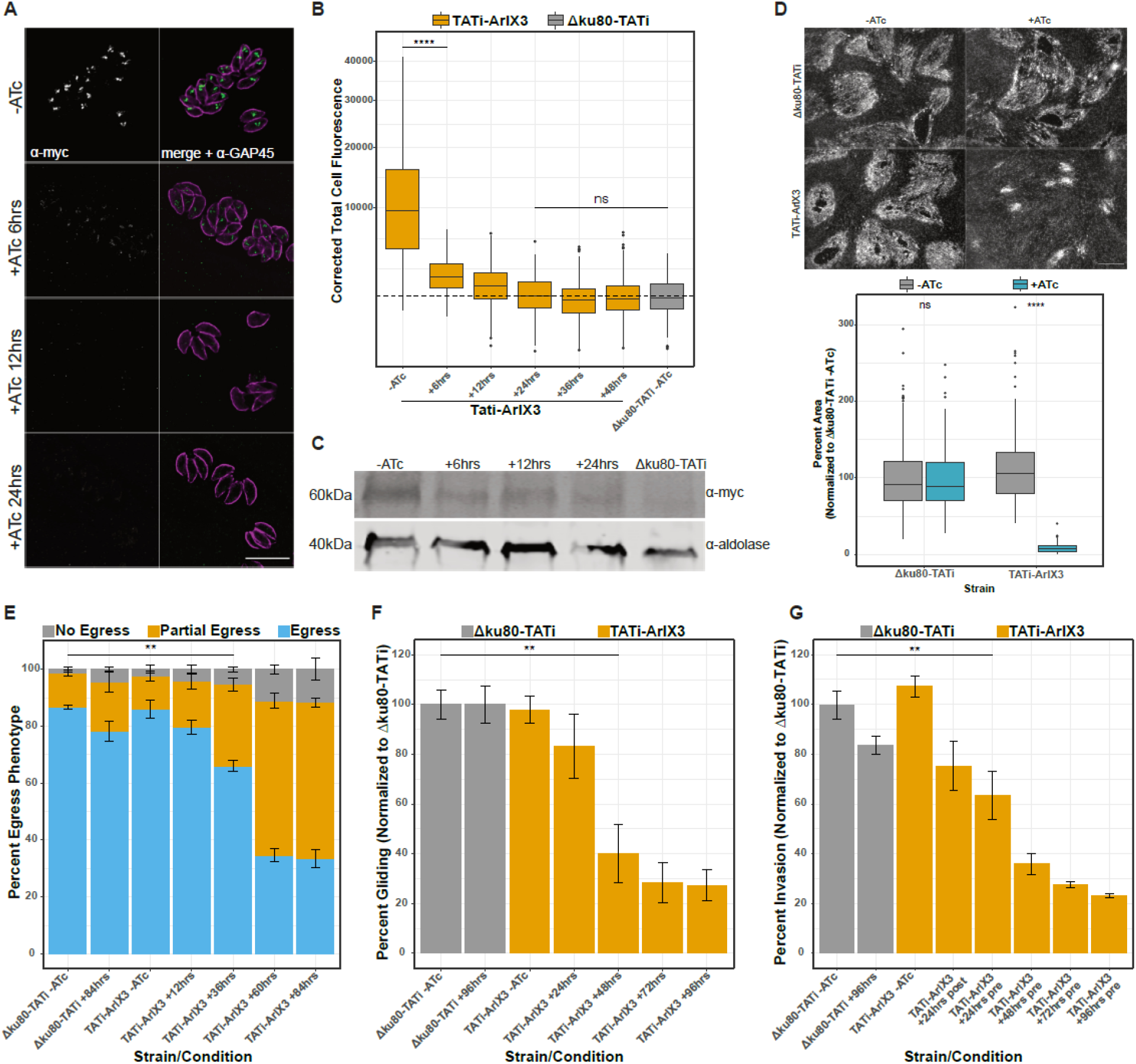
ArlX3-iKD impairs each step of the lytic cycle. This figure summarizes the effects of TgArlX3-iKD on the *T. gondii* lytic cycle. A) ATc induction causes a time-dependent decrease in ArlX3 signal. Scale bar is 10 μm. B) Background-normalized quantification of ArlX3 levels under ATc induction; dotted line represents background signal. Note that signal intensity by 24 hrs post-induction is not significantly different from the parental (Δku80-TATi) line. In this, and all subsequent bar graphs, the first instance of significant difference from controls is indicated. ns: non-significant differences, **** = *p*<0.0001 C) Western-bot depicting downregulation of ArlX3 expression in parasites induced with ATc for the indicated times. Aldolase was employed as loading control. Longer time points are shown in Figure S9D D) Plaque assay demonstrating the marked impact of ATc on TgArlX3-iKD parasite growth, with quantification shown below. E-G) TgArlX3-iKD impairs parasite egress (E), gliding (F), and invasion (G). In each case, the addition of ATc results in no significant difference in parental parasites, but causes a time-dependent decrease in the ability of TgArlX3-iKD parasites to complete each lytic cycle step. ** = *p*<0.01

As an initial characterization of our ArlX3-iKD line, we performed plaque assays, which assess the fitness of the parasite to carry out the lytic cycle and form clearing zones on HFF monolayers. While the parental parasites grew indistinguishably in the presence or absence of ATc, growth was almost completely abrogated in the presence of ATc (Figure 3D). We carried out further assays to assess the ability of ArlX3-iKD parasites to egress, glide, and invade in the presence/absence of ATc. While parental parasites were unaffected by the presence of ATc, ArlX3-iKD parasites show a time-dependent decrease in their ability to carry out each of these lytic cycle steps (Figure 3E-G).

### ArlX3 knockdown results in mislocalization of microneme and rhoptry cargo

As ArlX3-iKD impaired parasites in each step of the lytic cycle (Figure 3E-G), we examined the apical secretory organelles, i.e. the micronemes and rhoptries, which play key roles in egress, gliding, and invasion(45–48).

While parental parasites showed almost exclusively apical localization for each of the four microneme cargo proteins studied in the presence/absence of ATc, ArlX3-iKD parasites showed a time-dependent increase in cargo mislocalization (Figures 4A-E, S9E). We noted three distinct patterns for mislocalized cargo: “vesicular”, with punctate signal throughout the cell, “apical”, with a clear punctum at the apical tip of the cell, and “basal body/extracellular (BB/E)”, with signal concentrated within the basal body or diffusely in the parasitophorous vacuole, or both (Figure 4A). Interestingly, each cargo protein appeared to mislocalize in a distinct manner. AMA1 adopted a mainly apical signal (Figure 4B), M2AP mainly vesicular (Figure 4C), MIC3 overwhelmingly BB/E (Figure 4D), and MIC4 showed an approximately equal mix of all three patterns (Figure 4E). In each case though, mis-localization was moderate following 24 hours of induction, increasing sharply with 48 hours induction and then a gradual increase through 96 hours.

**Figure 4.**
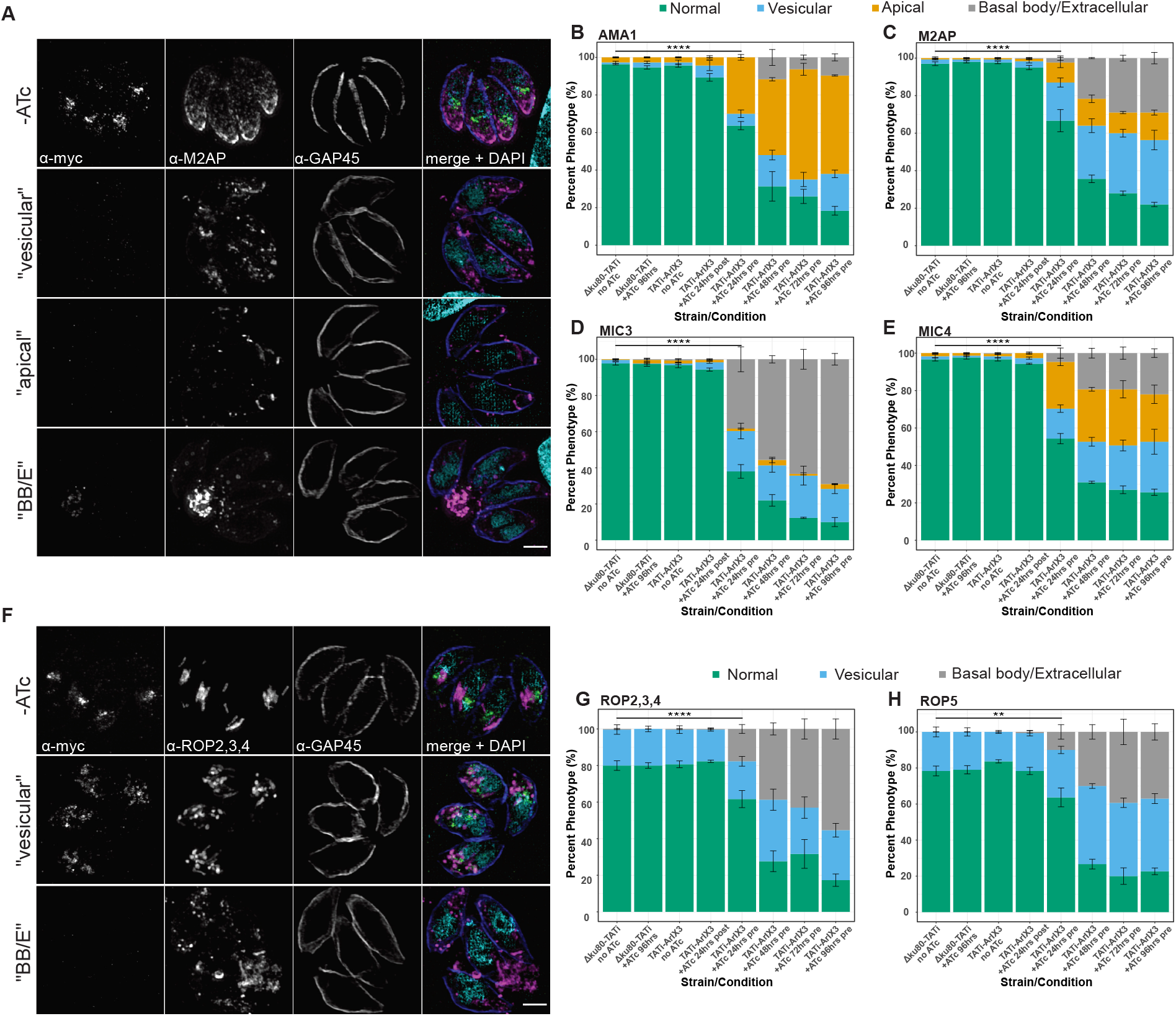
ArlX3-iKD results in mis-localization of microneme and rhoptry cargo. This figure shows the effect of TgArlX3-iKD on apical secretory organelles. A) In the absence of ATc, microneme protein staining adopts a characteristic apical pattern (top row). However, in the presence of ATc, the staining adopts one of three general patterns: scattered in puncta throughout the cell (“vesicular”), a single prominent dot at the apical tip (“apical”), or concentration in the basal body, parasitophorus vacuole, or beyond (“BB/E”). B-E) Quantification of staining patterns for the microneme proteins AMA1 (B), M2AP (C), MIC3 (D), AND MIC4 (E). Note the preference for apical and BB/E staining with AMA1 and MIC3, respectively. F) Staining patterns for rhoptry proteins, as in A; note the absence of apical staining. G, H) Quantification of staining patterns for the rhoptry proteins ROP2,3,4 (polyclonal) and ROP5. ** = *p*<0.01, **** = *p*<0.0001.

We observed a similar pattern with rhoptry markers (Figures 4F-H, S9E). Between ROP2/3/4 and ROP5, there was no clear preference for one pattern over another, and we observed similar temporal dynamics upon ArlX3 knockdown as for microneme cargoes (Figure 4G,H).

### ArlX3 knockdown has a delayed effect on the apicoplast

We also studied the effect of ArlX3-iKD on the endosymbiotic organelles, the mitochondrion and apicoplast (Figure 5). While there was no clear effect on the mitochondrion, there was a time-dependent increase in the mis-localization of the apicoplast protein CPN60, albeit delayed in comparison to the effect on microneme/rhoptry cargo (Figure 5A-C). To confirm this result, we carried out import/processing assays, as CPN60 is proteolytically processed during apicoplast import (49). While the ratio of processed to unprocessed protein was indistinguishable at 24- and 48-hours induction, it showed a marked increase at 72 hours, mirroring the localization data (Figure 5D, E).

**Figure 5.**
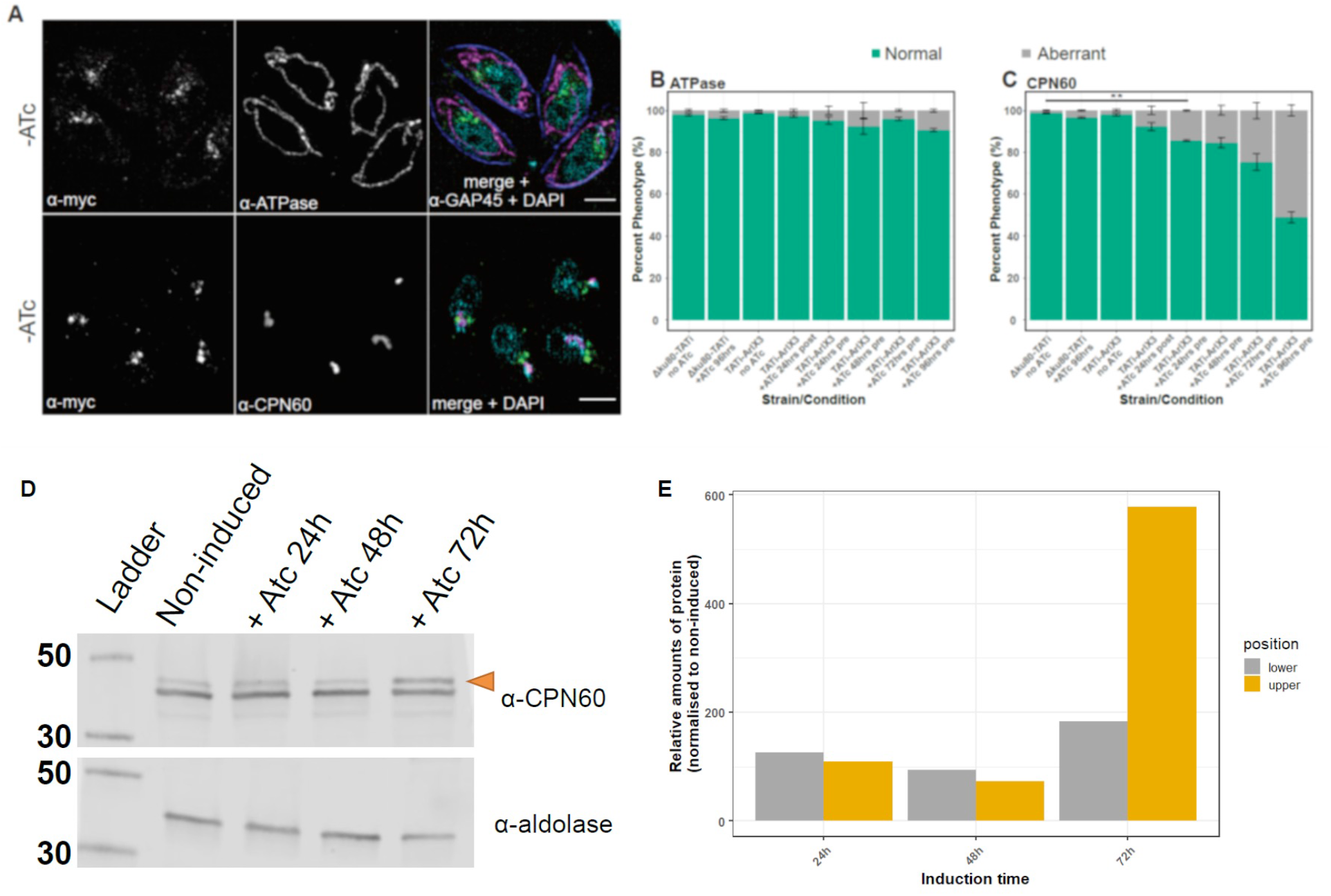
Disruption of post-Golgi trafficking in the ArlX3-KD alters the localization and import of an apicoplast resident protein. This figure shows the effect of TgArlX3-iKD on the *T. gondii* endosymbiotic organelles through an exemplar resident protein. A) Example images of “normal” localization for each marker. For the mitochondrion (ATPase), the classic “lasso” is depicted, although other morphologies have also been described in WT parasites. The apicoplast (CPN60) is typically a single punctum, which can be elongated in dividing parasites. B, C) Quantification of mitochondrion (B) and apicoplast (C) staining. While there was little effect on the mitochondrion, the apicoplast marker CPN60 became diffuse with longer induction times. ** = *p*<0.01. D) Western-blot of induced and non-induced parasites to see processing of the apicoplast resident protein CPN60. Orange arrow mark the preprocessed form of CPN60. Aldolase was used as loading control. E) Quantification of pre-CPN60 and CPN60 bands relative to the loading control in D. Values were normalised to the non-induced parasites.

### ArlX3 knockdown affects the Golgi and early secretory system

As ArlX3 is localized to the Golgi (Figure S6B) and ArlX3-iKD results in cargo trafficking defects (Figures 4, 5), we investigated the role of ArlX3-iKD on the Golgi itself. We first confirmed the close apposition of ArlX3 with the Golgi markers GRASP-RFP (cis-Golgi) and GalNAc-YFP (trans-Golgi), noting that the signal overlap appeared more dramatic with the latter (Figures S10A, B). While ArlX3-iKD had no apparent effect on GRASP-RFP localization (Figure S10A), by 48 hours post-induction the GalNAc-YFP signal appeared more diffuse (Figure S10B). However, the nature of the transient expression made these observations inconclusive.

To further study a possible effect, we tagged the cargo protein sortilin (SORTLR), which is known to localize to the *trans*-Golgi and early endosome-like compartment (ELC)(50), with a C-terminal YFP (Figure 6). As expected, in non-induced parasites ArlX3 signal overlaps with that of YFP (Pearson correlation *r* = 0.77 ± 0.07), confirming that ArlX3 primarily localizes at the trans-Golgi (Figure 6A). However, upon induction of ArlX3 knockdown, SORTLR signal ceased to be restricted to the Golgi-ELC area, appearing fragmented by 48 h post induction with ATc (Figures 6A-C).

**Figure 6.**
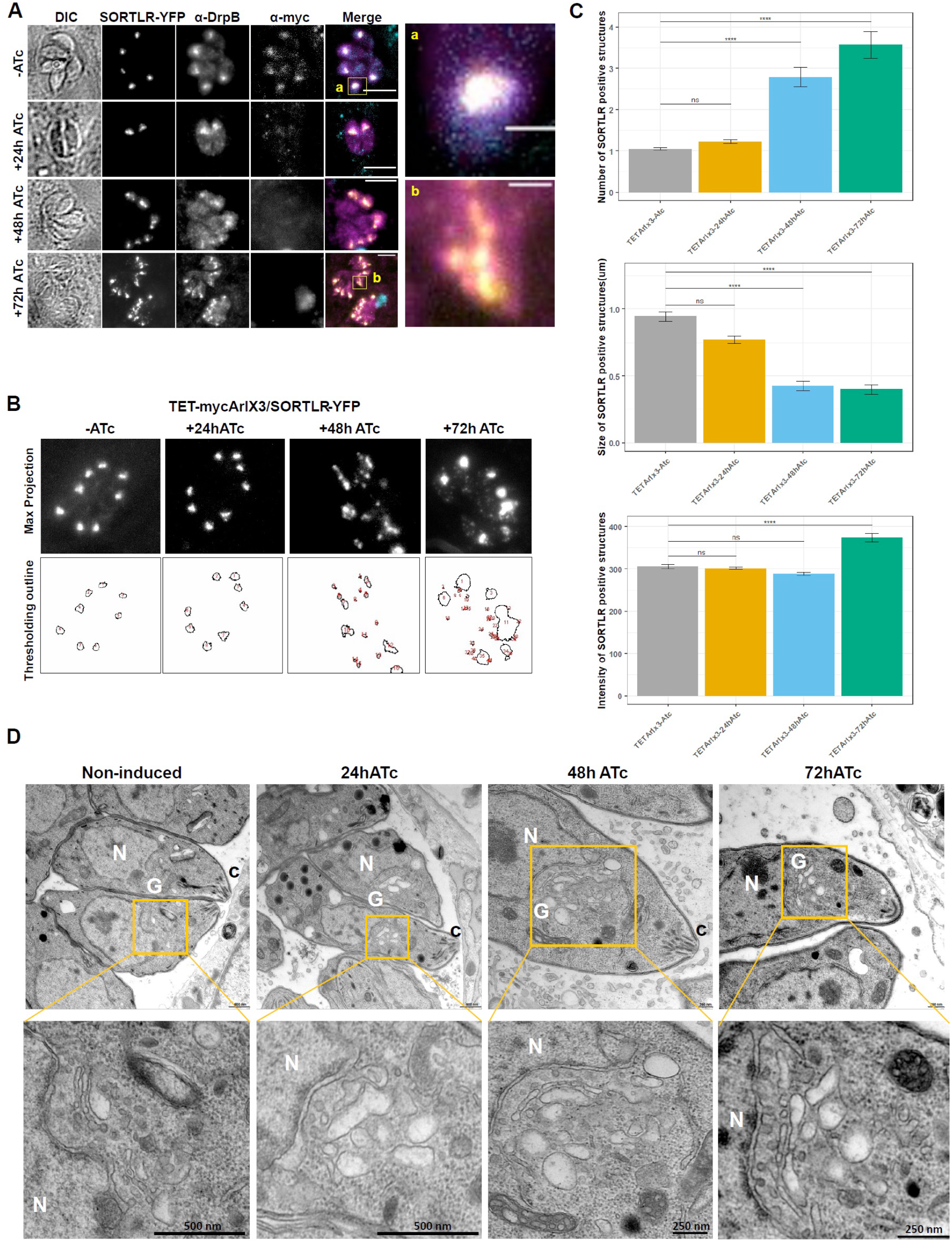
ArlX3-iKD impacts the organization of the Golgi and ELC. This figure shows the effect of the absence of ArlX3 in Golgi and ELC stability. A) Images depicting ArlX3-iKD-SORTLR-YFP parasites ± ATc for the indicated time-points and stained with α-myc and α-DrpB antibodies. Both Sortilin and DrpB are markers for Golgi. Insets shows the merge image of all 3 channels where there is colocalization of ArlX3 with the Golgi markers in non-induce parasites in a defined area that is the Golgi. After 72h of induction, the Golgi seemed to fragment and several vesicles were observed. B, C) Analysis of the vesicles with SORTLR signal. B) Representative images of the SORTLR channel and the outline of the threshold applied with FIJi. C) Bar graphs showing the quantification of vesicles, their size and the average intensity of vesicles analysed. Data shown as Mean ± SEM. D) Representative EM images of parasites induced with ATc. Insets show a closer image of the Golgi area. N: nucleus; G: Golgi.

These results are confirmed by transmission electron micrographs on ArlX3-iKD parasites in the presence/absence of ATc, which show a lack of recognizable stacked Golgi in the presence of ATc (Figure 6D).

## Discussion

In this study, we have confirmed the presence of 18 LSPs in Apicomplexa and related taxa, and explored the role of one LSP, ArlX3, in membrane-trafficking in *T. gondii*. These results suggest a previously unexplored wealth of unique trafficking factors in a lineage of important eukaryotic parasites, with implications for understanding apicomplexan cell biology and eukaryotic evolution.

Although some LSPs, such as Rabs 1A, 5B, 5C, and 11B and the TBS proteins have been reported previously(27–30), their unique phylogenetic relationship compared to other trafficking factors were not fully explored. We report here the presence of multiple additional LSPs, including examples from the Arl, Rab, SNARE, and TBC families (Figures 1B, S1). Through examination of co-regulation in transcriptomic datasets, we identified several relationships with proteins either previously studied in the Toxoplasma system or with orthologs of known function in opisthokont models indicative of these LSPs involved in endosomal/post-Golgi trafficking. Moreover, through endogenous tagging, we further demonstrate that at least three LSPs, ArlX1-3, are expressed in asexual *T. gondii* tachyzoites (Figures 1F, S5, S6), suggesting that other LSPs are also likely to be translated into mature proteins.

Most saliently, we have shown that ArlX3 plays an important role in *T. gondii* tachyzoites, as ArlX3-iKD abrogates parasite growth (Figure 3C) and impairs each step of the lytic cycle (Figure 3E-G). In agreement with these results, ArlX3-iKD severely impairs proper localization of multiple microneme and rhoptry cargo proteins (Figures 4, S9). Although some previous studies have noted the existence of alternative fates for mislocalized cargoes(51, 52), quantification of mislocalization frequently follows a binary approach (i.e., (mis)localized). In terms of micronemes, the vesicular staining may result from a failure to properly traffic microneme protein-containing vesicles, a failure of microneme recycling following endodyogeny, or some other defect. Similarly, the basal body staining may represent a recycling failure, while extracellular staining likely represents aberrant microneme protein inclusion in the “default constitutive” dense granule secretion pathway(53, 54). The apical signal is enigmatic; it was previously proposed to represent a novel trafficking pathway to a subgroup of apical micronemes(51, 55). However, this is inconsistent with the observation that TgSORTLR, responsible for forward translocation of microneme and rhoptry cargo from the TGN, also results in apical staining (see for example Figure 4, Sloves *et al*. 2012)(50). Although some vesicular rhoptry marker staining is always observed (Figure 4F-H), likely representing nascent rhoptries forming from a Rab5A-positive compartment(52), the ratio is increased in ArlX3-iKD parasites, suggesting a defect in trafficking, fusion, or both.

Compared with Rab5A/C, in which only some microneme cargo proteins were mislocalized with the overexpression of a dominant negative (DN) form of either Rab(29), all four cargo proteins tested here were mis-localized with ArlX3-iKD (Figure 4B-E). This included proteins that had a normal localization with DN Rab5A/C, such as AMA1 and M2AP, as well as a protein that was mis-localized (MIC3; MIC4 was not investigated). This may indicate that ArlX3 acts upstream of Rab5A/C in forward secretory trafficking, that it functions in multiple pathways, or some other explanation. The blanket effect of both Rab5A/C and ArlX3 disruption on rhoptry cargo localization supports distinct pathways for microneme and rhoptry biogenesis.

Our quantification of microneme marker mislocalization (Figure 4B-E) suggests that these proteins are differentially capable of unassisted forward trafficking and of entering alternate trafficking pathways. Several microneme cargoes contain a propeptide that is cleaved during trafficking, likely in the ELC(56). However, when these propeptides are removed through genetic modification, microneme protein trafficking arrests in different compartments: TgM2APΔpro arrests in the ELC(43), TgMIC3Δpro in the basal body/PV(57), and TgMIC5Δpro in the early secretory pathway including the ER(58). Of particular interest, the localization for M2AP (Figure 4C) and MIC3 (Figure 4D) upon ArlX3-iKD appear similar to the Δpro lines; MIC5 was not included here and so no comparisons can be drawn. These observations are consistent with the notion that blocking forward translocation of microneme proteins prior to the post-Golgi removal of their propeptides results in differential mis-localization, presumably by inclusion of the accumulating protein in non-physiological trafficking pathways. It also supports the idea that ArlX3 is involved in the forward trafficking of diverse microneme proteins.

The apicoplast is unique among red plastids in that it is present outside the ER limiting membrane and has four distinct membranes(19). This has spawned numerous questions around how resident proteins are trafficked to the apicoplast; the three main contending hypotheses include transient direct contacts with the ER, direct ER-apicoplast trafficking, and post-Golgi trafficking(21). Previous studies have alternatively favoured a direct pathway from the ER(59, 60), or a post-Golgi route(61). Though not explicitly designed to investigate this question, our localization and protein import studies with ArlX3-iKD could be consistent with the possibility of a post-Golgi trafficking route, at least for CPN60 (Figure 5). However, as the effect was delayed compared to apical organelles, it is also consistent with a knock-on effect, caused by disruption of the Golgi and therefore a general (delayed) traffic defect to other organelles (Figure 4). Future studies should further investigate the possible role of ArlX3, and other Golgi-localized trafficking factors, in apicoplast trafficking.

Although our study suggests a role for ArlX3 in post-Golgi trafficking in apicomplexans, the exact function of ArlX3 remains unclear. This is in part hampered by the lack of a clear origin for ArlX3 (Figures 1, S3, Supplementary text). In cases where an apicomplexan LSP has a clear origin, for example both Rab5C and 11B, the function has remained similar to that of its pan-eukaryotic paralog(28, 29). In addition, Arl biology remains comparatively understudied, with diverse functions in membrane trafficking, cytoskeletal organization, and ciliogenesis/intraflagellar transport posited for various family members(32). The notion that ArlX3-iKD impairs trafficking is suggested by the extensive mis-localization of microneme and rhoptry proteins (Figures 4, 5). In addition, our observation of a more general disruption on Golgi/ELC morphology (Figures 6, S10) bears striking similarity to disruption of another TGN resident trafficking protein in *T. gondii*, Stx6 (62). We have recently published an extensive model for trafficking in *T. gondii* (17); here, we suggest the possible inclusion of ArlX3 (Figure 7). Future studies should elaborate the precise role of ArlX3 in the apicomplexan MTS.

**Figure 7.**
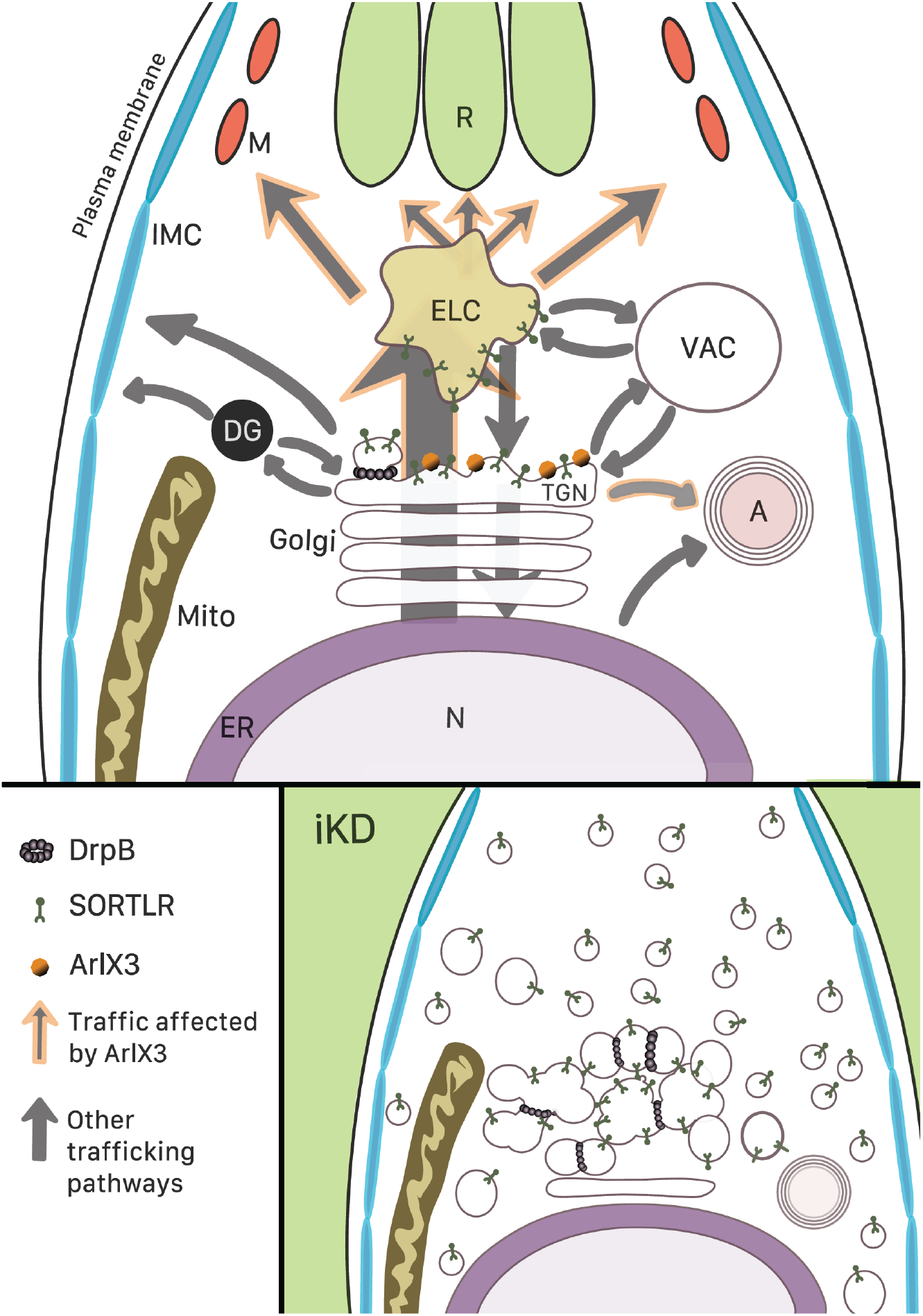
A comprehensive model of *T. gondii* membrane trafficking. This figure outlines a current view of *T. gondii* membrane trafficking, including insights from this study regarding the localization and possible function of ArlX3. In the main panel, highlighted arrows show traffic routes towards micronemes and rhoptries where ArlX3 is involved. In the lower panel, depletion of ArlX3-KD (iKD) leads to the disruption of the trans-Golgi. M: micronemes, R: rhoptries, IMC: inner membrane complex, ELC: endosomal-like compartment, VAC: plant-like vacuole, DG: dense granules, TGN: trans-Golgi network, A: apicoplast, Mito: mitochondria, ER: endoplasmic reticulum, N: nucleus.

One larger question remains regarding the evolutionary cell biological implications of the LSPs themselves. We have previously argued that the OPH, which was conceived to explain the proliferation of distinct organelles during the FECA-LECA transition, is just as relevant to modern-day eukaryotes(12). In this view, continued paralogous duplication and diversification of trafficking factors would allow for the emergence of additional unique organelles in eukaryotic lineages. The expectation under this model is therefore that the LSP should be taxonomically restricted to only those lineages in which the unique structure or organelle is present. This is supported by previous studies of some of the LSPs identified here. Rab5C, which we confirm is present across the Myzozoa (Table S1, Figures 1B, S1, S2Q), is involved in trafficking to apical secretory organelles(29), which are conserved across this group(20). Rab11B, which we confirm is conserved across alveolates (Table S1, Figures 1B, S1, S2R), is involved in trafficking to the IMC in *T. gondii* (28), which is homologous to the alveoli of all alveolates(21). Notably, our results regarding ArlX3 also follow this pattern. ArlX3 is restricted to myzozoans (Table S1, Figures 1B, S1,S3) and, like Rab5C, plays a role in proper localization of microneme and rhoptry cargoes (Figure 4). Hence, our results lend weight to the idea that the OPH continues to operate in extant eukaryotes, giving rise to unique organelles not conserved across eukaryotic diversity.

In this study, we have confirmed the presence of 18 LSPs in Apicomplexa and their close relatives and showed that one such LSP, ArlX3, plays an important role in asexual *T. gondii* tachyzoites. These results have important implications not only for apicomplexan cell biology but for understanding the conservation and emergence of eukaryotic trafficking factors. Future studies should investigate the correlation between phylogenetic distribution and function of trafficking factors more closely. Our results provide a list of potential targets for those working in apicomplexan cell models and, on a more fundamental level, a general template for uncovering and exploring novel cell biology applicable to any eukaryote.

## Materials and Methods

### Homology searching and phylogenetic analysis

Predicted proteomes of all organisms under study were downloaded from relevant public databases; information regarding all datasets is found in Table S3. A full description of the methodoloy can be found in the Supplementary Text. Briefly, homology detection involved HMMer v3.1b1 (63) followed by reciprocal BLASTp (v2.8.1) (64) searches with an e-value cut off of 0.05.

Sequences were aligned with MAFFT v7.407 (66)□ and subjected to maximum likelihood analysis with IQ-TREE v1.6.11 (67) and RAxML v8.2.12 (68) or Bayesian analysis using MrBayes v3.2.7a (70).

### Protein structural prediction

*De novo* structural prediction used RoseTTAFold (71) hosted at https://robetta.bakerlab.org/. Starting sequences were searched against the UniRef30 (2022) database using HHblits (72) as hosted on the MPI Bioinformatics Toolkit webserver (73) with default parameters. The full a3m alignment from HHblits was used together with the starting sequence as inputs to RoseTTAFold. Human structures for comparison were retrieved directly from the AlphaFold protein structure database (74, 75). All protein structures were visualized using ChimeraX (76) and coloured using the default “rainbow” colouration.

### Analysis of Gene Co-expression

RNA-Seq gene expression data sets were collected from ToxoDB as previously described(77). Pearson correlation coefficients between genes were calculated from the FPKM values in R (78) and plotted using the gplots package (v3.1.1).

### Parasite and host cell culture

Human foreskin fibroblast (HFF; ATCC^®^ designation SCRC-1041 ™) cells were grown in Dulbecco’s modified Eagle’s medium (DMEM; Sigma, D6546) supplemented with 10% foetal bovine serum (FBS; BioSell FBS.US.0500), 2 mM L-glutamine (Sigma, G7513), and 25 mg/mL gentamicin (Sigma G1397). *T. gondii* strain RH parasites were cultured on confluent HFF monolayers in the same supplemented media. All cells were maintained at 37°C and 5% CO2.

### Genomic DNA Isolation, cloning, and PCR

To isolate genomic DNA from parasites, roughly 1×10^6^ fully egressed parasites were collected and then gDNA was isolated using Qiagen DNeasy Blood and Tissue Kit (Cat. No. 69504) or Extractme genomic DNA kit (Blirt, EM13), as per the manufacturer’s instructions. Amplification of DNA segments for cloning used Q5^®^ high-fidelity DNA polymerase (NEB) whereas diagnostic PCR used standard *Taq* DNA polymerase (NEB).

All restriction enzymes for cloning were from NEB, using the high-fidelity (HF) versions, where available. Plasmid preps were made using Qiagen QIAprep Spin Miniprep Kit or Extractme Plasmid Mini Kit (Blirt, EMO1.1), as per the manufacturer’s instructions. All primers used in this study for cloning and diagnostic PCR confirmation of stably transfected cell lines are provided in Table S4. Details on the generation of tagged lines for each Arl LSP using ligation-independent cloning (LIC) (79) are described in Supplementary text.

CRISPR/CAS9 modification of parasites used a single vector encoding both CAS9-NLS-YFP enzyme and pTgU6-gRNA (79)(Supplementary text).

Repair template for integration of a tag in the *sortlr* locus was generated as previously described (80).The repair template was purified using a PCR purification kit (Blirt; EM26.1).

### Transfection of parasites

Transfections were carried out using an AMAXA 4DNucleofector ™ (Lonza; AAF-1003X) and the P3 primary cells kit (Lonza; V4XP-3024) as previously described (81) (see Supplementary text for further details).

Generation of ArlX3-iKD was achieved by integration of a PCR fragment containing the p*dhfr*-*hxgprt*-3’UTR*dhfr*-TetO7-myc-pSag1 cassettes with 50 nucleotides (nt) of homology to the promoter of ArlX3 and 50nt of homology to the n-terminus of ArlX3 in each side of the amplicon. Parasites were transfected with both donor DNA and a CAS9-YFP vector a gRNA targeting the N-terminus of this gene. Parasites were selected with 78μM mycophenolic acid (MPA; Sigma; M3536) and 230μM xanthine (Xan. Sigma; X3627)(82); selected pools were then cloned by limiting dilution in 96 well plates and individual clones picked and analysed.

Tagging of SORTLR with the yellow fluorescent protein (YFP) was achieved by simultaneous transfection of a specific CAS9-YFP-gRNA vector with a PCR amplicon as donor containing 50 nt of homology to *sortlr* in each side of the YFP cassette. Parasites expressing Cas9-YFP were sorted via FACS (FACSARIA III, BD Biosciences) into 96-well plates (a minimum of 3 events per well). Resultant clonal parasites were screened by PCR and repair template integration confirmed by sequencing(81).

### Immunofluorescence assays

HFF cell confluent monolayers on glass coverslips were infected with parasites. Parasites were fixed using 4% paraformaldehyde (PFA) at room temperature for 20 minutes before being washed three times with 1X phosphate-buffered saline (PBS). Subsequently, coverslips were permeabilized and blocked using blocking buffer (3% BSA and 0.2% Triton X-100 in 1X PBS (PBS-TX-100)) for one hour at room temperature. Primary antibodies were added to blocking buffer at the dilutions indicated in Table S4, and cells stained for one hour at room temperature before being washed three times with PBS-TX-100. Similarly, secondary antibodies (Life Technologies) were added to blocking buffer and cells stained for one hour at room temperature in the dark. Samples were washed three more times with PBS-TX-100 and then coverslips were mounted using either mounting media alone or mounting media supplemented with DAPI. Non permeabilizing IFAs were performed as above, but without the addition of TX-100 to blocking and wash buffers.

### Plaque assay

For plaque assays, 1×10^3^ freshly egressed parasites were added to a confluent HFF monolayer with or without addition of 1 μg/mL ATc. After five days, cultures were washed once with PBS and then fixed with ice-cold methanol for 20 minutes. Methanol was removed and cells stained with Giemsa, followed by three washes with PBS. All plaques in 10 random fields of view were measured using Fiji for three independent experiments. Raw data are provided in Source data.

### Gliding assay

1×10^6^ freshly egressed parasites were suspended in pre-warmed gliding buffer (1mM EDTA and 100mM HEPES) and allowed to glide on glass coverslips coated with FBS for 30 minutes prior to fixation with 3% PFA. An IFA was performed using α-SAG1 primary antibody under non-permeabilising conditions to label deposited trails. One hundred random parasites were assessed for the presence/absence of trails in three independent experiments and the mean and SEM were calculated. Raw data are provided in Source data.

### Invasion-replication assay

5×10^4^ freshly egressed parasites were allowed to invade confluent HFFs pre-seeded onto coverslips for one hour before several washes with PBS were performed to remove uninvaded parasites. 24 hours later, cells were fixed with 4% PFA and an IFA was done with α-GAP45 primary antibody. For the invasion assays, the number of vacuoles in 15 random 40X fields of view were counted for three independent experiments, and the mean and SEM calculated. For the replication assays, the number of parasites within 100 random vacuoles were counted in 3 biologically independent experiments. Raw data are provided in Source data.

### Egress assay

For egress assays, 5×10^4^ freshly egressed parasites were allowed to invade confluent HFFs for an hour. 36 hours later, culture media was exchanged for pre-warmed DMEM without FBS with 2 μM calcium ionophore (A23187) to induce egress. Five minutes after media exchange, cells were fixed with 3% PFA and subsequently stained with α-SAG1 antibody under non-permeabilising conditions (together with DAPI to assess intracellular vacuoles). One hundred random vacuoles were assessed for egress ability in three independent experiments, and the mean and SEM calculated. Raw data are provided in Source Data.

### Western blotting and protein detection

Approximately 1×10^7^ freshly egressed parasites were harvested and pelleted by centrifugation at 5000 rpm followed by a single wash with 1X PBS. The parasite pellet was lysed on ice with NP-40 buffer (50 mM Tris-HCl pH 7.5, 150 mM NaCl, 1% Nonident P-40, and 4mM EDTA) and incubated for five minutes on ice. Insoluble material was pelleted by centrifugation at 14,000 rpm at 4°C. The supernatant was placed in a new tube together with 10X NuPage™ Sample Reducing Agent (Invitrogen™) and 4X loading buffer (125mM Tris-HCl pH 6.5, 50% v/v glycerol, 4% w/v SDS, 0.2% w/v orange G). Samples were boiled for 10 minutes at 95°C, loaded onto 12% Mini-Protean^®^ TGX™ Precast polyacrylamide gels (BioRad), and run at 130V. Samples were transferred to nitrocellulose using a Mini-Protean^®^ transfer tank containing 1L of transfer buffer (48mM Tris, 39mM glycine, and 20% methanol) running at 400mA for one hour. Membranes were blocked using 5% skim milk powder in 1X PBS at room temperature for one hour. Primary antibodies were added at the appropriate concentration (Table S4) in blocking buffer (5% skim milk powder in 1X PBS + 0.2% Tween-20 (PBS-TW-20) for one hour. Membranes were washed three times with PBSTW-20, before addition of IRDye680RD and IRDye800RD secondary antibodies (Li-Cor) in blocking buffer for a further hour. Membranes were washed three times in PBS-TW-20 followed by an additional wash in 1X PBS to remove Tween-20 prior to imaging. Detection of infrared signal was performed using a Li-Cor Odyssey with Image Studio 5.0 software (Li-Cor).

### Structured illumination microscopy (SIM)

For SIM imaging we used an ELYRA PS.1 microscope (Zeiss) equipped with a Plan Apochromat 63x, 1.4 NA oil immersion lens and CoolSNAP HQ camera (Photometrics). SIM processing of captured images used ZEN Black software (Zeiss) and all subsequent processing used Fiji ImageJ (83).

### Electron Microscopy

Induced and non-induced intracellular parasites were fixed with 2.5% (v/v) glutaraldehyde in 0.1 M phosphate buffer, pH 7.4, after the indicated incubation. The parasites were washed three times at room temperature with PBS (137 mM NaCl2, 2.7 mM KCl, 10 mM Na2HPO4, 1.8 mM KH2PO4, pH 7.4) and postfixed with 1% (w/v) osmium tetroxide for 1 hour. Subsequent to washing with PBS and water, the samples were stained *en bloc* with 1% (w/v) uranyl acetate in 20% (v/v) acetone for 30 minutes. Samples were dehydrated in a series of graded acetone and embedded in Epon 812 resin. Ultrathin sections (thickness: 60 nm) were cut using a diamond knife on a Reichert Ultracut-E ultramicrotome. Sections were mounted on collodium-coated copper grids, post-stained with lead citrate (80 mM, pH 13), and examined with an EM 912 transmission electron microscope (Zeiss, Oberkochen, Germany) equipped with an integrated OMEGA energy filter operated in the zero-loss mode at 80 kV. Images were acquired using a 2k × 2k slow-scan CCD camera (Tröndle Restlichtverstärkersysteme, Moorenweis, Germany).

For cryo-immunolabeling, the samples were fixed in phosphate buffer, pH 7.2, containing 4% freshly prepared formaldehyde. After several washes in the same buffer, they were embedded in 10% gelatin at 37° C for 30 minutes. The material was spun down and the samples were left on ice for 30 minutes. After confirming the gelatine was solid, the pellet was removed from the tubes and infiltrated overnight in 2.1 M sucrose and rapidly frozen by immersion in liquid nitrogen. Cryo-sections (70 nm thick) of the frozen material were obtained at −120°C using an Ultracut cryo-ultramicrotome (Leica Microsystems). The cryo-sections were collected on formvar-coated nickel grids, thawed, and put on a cushion of 2% gelatine. The grids were left for 20 minutes at 37°C and then blocked in PBS containing 3% bovine serum albumin for 1 hour. After this time, they were incubated in the presence of primary antibody. Then they were washed several times in blocking buffer and incubated with 15 nm gold-conjugated Protein A (Aurion). The grids were washed several times in the blocking buffer, dried, and contrasted in a mixture of methylcellulose/uranyl acetate. All images were captured on a Jeol 1200 transmission electron microscope (JEOL, Japan) operating at 80kV and analysed/processed with Fiji software.

### Statistical and image analysis

All statistical analysis was performed in R v3.6.1 and v4.2.1 (78). Comparisons among multiple means used one-way ANOVA followed by post-hoc Tukey’s HSD test when assumptions of normality and equal variance were not significantly violated; in cases where violation did occur, Kruskal-Wallis followed by post-hoc Dunn’s test was used instead. Comparison between plaque sizes in plaque assay used two-way ANOVA and means within each group were compared by a Wilcoxon signed rank test. Comparison of multiple populations within a single group (e.g. for phenotypic analysis) used Chi-square followed by post-hoc Fisher’s exact test. All plots were made in R and the first instance of significant difference from controls indicated.

Quantitative fluorescence analysis were performed in images captured using the same excitation parameters on a Leica DiM8 widefield fluorescence microscope equipped with a HC PL APO 100x/1.44 oil immersion lens (Leica) and C13440-20C CMOS camera (Hamamatsu). They were then analysed with ImageJ Fiji v1.53q as detailed in Supplementary text.

## Acknowledgments

We wish to thank Jennifer Grünert for technical assistance in electron microscopy. C.M.K. was funded by an Alberta Innovates Health Solutions Fulltime Studentship and a Canada Vanier Graduate Scholarship. His research has been funded in part by the generosity of the Stollery Children’s Hospital Foundation and supporters of the Lois Hole Hospital for Women through the Women and Children’s Health Research Institute. AP and TM were funded by a KAUST faculty baseline fund (BAS/ 1/1020-01-01). Research in the Dacks Lab is funded by NSERC Discovery Grants (RES0043758, RES0046091). Research in the Meissner lab is funded by a DFG Programme Grant (ME 2675/6-2).

## Supplementary Figure Legends

Figure S1. MTS machinery across taxa.

This figure provides an overview of MTS machinery across all study taxa. Each row corresponds to a single taxon written in abbreviated form. Each column corresponds to a grouping of related proteins as indicated by the column title. Groups of proteins are represented by pie charts, with each wedge representing a single orthologue as indicated by the legend at the top of each column. Within each row, a filled circle represents the presence of at least one orthologue of the relevant protein in the relevant taxon; multiple gene copies are not indicated but may be deduced by inspecting Table S1. The cladogram to the left of the taxon names indicates known relationships between taxa. Groups are colour-coded for ease of interpretation: red (apicomplexans), cyan (chromerids), green (dinoflagellates), purple (ciliates), brown (stramenopilles), orange (chlorarachniophytes), light blue (haptophytes/cryptomonads), green (archaeplastids).

Figure S2. Phylogenetic analysis of paralogous gene families.

This figure shows the results of phylogenetic analyses across paralogous membrane-trafficking gene families in Apicomplexa and close outgroup taxa. For this and future figures depicting phylogenetic trees, the best Bayesian topology is shown, with bootstrap support for clades derived from RAxML and IQ-Tree mapped to the appropriate node. For larger datasets where Bayesian analysis was not conducted, the best RAxML topology is shown. In all cases, the support for important nodes is denoted. All other node support values are provided as symbols; the figure legend in each case denotes the minimum support across all methods required for the node to carry the relevant symbol. Clades corresponding to specific paralogues are shaded in alternating colours and are labelled using the appropriate abbreviated name. Taxa are colour-coded as in Figure S1. A) Analysis of Qa SNARE homologues. The Qa SNAREs Stx2, 5, 12, 16, and 18 resolve into strongly supported monophyletic clades. Notably, the hematozoan duplication of Stx2 and the clear division into Stx12A and Stx12B paralogues are evident B) Analysis of Stx2 homologues with Stx16 as an outgroup. The Stx2 duplication in hematozoans is labelled. C) Analysis of Stx12 paralogues with Stx16 as an outgroup. Note the clear separation into pan-eukaryotic Stx12A and myzozoan Stx12B paralogues. D) Analysis of Qb SNARE homologues. The Qb SNAREs GOSR1, GOSR2, NPSN11, and Sec20 resolve into strongly supported monophyletic clades. Although the Vti1 clade lacks RAxML bootstrap support, Vti1 homologues are clearly distinct from other Qb SNAREs. E) Analysis of Vti1 homologues with NPSN11 as an outgroup. Note the presence of multiple Vti1 homologues across taxa, including the Apicomplexa, but the overall lack of support for internal Vti1 clade nodes. F) Analysis of Qb SNARE homologues with reduced Vti1 sequences. The Qb SNAREs GOSR1, GOSR2, NPSN11, Sec20, and Vti1 resolve into strongly supported monophyletic clades. Note that, compared to (D), the Vti1 clade corresponding to a smaller number of less divergent sequences (as chosen by scrollsaw) is better supported. G) Analysis of Qc SNARE homologues. The Qc SNAREs Bet1, Stx6, Stx8, SYP71, and Use1 resolve into strongly supported monophyletic clades. H) Analysis of the Qb domain of Qbc SNARE homologues. Although the support for NPSN11 as a clade dropped with the inclusion of the Qb domain of Qbc SNAREs (0.88/28/94), the combined clade of Qbc/NPSN11 was moderately supported (0.9/48/81). I) Analysis of the Qc domain of Qbc SNARE homologues. Note that the Qc domain of Qbc SNAREs does not form a single monophyletic clade, instead grouping between SYP71 and Use1 homologues (0.82/35/99 for the group as a whole). J) Analysis of R SNARE homologues. The R SNAREs Sec22, VAMP7, and Ykt6 resolve into strongly supported monophyletic clades. Note the clear duplications of VAMP7 and Ykt6. K) Analysis of VAMP7 paralogues with Sec22 as an outgroup. There is a clear separation into pan-eukaryotic VAMP7A and myzozoan VAMP7B paralogues. L) Analysis of Ykt6 paralogues with Sec22 as an outgroup. There is a clear separation into pan-eukaryotic Ykt6A and myzozoan Ykt6B paralogues. M) Analysis of Rab homologues. Note that, while bootstrap support for each clade overall is good, some clades fail to reach significance. N) Scrollsaw analysis of Rab homologues. Each clade is represented by the 15 least divergent representatives, based on scrollsaw analysis. Note the similar topology compared to (M) but overall increased support. O) Analysis of Rab1 paralogues with Rab18 as an outgroup. There is a clear separation into pan-eukaryotic Rab1B, SAR-specific Rab1A, and myzozoan Rab1K paralogues. The protein structure is an exemplar predicted structure of TgRab1K, showing the N-terminal β-propeller. P) Analysis of Rab5-related sequences. Note that all clades are at least weakly supported. Q) Analysis of Rab5 paralogues with Rab6 as an outgroup. There is a clear separation into pan-eukaryotic Rab5A, alveolate-specific Rab5B, and myzozoan Rab5C paralogues. R) Analysis of Rab11 paralogues with Rab18 as an outgroup. There is a clear separation into pan-eukaryotic Rab11A and alveolate-specific Rab11B paralogues. The included alignment shows the conserved Rab11B insertion along with its amino acid conservation. S) Analysis of RabX1 and related sequences. Note the relationship between myzozoan RabX1 sequences and RabL2/RTW sequences. T) Analysis of TBC homologues. Note the overall strong support for individual clades as well as larger groupings. U) Scrollsaw analysis of TBC homologues. Each clade is represented by the 15 least divergent representatives, based on scrollsaw analysis. Note the similar topology compared to (T) but overall increased support. V) Analysis of the TBC domain of TBS sequences. Note the clear grouping of TBS sequences apart from the lone E. huxleyi sequence, which groups with other TBC-N sequences (asterisk). W) Analysis of TBC-X2 homologues, demonstrating their relationship with archaeplastid TBC-PI orthologues. X) Analysis of TBC-X homologues apart from TBC-X2. Note the relationship between pan-eukaryotic TBC-Q and alveolate TBC-X1, 3, 4, and 5 clades.

Figure S3. Phylogenetic analysis of ARF/Arl/Sar proteins and their regulators.

This figure shows the results of phylogenetic analyses of the ARF/Arl/Sar gene family and their regulators in Apicomplexa and close outgroup taxa. A) Analysis of ARF/Arl/Sar homologues. Note that, although the expected clades are present, support for each clade varies considerably. B) Analysis of ARF/Arl/Sar homologues with the exception of the highly divergent ArlPlasmo homologues, the omission of which improves the overall support values. C) Scrollsaw analysis of ARF/Arl/Sar homologues. Each pan-eukaryotic clade is represented by the 15 least divergent representatives, based on scrollsaw analysis. Note the similar topology compared to (B) but overall increased support. D) Analysis of ArlX1 homologues. The same dataset as in (C) was used, but ArlX2 and ArlX3 homologues were omitted. Note the supported sister relationship between Arl16 and ArlX1. E) Analysis of ArlX2 homologues. The same dataset as in (C) was used, but ArlX1 and ArlX3 homologues were omitted. Note the supported sister relationship between Arl16 and ArlX2. F) Analysis of ArlX3 homologues. The same dataset as in (C) was used, but ArlX1 and ArlX2 homologues were omitted. Note the sister relationship between Arl16 and ArlX3, though without bootstrap support. G) Analysis of ArlX homologues with closely related sequences. Note the close relationship between Arl16 and ArlX1, which lacks RAxML bootstrap support but is otherwise supported. H) Analysis of ArlX1 homologues with closely related sequences. Note that the ArlX1/Arl16 relationship in (G) is now supported in all three methods. I) Analysis of ArlX2 homologues with closely related sequences. Note the sister relationship between Arl6 and ArlX2, albeit without statistical support. J) Analysis of ArlX3 homologues with closely related sequences. Note the sister relationship between Arl16 and ArlX3 sequences, albeit without RAxML bootstrap support. K) Analysis separating BIG and GBF homologues. L) Analysis of Cytohesin-like sequences. Note the clear grouping of TBS sequences apart from E. huxleyi sequences, which group with other cytohesin sequences (asterisks). Similarly, dinoflagellate ARCC sequences group apart from the long E. huxleyi representative (dollar sign). M) Analysis of ArfGAP homologues. Note the grouping of putative ArfGAPC2 homologues, albeit without RAxML bootstrap support.

Figure S4. Co-regulation heat map of LSP genes and known Microneme and Rhoptry genes.

Heat map showing Pearson correlation coefficients between gene expression profiles obtained from publicly available RNA-Seq data from various T. gondii developmental stages. Correlations are shown for LSP genes and known genes of the Microneme and Rhoptry machinery (highlighted in light blue and orange, respectively). Colors correspond to levels of correlation as indicated to the right of the figure. Two clusters of co-regulated genes (1 and 2) are outlined in black and described in the main text.

Figure S5. Tagging of Arl LSPs.

A) Scheme of LIC tagging strategy. P1 and P1’ are the primers used for confirmation of integration (see Table S4 for specific primers). B) Agarose gel showing integration of 3xHA tag in ArlX1 locus. C) Western blot showing expression of ArlX1-HA. D) Agarose gel showing integration of 3xHA and YFP tags in ArlX2 locus. E-F) Western blot showing expression of ArlX2-HA (E) and ArlX2-YFP (F). G) Agarose gel showing integration of 3xHA tag in ArlX3 locus. H) Western blot showing expression of ArlX3-HA.

Figure S6: Localisation assays of Arl LSPs proteins.

A) ArlX1 localised apical to the conoid and in some occasions at the filamentous intravacuolar network. mCherry-α-Tubulin: microtubules; mCherry-Morn1: basal ring in mother and daughter cells; α-ISP1: apical cap. B) ImmunoEM performed on intracellular ArlX1-HA parasites. Arrows point to gold labeling in each section. Abbreviations: M, microneme; R, rhoptry; v, vesicle. Scale bars are as indicated on each image. C) ArlX3 showed a localisation apical to the nucleus but did not localise with the ER (P30-GFP-HDEL) but localised with Golgi markers (GRASP-RFP and GalNac-YFP). D) ImmunoEM performed on intracellular ArlX3-HA parasites. Arrows point to gold labeling in each section. Abbreviations: M, microneme; R, rhoptry; v, vesicle. Scale bars are as indicated on each image E) Scheme showing the position of the Cas9 targeted sequence in arlX loci. Next to the name of the gene is the phenotypic score obtained by Sidik et al (2016). On top of each arrow is the specific score for the selected gRNA.

Figure S7: Transient Cas9 targeting Arl LSPs show a possible effect on micronemes, Golgi and ECL when disrupting ArlX3.

A) Effect on M2AP after depletion of Arl LSPs. B) Effect on Mic4. Note that Mic 4 localisation shifts to an apical position in the parasite. C-D) Effect on rhoptries. E) Effect on Golgi. α-DrpB antibody was used as Golgi marker. F) Effect on Endosomal like compartment (ECL) α-Pro-M2AP antibody used as a marker for ECL. Scale bars = 10μm

Figure S8: ArlX1-KO and ArlX2-KO characterisation.

A) After transient transfection of gRNA targeting ArlX2, 5 clones were isolated and PCR around the targeted region. Agarose gel and sequencing of PCR products are shown here. B) Plaque assay of ArlX2KO showed no difference when compared to wild type parasites. C) Quantification of plaque size shown in B. D) Images depicting normal localisation of micronemes (α-Mic6 and α-Mic8 antibodies), dense granules (α-GRA1 and α-GRA7 antibodies) and ECL (α-ProM2AP antibody). Scale bar = 5μm. E) Scheme showing strategy followed for the integration of STOP codons in the ArlX1 locus. F) agarose gel showing PCR integration of STOP codons G) Sequences of the targeted region in wildtype and ArlX1-KO. H) Plaque assay of ArlX1-KO showing a slight reduction in parasite growth that was quantified in I. J) Invasion assay showed a discrete drop in parasite invasion. K) Replication assay showed no significant changes in parasite replication regarding the wild type. L) Parasites were able to egress normally when compared to the wildtype. Mean ± SEM are represented in all graphs. ns: not significant differences, **** = *p*< 0.0001

Figure S9: Generation of ArlX3-iKD.

A) Scheme showing the strategy followed to achieve promotor replacement. B) Integration PCR showed correct integration of the TetO7-pSag1-myc cassette (T7S1) in the ArlX3 locus. C) Western blot showing protein expression of T7S1-ArlX3. D) Western blot showing downregulation of ArlX3 expression as soon as 24h after addition of ATc. α-aldolase antibody was employed as loading control. E) STED images of micronemes upon induction with ATc at the indicated time points. F) STED images of rhoptries after addition of ATc at the indicated time points. Scale bar = 2 μm

Figure S10: Golgi fragmentation in ArlX3-iKD parasites.

A) Transient transfection of GRASP-RFP (cis Golgi) in ArlX3-iKD parasites. B) transient transfection of GalNac-YFP in ArlX3-iKD parasites. Scale bars = 10 μm C) STED images of ArlX3-iKD stained with α-myc antibody (ArlX3) or α-GFP antibody (SORTLR). Scale bars = 2 μm.

## Notes

**Competing Interest Statement:** Authors have no competing interests.

### Competing Interest Statement

The authors have declared no competing interest.

