## Supplementary text for "Evolution of lineage-specific trafficking proteins and a novel post-Golgi trafficking pathway in Apicomplexa"

#### S1. Homology Searching and Phylogenetic Analysis

Predicted proteomes of all organisms under study were downloaded from relevant public databases; information regarding all datasets is found in Table S3. Initial identification of homologs was performed using HMMer v3.1b1 (63) followed by reciprocal BLASTp (v2.8.1) (64) searches against the *Homo sapiens* predicted proteome to check for false positives. All homology searching employed an e-value cut off of 0.05; discrimination between positive and negative hits in reciprocal BLAST used a two order of magnitude cut off, whereby true homologs were considered to hit a relevant homolog in *H. sapiens* with an e-value at least two orders of magnitude greater than the first non-homologous hit. In some cases, additional homologs were identified by reciprocal BLASTp analysis (64) using an identified homolog from the most closely related taxon within the dataset. Domain prediction used PfamScan v1.6 (65) with an e-value cut off of 0.01; reported start and stop positions represent the domain "envelope". All results of homology searching and domain prediction analysis can be found in Table S1.

All alignments were carried out using MAFFT v7.407 (66); for alignments less than ~250 sequences, the slow and accurate L-INS-i method was used, while larger alignments used the —auto option. Alignments were manually inspected and trimmed by hand. IQ-TREE v1.6.11 (67) was used for rapid inference of large datasets, under the best model as inferred by each program run and performing 1000 replicates for ultra-fast bootstrapping. RAxML v8.2.12 (68) was used both for initial phylogenies during sequence classification, employing maximum-likelihood tree inference and rapid bootstrapping with 100 replicates for each run, and for final bootstrapping of datasets, using the autoMRE criterion to determine a sufficient number of bootstrap replicates for each dataset (69). In all cases, model selection was performed by RAxML during each program run. Bayesian phylogenies were performed using MrBayes v3.2.7a (70). Four independent runs of four chains were run for 1,000,000 MCMC generations, sampling every 500 generations under a mixed amino acid model. The consensus tree and statistics were calculated following removal of the first 20% of samples from each run as burn-in. For all phylogenetic inference, rate variation among sites was modeled using a discrete gamma distribution with four rate categories. All trees were viewed using FigTree v1.4.4 (https://github.com/rambaut/figtree) and tree figures manually modified using Affinity Designer v1.6.1 for Mac.

Further sequence filtering used the scrollsaw method of choosing the N best representative sequences from each group among several groups using mutual minimal genetic distance as a selection metric (1). For simplicity, we wrote a scrollsaw implementation in the Python 3 programming language, available at https://github.com/chris-klinger/scrollpy. Our implementation first aligns sequences before calling the -f x option in RAxML (v8.2.12, run locally with the LG+$\Gamma$ substitution model) (2) to calculate pairwise maximum-likelihood distances. The program ranks all sequences by mutual minimal ML distance and optionally returns the top N sequences.

To validate this implementation, we ran simulations to generate sequences according to various branch lengths. For each set of branch lengths we generated a random tree with five clades of either three, four, or five sequences each, and then evolved a representative sequence (H. sapiens KLHL8; NP_065854.3) from the root to the tips using Pyvolve v1.0.3 (3) (using the LG+$\Gamma$ with four rate categories and an alpha of 0.5). We calculated all-vs-all distances using the tree branch lengths and between the simulated sequences using our scrollpy implementation; mismatches between the order of each were compared to the total possible number of mismatches given the number of sequences per clade. Running 100 simulations for each yielded an accuracy of 96.67 ± 3.78, 95.64 ± 3.22, and 95.58 ± 2.21% for three, four, and five sequences per clade, respectively (Table S5). The small differences between tree and sequence distances may be due to stochasticity in the simulation results; regardless, the simulation results demonstrate that our method accurately selects sequences with mutual minimal distances.

### S2. SNAREs

Soluble *N*-ethylmaleimide-Sensitive Factor Attachment Protein (SNAP) receptors (SNAREs) are relatively small (~200 amino acid) coiled-coil proteins involved in vesicular fusion (4). Previous studies have demonstrated four families within the SNARE superfamily; the classification depends to some extent on whether a glutamine (Q) or asparagine (R) is present at the 0-layer of the SNARE domain. These families are the Qa, Qb, Qc, and R families (5, 6) homologues of all these families were investigated (Figure S1, Table S1).

### S2.1 Qa SNAREs

We identified putative homologues of the Qa SNAREs Syntaxin-2 (Stx2), Stx5, Stx12, Stx16, and Stx18 in our study taxa (Figure S1, Table S1). Conservation across study taxa was excellent, apart from the lack of Stx12A homologues in *Cryptosporidium spp.* and sporadic absence of one or two of the identified Qa SNAREs in other taxa. In these latter cases, it is unclear whether the absence represents true absence of false negatives.

Initial phylogenetic reconstruction (Figure S2A) separated the identified homologues into robust clades: Stx2 (Bayesian posterior probability 0.99/RAxML rapid bootstrap 76/IQ-TREE ultrafast bootstrap 98), Stx5 (1/100/100), Stx12 (0.99/50/99), Stx16 (1/99/100), and Stx18 (1/85/99). Stx2 and Stx12 were each other's closest neighbours (1/81/100) while Stx5 and Stx18 were also more closely related to each other (1/87/100), with Stx16 branching between these groups.

We also noted some internal duplications within the Stx2 and Stx12 clades. Though most apicomplexans possess a single orthologue of each Qa SNARE, the piroplasmids and various *Plasmodium* species possess a second orthologue of Stx2 (0.96/71/99, Figure S2A). Compared to this later duplication, an earlier duplication in Stx12 (referred to as Stx12B to delineate it from the pan-eukaryotic Stx12A), likely at the base of the myzozoa, was also observed (0.99/78/100, Figure S2A).

To confirm each of these, the relevant clade (Stx2 or Stx12) was run in a separate analysis with Stx16 as an outgroup. These analyses confirmed the deep duplication of Stx2 within Apicomplexa (0.95/60/96, Figure S2B) and the ancestral myzozoan duplication of Stx12 (1/95/100, Figure S2C). Hence, Stx12B represents a lineage-specific paralogue (LSP) by our criteria.

### S2.2 Qb SNAREs

We identified putative homologues of the Qb SNAREs GOSR1, GOSR2, NPSN11, Sec20, and Vti1 in our study taxa (Figure S1, Table S1). Conservation within apicomplexans was excellent, although GOSR2 appears absent from piroplasmids. Outside of the Apicomplexa, we failed to identify several of these Qb SNAREs across taxa, including the dinoflagellate *S. kawagutii*, the ciliates *I. multifiliis* and *P. tetraurelia,* and the archaeplastid *C. cripsus*. The presence of orthologues in closely related taxa suggest that at least some of these absences may be artefactual.

An initial phylogenetic reconstruction (Figure S2D) reconstructed each clade: GOSR1 (0.79/88/100), GOSR2 (0.86/85/100), NPSN11 (0.99/90/100), Sec20 (0.97/86/100), and Vti1 (0.84/46/99). GOSR1 and GOSR2 were consistently observed as sister clades (0.89/62/98 in Figure S2D), while the exact topology of NPSN11, Sec20, and Vti1 varied. The large number of Vti1 homologues, including up to three in some apicomplexans like *T. gondii* (Table S1), hinted at the possibility of additional LSPs. However, a separate analysis of Vti1 homologues using NPSN11 as an outgroup revealed very little supported internal structure (Figure S2E). Though apicomplexans encode multiple Vti1 homologues, their relationships remain poorly resolved.

To improve resolution in our Qb analyses, we took the 20 least divergent Vti1 homologues, as assessed by scrollsaw (materials and methods) and re-ran our Qb phylogeny (Figure S2F). Overall, support for each clade increased: GOSR1 (1/92/100), GOSR2 (0.93/84/98), NPSN11 (1/95/100), Sec20 (1/88/100), and Vti1 (1/94/100). The sister relationship between GOSR1 and GOSR2 remained unchanged (0.98/67/98), while NPSN11 and Vti1 branched as sisters (0.82/62/98).

Overall, these results confirm the presence of at least one homologue each of the Qb SNAREs GOSR1, GOSR2, NPSN11, Sec20, and Vti1 in apicomplexans. Additional Vti1 homologues were noted, though the pattern of duplications that gave rise to these paralogues is unclear.

#### S2.3 Qc SNAREs

We identified putative homologues of the Qc SNAREs Bet1, Stx6, Stx8, SYP71, and Use1 in our study taxa (Figure S1, Table S1). Conservation within apicomplexans was excellent, although Use1 appears absent from cryptosporidians. Outside of the Apicomplexa, we failed to identify several of these Qc SNAREs across taxa, including the dinoflagellate *S. kawagutii*, the ciliate *I. multifiliis,* the haptophyte *E. huxleyi*, and the archaeplastid *C. cripsus*. The presence of orthologues in closely related taxa suggest that at least some of these absences may be artefactual.

An initial phylogenetic reconstruction (Figure S2G) reconstructed each clade: Bet1 (1/86/100), Stx6 (0.94/62/96), Stx8 (0.97/78/100), SYP71 (1/93/100), and Use1 (0.89/91/100). There appears to be two distinct clades of Stx6 homologues in stramenopiles, the first branching on its own (1/89/100) and the latter branching with haptophytes and other taxa (1/84/99). No other duplications outside of single taxa were evident and no LSPs were identified.

Overall, these results confirm the presence of the Qc SNAREs Bet1, Stx6, Stx8, SYP71, and Use1 in apicomplexans and their close relatives.

#### S2.4 Qbc SNAREs

During our homology searching and phylogenetic analyses, we identified 41 sequences comprising two SNARE domains and which appeared to group together in phylogenetic analyses of Qb and Qc SNAREs. Such Qbc SNAREs have been described previously (6). Within our dataset, we identified Qbc homologues across apicomplexans, with a likely secondary loss in piroplasmids, and chromerids, with some conservation in more distantly related groups (Figure S1, Table S1).

To better understand the origins of these Qbc SNAREs, we extracted the relevant Qb and Qc SNARE domains from each and ran them together with identified Qb/Qc homologues from our datasets. The overall topology of Qb SNAREs remained unchanged with the addition of the Qbc Qb SNARE domains (Figure S2H), with GOSR1 and GOSR2 remaining as sister clades (0.99/49/95). NPSN11 and Vti1 remained closely related, with the Qb domain of Qbc SNAREs placing basal to NPSN11 (0.9/48/81); the group as a whole had similar support (0.82/45/93). The clades themselves had variable support: Qbc (0.84/40/80), NPSN11 (0.88/28/94), and Vti1 (0.95/71/94).

Similarly, the addition of the Qc domains of Qbc proteins did not drastically alter the topology of Qc SNARE homologues (Figure S2I). Stx6 and Stx8 branched as sister clades, albeit with very low RAxML bootstrap support (0.81/28/99). Unlike the Qbc Qb domain, the Qc domain did not readily form a monophyletic clade, with the myzozoan sequences branching separately from the other Qc domains. SYP71, which was previously well supported (Figure S2G), lacked support (0.55/15/62). The overall group of the Qbc Qc domains, SYP71, and Use1 had better support (0.82/35/99) and the Use1 clade specifically was a well-supported clade within this larger group (0.87/80/100).

These results demonstrate the presence of Qbc proteins in the myzozoa, with sparse retention in a variety of more distantly related taxa. Overall, these proteins appear to be the result of a single gene fusion, likely involving the “plant syntaxins” NPSN11 (the Qb domain) and either SYP71 or Use1 (the Qc domain).

#### S2.5 R SNAREs

We identified putative homologues of the R SNAREs Sec22, Ykt6, and VAMP7 across our study taxa (Figure S1, Table S1). Additional poorly defined VAMP-like proteins were also detected (termed “VAMPX”), but were excluded from phylogenetic analyses due widespread variation in length and their generally divergent nature. Conservation was excellent overall, with most taxa including at least one homologue of each identified R SNARE.

Initial phylogenetic reconstruction (Figure S2J) separated the identified homologues into robust clades: Sec22 (1/99/100), Ykt6 (1/100/100), and VAMP7 (0.99/92/99). Careful inspection of the VAMP7 and Ykt6 clades revealed groupings of myzozoan sequences in each, which we refer to as VAMP7B/Ykt6B to separate them from the pan-eukaryotic VAMP7A/Ykt6A. Both the VAMP7B (0.95/95/100) and Ykt6B (0.98/85/99) clades were well-supported.

To further explore the VAMP7A/B sequences, we constructed a separate phylogeny using Sec22 as an outgroup (Figure S2K). The VAMP7A clade on the whole lacks strong internal structure and the branching order of groups does not reflect the taxonomic groups; this is reflected in the short branch lengths and low bootstrap support for most nodes. Of interest, there appears to be a duplication in archaeplastids; *B. natans* and the two *Phytophthora* species branch basal to the first (0.95/46/88) group while *G. theta* is basal to the latter (0.83/50/95). Despite the overall low support, the VAMP7B clade is well-supported (0.94/87/100). The conservation pattern within Apicomplexa was also peculiar. All *Plasmodium* and *Cryptosporidium* species possess VAMP7A but not VAMP7B while all piroplasmids lack VAMP7A but encode VAMP7B.

We conducted a similar analysis of Ykt6A/B sequences, again using Sec22 as an outgroup (Figure S2L). There appeared to be two distinct ciliate Ykt6A clades, one that grouped with the majority of myzozoan sequences (0.98/83/100) and the other that branched basal to the divergent piroplasmid Ykt6A sequences (1/100/100). Although the topology obtained could suggest a third Ykt6 paralogue, the low support for each separate group (0.89/29/90 and 0.73/33/89) and the variety of topologies observed in separate analyses favours the view that these represent divergent Ykt6A orthologues instead. Comparatively, the Ykt6B clade was clearly distinct (0.98/72/96) and was conserved across chromerids and apicomplexans. We failed to identify a dinoflagellate homologue.

Our results suggest the presence of two novel R SNARE paralogues in the myzozoa: VAMP7B and Ykt6B. VAMP7B likely arose at the base of the myzozoa and was retained variably across Apicomplexa. Curiously, piroplasmids have retained this novel paralogue but have lost the ancestral VAMP7A paralogue. Ykt6B likely arose after the divergence of dinoflagellates, but is well conserved in both chromerids and apicomplexans.

### S3. Rab GTPases

Ras-like proteins from rat brain (Rab) GTPases are members of the Ras G protein superfamily with diverse roles including in membrane trafficking (7). We identified 1,590 putative Rab homologues across our dataset, including at least one representative of each of the 23 ancient Rabs identified previously (1, 8). As reported previously, apicomplexans encode homologues of Rabs 1, 2, 4, 5, 6, 7, 8, 11, 18, and Rab23 (Figure S1, Table S1). Chromerids additionally encode homologues of Rabs 23, 28, 32, L2/RTW, and L4/IFT27 and dinoflagellates Rab21 and Rab34, suggesting that the common ancestor of myzozoans encoded almost all known pan-eukaryotic Rabs (Figure S1, Table S1).

An initial ML-only phylogenetic analysis revealed a topology largely consistent with that observed previously (1), though without bootstrap support for many nodes (Figure S2M). The various Rab5 homologues grouped with other “endocytic” Rabs as expected, with Rab6 branching basal. Another grouping consisted of Rabs 7, 23, 28, 32, Titan, L2/RTW, and L4/IFT27, though again without strong support. The various Rab1 homologues grouped together with Rabs 8 and 18 with some measure of support (39/98). Finally, Rabs 2, 4, 11, and 14 branched together with a previously undescribed Rab-like protein (RabX1).

Despite the overall topology resembling previous Rab phylogenies, the large number of sequences and presence of divergent homologues impaired the resolution and statistical support for almost all backbone nodes. In order to overcome this obstacle, we took the 15 least divergent representatives of each clade assessed using the scrollsaw method (materials and methods) and reconstructed the phylogeny (Figure S2N). Three clear groups emerged: Rab5-related ("endocytic Rabs") sequences with Rab6 as an outgroup (0.69/39/92), Rab1/2/11-related ("exocytic Rabs", 0.99/56/100), and all other Rabs (0.7/19/83). Though only the exocytic Rab group was supported by all three methods, all three groups were consistently observed across analyses and closely mirror groupings observed previously (1).

In general, despite a lack of backbone support, the individual Rab clades were well supported in both the scrollsaw (Figure S2N) and all sequence (Figure S2M) analyses. It was clear that several LSPs were present in our Rab dataset; the consistent relationships we observed allowed us to run additional phylogenies with smaller subsets of our full Rab dataset in order to solidify the identity of each LSP.

#### S3.1 Rab1-like

A previous study identified a novel paralogue of Rab1 present in various SAR taxa as well as the cryptophyte *G. theta* (1). In our large phylogenetic analysis, we noticed several putative Rab1-like clades (Figure S2M,N), including one (which we refer to as Rab1A to keep the nomenclature consistent) that corresponds to this previously identified paralogue. Our scrollsaw analysis confirmed the grouping of all Rab1-like sequences with Rabs 8 and 18 (Figure S2N, 1/83/100).

To further investigate the Rab1-like sequences in our dataset, we constructed an additional tree using the closely related Rab18 as an outgroup (Figure S2O). As expected, the pan-eukaryotic Rab1B was basal to the more restricted Rab1A clade (0.99/86/100), which contained stramenopiles, alveolates, and the cryptophyte *G. theta* (Figures S1,

S2O). These results confirm those obtained previously, namely a Rab1 duplication deep in eukaryotic evolutionary history.

We noticed an additional clade, which was restricted to just dinoflagellates, chromerids, and coccidians, which we refer to as Rab1K (1/100/100). These sequences are unusual, generally around 600 amino acids compared to ~200 for other Rabs, with an extended N-terminus that harbours several Kelch domain repeats (hence, Rab1K). As multiple Kelch domains are known to adopt a β-propeller fold, we carried out template-based modelling of the *T. gondii* orthologue (Figure S2O). The resulting structure revealed an N-terminal six-bladed β-propeller connected to a structurally canonical Ras domain and a largely unstructured C-terminal end (the C-terminus alternatively adopted an extended α-helical fold or was unstructured across the five returned models).Our results confirm the presence of two Rab1-like LSPs, Rab1A and Rab1K, with the former representing a deep duplication in eukaryotic history. Based on the topology observed (Figure S2O), Rab1A further duplicated in the myzozoa and accrued multiple N-terminal Kelch domains to yield the more taxonomically restricted Rab1K.

### S3.2 Rab5-like

Previous studies have demonstrated the presence of three Rab5-like proteins in *T. gondii* and other apicomplexans, known as Rab5A, Rab5B, and Rab5C (9).

Based on previous phylogenies, as well as our own large-scale (Figure S2M) and scrollsaw (Figure S2N) analyses, we constructed trees with Rab5-like GTPases: Rab5A, Rab5B, Rab5C, Rab20, Rab21, Rab22, Rab24, and Rab50 (Figure S2P). Rab6 was included as an outgroup. The tree was well resolved overall, with statistical support for each individual clade.

In these analyses, the Rab5A-C clades grouped together, but with comparatively low support (0.7/25/77). Within this group, Rab5B branched as basal to a combined Rab5A/5C clade (0.98/56/94), wherein the two clades were sister and were distinct with moderate support (Rab5A 1/52/94 and Rab5C 1/97/94). Although the grouping of all three Rab5 paralogues together was not well-supported it was consistently observed across numerous analyses, suggesting that Rab5B and Rab5C represent *bona fide* Rab5 paralogues.

A smaller analysis of the three Rab5 paralogues alone with Rab6 as an outgroup yielded a very similar topology as before with Rab5B basal to a single Rab5A/5C split (Figure S2Q). Support for the Rab5A/5C grouping was higher (1/60/96), while support for each of the Rab5A (1/45/96), Rab5B (1/91/100), and Rab5C (1/95/100) clades varied.

Our results confirm the presence of three Rab5 paralogues, Rab5A-C, in Apicomplexa, and further confirm that Rab5A and Rab5C are each other's closest relatives.

#### S3.3 Rab11-like

Previous studies have demonstrated the presence of two Rab11 paralogues in apicomplexans and their ciliate relatives, known as Rab11A and Rab11B (10, 11). These two paralogues formed a clear clade in our scrollsaw phylogenetic analysis (1/98/100, Figure S2N), with each paralogue having excellent support (0.95/84/100 for Rab11A and 1/100/100 for Rab11B).

To further investigate the Rab11 sequences in our dataset, we constructed an additional tree using the closely related Rab18 as an outgroup (Figure S2R). The topologies we constructed varied slightly depending on taxon and site selection, with Rab11B branching either as sister to, or within, the pan-eukaryotic Rab11A clade. However, the Rab11B clade was always strongly supported (1/94/100 in Figure S2R).

In the course of our analyses, we also noted the presence of a conserved four amino acid insertion in all Rab11B homologues (Figure S2R). By inspecting secondary structural predictions and conserved sequence domains, this insertion appears after the G4 motif (between the β5 and α4 conserved structural elements). The first two positions show a strong preference for DP, with less conservation in the following two positions. Future studies should examine the potential relevance of this conserved insertion.

Our results confirm the presence of an alveolate-specific Rab11 paralogue, Rab11B, whose members share a conserved four amino acid insertion compared to the pan-eukaryotic Rab11A paralogue.

### S3.4 RabX1

One Rab-like protein in our dataset was not clearly homologous to any existing human Rab protein, with the top hit in reverse BLAST varying between Rabs 4, 13, 27, 36, and other seemingly unrelated homologues (Table S1). Our initial Rab scrollsaw phylogenetic analysis placed this Rab, which we refer to as RabX1, in a large group like that identified in (Elias 2012) together with Rabs 7, 23, 28, 32, 34, RTW/RabL2, IFT27/RabL4, and Titan, albeit with very weak support (0.7/19/83, Figure S2N). Despite this low support, the sister relationship between RabX1 and RabL2 was appeared to be supported (0.98/42/88).

To further investigate the possible origin of RabX1, we constructed another phylogeny using this more restricted set of sequences (Figure S2S). Overall, the backbone of the phylogeny remained poorly resolved, though we did note a sister relationship between IFT27 and Rab28 that had been reported previously (1); however, similar to the previous report, this relationship did not reach statistical significance (0.63/41/81). Conversely, we did not observe a previously reported sister relationship between Rab32 and RabTitan, though this was also not significant in the previous analysis (1).

Surprisingly, the same sister relationship was observed between RabL2 and RabX1 that we observed in the scrollsaw dataset (Figures S2N, S), albeit with improved support (0.99/61/96). In addition, Rab7 remained the most basal clade to this sister grouping, though this lacked significant bootstrap support (0.83/31/79).

Hence, it appears that RabL2, which is conserved in chromerids and dinoflagellates, duplicated at the base of the myzozoa to yield the enigmatic RabX1; the former was lost in apicomplexans while the latter was retained.

### S4. TBCs

Rab GAPs primarily belong to a large family of Tre-2/Bub2/Cdc16 (TBC) proteins that contribute both an arginine and a glutamine finger to mediate GTP hydrolysis (12). A previous analysis identified 13 TBCs conserved across eukaryotes: TBC-B, -D, -E, -F, -G, -H, -I, -K, -L, -M, -N, -Q, and -RootA, albeit with a patchy distribution (12, 13).

We identified 1,047 TBC homologues across our dataset, including homologues of all previously identified pan-eukaryotic TBCs (Figure S1, Table S1). Apicompexans retain the majority of pan-eukaryotic TBCs, only lacking orthologues of TBC-B, TBC-H, and TBC-L. Chromerids encode TBC-H and TBC-L; combined with the observed retention in dinoflagellates, it is clear that the myzozoan ancestor encoded a homologue of every TBC except TBC-B, which was previously reported as absent from SAR (13).

Previous phylogenetic analysis supported three groups of TBCs: one consisting of TBC-A, -N, -Q, and -RootA, the second TBC-G and -M, and the third comprising TBC-B, -D, -E, and -F (13). To further investigate the relationships between TBC proteins, we constructed a phylogeny of all TBC homologues (Figure S2T). We observed similar large groups as those observed previously: group 1, comprising TBC-K, -N, -Q, -RootA and other related proteins (94/100), group 2, with TBC-G, -H, -I, -L, and -M (68/97), and group 3, with TBC-B, -D, -E, and -F (79/99).

Despite a large number of sequences our first ML-only tree was well-resolved. We also decided to run a scrollsaw tree using the 15 least divergent representative sequences for each clade (Figure S2U). Encouragingly, the same groups were observed with good support: group 1 (1/85/99), group 2 (0.98/50/90), and group 3 (0.96/65/98).

#### S4.1 TBC-N and TBS proteins

A previous analysis reported that the TBC domain of TBS proteins is derived from the TBC-N clade, with independent fusions in both alveolates and the haptophyte *E. huxleyi (14)*. Our large-scale analyses appeared to support this relationship, with the combined grouping of TBC-N with the TBC domain of TBS proteins being well-supported in both conventional (92/93, Figure S2T) and scrollsaw (1/97/100, Figure S2U) analyses.

To further characterize these relationships, we constructed a separate phylogeny of the TBC domain of TBS proteins and all TBC-N homologues in our dataset, as well as TBC-K as an outgroup (Figure S2V). As expected, the alveolate TBS TBC domains formed a single well-supported clade that branched within the larger TBC-N clade (0.91/55/93), while the lone *E. huxleyi* representative formed a deep branch within a large sub-group of TBC-N proteins containing stramenopile, cryptophyte, and some alveolate sequences (0.63/1/52).

Within the myzozoan TBS grouping, there was a clear duplication in *T. gondii* and closely related coccidian species with one clade branching basal to the other sequences

(1/99/100) and the other within (1/100/100); *V. brassicaformis* also appeared to have two distinct paralogues (Figure S2V).

Our results confirm those reported previously (14) and suggest a further deeper duplication of TBS proteins within the Apicomplexa themselves.

#### S4.2 TBC-PI

A previous large-scale study investigating the conservation of TBC proteins across eukaryotes identified two paralogues, TBC-PIA and TBC-PIB, conserved only in archaeplastids. We similarly identified TBC-PI homologues in our dataset (Table S1). Curiously, in our large-scale TBC phylogenetic analyses, we observed grouping of TBC-PI homologues with identified sequences from the myzozoa that we termed TBC-X2 (72/100, Figure S2T).

To understand the relationship between TBC-PI and TBC-X2 homologues, we performed a separate phylogenetic analysis using TBC-K as an outgroup (Figure S2W). Surprisingly, the archaeplastid TBC-PI homologues branched within the larger TBC-X2 clade between the larger myzozoan grouping and an apparent duplicated group including dinoflagellates and *V. brassicaformis*. However, while the archaeplastid TBC-PI grouping itself was well-supported (0.99/75/95), the relevant backbone node lacked support by RAxML (0.84/43/82).

Our data suggest that TBC-PI, previously reported as an archaeplastid-specific TBC protein, represents a larger clade with a sparse distribution in the diaphoretickes.

#### S4.3 TBC-Q-like

We identified 147 sequences within the alveolates that appeared to form several distinct clades, similar to TBC-Q but clearly distinct due to the clear presence of alveolate TBC-Q homologues (Figure S2T). Despite variable numbers of homologues within each clade, sufficient sequence converage was present to determine a likely origin at the base of alveolates in each case.

We constructed a separate tree to determine the relationship of these sequences to one another and to TBC-Q, using TBC-K as an outgroup (Figure S2X). The resulting tree confirmed the separation of these sequences into clades: TBC-Q (1/71/91), TBC-X1

(1/99/100), TBC-X3 (1/96/100), TBC-X4 (1/83/99), and TBC-X5 (0.63/35/92). Although TBC-X5 lacked statistical support in this analysis, we consistently observed this relationship, and these sequences are confidently excluded from all other related clades. Together, the TBC-X clades were distinct from TBC-Q (0.52/44/86), though clearly related to it.

Thus, we have identified four novel clades of TBC-Q-related proteins conserved within alveolates: TBC-X1, TBC-X3, TBC-X4, and TBC-X5. In all cases except TBC-X4, there appears to have been massive expansions of each homologue within the ciliates with variable retention within the myzozoa.

### S5. Arf/Arl/Sar G proteins

The Arf-related G protein superfamily, comprising Sar, ARF, and ARF-like (Arl) proteins, is another part of the Ras superfamily. Similar to Rabs, Arf family proteins have diverse roles in membrane trafficking and other non-trafficking roles as well (15).

We identified 700 Arf-related proteins across our dataset, including putative homologues of Sar, ARF, ARF-related protein (ARFRP), and Arls 1, 2, 3, 5, 6, 8, 13, and 16 (Figure S1, Table S1). Whereas all study taxa encoded at least one homologue of both ARF and Sar, conservation among Arl proteins was highly variable. Apicomplexans universally retain Arl2, with some groups also encoding Arl1 and ARFRP. Chromerids and dinoflagellates encode all Arls except Arls 8, 13, and 16, suggesting that three Arls (Arls 3, 5, and 6) were lost during the chromerid-apicomplexan transition. Numerous organisms outside of the myzozoa, such as *P. sojae, P. ultimum*, *B. natans*, *E. huxleyi*, *G. theta*, *C. reinhardtii*, and *C. paradoxa*, had identifiable homologues of all or nearly all Arls, suggesting a complete complement at the base of diaphoretickes.

A phylogenetic analysis of all identified homologues (Figure S3A) was generally well-resolved, with support for most individual clades and some small groupings: Arf/Arl1/Arl2/Arl3/Arl5 (40/98), Arl6/Arl8 (49/93), and ARFRP/Arl13 (31/98). One Arf-like clade that did not readily place anywhere was that of ArlPlasmo (100/100), a group of enigmatic sequences conserved only in *Plasmodium* roughly four times longer than most other Arls (~800 amino acids) and containing only an identifiable N-terminal Arf (PF00025) domain (Table S1). This group was never clearly related to any other Arf/Arl clade in any phylogenetic analysis, and, due to its divergent nature, also tended to have an adverse effect on tree topology and support values.

A separate analysis without the divergent ArlPlasmo sequences (Figure S3B) yielded a similar overall topology. All pan-eukaryotic homologues were resolved: Sar (100/100), ARF (52/94), Arl1 (66/90), Arl2 (97/100), Arl3 (93/100), Arl5 (87/97), Arl6 (91/100), Arl8 (98/100), Arl16 (71/98), and a combined ARFRP/Arl13 clade (46/100). In the course of our analyses, we also identified three putative LSPs, which we refer to as ArlX1 (73/99), ArlX2 (41/94), and ArlX3 (68/98), and which did not obviously group with any clades in large-scale phylogenies.

### S5.1 ArlX Proteins

In order to further investigate the evolutionary origins of the ArlX proteins, we obtained the 15 least divergent representative sequences for each pan-eukaryotic clade and constructed phylogenies with all ArlX homologues (Figure S3C). As before, the ArlX homologues branched outside of the consistent groupings of ARF/Arl1/Arl5, Arl2/3, Arl6/8, and ARFRP/Arl13. ArlX1 branched with Arl16, albeit with low bootstrap support (0.89/36/75), while ArlX2 and ArlX3 grouped with each other with low support (0.87/30/18). Curiously, despite relevant support in most analyses, Arls 6 and 8 grouped with very low support in this analysis (0.4/22/14).

We ran additional analyses, including only sequences corresponding to one ArlX clade each time. A tree with only ArlX1 homologues (Figure S3D) was largely the same, with ArlX1 branching together with Arl16; however, this relationship now achieved support across all three methods (0.92/57/94). When only ArlX2 homologues were included (Figure S3E), they also grouped with Arl16 (0.87/40/96). However, the topology was unusual, with this combined group branching sister to Arl6 (0.66/12/84) and Arl8 as basal rather than sister to Arl6 (0.89/24/94 for the entire group). Finally, when only ArlX3 homologues were included (Figure S3F), the topology was similar to that observed for ArlX1. Indeed, ArlX3, like both ArlX1 and ArlX2, grouped with Arl16, though without clear bootstrap support (0.82/31/58).

To further investigate the relationships, we returned to using all homologues in phylogenetic analyses, but limited our sequence selection to only those Arls that had potential relationships with the ArlX homologues: Arl6, Arl8, and Arl16; Arls 1 and 5 were included as outgroups (Figure S3G). In the resulting phylogeny, ArlX1 was again the closest sister to Arl16 (0.9/46/90), with ArlX3 branching basal to this pair (0.85/17/48). Conversely, ArlX2 branched basal (0.82/19/46) to a combined Arl6/Arl8 clade (0.83/61/92).

Including only ArlX1 (Figure S3H) improved the support for the relationship with Arl16 (1/64/96) but otherwise left the topology unchanged. Conversely, including only ArlX2 (Figure S3I), resulted in ArlX2 branching as sister to Arl6 (0.65/44/41) rather than Arl8. The tree including only ArlX3 homologues (Figure S3J) was similar to that for ArlX1, with ArlX3 branching as sister to Arl16 (0.98/46/94).

Overall, these data support the existence of three Arl LSPs, ranging in taxonomic distribution from alveolates + rhizarians (ArlX1) to Apicomplexa only (ArlX2). The origin of these Arls remains enigmatic; ArlX1 and ArlX3 appear to associate closely with Arl16, although the latter tends to be less well-supported, while ArlX2 appears loosely related to both Arl6 and Arl16. This lack of concrete relationships is reflected in the use of topology testing for the individual datasets of each ArlX homologue with the subset of pan-eukaryotic Arl homologues selected by scrollsaw (Figures SD-F).

Testing against all possible sister relationships, as well as relationships with consistently observed groups of ARF/Arl1/Arl5, Arl2/3, Arl6/8, and ARFRP/Arl13, revealed that none of the alternative topologies could be confidently rejected across all methods (Table S6). Focussing on the AU test, topologies grouping ArlX1 with Arl13 or ARFRP were rejected. Similarly, the grouping of ArlX3 and ARFRP could be rejected. Alternative placement of ArlX2 with any group could not be rejected.

Our inability to confidently place these Arl LSPs by any method mirrors their general lack of feature conservation with canonical eukaryotic homologues (for example, see Figure 1C and 1D). Hence, although these ArlXs are clearly distinct paralogues within the Arf superfamily, they represent such divergent members that it is unclear exactly how they arose.

### S6. ArfGEFs

ArfGEFs facilitate the swapping of GDP and loading of GTP on Arf GTPases (15). We identified putative homologues of the ARCC, Cytohesin, BIG, and GBF1 ArfGEFS in our dataset (Figure S1, Table S1).

BIG and GBF1 are similar large ArfGEFs, each containing characteristic DCB, HUS, and HDS1-3 domains in addition to a Sec7 domain (14). To confirm our classifications made through homology searching, we ran a phylogenetic analysis of BIG/GBF1 homologues (Figure S3K). As expected, BIG and GBF1 homologues grouped together (0.9/82/99). Within the Apicomplexa, piroplasmids and *Plasmodium* species retain only GBF1 (Figures S1, 3K). Though cryptosporidians and *G. niphandrodes* retain orthologues of both BIG and GBF1, their BIG sequences appear divergent, branching basal to all other BIG orthologues (Figure S3K).

A previous analysis of ArfGEFs across eukaryotes established that all other homologues are more closely related to cytohesin, including the ARCC and TBS Sec7 domains (14). Hence, we constructed an additional phylogeny to investigate the relationships among the remaining ArfGEFs in our dataset (Figure S3L). All myzozoan ARCC proteins grouped together as expected (1/93/98), but the lone *E. huxleyi* ARCC homologue, identified by the presence of N-terminal ankyrin repeats (Table S1), branched together with other *E. huxleyi* cytohesin homologues albeit with low support (0.52/41/58). Similarly, as previously reported (14), the Sec7 domain of alveolate TBS proteins grouped together (0.9/65/91) while the domains of TBS proteins from *E. huxleyi* branched within other cytohesin homologues (Figure S3L).

Our results confirm the presence of ARCC, BIG, Cytohesin, GBF1, and TBS proteins in our study taxa. While chromerids and dinoflagellates possess homologues of all ArfGEFs except Cytohesin, apicomplexans additionally lack ARCC homologues. At the extreme, piroplasmids and *Plasmodium* only encode a lone GBF1 protein.

### S7. ArfGAPs

ArfGAPs represent the known GTPase activating proteins (GAPs) of the Arf family, united by a common ArfGAP domain (16). We identified putative homologues of ACAP, AGFG, ArfGAP1, ArfGAP2/3, ArfGAPC2, and SMAP in our dataset (Figure S1, Table S1).

To confirm the classifications based on homology searching and investigate the relationships between the identified ArfGAP homologues, we conducted a phylogenetic analysis (Figure S3M). Several ArfGAPs resolved into well-supported monophyletic clades: ACAP (0.98/73/100), AGFG (0.98/100/100), and ArfGAPC2 (0.97/43/98). ACAP and ArfGAPC2 branched from within a polyphyletic grouping of SMAP homologues, which generally did not form a single clade in any of our analyses but which were confidently classified by homology searching and domain analysis. Curiously, there appears to be an internal AGFG duplication shared among stramenopiles and dinoflagellates (though not with chromerids or other apicomplexans, 1/100/100 for the internal clade).

The canonical ArfGAP1/2/3 sequences formed a single well-supported group (0.97/99/100). Within this group there was a single bifurcation into two separate clades, one corresponding to ArfGAP1 orthologues (0.88/61/95) and another to sequences that were indistinguishable between ArfGAP2 and 3 (0.54/45/87). A strict bifurcation was not always observed however; in other analyses, ArfGAP1 emerged within the ArfGAP2-3 sequences, albeit always with clear support.

Our results confirm the presence of ACAP, AGFG, ArfGAP1, ArfGAP2/3, ArfGAPC2, and SMAP proteins in our study taxa. Most apicomplexans encode only ArfGAP1, ArfGAP2-3, and SMAP, while cryptosporidians additionally encode AGFG (Figure S1, Table S1). In addition, we confirm the existence of ArfGAPC2 orthologues in the diaphoretickes, including in the dinoflagellates, stramenopiles, archaeplastids, as well as *B. natans* and *E. huxleyi*.

### S8. Generation of Arl LSP tagged lines and additional localization

We generated tagged lines for each Arl LSP using ligation-independent cloning (LIC), the scheme of which is shown in Figure S5A (17). Briefly, a C-terminal fragment of each gene to be tagged was amplified by PCR to contain a unique restriction site not present in the LIC vector backbone (for each gene, primers LIC fwd and LIC rev, Table S4). The LIC vector (pG514, Supplementary Table S4.2) was digested with PacI and then both backbone and insert were treated with T4 DNA polymerase (NEB) prior to ligation. For the vector, 6µl 10X NEB buffer 2, 3µl 100mM DTT, 2.4µl 100mM dGTP, 1.5µl T4 DNA polymerase, 0.6µl 100X BSA, and 1.2µg of digested vector prep were mixed on ice and the final volume adjusted to 60µl. For the PCR insert, 2µl NEB buffer 2, 1µl 100mM DTT, 0.8µl 100mM dCTP, 0.5µl T4 DNA polymerase, 0.2µl 100X BSA, and 0.2pm PCR product were mixed on ice and the final volume adjusted to 20µl. Each separate prep was then incubated in a thermocycler: 30 minutes at 22ºC, 20 minutes at 75ºC, 4ºC hold; reactions were held on ice prior to annealing. To anneal, 1µl of treated vector and 2µl of treated insert were mixed and incubated for 10 minutes at room temperature before addition of 1µl 25mM EDTA and five minutes additional incubation. Annealed vectors were held on ice and used to transform competent bacteria. Each vector was linearized using the corresponding unique restriction enzyme prior to transfection.

The programme FI-158 was used for electroporation. Transient transfections used ~10 µg of purified DNA and ~1x105 freshly egressed parasites, whereas stable transfections used ~20-30 µg of purified DNA and ~1x106 freshly egressed parasites. DNA for transfection was ethanol precipitated and resuspended in P3 Buffer. In the case of stable transfection, integration was selected for by supplementing culture medium with 78µM mycophenolic acid (MPA; Sigma; M3536) and 230µM xanthine (Sigma; X3627)(18); selected pools were then cloned by limiting dilution in 96 well plates and individual clones picked and analysed. All vectors used for transfection are listed in Table S4.

This tagging was carried out in parasites lacking ku80 but stably expressing split Cre recombinase (Δku80-diCre); this parental line was chosen in case Cre-mediated gene excision was required (though this option was not explored in this study). For each line, genomic DNA and protein was collected in order to confirm proper integration and expression by both integration PCR and Western blotting (Figure S5B-H).

The canonical localization of each Arl LSP is shown in Figure 1F. For ArlX2 we were able to tag with both 3xHA and YFP as shown in Figures 5 D and E. Localisation of ArlX2 was identical regardless of the tag employed.

Additional localization using various markers as a co-stain was carried out for ArlX1 (Figure S6A), though none greatly assisted in pinning down its localization. To this end, we also performed cryo-iEM labelling of ArlX1 tagged cells (Figure S6B). Mirroring the fluorescence imaging, we observed labelling at the extreme apical end of cells, but also some signal associated with rhoptries and unclassifiable vesicles. As discussed in the main text, the ArlX3 localization is consistent with a rough localization in or around the Golgi, which is supported by overlap with ERD-GFP, GRASP-RFP, and GalNAc-YFP markers (Figure S6C). Similar cryo-iEM analysis of ArlX3 tagged cells showed labelling of the Golgi, but also frequent labelling of vesicular structures and micronemes (Figure S6D).

### Generation of Arl LSPs KOs via CRISPR/Cas 9 mediated disruption

As a first step to analyse the phenotype of these Arl LSPs proteins, we decided to disrupt the genes by targeting them in parasites with intact non-homologous end joining repair (NHEJ), RHΔHx strain. To generate vectors containing a specific gRNA, the gRNA was synthesized as complementary primers (for each gene, primers gRNA fwd and rev, Supplementary Table S4). Primers were suspended in annealing buffer (10mM Tris pH 7.5, 50mM NaCl, and 1mM EDTA), heated to 95ºC, and then allowed to cool to room temperature. The parental vector was digested with BsaI and then gRNA inserts were ligated into the digested vector using T4 DNA ligase (NEB).

We transiently transfected vectors containing a fusion of Cas9 and YFP, that allows identification of transfected parasites, together with the cassette for the expression of the single guide RNA (sgRNA) targeting the gene of interest (GOI) (19, 20). To find an effective guide RNA (gRNA), we selected one already employed in Sidik et al 2016 (21). The localisation of the targeted sequence and the phenotypic score annotated for that specific sgRNA is displayed in Figure S6E. All sgRNA targeted the main Arf-ADF ribosylation factor domain. After 48 h post transfection, parasites were fixed and labelled with different antibodies targeting secretory organelles such as micronemes (MIC4 and Mic8), rhoptries (ROP1 and ROP2,4), endosomal like compartment (ECL, Pro-M2AP) and Trans-Golgi network (TGN; DrpB). While for ArlX1 and ArlX2 these markers remained unaffected, ArlX3 showed a remarkable decrease in micronemes signal and an alteration of ECL and TGN (Figure S7).

We attempted to isolate Knock-out clones (KO) after transient transfection of the Cas9-YFP-sgRNA vector by cloning parasites via limited dilution but only in the case of ArlX2 we were able to isolate parasites and confirmed indels via sequencing (Figure S8A). A further characterisation of this clone revealed that ArlX2 is not important for the growth of the parasite (Figure S8B,C) and does not affect the secretion of proteins such as dense granule GRA1 and 7, or the biogenesis of apical secretory organelles (Figure S8D).

In the case of ArlX1, a KO line was obtained after insertion of STOP codons in the three reading frames within the first exon (Figure S8E-F). This KO strain showed a very discrete growth defect when compared to the tagged line (Figure S8H,I). A more in depth characterisation of ArlX1-KO revealed a mild deficiency in parasite invasion (Figure S8J) but not effect in replication or egress when compared to the wild type (Figure S8K,L).

### Quantitative fluorescence microscopy

Time-course quantification of TATi-ArlX3 knockdown protein levels was carried out as follows. Parental Δku80-TATi and TATi-ArlX3 parasite lines were induced for the relevant time periods with 1 µg/mL ATc and processed for IFA using α-myc and α-GAP45 antibodies. Image files were loaded into Fiji and z-stacks collapsed into 2D images by summation of individual slices. For each vacuole, a region of interest (ROI) was traced in Fiji, using the α-Gap45 signal to indicate the bounding region of parasites in each vacuole. These ROIs were subsequently used to measure area, integrated density, and mean grey value in the α-myc channel. For each ROI, similar measurements were also obtained for the local background in the α-myc channel where no vacuoles were present. Subsequently, corrected total cell fluorescence was calculated as integrated density – (vacuole area x mean background fluorescence), as described previously (22). One hundred random vacuoles were quantified for each of three independent experiments. Raw data are provided in Source data.

For the analysis of Golgi vesiculation with the SORTLR marker, parasites were induced with or without ATc for 24, 48 and 72h, fixed and imaged as described above. Images were processed in Fiji and Z-stacks were collapsed into 2D images by applying maximum projection. A global thresholding was employed to mark vesicles in single vacuoles and particles bigger than 0.01 µm were analysed to calculate number of vesicles, area and mean intensity. Outlines of thresholding were obtained. At least 10 vacuoles per time-point per replicate (3 biologically independent replicates) were analysed. Mean and SEM were calculated and plotted. Raw data are provided in the Source data file.

Pearson's correlation (*r*) was calculated in Fiji using the plugin JACoP (23). Parasites were induced with or without ATc for 24, 48 and 72h fixed and α-myc antibody was used to visualised ArlX3. A single stack was isolated for the analysis of the correlation. 90 vacuoles from 3 biologically independent replicates in each time-point. Mean and SEM were calculated. Raw data are provided in the Source data file.
