## Supplementary figures and images for "Evolution of lineage-specific trafficking proteins and a novel post-Golgi trafficking pathway in Apicomplexa"

### Supplementary Figure 1

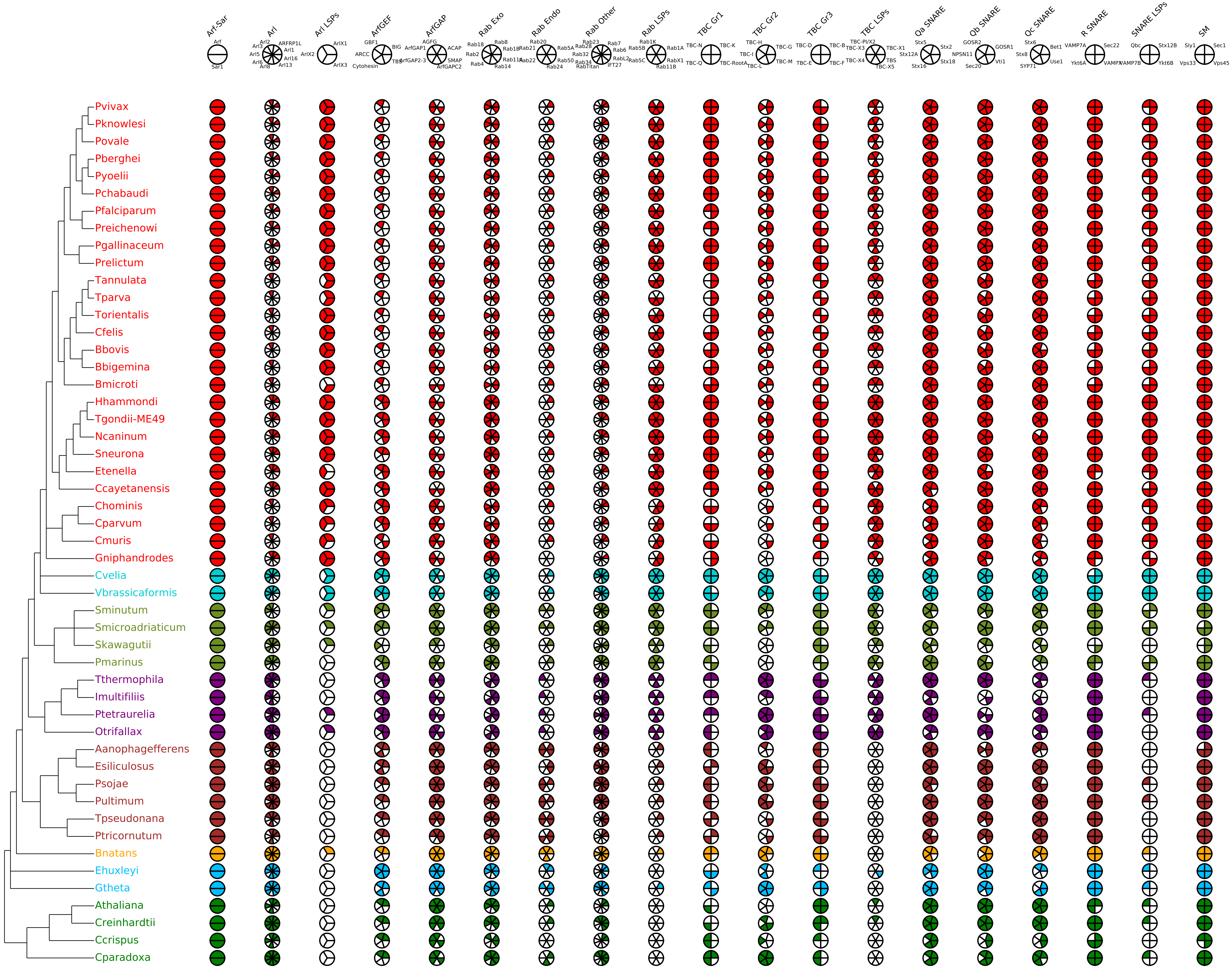

### Supplementary Figure 4

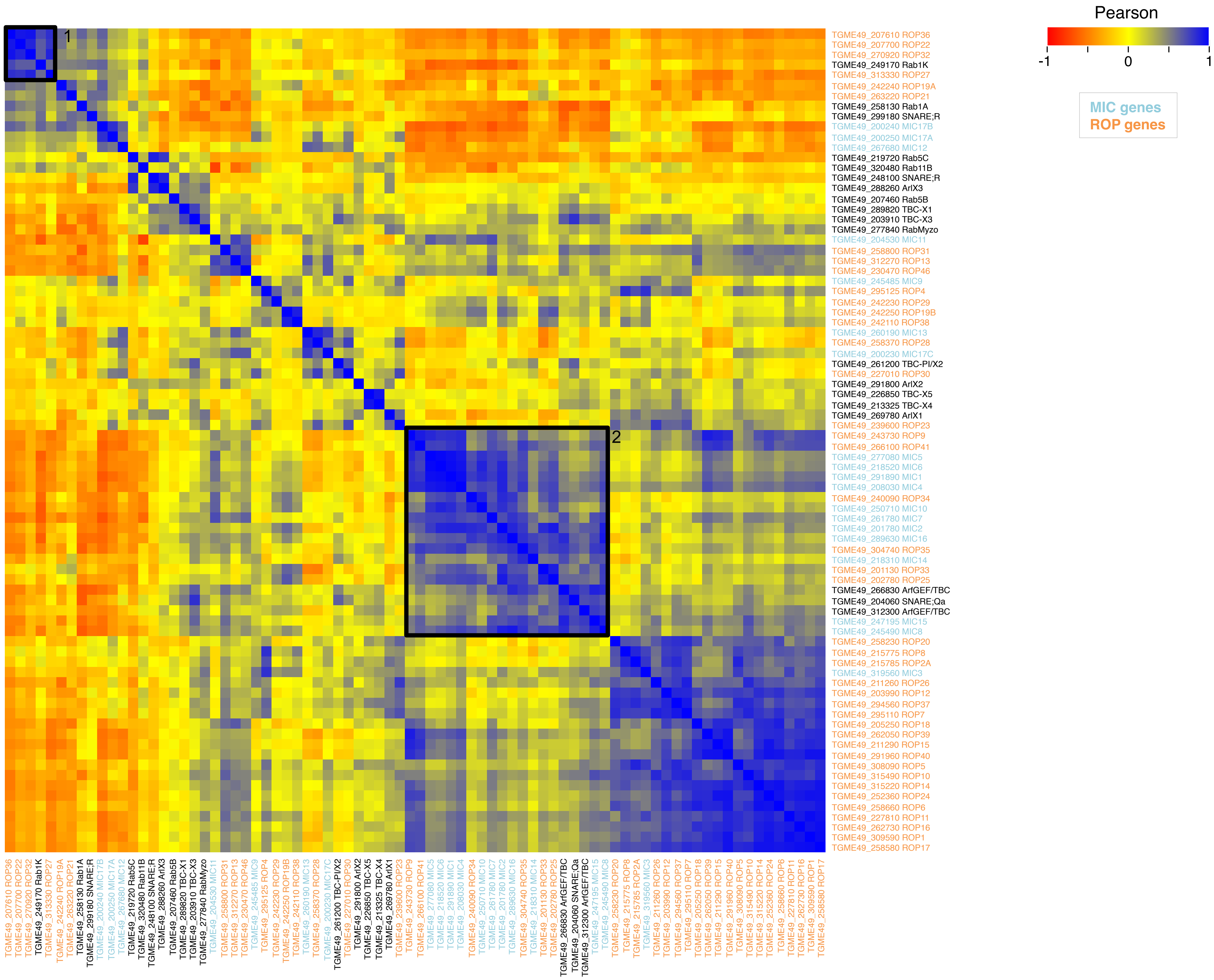

### Supplementary Figure 5

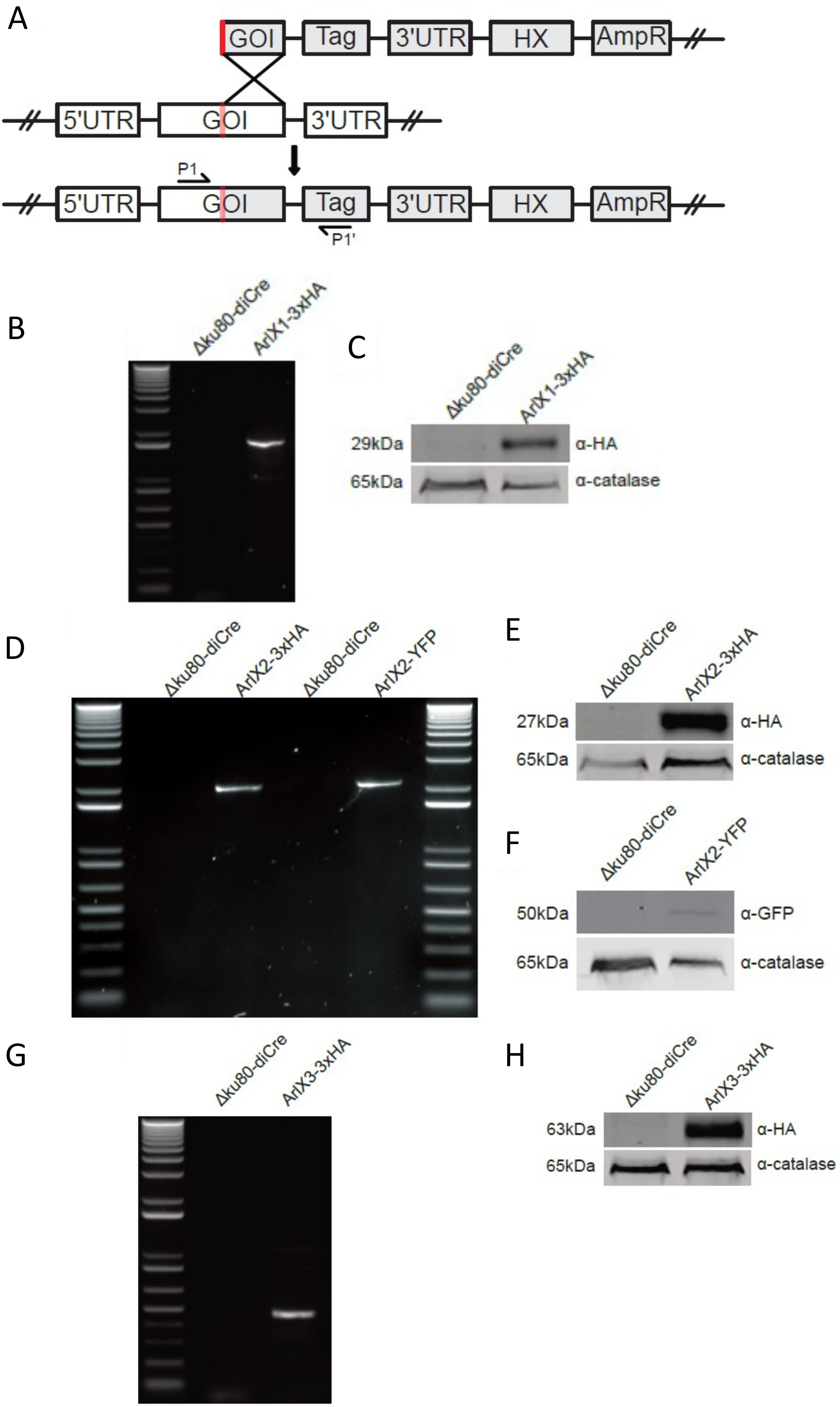

### Supplementary Figure 6

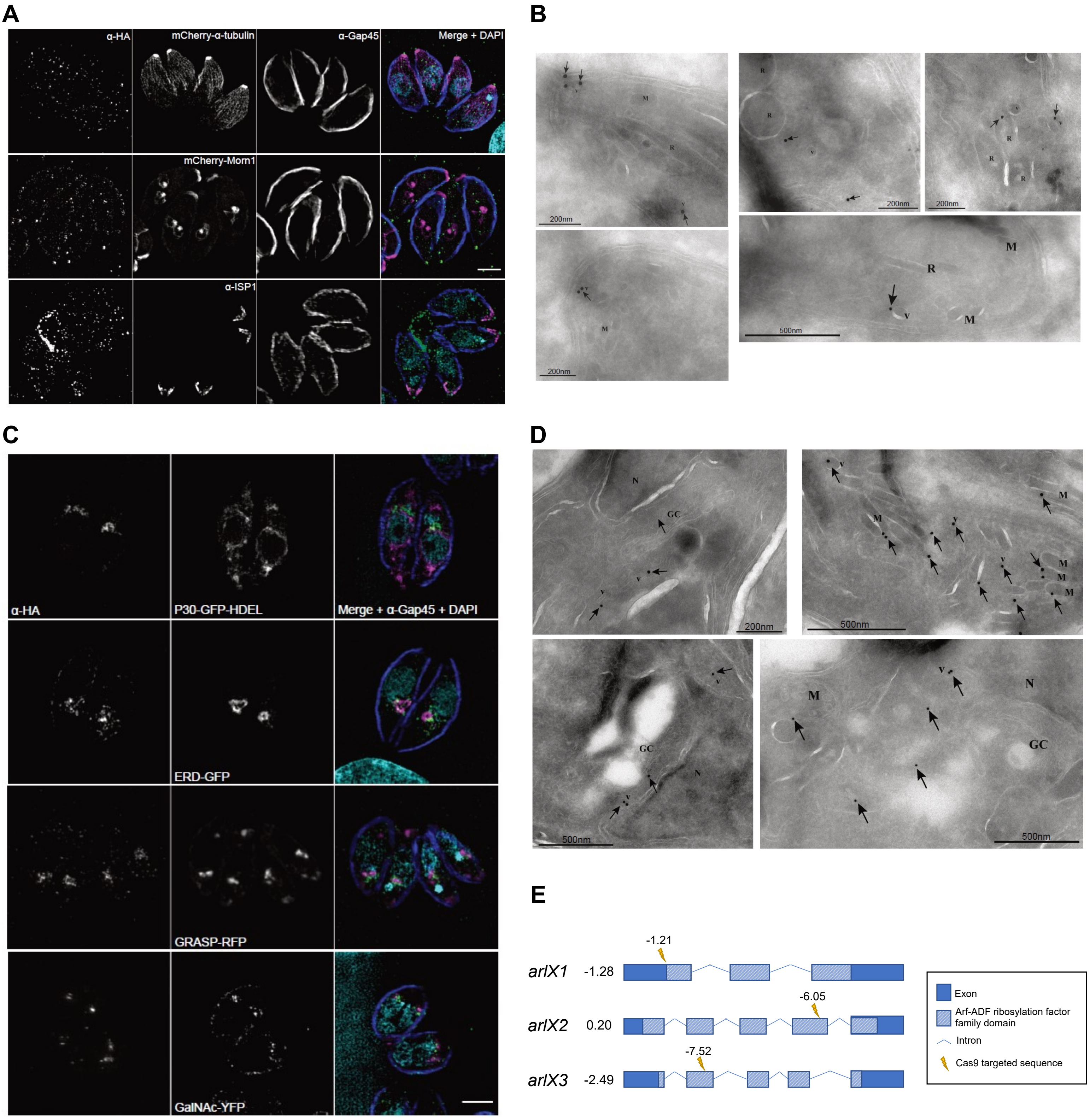

### Supplementary Figure 7

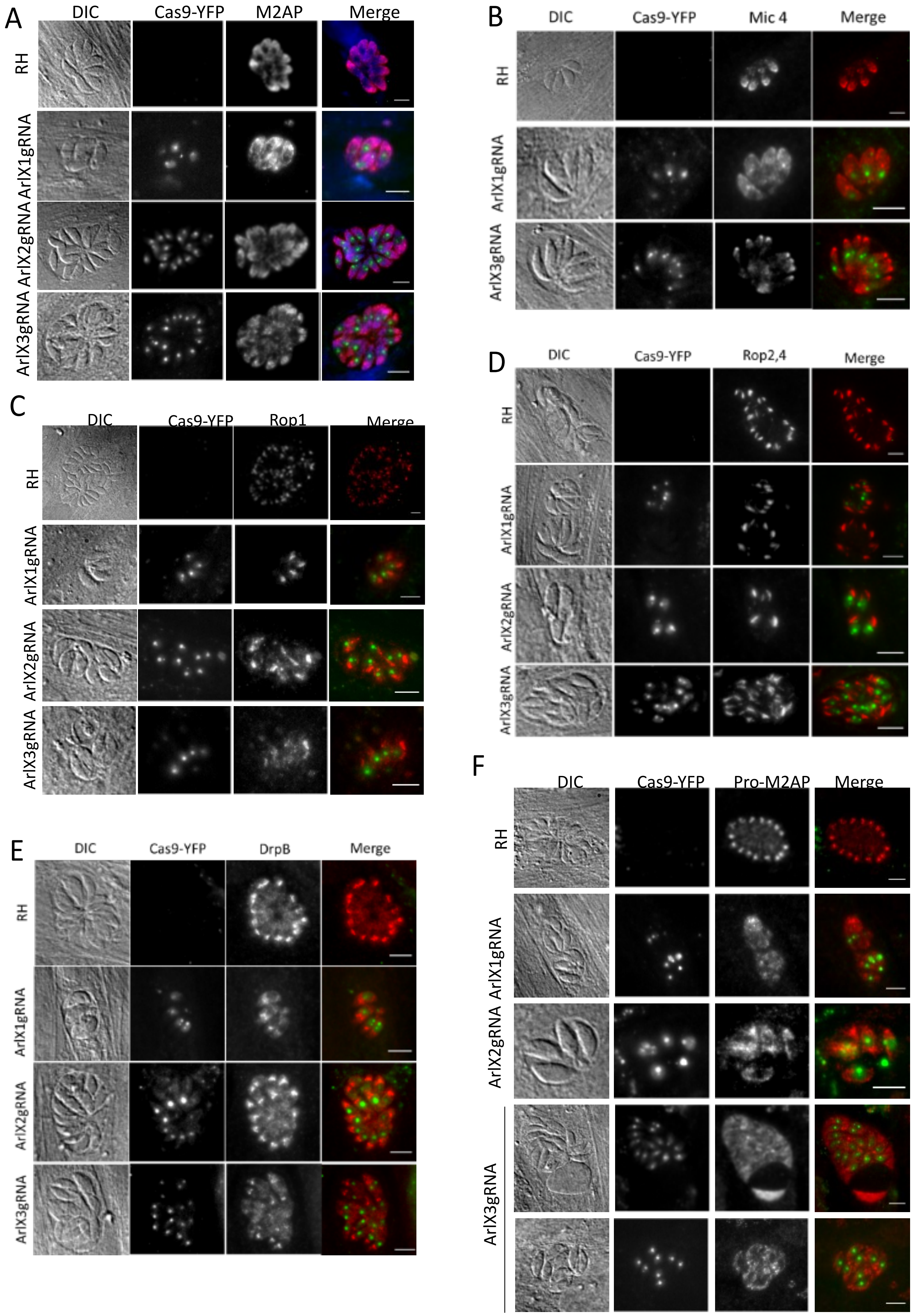

### Supplementary Figure 8

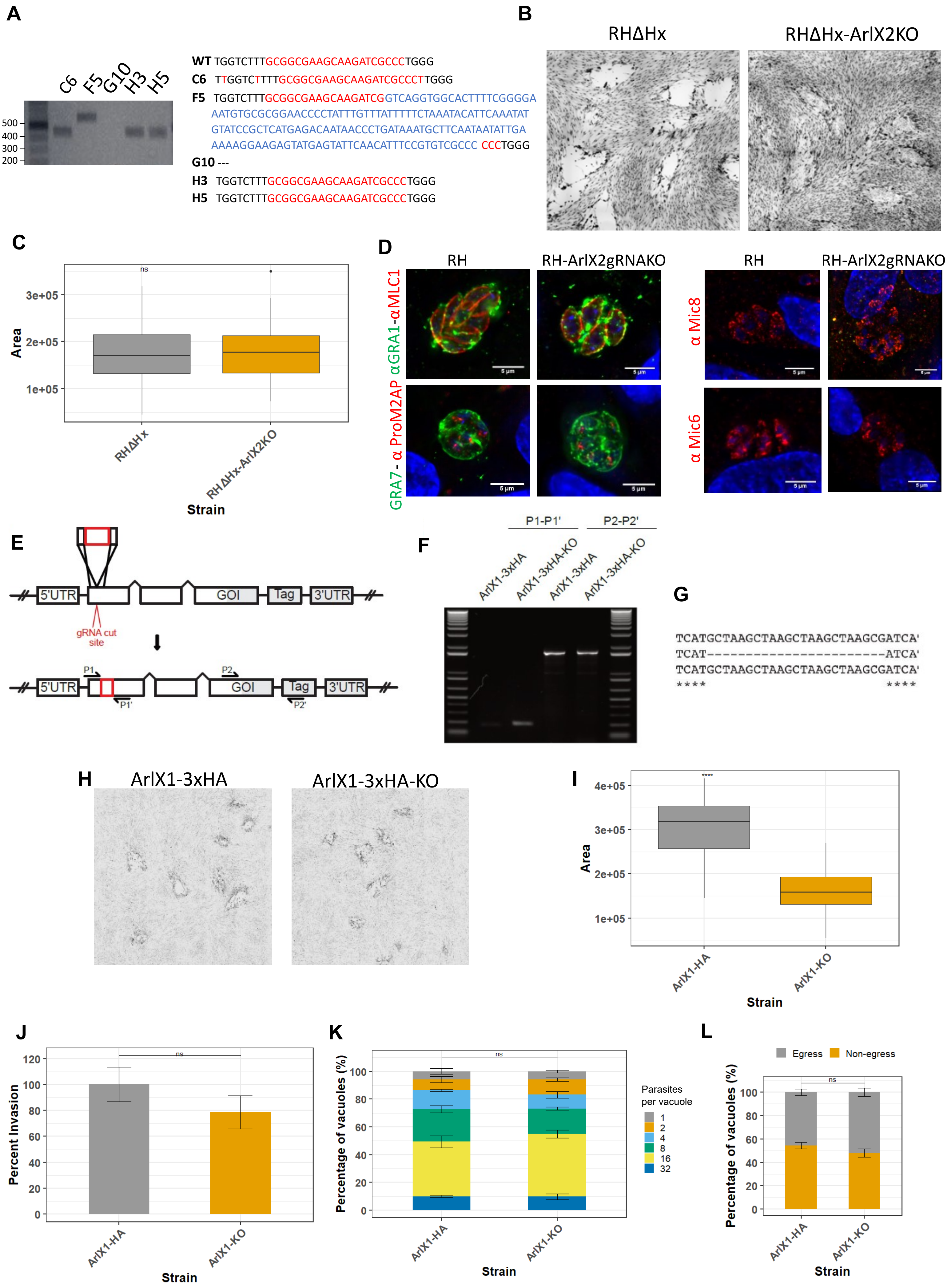

### Supplementary Figure 9

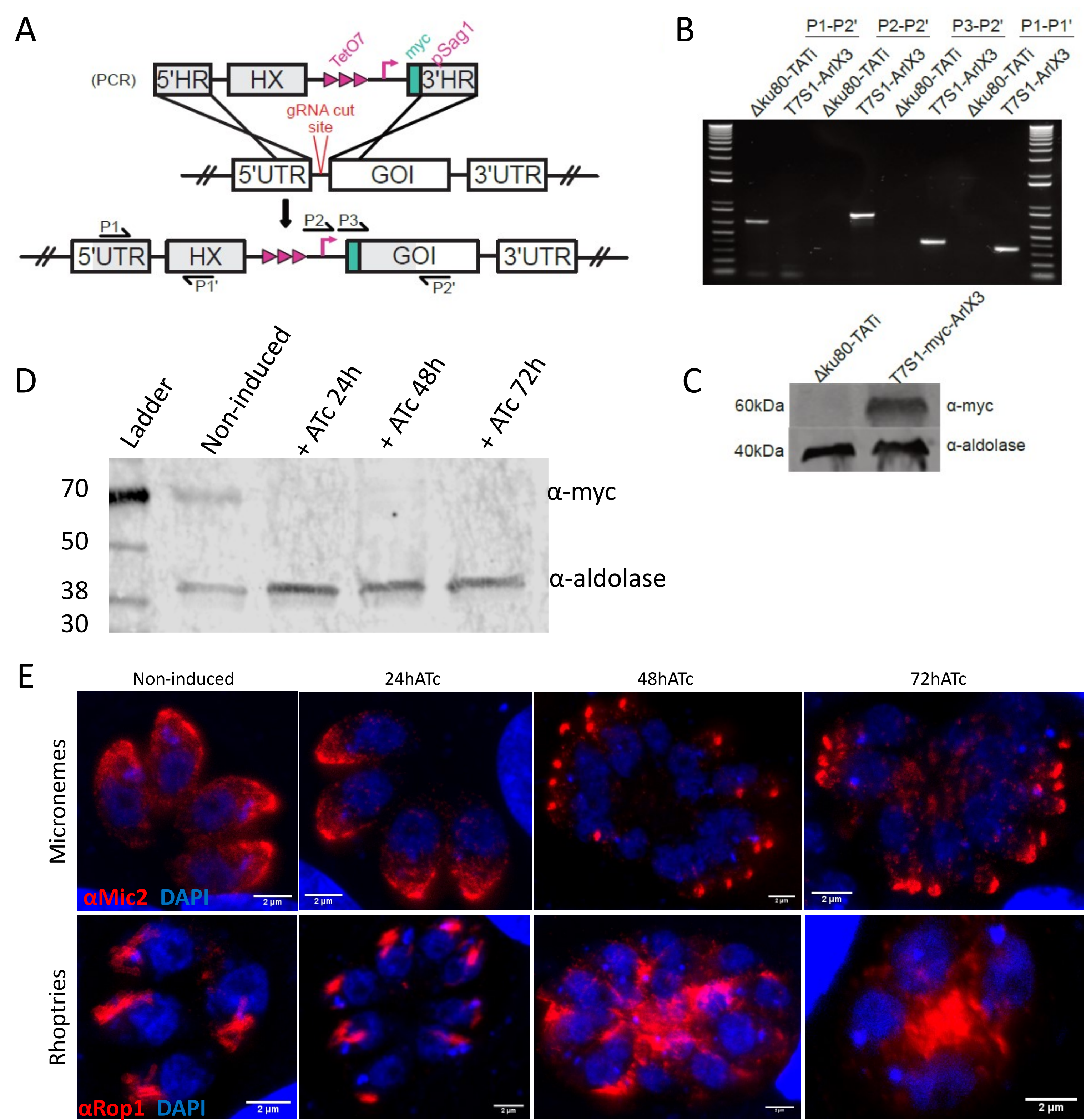

### Supplementary Figure 10

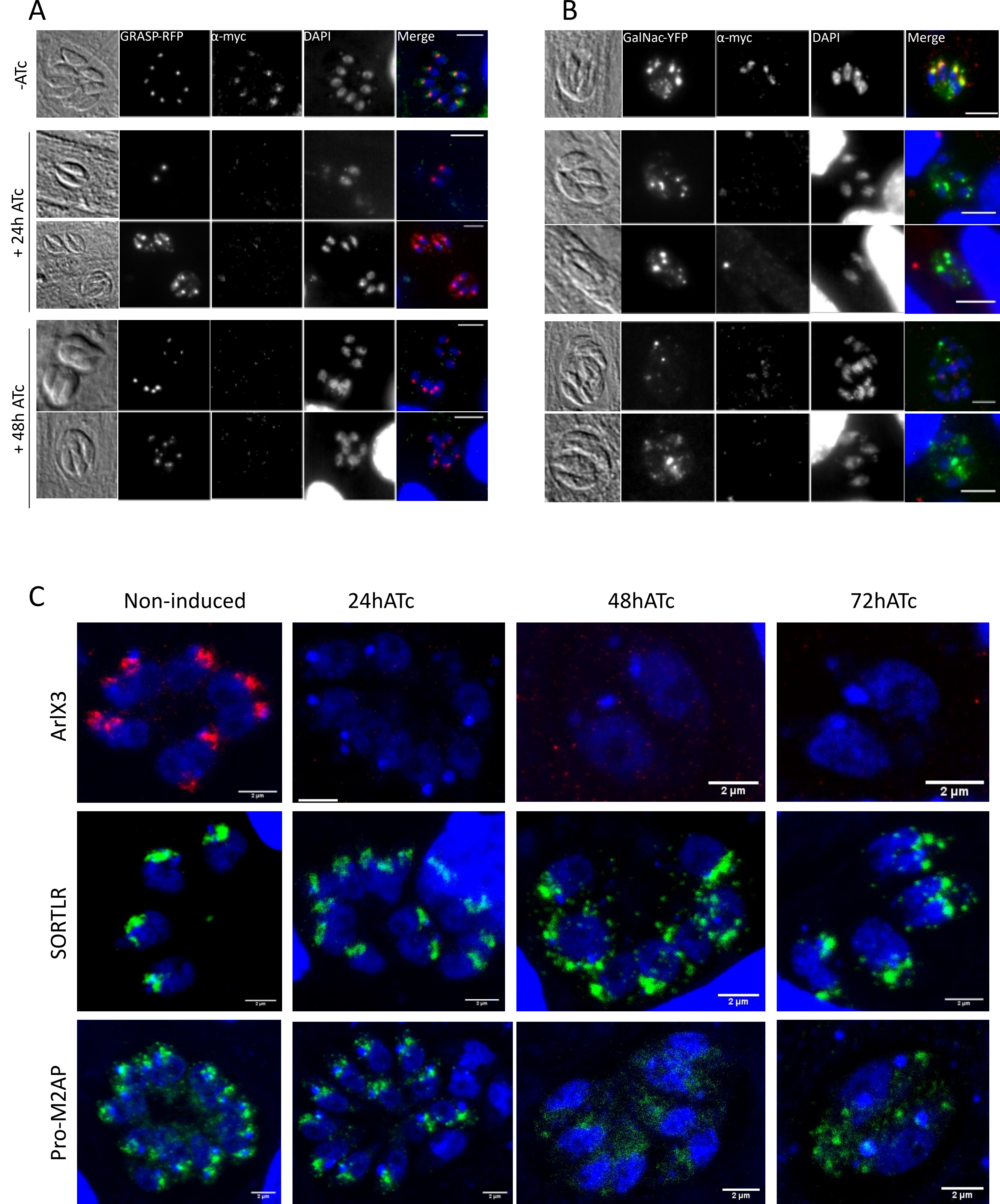
