## Supplementary Figure 2 for "Evolution of lineage-specific trafficking proteins and a novel post-Golgi trafficking pathway in Apicomplexa"

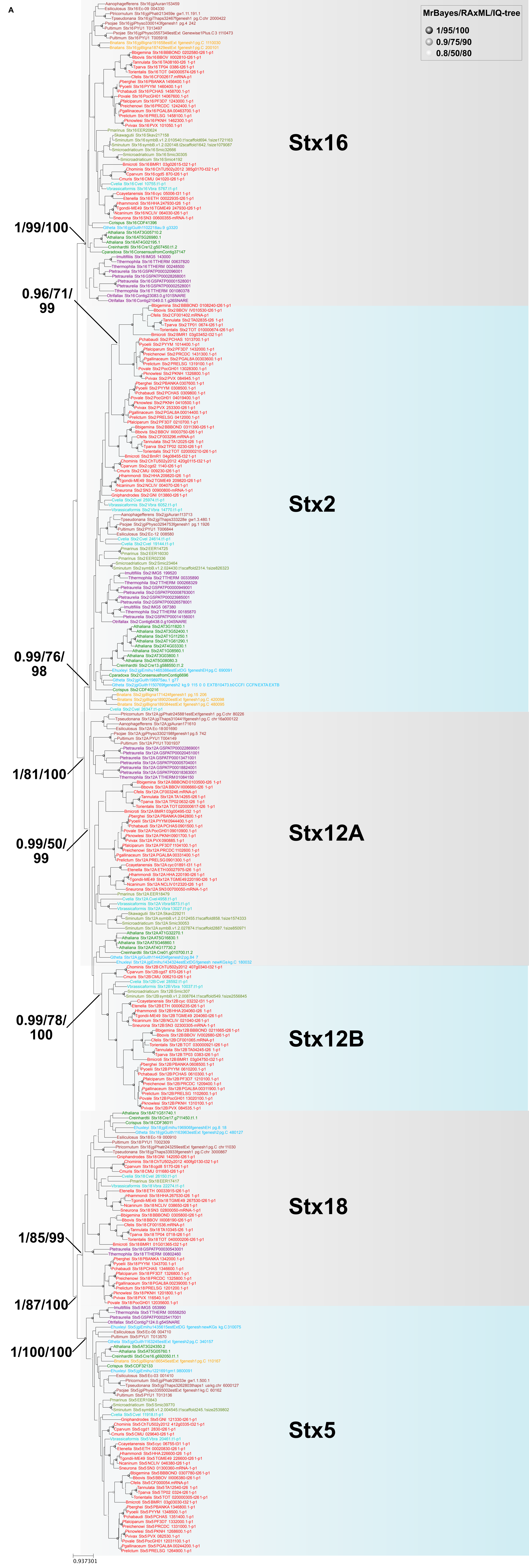
A
MrBayes/RAxML/IQ-tree
1/95/100
0.9/75/90
0.8/50/80
Stx16
Stx2
Stx12A
Stx12B
Stx18
Stx5
1/99/100
0.96/71/99
0.99/76/98
1/81/100
0.99/50/99
0.99/78/100
1/85/99
1/87/100
1/100/100
0.937301

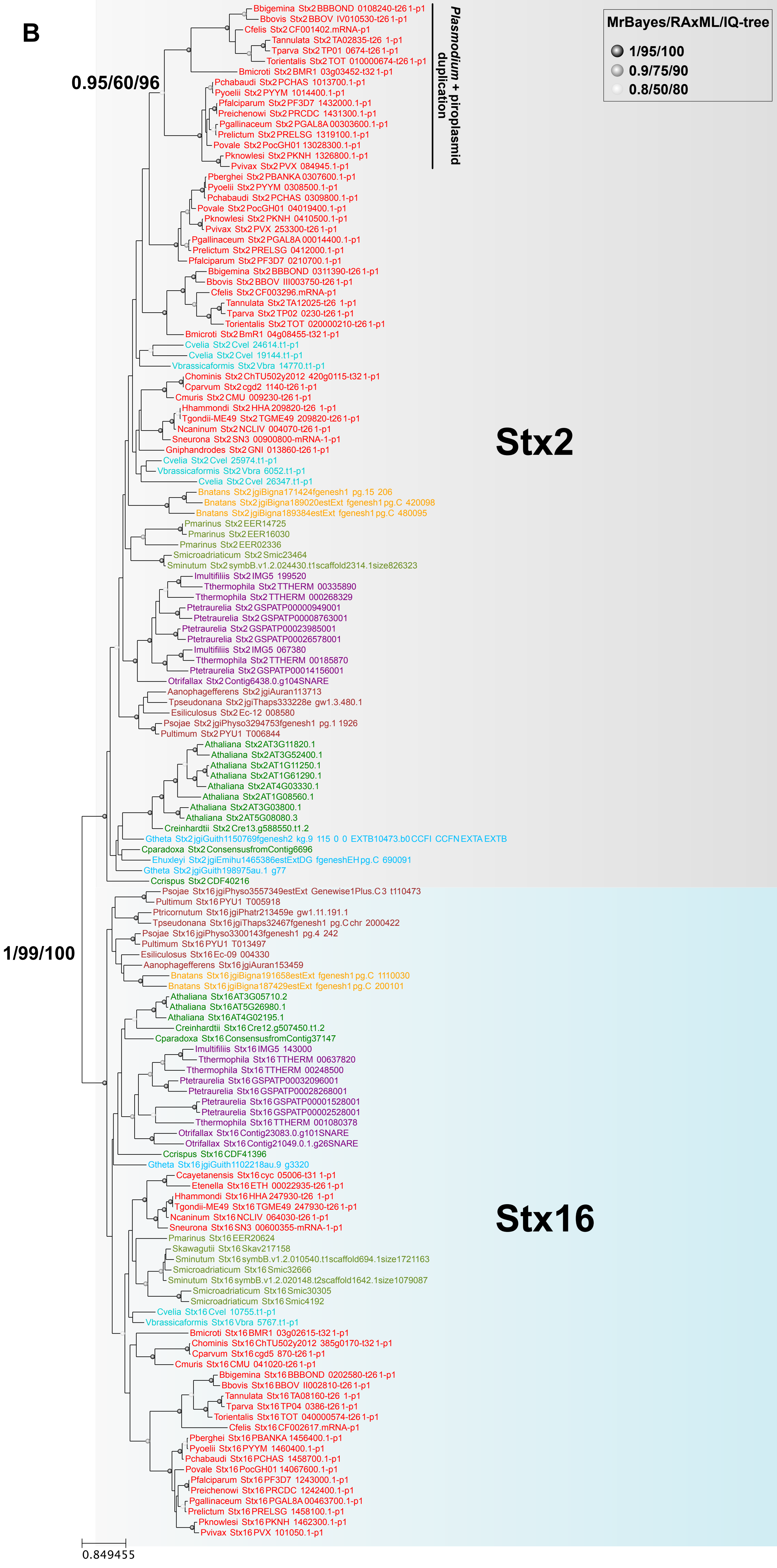

B
MrBayes/RAxML/IQ-tree
1/95/100
0.9/75/90
0.8/50/80
0.95/60/96
1/99/100
Plasmodium + piroplasmid duplication
Stx2
Stx16
0.849455
Bbigemina Stx2 BBBOND 0108240-t26 1-p1
Bbovis Stx2 BBOV IV010530-t26 1-p1
Cfelis Stx2 CF001402.mRNA-p1
Tannulata Stx2 TA02835-t26 1-p1
Tparva Stx2 TP01 0674-t26 1-p1
Torientalis Stx2 TOT 010000674-t26 1-p1
Bmicroti Stx2 BMR1 03g03452-t32 1-p1
Pchabaudi Stx2 PCHAS 1013700.1-p1
Pyoelii Stx2 PYYM 1014400.1-p1
Pfalciparum Stx2 PF3D7 1432000.1-p1
Preichenowi Stx2 PRCDC 1431300.1-p1
Pgallinaceum Stx2 PGAL8A 00303600.1-p1
Prelictum Stx2 PRELSG 1319100.1-p1
Povale Stx2 PocGH01 13028300.1-p1
Pknowlesi Stx2 PKNH 1326800.1-p1
Pvivax Stx2 PVX 084945.1-p1
Pberghei Stx2 PBANKA 0307600.1-p1
Pyoelii Stx2 PYYM 0308500.1-p1
Pchabaudi Stx2 PCHAS 0309800.1-p1
Povale Stx2 PocGH01 04019400.1-p1
Pknowlesi Stx2 PKNH 0410500.1-p1
Pvivax Stx2 PVX 253300-t26 1-p1
Pgallinaceum Stx2 PGAL8A 00014400.1-p1
Prelictum Stx2 PRELSG 0412000.1-p1
Pfalciparum Stx2 PF3D7 0210700.1-p1
Bbigemina Stx2 BBBOND 0311390-t26 1-p1
Bbovis Stx2 BBOV III003750-t26 1-p1
Cfelis Stx2 CF003296.mRNA-p1
Tannulata Stx2 TA12025-t26 1-p1
Tparva Stx2 TP02 0230-t26 1-p1
Torientalis Stx2 TOT 020000210-t26 1-p1
Bmicroti Stx2 BmR1 04g08455-t32 1-p1
Cvelia Stx2 Cvel 24614.t1-p1
Cvelia Stx2 Cvel 19144.t1-p1
Vbrassicaformis Stx2 Vbra 14770.t1-p1
Chominis Stx2 ChTU502y2012 420g0115-t32 1-p1
Cparvum Stx2 cgd2 1140-t26 1-p1
Cmuris Stx2 CMU 009230-t26 1-p1
Hhammondi Stx2 HHA 209820-t26 1-p1
Tgondii-ME49 Stx2 TGME49 209820-t26 1-p1
Ncaninum Stx2 NCLIV 004070-t26 1-p1
Sneurona Stx2 SN3 00900800-mRNA-1-p1
Gniphandrodes Stx2 GNI 013860-t26 1-p1
Cvelia Stx2 Cvel 25974.t1-p1
Vbrassicaformis Stx2 Vbra 6052.t1-p1
Cvelia Stx2 Cvel 26347.t1-p1
Bnatans Stx2 jgiBigna171424fgenesh1 pg.15 206
Bnatans Stx2 jgiBigna189020estExt fgenesh1 pg.C 420098
Bnatans Stx2 jgiBigna189384estExt fgenesh1 pg.C 480095
Pmarinus Stx2 EER14725
Pmarinus Stx2 EER16030
Pmarinus Stx2 EER02336
Smicroadriaticum Stx2 Smic23464
Sminutum Stx2 symbB.v1.2.024430.t1scaffold2314.1size826323
Imultifiliis Stx2 IMG5 199520
Tthermophila Stx2 TTHERM 00335890
Tthermophila Stx2 TTHERM 000268329
Ptetraurelia Stx2 GSPATP00000949001
Ptetraurelia Stx2 GSPATP00008763001
Ptetraurelia Stx2 GSPATP00023985001
Ptetraurelia Stx2 GSPATP00026578001
Imultifiliis Stx2 IMG5 067380
Tthermophila Stx2 TTHERM 00185870
Ptetraurelia Stx2 GSPATP00014156001
Otrifallax Stx2 Contig6438.0.g104SNARE
Aanophagefferens Stx2 jgiAuran113713
Tpseudonana Stx2 jgiThaps333228e gw1.3.480.1
Esiliculosus Stx2 Ec-12 008580
Psojae Stx2 jgiPhyso3294753fgenesh1 pg.1 1926
Pultimum Stx2 PYU1 T006844
Athaliana Stx2 AT3G11820.1
Athaliana Stx2 AT3G52400.1
Athaliana Stx2 AT1G11250.1
Athaliana Stx2 AT1G61290.1
Athaliana Stx2 AT4G03330.1
Athaliana Stx2 AT1G08560.1
Athaliana Stx2 AT3G03800.1
Athaliana Stx2 AT5G08080.3
Creinhardtii Stx2 Cre13.g588550.t1.2
Gtheta Stx2 jgiGuith1150769fgenesh2 kg.9 115 0 0 EXTB10473.b0 CCFI CCFN EXTA EXTB
Cparadoxa Stx2 ConsensusfromContig6696
Ehuxleyi Stx2 jgiEmihu1465386estExtDG fgeneshEH pg.C 690091
Gtheta Stx2 jgiGuith198975au.1 g77
Ccrispus Stx2 CDF40216
Psojae Stx16 jgiPhyso3557349estExt Genewise1Plus.C3 t110473
Pultimum Stx16 PYU1 T005918
Ptricornutum Stx16 jgiPhatr213459e gw1.11.191.1
Tpseudonana Stx16 jgiThaps32467fgenesh1 pg.C chr 2000422
Psojae Stx16 jgiPhyso3300143fgenesh1 pg.4 242
Pultimum Stx16 PYU1 T013497
Esiliculosus Stx16 Ec-09 004330
Aanophagefferens Stx16 jgiAuran153459
Bnatans Stx16 jgiBigna191658estExt fgenesh1 pg.C 1110030
Bnatans Stx16 jgiBigna187429estExt fgenesh1 pg.C 200101
Athaliana Stx16 AT3G05710.2
Athaliana Stx16 AT5G26980.1
Athaliana Stx16 AT4G02195.1
Creinhardtii Stx16 Cre12.g507450.t1.2
Cparadoxa Stx16 ConsensusfromContig37147
Imultifiliis Stx16 IMG5 143000
Tthermophila Stx16 TTHERM 00637820
Tthermophila Stx16 TTHERM 00248500
Ptetraurelia Stx16 GSPATP00032096001
Ptetraurelia Stx16 GSPATP00028268001
Ptetraurelia Stx16 GSPATP00001528001
Ptetraurelia Stx16 GSPATP00002528001
Tthermophila Stx16 TTHERM 001080378
Otrifallax Stx16 Contig23083.0.g101SNARE
Otrifallax Stx16 Contig21049.0.1.g26SNARE
Ccrispus Stx16 CDF41396
Gtheta Stx16 jgiGuith1102218au.9 g3320
Ccayetanensis Stx16 cyc 05006-t31 1-p1
Etenella Stx16 ETH 00022935-t26 1-p1
Hhammondi Stx16 HHA 247930-t26 1-p1
Tgondii-ME49 Stx16 TGME49 247930-t26 1-p1
Ncaninum Stx16 NCLIV 064030-t26 1-p1
Sneurona Stx16 SN3 00600355-mRNA-1-p1
Pmarinus Stx16 EER20624
Skawagutii Stx16 Skav217158
Sminutum Stx16 symbB.v1.2.010540.t1scaffold694.1size1721163
Smicroadriaticum Stx16 Smic32666
Sminutum Stx16 symbB.v1.2.020148.t2scaffold1642.1size1079087
Smicroadriaticum Stx16 Smic30305
Smicroadriaticum Stx16 Smic4192
Cvelia Stx16 Cvel 10755.t1-p1
Vbrassicaformis Stx16 Vbra 5767.t1-p1
Bmicroti Stx16 BMR1 03g02615-t32 1-p1
Chominis Stx16 ChTU502y2012 385g0170-t32 1-p1
Cparvum Stx16 cgd5 870-t26 1-p1
Cmuris Stx16 CMU 041020-t26 1-p1
Bbigemina Stx16 BBBOND 0202580-t26 1-p1
Bbovis Stx16 BBOV II002810-t26 1-p1
Tannulata Stx16 TA08160-t26 1-p1
Tparva Stx16 TP04 0386-t26 1-p1
Torientalis Stx16 TOT 040000574-t26 1-p1
Cfelis Stx16 CF002617.mRNA-p1
Pberghei Stx16 PBANKA 1456400.1-p1
Pyoelii Stx16 PYYM 1460400.1-p1
Pchabaudi Stx16 PCHAS 1458700.1-p1
Povale Stx16 PocGH01 14067600.1-p1
Pfalciparum Stx16 PF3D7 1243000.1-p1
Preichenowi Stx16 PRCDC 1242400.1-p1
Pgallinaceum Stx16 PGAL8A 00463700.1-p1
Prelictum Stx16 PRELSG 1458100.1-p1
Pknowlesi Stx16 PKNH 1462300.1-p1
Pvivax Stx16 PVX 101050.1-p1

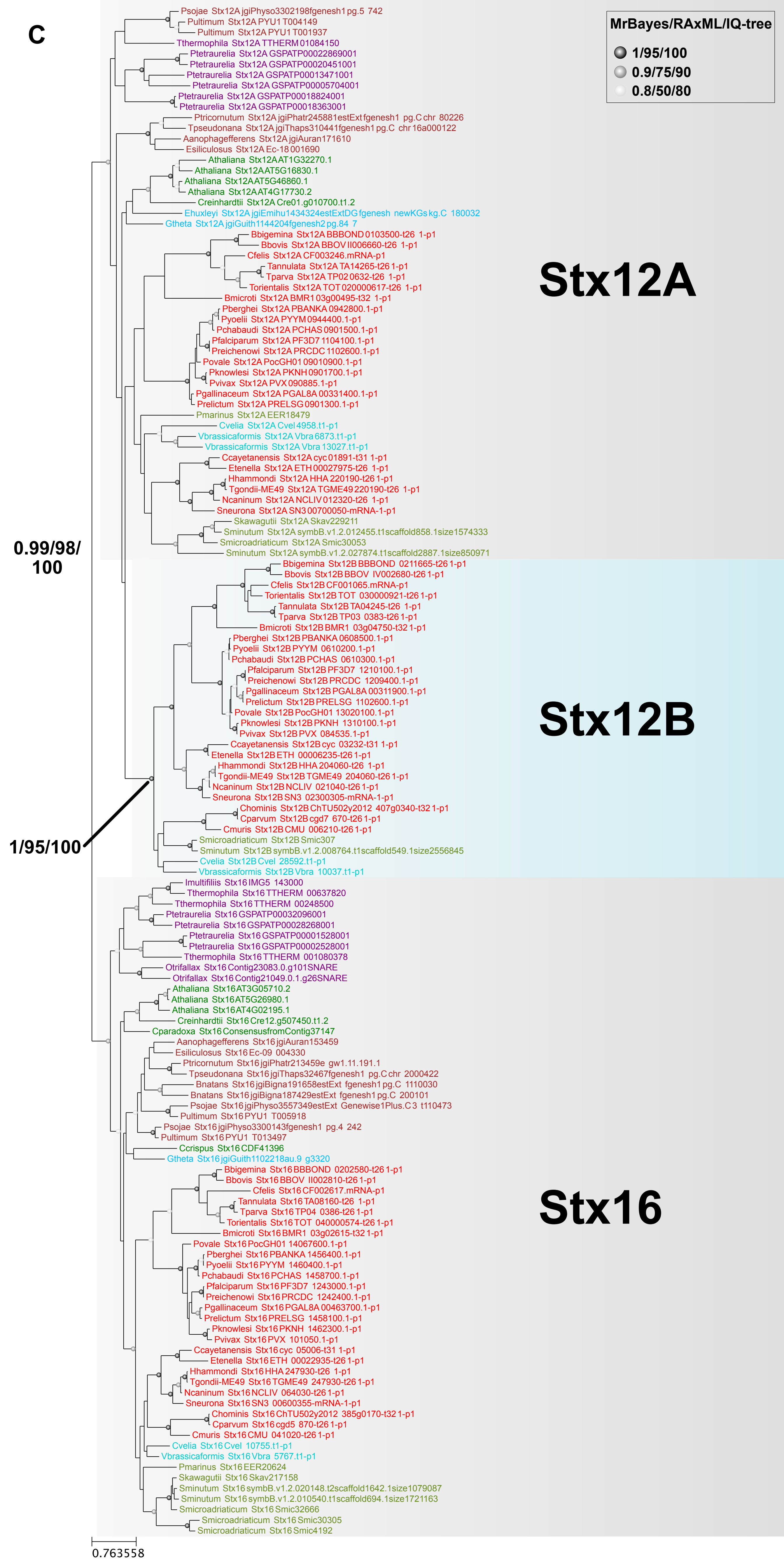

C
MrBayes/RAxML/IQ-tree
1/95/100
0.9/75/90
0.8/50/80
Psojae Stx12A jgiPhyso3302198fgenesh1 pg.5 742
Pultimum Stx12A PYU1 T004149
Pultimum Stx12A PYU1 T001937
Tthermophila Stx12A TTHERM 01084150
Ptetraurelia Stx12A GSPATP00022869001
Ptetraurelia Stx12A GSPATP00020451001
Ptetraurelia Stx12A GSPATP00013471001
Ptetraurelia Stx12A GSPATP00005704001
Ptetraurelia Stx12A GSPATP00018824001
Ptetraurelia Stx12A GSPATP00018363001
Ptricornutum Stx12A jgiPhatr245881estExtfgenesh1 pg.C chr 80226
Tpseudonana Stx12A jgiThaps310441fgenesh1 pg.C chr 16a000122
Aanophagefferens Stx12A jgiAuran171610
Esiliculosus Stx12A Ec-18 001690
Athaliana Stx12A AT1G32270.1
Athaliana Stx12A AT5G16830.1
Athaliana Stx12A AT5G46860.1
Athaliana Stx12A AT4G17730.2
Creinhardtii Stx12A Cre01.g010700.t1.2
Ehuxleyi Stx12A jgiEmihu1434324estExtDG fgenesh newKGs kg.C 180032
Gtheta Stx12A jgiGuith1144204fgenesh2 pg.84 7
Bbigemina Stx12A BBBOND 0103500-t26 1-p1
Bbovis Stx12A BBOV II006660-t26 1-p1
Cfelis Stx12A CF003246.mRNA-p1
Tannulata Stx12A TA14265-t26 1-p1
Tparva Stx12A TP02 0632-t26 1-p1
Torientalis Stx12A TOT 020000617-t26 1-p1
Bmicroti Stx12A BMR1 03g00495-t32 1-p1
Pberghei Stx12A PBANKA 0942800.1-p1
Pyoelii Stx12A PYYM 0944400.1-p1
Pchabaudi Stx12A PCHAS 0901500.1-p1
Pfalciparum Stx12A PF3D7 1104100.1-p1
Preichenowi Stx12A PRCDC 1102600.1-p1
Povale Stx12A PocGH01 09010900.1-p1
Pknowlesi Stx12A PKNH 0901700.1-p1
Pvivax Stx12A PVX 090885.1-p1
Pgallinaceum Stx12A PGAL8A 00331400.1-p1
Prelictum Stx12A PRELSG 0901300.1-p1
Pmarinus Stx12A EER18479
Cvelia Stx12A Cvel 4958.t1-p1
Vbrassicaformis Stx12A Vbra 6873.t1-p1
Vbrassicaformis Stx12A Vbra 13027.t1-p1
Ccayetanensis Stx12A cyc 01891-t31 1-p1
Etenella Stx12A ETH 00027975-t26 1-p1
Hhammondi Stx12A HHA 220190-t26 1-p1
Tgondii-ME49 Stx12A TGME49 220190-t26 1-p1
Ncaninum Stx12A NCLIV 012320-t26 1-p1
Sneurona Stx12A SN3 00700050-mRNA-1-p1
Skawagutii Stx12A Skav229211
Sminutum Stx12A symbB.v1.2.012455.t1scaffold858.1size1574333
Smicroadriaticum Stx12A Smic30053
Sminutum Stx12A symbB.v1.2.027874.t1scaffold2887.1size850971
Stx12A
0.99/98/100
Bbigemina Stx12B BBBOND 0211665-t26 1-p1
Bbovis Stx12B BBOV IV002680-t26 1-p1
Cfelis Stx12B CF001065.mRNA-p1
Torientalis Stx12B TOT 030000921-t26 1-p1
Tannulata Stx12B TA04245-t26 1-p1
Tparva Stx12B TP03 0383-t26 1-p1
Bmicroti Stx12B BMR1 03g04750-t32 1-p1
Pberghei Stx12B PBANKA 0608500.1-p1
Pyoelii Stx12B PYYM 0610200.1-p1
Pchabaudi Stx12B PCHAS 0610300.1-p1
Pfalciparum Stx12B PF3D7 1210100.1-p1
Preichenowi Stx12B PRCDC 1209400.1-p1
Pgallinaceum Stx12B PGAL8A 00311900.1-p1
Prelictum Stx12B PRELSG 1102600.1-p1
Povale Stx12B PocGH01 13020100.1-p1
Pknowlesi Stx12B PKNH 1310100.1-p1
Pvivax Stx12B PVX 084535.1-p1
Ccayetanensis Stx12B cyc 03232-t31 1-p1
Etenella Stx12B ETH 00006235-t26 1-p1
Hhammondi Stx12B HHA 204060-t26 1-p1
Tgondii-ME49 Stx12B TGME49 204060-t26 1-p1
Ncaninum Stx12B NCLIV 021040-t26 1-p1
Sneurona Stx12B SN3 02300305-mRNA-1-p1
Chominis Stx12B ChTU502y2012 407g0340-t32 1-p1
Cparvum Stx12B cgd7 670-t26 1-p1
Cmuris Stx12B CMU 006210-t26 1-p1
Smicroadriaticum Stx12B Smic307
Sminutum Stx12B symbB.v1.2.008764.t1scaffold549.1size2556845
Cvelia Stx12B Cvel 28592.t1-p1
Vbrassicaformis Stx12B Vbra 10037.t1-p1
Stx12B
1/95/100
Imultifiliis Stx16 IMG5 143000
Tthermophila Stx16 TTHERM 00637820
Tthermophila Stx16 TTHERM 00248500
Ptetraurelia Stx16 GSPATP00032096001
Ptetraurelia Stx16 GSPATP00028268001
Ptetraurelia Stx16 GSPATP00001528001
Ptetraurelia Stx16 GSPATP00002528001
Tthermophila Stx16 TTHERM 001080378
Otrifallax Stx16 Contig23083.0.g101SNARE
Otrifallax Stx16 Contig21049.0.1.g26SNARE
Athaliana Stx16 AT3G05710.2
Athaliana Stx16 AT5G26980.1
Athaliana Stx16 AT4G02195.1
Creinhardtii Stx16 Cre12.g507450.t1.2
Cparadoxa Stx16 ConsensusfromContig37147
Aanophagefferens Stx16 jgiAuran153459
Esiliculosus Stx16 Ec-09 004330
Ptricornutum Stx16 jgiPhatr213459e gw1.11.191.1
Tpseudonana Stx16 jgiThaps32467fgenesh1 pg.C chr 2000422
Bnatans Stx16 jgiBigna191658estExt fgenesh1 pg.C 1110030
Bnatans Stx16 jgiBigna187429estExt fgenesh1 pg.C 200101
Psojae Stx16 jgiPhyso3557349estExt Genewise1Plus.C3 t110473
Pultimum Stx16 PYU1 T005918
Psojae Stx16 jgiPhyso3300143fgenesh1 pg.4 242
Pultimum Stx16 PYU1 T013497
Ccrispus Stx16 CDF41396
Gtheta Stx16 jgiGuith1102218au.9 g3320
Bbigemina Stx16 BBBOND 0202580-t26 1-p1
Bbovis Stx16 BBOV II002810-t26 1-p1
Cfelis Stx16 CF002617.mRNA-p1
Tannulata Stx16 TA08160-t26 1-p1
Tparva Stx16 TP04 0386-t26 1-p1
Torientalis Stx16 TOT 040000574-t26 1-p1
Bmicroti Stx16 BMR1 03g02615-t32 1-p1
Povale Stx16 PocGH01 14067600.1-p1
Pberghei Stx16 PBANKA 1456400.1-p1
Pyoelii Stx16 PYYM 1460400.1-p1
Pchabaudi Stx16 PCHAS 1458700.1-p1
Pfalciparum Stx16 PF3D7 1243000.1-p1
Preichenowi Stx16 PRCDC 1242400.1-p1
Pgallinaceum Stx16 PGAL8A 00463700.1-p1
Prelictum Stx16 PRELSG 1458100.1-p1
Pknowlesi Stx16 PKNH 1462300.1-p1
Pvivax Stx16 PVX 101050.1-p1
Ccayetanensis Stx16 cyc 05006-t31 1-p1
Etenella Stx16 ETH 00022935-t26 1-p1
Hhammondi Stx16 HHA 247930-t26 1-p1
Tgondii-ME49 Stx16 TGME49 247930-t26 1-p1
Ncaninum Stx16 NCLIV 064030-t26 1-p1
Sneurona Stx16 SN3 00600355-mRNA-1-p1
Chominis Stx16 ChTU502y2012 385g0170-t32 1-p1
Cparvum Stx16 cgd5 870-t26 1-p1
Cmuris Stx16 CMU 041020-t26 1-p1
Cvelia Stx16 Cvel 10755.t1-p1
Vbrassicaformis Stx16 Vbra 5767.t1-p1
Pmarinus Stx16 EER20624
Skawagutii Stx16 Skav217158
Sminutum Stx16 symbB.v1.2.020148.t2scaffold1642.1size1079087
Sminutum Stx16 symbB.v1.2.010540.t1scaffold694.1size1721163
Smicroadriaticum Stx16 Smic32666
Smicroadriaticum Stx16 Smic30305
Smicroadriaticum Stx16 Smic4192
Stx16
0.763558

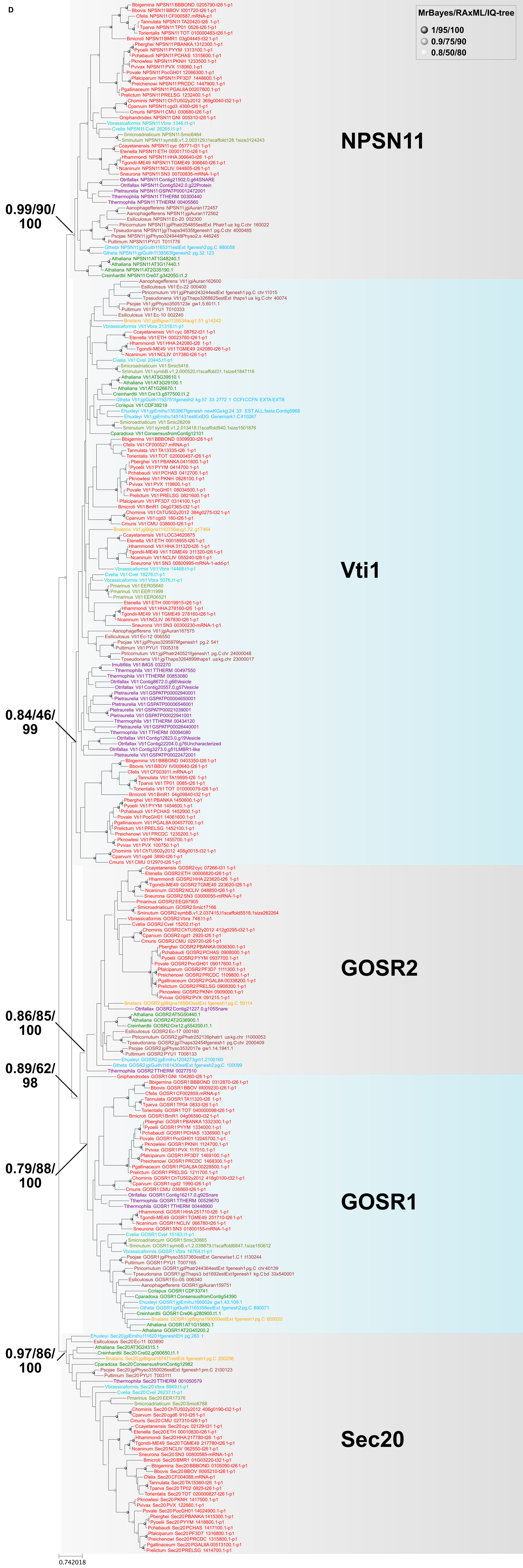
D
MrBayes/RAxML/IQ-tree
1/95/100
0.9/75/90
0.8/50/80
NPSN11
0.99/90/
100
Vti1
0.84/46/
99
GOSR2
0.86/85/
100
0.89/62/
98
0.79/88/
100
GOSR1
0.97/86/
100
Sec20
0.742018

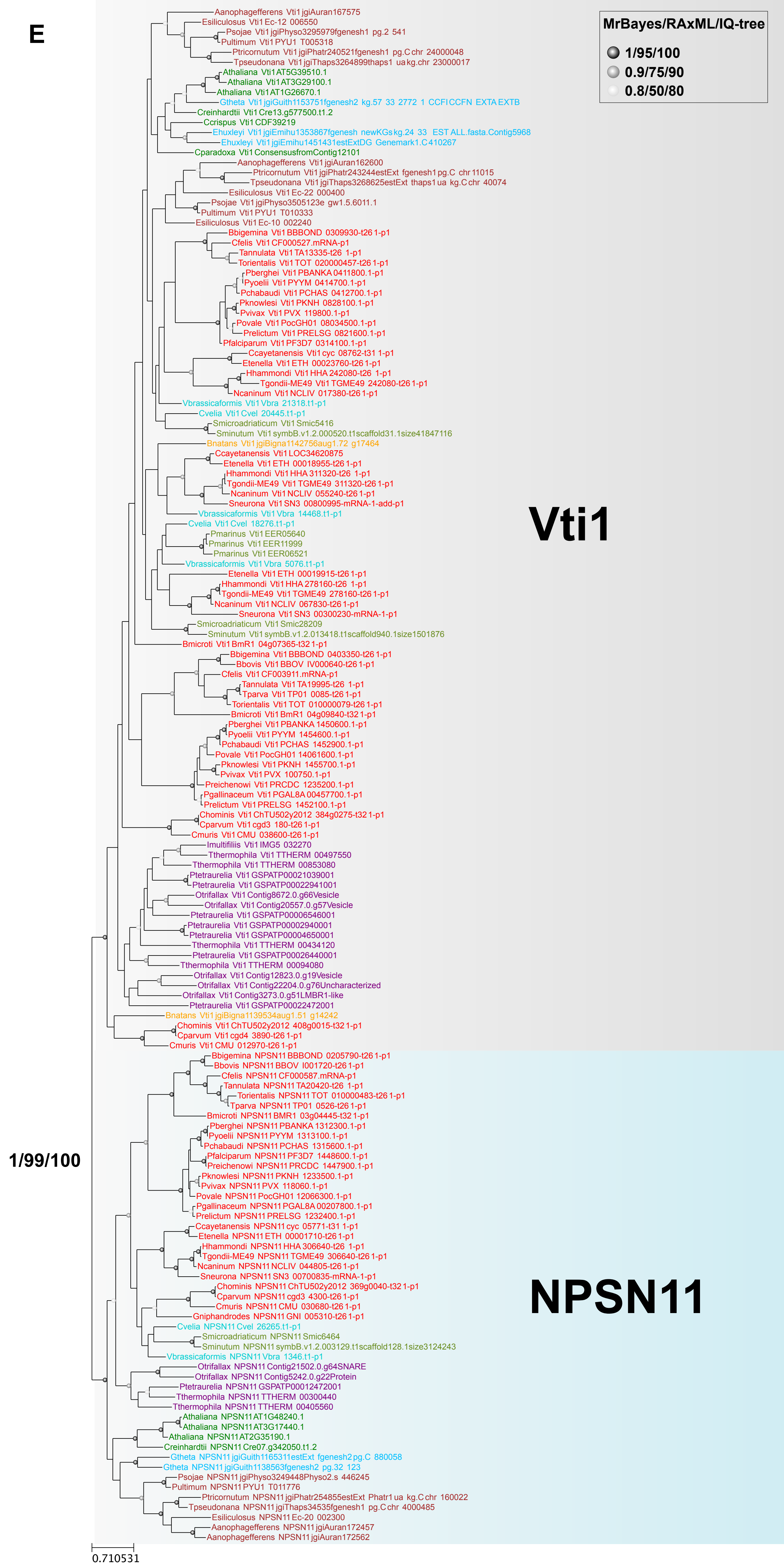
E
MrBayes/RAxML/IQ-tree
1/95/100
0.9/75/90
0.8/50/80
Vti1
NPSN11
1/99/100
0.710531
Aanophagefferens Vti1 jgiAuran167575
Esiliculosus Vti1 Ec-12 006550
Psojae Vti1 jgiPhyso3295979fgenesh1 pg.2 541
Pultimum Vti1 PYU1 T005318
Ptricornutum Vti1 jgiPhatr240521fgenesh1 pg.C chr 24000048
Tpseudonana Vti1 jgiThaps3264899thaps1 ua kg.chr 23000017
Athaliana Vti1 AT5G39510.1
Athaliana Vti1 AT3G29100.1
Athaliana Vti1 AT1G26670.1
Gtheta Vti1 jgiGuith1153751fgenesh2 kg.57 33 2772 1 CCFI CCFN EXTA EXTB
Creinhardtii Vti1 Cre13.g577500.t1.2
Ccrispus Vti1 CDF39219
Ehuxleyi Vti1 jgiEmihu1353867fgenesh newKGs kg.24 33 EST ALL.fasta.Contig5968
Ehuxleyi Vti1 jgiEmihu1451431estExtDG Genemark1.C410267
Cparadoxa Vti1 ConsensusfromContig12101
Aanophagefferens Vti1 jgiAuran162600
Ptricornutum Vti1 jgiPhatr243244estExt fgenesh1 pg.C chr 11015
Tpseudonana Vti1 jgiThaps3268625estExt thaps1 ua kg.C chr 40074
Esiliculosus Vti1 Ec-22 000400
Psojae Vti1 jgiPhyso3505123e gw1.5.6011.1
Pultimum Vti1 PYU1 T010333
Esiliculosus Vti1 Ec-10 002240
Bbigemina Vti1 BBBOND 0309930-t26 1-p1
Cfelis Vti1 CF000527.mRNA-p1
Tannulata Vti1 TA13335-t26 1-p1
Torientalis Vti1 TOT 020000457-t26 1-p1
Pberghei Vti1 PBANKA 0411800.1-p1
Pyoelii Vti1 PYYM 0414700.1-p1
Pchabaudi Vti1 PCHAS 0412700.1-p1
Pknowlesi Vti1 PKNH 0828100.1-p1
Pvivax Vti1 PVX 119800.1-p1
Povale Vti1 PocGH01 08034500.1-p1
Prelictum Vti1 PRELSG 0821600.1-p1
Pfalciparum Vti1 PF3D7 0314100.1-p1
Ccayetanensis Vti1 cyc 08762-t31 1-p1
Etenella Vti1 ETH 00023760-t26 1-p1
Hhammondi Vti1 HHA 242080-t26 1-p1
Tgondii-ME49 Vti1 TGME49 242080-t26 1-p1
Ncaninum Vti1 NCLIV 017380-t26 1-p1
Vbrassicaformis Vti1 Vbra 21318.t1-p1
Cvelia Vti1 Cvel 20445.t1-p1
Smicroadriaticum Vti1 Smic5416
Sminutum Vti1 symbB.v1.2.000520.t1scaffold31.1size41847116
Bnatans Vti1 jgiBigna1142756aug1.72 g17464
Ccayetanensis Vti1 LOC34620875
Etenella Vti1 ETH 00018955-t26 1-p1
Hhammondi Vti1 HHA 311320-t26 1-p1
Tgondii-ME49 Vti1 TGME49 311320-t26 1-p1
Ncaninum Vti1 NCLIV 055240-t26 1-p1
Sneurona Vti1 SN3 00800995-mRNA-1-add-p1
Vbrassicaformis Vti1 Vbra 14468.t1-p1
Cvelia Vti1 Cvel 18276.t1-p1
Pmarinus Vti1 EER05640
Pmarinus Vti1 EER11999
Pmarinus Vti1 EER06521
Vbrassicaformis Vti1 Vbra 5076.t1-p1
Etenella Vti1 ETH 00019915-t26 1-p1
Hhammondi Vti1 HHA 278160-t26 1-p1
Tgondii-ME49 Vti1 TGME49 278160-t26 1-p1
Ncaninum Vti1 NCLIV 067830-t26 1-p1
Sneurona Vti1 SN3 00300230-mRNA-1-p1
Smicroadriaticum Vti1 Smic28209
Sminutum Vti1 symbB.v1.2.013418.t1scaffold940.1size1501876
Bmicroti Vti1 BmR1 04g07365-t32 1-p1
Bbigemina Vti1 BBBOND 0403350-t26 1-p1
Bbovis Vti1 BBOV IV000640-t26 1-p1
Cfelis Vti1 CF003911.mRNA-p1
Tannulata Vti1 TA19995-t26 1-p1
Tparva Vti1 TP01 0085-t26 1-p1
Torientalis Vti1 TOT 010000079-t26 1-p1
Bmicroti Vti1 BmR1 04g09840-t32 1-p1
Pberghei Vti1 PBANKA 1450600.1-p1
Pyoelii Vti1 PYYM 1454600.1-p1
Pchabaudi Vti1 PCHAS 1452900.1-p1
Povale Vti1 PocGH01 14061600.1-p1
Pknowlesi Vti1 PKNH 1455700.1-p1
Pvivax Vti1 PVX 100750.1-p1
Preichenowi Vti1 PRCDC 1235200.1-p1
Pgallinaceum Vti1 PGAL8A 00457700.1-p1
Prelictum Vti1 PRELSG 1452100.1-p1
Chominis Vti1 ChTU502y2012 384g0275-t32 1-p1
Cparvum Vti1 cgd3 180-t26 1-p1
Cmuris Vti1 CMU 038600-t26 1-p1
Imultifiliis Vti1 IMG5 032270
Tthermophila Vti1 TTHERM 00497550
Tthermophila Vti1 TTHERM 00853080
Ptetraurelia Vti1 GSPATP00021039001
Ptetraurelia Vti1 GSPATP00022941001
Otrifallax Vti1 Contig8672.0.g66Vesicle
Otrifallax Vti1 Contig20557.0.g57Vesicle
Ptetraurelia Vti1 GSPATP00006546001
Ptetraurelia Vti1 GSPATP00002940001
Ptetraurelia Vti1 GSPATP00004650001
Tthermophila Vti1 TTHERM 00434120
Ptetraurelia Vti1 GSPATP00026440001
Tthermophila Vti1 TTHERM 00094080
Otrifallax Vti1 Contig12823.0.g19Vesicle
Otrifallax Vti1 Contig22204.0.g76Uncharacterized
Otrifallax Vti1 Contig3273.0.g51LMBR1-like
Ptetraurelia Vti1 GSPATP00022472001
Bnatans Vti1 jgiBigna1139534aug1.51 g14242
Chominis Vti1 ChTU502y2012 408g0015-t32 1-p1
Cparvum Vti1 cgd4 3890-t26 1-p1
Cmuris Vti1 CMU 012970-t26 1-p1
Bbigemina NPSN11 BBBOND 0205790-t26 1-p1
Bbovis NPSN11 BBOV I001720-t26 1-p1
Cfelis NPSN11 CF000587.mRNA-p1
Tannulata NPSN11 TA20420-t26 1-p1
Torientalis NPSN11 TOT 010000483-t26 1-p1
Tparva NPSN11 TP01 0526-t26 1-p1
Bmicroti NPSN11 BMR1 03g04445-t32 1-p1
Pberghei NPSN11 PBANKA 1312300.1-p1
Pyoelii NPSN11 PYYM 1313100.1-p1
Pchabaudi NPSN11 PCHAS 1315600.1-p1
Pfalciparum NPSN11 PF3D7 1448600.1-p1
Preichenowi NPSN11 PRCDC 1447900.1-p1
Pknowlesi NPSN11 PKNH 1233500.1-p1
Pvivax NPSN11 PVX 118060.1-p1
Povale NPSN11 PocGH01 12066300.1-p1
Pgallinaceum NPSN11 PGAL8A 00207800.1-p1
Prelictum NPSN11 PRELSG 1232400.1-p1
Ccayetanensis NPSN11 cyc 05771-t31 1-p1
Etenella NPSN11 ETH 00001710-t26 1-p1
Hhammondi NPSN11 HHA 306640-t26 1-p1
Tgondii-ME49 NPSN11 TGME49 306640-t26 1-p1
Ncaninum NPSN11 NCLIV 044805-t26 1-p1
Sneurona NPSN11 SN3 00700835-mRNA-1-p1
Chominis NPSN11 ChTU502y2012 369g0040-t32 1-p1
Cparvum NPSN11 cgd3 4300-t26 1-p1
Cmuris NPSN11 CMU 030680-t26 1-p1
Gniphandrodes NPSN11 GNI 005310-t26 1-p1
Cvelia NPSN11 Cvel 26265.t1-p1
Smicroadriaticum NPSN11 Smic6464
Sminutum NPSN11 symbB.v1.2.003129.t1scaffold128.1size3124243
Vbrassicaformis NPSN11 Vbra 1346.t1-p1
Otrifallax NPSN11 Contig21502.0.g64SNARE
Otrifallax NPSN11 Contig5242.0.g22Protein
Ptetraurelia NPSN11 GSPATP00012472001
Tthermophila NPSN11 TTHERM 00300440
Tthermophila NPSN11 TTHERM 00405560
Athaliana NPSN11 AT1G48240.1
Athaliana NPSN11 AT3G17440.1
Athaliana NPSN11 AT2G35190.1
Creinhardtii NPSN11 Cre07.g342050.t1.2
Gtheta NPSN11 jgiGuith1165311estExt fgenesh2 pg.C 880058
Gtheta NPSN11 jgiGuith1138563fgenesh2 pg.32 123
Psojae NPSN11 jgiPhyso3249448Physo2.s 446245
Pultimum NPSN11 PYU1 T011776
Ptricornutum NPSN11 jgiPhatr254855estExt Phatr1 ua kg.C chr 160022
Tpseudonana NPSN11 jgiThaps34535fgenesh1 pg.C chr 4000485
Esiliculosus NPSN11 Ec-20 002300
Aanophagefferens NPSN11 jgiAuran172457
Aanophagefferens NPSN11 jgiAuran172562

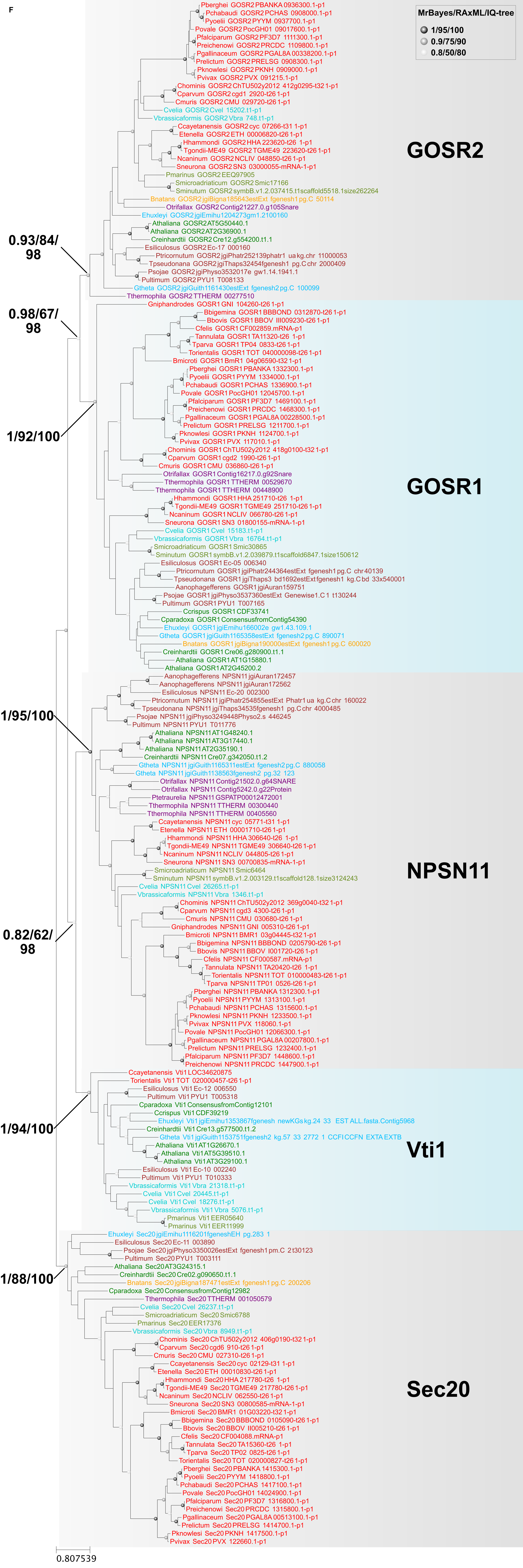
F
MrBayes/RAxML/IQ-tree
1/95/100
0.9/75/90
0.8/50/80
Pberghei GOSR2 PBANKA 0936300.1-p1
Pchabaudi GOSR2 PCHAS 0908000.1-p1
Pyoelii GOSR2 PYYM 0937700.1-p1
Povale GOSR2 PocGH01 09017600.1-p1
Pfalciparum GOSR2 PF3D7 1111300.1-p1
Preichenowi GOSR2 PRCDC 1109800.1-p1
Pgallinaceum GOSR2 PGAL8A 00338200.1-p1
Prelictum GOSR2 PRELSG 0908300.1-p1
Pknowlesi GOSR2 PKNH 0909000.1-p1
Pvivax GOSR2 PVX 091215.1-p1
Chominis GOSR2 ChTU502y2012 412g0295-t32 1-p1
Cparvum GOSR2 cgd1 2920-t26 1-p1
Cmuris GOSR2 CMU 029720-t26 1-p1
Cvelia GOSR2 Cvel 15202.t1-p1
Vbrassicaformis GOSR2 Vbra 748.t1-p1
Ccayetanensis GOSR2 cyc 07266-t31 1-p1
Etenella GOSR2 ETH 00006820-t26 1-p1
Hhammondi GOSR2 HHA 223620-t26 1-p1
Tgondii-ME49 GOSR2 TGME49 223620-t26 1-p1
Ncaninum GOSR2 NCLIV 048850-t26 1-p1
Sneurona GOSR2 SN3 03000055-mRNA-1-p1
Pmarinus GOSR2 EEQ97905
Smicroadriaticum GOSR2 Smic17166
Sminutum GOSR2 symbB.v1.2.037415.t1scaffold5518.1size262264
Bnatans GOSR2 jgiBigna185643estExt fgenesh1 pg.C 50114
Otrifallax GOSR2 Contig21227.0.g105Snare
Ehuxleyi GOSR2 jgiEmihu1204273gm1.2100160
Athaliana GOSR2 AT5G50440.1
Athaliana GOSR2 AT2G36900.1
Creinhardtii GOSR2 Cre12.g554200.t1.1
Esiliculosus GOSR2 Ec-17 000160
Ptricornutum GOSR2 jgiPhatr252139phatr1 ua kg.chr 11000053
Tpseudonana GOSR2 jgiThaps32454fgenesh1 pg.C chr 2000409
Psojae GOSR2 jgiPhyso3532017e gw1.14.1941.1
Pultimum GOSR2 PYU1 T008133
Gtheta GOSR2 jgiGuith1161430estExt fgenesh2 pg.C 100099
Tthermophila GOSR2 TTHERM 00277510
GOSR2
0.93/84/
98
0.98/67/
98
Gniphandrodes GOSR1 GNI 104260-t26 1-p1
Bbigemina GOSR1 BBBOND 0312870-t26 1-p1
Bbovis GOSR1 BBOV III009230-t26 1-p1
Cfelis GOSR1 CF002859.mRNA-p1
Tannulata GOSR1 TA11320-t26 1-p1
Tparva GOSR1 TP04 0833-t26 1-p1
Torientalis GOSR1 TOT 040000098-t26 1-p1
Bmicroti GOSR1 BmR1 04g06590-t32 1-p1
Pberghei GOSR1 PBANKA 1332300.1-p1
Pyoelii GOSR1 PYYM 1334000.1-p1
Pchabaudi GOSR1 PCHAS 1336900.1-p1
Povale GOSR1 PocGH01 12045700.1-p1
Pfalciparum GOSR1 PF3D7 1469100.1-p1
Preichenowi GOSR1 PRCDC 1468300.1-p1
Pgallinaceum GOSR1 PGAL8A 00228500.1-p1
Prelictum GOSR1 PRELSG 1211700.1-p1
Pknowlesi GOSR1 PKNH 1124700.1-p1
Pvivax GOSR1 PVX 117010.1-p1
Chominis GOSR1 ChTU502y2012 418g0100-t32 1-p1
Cparvum GOSR1 cgd2 1990-t26 1-p1
Cmuris GOSR1 CMU 036860-t26 1-p1
Otrifallax GOSR1 Contig16217.0.g92Snare
Tthermophila GOSR1 TTHERM 00529670
Tthermophila GOSR1 TTHERM 00448900
Hhammondi GOSR1 HHA 251710-t26 1-p1
Tgondii-ME49 GOSR1 TGME49 251710-t26 1-p1
Ncaninum GOSR1 NCLIV 066780-t26 1-p1
Sneurona GOSR1 SN3 01800155-mRNA-1-p1
Cvelia GOSR1 Cvel 15183.t1-p1
Vbrassicaformis GOSR1 Vbra 16764.t1-p1
Smicroadriaticum GOSR1 Smic30865
Sminutum GOSR1 symbB.v1.2.039879.t1scaffold6847.1size150612
Esiliculosus GOSR1 Ec-05 006340
Ptricornutum GOSR1 jgiPhatr244364estExt fgenesh1 pg.C chr 40139
Tpseudonana GOSR1 jgiThaps3 bd1692estExt fgenesh1 kg.C bd 33x540001
Aanophagefferens GOSR1 jgiAuran159751
Psojae GOSR1 jgiPhyso3537360estExt Genewise1.C 1 t130244
Pultimum GOSR1 PYU1 T007165
Ccrispus GOSR1 CDF33741
Cparadoxa GOSR1 ConsensusfromContig54390
Ehuxleyi GOSR1 jgiEmihu166002e gw1.43.109.1
Gtheta GOSR1 jgiGuith1165358estExt fgenesh2 pg.C 890071
Bnatans GOSR1 jgiBigna190000estExt fgenesh1 pg.C 600020
Creinhardtii GOSR1 Cre06.g280900.t1.1
Athaliana GOSR1 AT1G15880.1
Athaliana GOSR1 AT2G45200.2
GOSR1
1/92/100
Aanophagefferens NPSN11 jgiAuran172457
Aanophagefferens NPSN11 jgiAuran172562
Esiliculosus NPSN11 Ec-20 002300
Ptricornutum NPSN11 jgiPhatr254855estExt Phatr1 ua kg.C chr 160022
Tpseudonana NPSN11 jgiThaps34535fgenesh1 pg.C chr 4000485
Psojae NPSN11 jgiPhyso3249448Physo2.s 446245
Pultimum NPSN11 PYU1 T011776
Athaliana NPSN11 AT1G48240.1
Athaliana NPSN11 AT3G17440.1
Athaliana NPSN11 AT2G35190.1
Creinhardtii NPSN11 Cre07.g342050.t1.2
Gtheta NPSN11 jgiGuith165311estExt fgenesh2 pg.C 880058
Gtheta NPSN11 jgiGuith138563fgenesh2 pg.32 123
Otrifallax NPSN11 Contig21502.0.g64SNARE
Otrifallax NPSN11 Contig5242.0.g22Protein
Ptetraurelia NPSN11 GSPATP00012472001
Tthermophila NPSN11 TTHERM 00300440
Tthermophila NPSN11 TTHERM 00405560
Ccayetanensis NPSN11 cyc 05771-t31 1-p1
Etenella NPSN11 ETH 00001710-t26 1-p1
Hhammondi NPSN11 HHA 306640-t26 1-p1
Tgondii-ME49 NPSN11 TGME49 306640-t26 1-p1
Ncaninum NPSN11 NCLIV 044805-t26 1-p1
Sneurona NPSN11 SN3 00700835-mRNA-1-p1
Smicroadriaticum NPSN11 Smic6464
Sminutum NPSN11 symbB.v1.2.003129.t1scaffold128.1size3124243
Cvelia NPSN11 Cvel 26265.t1-p1
Vbrassicaformis NPSN11 Vbra 1346.t1-p1
Chominis NPSN11 ChTU502y2012 369g0040-t32 1-p1
Cparvum NPSN11 cgd3 4300-t26 1-p1
Cmuris NPSN11 CMU 030680-t26 1-p1
Gniphandrodes NPSN11 GNI 005310-t26 1-p1
Bmicroti NPSN11 BMR1 03g04445-t32 1-p1
Bbigemina NPSN11 BBBOND 0205790-t26 1-p1
Bbovis NPSN11 BBOV I001720-t26 1-p1
Cfelis NPSN11 CF000587.mRNA-p1
Tannulata NPSN11 TA20420-t26 1-p1
Torientalis NPSN11 TOT 010000483-t26 1-p1
Tparva NPSN11 TP01 0526-t26 1-p1
Pberghei NPSN11 PBANKA 1312300.1-p1
Pyoelii NPSN11 PYYM 1313100.1-p1
Pchabaudi NPSN11 PCHAS 1315600.1-p1
Pknowlesi NPSN11 PKNH 1233500.1-p1
Pvivax NPSN11 PVX 118060.1-p1
Povale NPSN11 PocGH01 12066300.1-p1
Pgallinaceum NPSN11 PGAL8A 00207800.1-p1
Prelictum NPSN11 PRELSG 1232400.1-p1
Pfalciparum NPSN11 PF3D7 1448600.1-p1
Preichenowi NPSN11 PRCDC 1447900.1-p1
NPSN11
1/95/100
0.82/62/
98
Ccayetanensis Vti1 LOC34620875
Torientalis Vti1 TOT 020000457-t26 1-p1
Esiliculosus Vti1 Ec-12 006550
Pultimum Vti1 PYU1 T005318
Cparadoxa Vti1 ConsensusfromContig12101
Ccrispus Vti1 CDF39219
Ehuxleyi Vti1 jgiEmihu1353867fgenesh newKGs kg.24 33 EST ALL.fasta.Contig5968
Creinhardtii Vti1 Cre13.g577500.t1.2
Gtheta Vti1 jgiGuith1153751fgenesh2 kg.57 33 2772 1 CCFICCFN EXTA EXTB
Athaliana Vti1 AT1G26670.1
Athaliana Vti1 AT5G39510.1
Athaliana Vti1 AT3G29100.1
Esiliculosus Vti1 Ec-10 002240
Pultimum Vti1 PYU1 T010333
Vbrassicaformis Vti1 Vbra 21318.t1-p1
Cvelia Vti1 Cvel 20445.t1-p1
Cvelia Vti1 Cvel 18276.t1-p1
Vbrassicaformis Vti1 Vbra 5076.t1-p1
Pmarinus Vti1 EER05640
Pmarinus Vti1 EER11999
Vti1
1/94/100
Ehuxleyi Sec20 jgiEmihu1116201fgeneshEH pg.283 1
Esiliculosus Sec20 Ec-11 003890
Psojae Sec20 jgiPhyso3350026estExt fgenesh1 pm.C 2t30123
Pultimum Sec20 PYU1 T003111
Athaliana Sec20 AT3G24315.1
Creinhardtii Sec20 Cre02.g090650.t1.1
Bnatans Sec20 jgiBigna187471estExt fgenesh1 pg.C 200206
Cparadoxa Sec20 ConsensusfromContig12982
Tthermophila Sec20 TTHERM 001050579
Cvelia Sec20 Cvel 26237.t1-p1
Smicroadriaticum Sec20 Smic6788
Pmarinus Sec20 EER17376
Vbrassicaformis Sec20 Vbra 8949.t1-p1
Chominis Sec20 ChTU502y2012 406g0190-t32 1-p1
Cparvum Sec20 cgd6 910-t26 1-p1
Cmuris Sec20 CMU 027310-t26 1-p1
Ccayetanensis Sec20 cyc 02129-t31 1-p1
Etenella Sec20 ETH 00010830-t26 1-p1
Hhammondi Sec20 HHA 217780-t26 1-p1
Tgondii-ME49 Sec20 TGME49 217780-t26 1-p1
Ncaninum Sec20 NCLIV 062550-t26 1-p1
Sneurona Sec20 SN3 00800585-mRNA-1-p1
Bmicroti Sec20 BMR1 01G03220-t32 1-p1
Bbigemina Sec20 BBBOND 0105090-t26 1-p1
Bbovis Sec20 BBOV II005210-t26 1-p1
Cfelis Sec20 CF004088.mRNA-p1
Tannulata Sec20 TA15360-t26 1-p1
Tparva Sec20 TP02 0825-t26 1-p1
Torientalis Sec20 TOT 020000827-t26 1-p1
Pberghei Sec20 PBANKA 1415300.1-p1
Pyoelii Sec20 PYYM 1418800.1-p1
Pchabaudi Sec20 PCHAS 1417100.1-p1
Povale Sec20 PocGH01 14024900.1-p1
Pfalciparum Sec20 PF3D7 1316800.1-p1
Preichenowi Sec20 PRCDC 1315800.1-p1
Pgallinaceum Sec20 PGAL8A 00513100.1-p1
Prelictum Sec20 PRELSG 1414700.1-p1
Pknowlesi Sec20 PKNH 1417500.1-p1
Pvivax Sec20 PVX 122660.1-p1
Sec20
1/88/100
0.807539

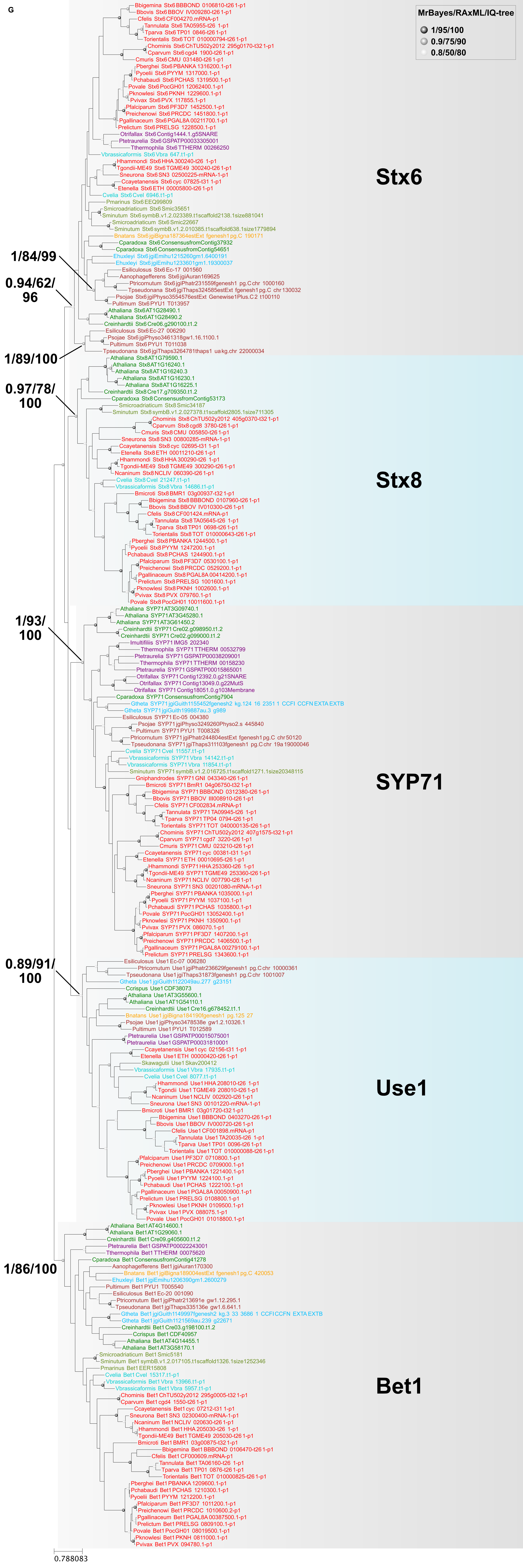
G
MrBayes/RAxML/IQ-tree
1/95/100
0.9/75/90
0.8/50/80
Stx6
Stx8
SYP71
Use1
Bet1
1/84/99
0.94/62/96
1/89/100
0.97/78/100
1/93/100
0.89/91/100
1/86/100
0.788083

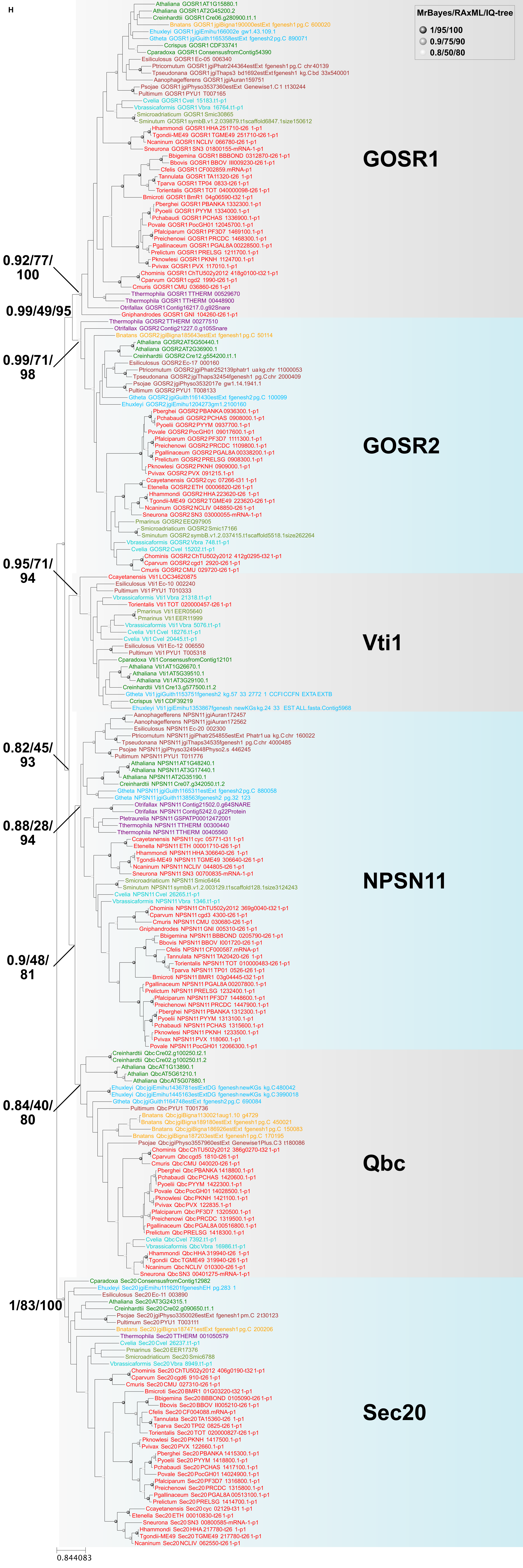
H
MrBayes/RAxML/IQ-tree
1/95/100
0.9/75/90
0.8/50/80
GOSR1
GOSR2
Vti1
NPSN11
Qbc
Sec20
0.92/77/100
0.99/49/95
0.99/71/98
0.95/71/94
0.82/45/93
0.88/28/94
0.9/48/81
0.84/40/80
1/83/100
0.844083

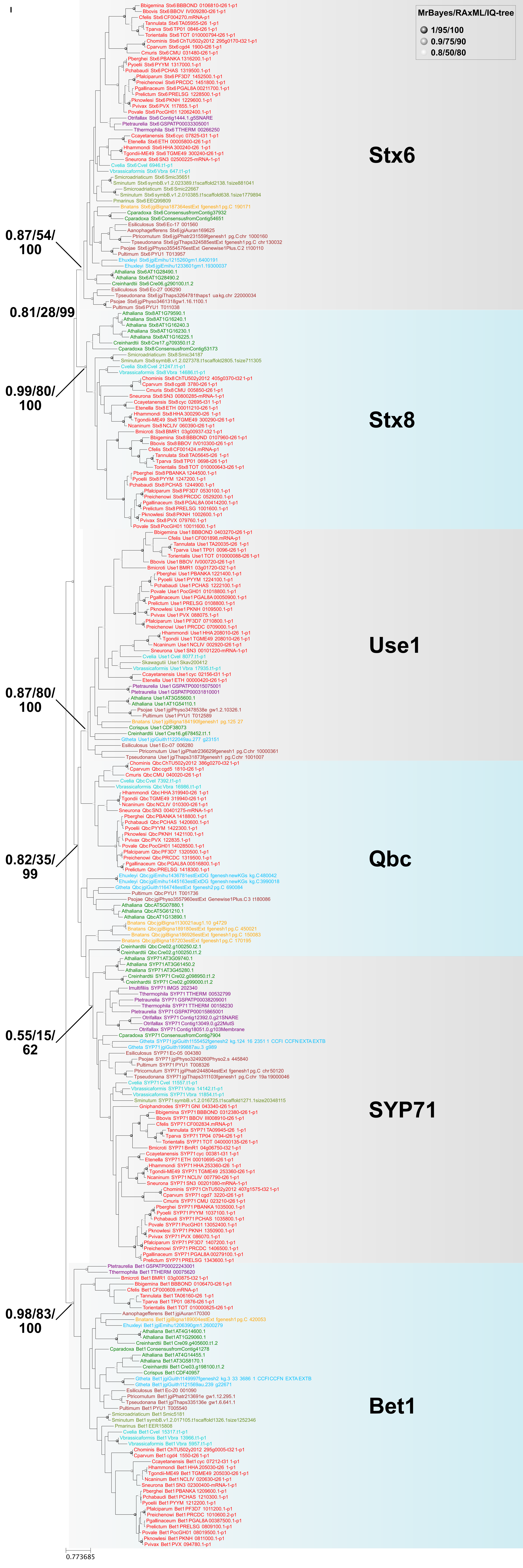

I
MrBayes/RAxML/IQ-tree
1/95/100
0.9/75/90
0.8/50/80
Stx6
0.87/54/
100
0.81/28/99
0.99/80/
100
Stx8
Use1
0.87/80/
100
Qbc
0.82/35/
99
0.55/15/
62
SYP71
0.98/83/
100
Bet1
0.773685

J

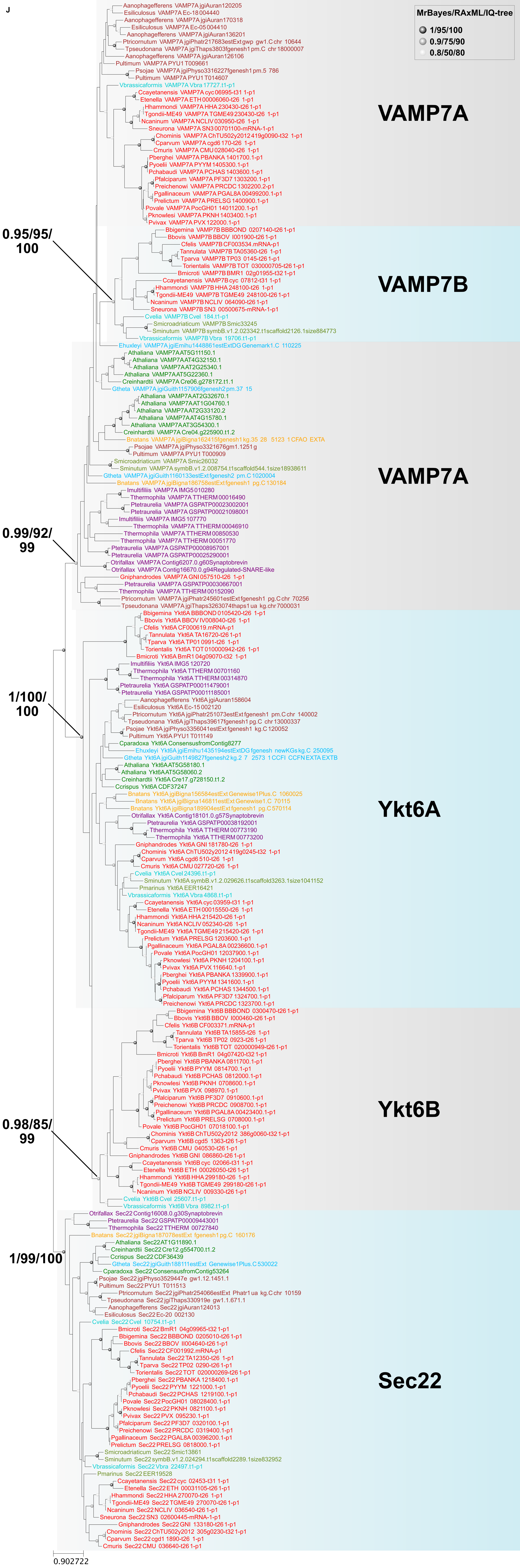
MrBayes/RAxML/IQ-tree
1/95/100
0.9/75/90
0.8/50/80
VAMP7A
VAMP7B
VAMP7A
Ykt6A
Ykt6B
Sec22
0.95/95/
100
0.99/92/
99
1/100/
100
0.98/85/
99
1/99/100
0.902722

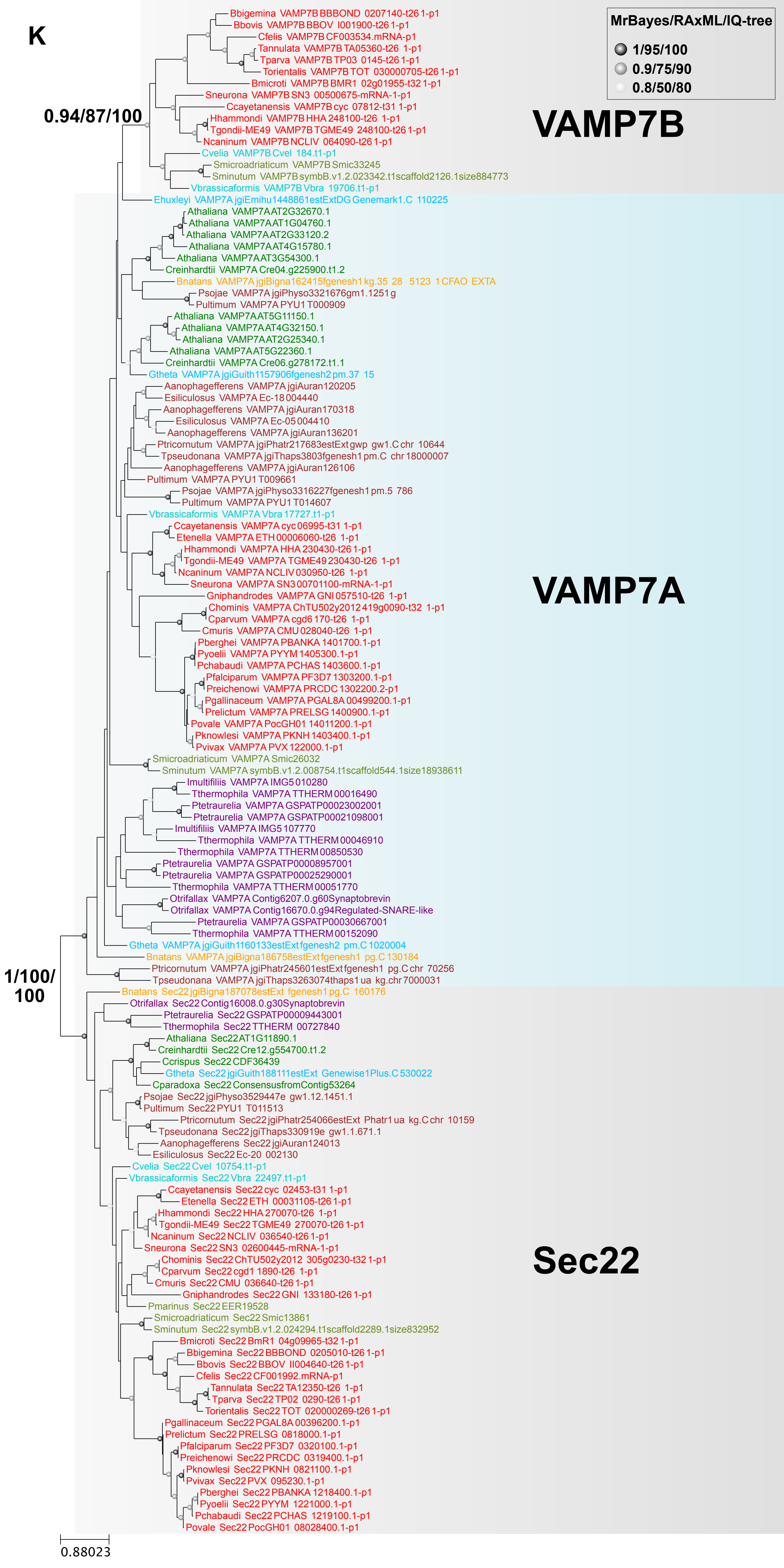

K
MrBayes/RAxML/IQ-tree
1/95/100
0.9/75/90
0.8/50/80
0.94/87/100
1/100/
100
VAMP7B
VAMP7A
Sec22
Bbigemina VAMP7B BBBOND 0207140-t26 1-p1
Bbovis VAMP7B BBOV I001900-t26 1-p1
Cfelis VAMP7B CF003534.mRNA-p1
Tannulata VAMP7B TA05360-t26 1-p1
Tparva VAMP7B TP03 0145-t26 1-p1
Torientalis VAMP7B TOT 030000705-t26 1-p1
Bmicroti VAMP7B BMR1 02g01955-t32 1-p1
Sneurona VAMP7B SN3 00500675-mRNA-1-p1
Ccayetanensis VAMP7B cyc 07812-t31 1-p1
Hhammondi VAMP7B HHA 248100-t26 1-p1
Tgondii-ME49 VAMP7B TGME49 248100-t26 1-p1
Ncaninum VAMP7B NCLIV 064090-t26 1-p1
Cvelia VAMP7B Cvel 184.t1-p1
Smicroadriaticum VAMP7B Smic33245
Sminutum VAMP7B symbB.v1.2.023342.t1scaffold2126.1size884773
Vbrassicaformis VAMP7B Vbra 19706.t1-p1
Ehuxleyi VAMP7A jgiEmihu1448861estExtDG Genemark1.C 110225
Athaliana VAMP7A AT2G32670.1
Athaliana VAMP7A AT1G04760.1
Athaliana VAMP7A AT2G33120.2
Athaliana VAMP7A AT4G15780.1
Athaliana VAMP7A AT3G54300.1
Creinhardtii VAMP7A Cre04.g225900.t1.2
Bnatans VAMP7A jgiBigna162415fgenesh1 kg.35 28 5123 1CFAO EXTA
Psojae VAMP7A jgiPhyso3321676gm1.1251 g
Pultimum VAMP7A PYU1 T000909
Athaliana VAMP7A AT5G11150.1
Athaliana VAMP7A AT4G32150.1
Athaliana VAMP7A AT2G25340.1
Athaliana VAMP7A AT5G22360.1
Creinhardtii VAMP7A Cre06.g278172.t1.1
Gtheta VAMP7A jgiGuith1157906fgenesh2 pm.37 15
Aanophagefferens VAMP7A jgiAuran120205
Esiliculosus VAMP7A Ec-18 004440
Aanophagefferens VAMP7A jgiAuran170318
Esiliculosus VAMP7A Ec-05 004410
Aanophagefferens VAMP7A jgiAuran136201
Ptricornutum VAMP7A jgiPhatr217683estExt gwp gw1.C chr 10644
Tpseudonana VAMP7A jgiThaps3803fgenesh1 pm.C chr 18000007
Aanophagefferens VAMP7A jgiAuran126106
Pultimum VAMP7A PYU1 T009661
Psojae VAMP7A jgiPhyso3316227fgenesh1 pm.5 786
Pultimum VAMP7A PYU1 T014607
Vbrassicaformis VAMP7A Vbra 17727.t1-p1
Ccayetanensis VAMP7A cyc 06995-t31 1-p1
Etenella VAMP7A ETH 00006060-t26 1-p1
Hhammondi VAMP7A HHA 230430-t26 1-p1
Tgondii-ME49 VAMP7A TGME49 230430-t26 1-p1
Ncaninum VAMP7A NCLIV 030950-t26 1-p1
Sneurona VAMP7A SN3 00701100-mRNA-1-p1
Gniphandrodes VAMP7A GNI 057510-t26 1-p1
Chominis VAMP7A ChTU502y2012 419g0090-t32 1-p1
Cparvum VAMP7A cgd6 170-t26 1-p1
Cmuris VAMP7A CMU 028040-t26 1-p1
Pberghei VAMP7A PBANKA 1401700.1-p1
Pyoelii VAMP7A PYYM 1405300.1-p1
Pchabaudi VAMP7A PCHAS 1403600.1-p1
Pfalciparum VAMP7A PF3D7 1303200.1-p1
Preichenowi VAMP7A PRCDC 1302200.2-p1
Pgallinaceum VAMP7A PGAL8A 00499200.1-p1
Prelictum VAMP7A PRELSG 1400900.1-p1
Povale VAMP7A PocGH01 14011200.1-p1
Pknowlesi VAMP7A PKNH 1403400.1-p1
Pvivax VAMP7A PVX 122000.1-p1
Smicroadriaticum VAMP7A Smic26032
Sminutum VAMP7A symbB.v1.2.008754.t1scaffold544.1size18938611
Imultifiliis VAMP7A IMG5 010280
Tthermophila VAMP7A TTHERM 00016490
Ptetraurelia VAMP7A GSPATP00023002001
Ptetraurelia VAMP7A GSPATP00021098001
Imultifiliis VAMP7A IMG5 107770
Tthermophila VAMP7A TTHERM 00046910
Tthermophila VAMP7A TTHERM 00850530
Ptetraurelia VAMP7A GSPATP00008957001
Ptetraurelia VAMP7A GSPATP00025290001
Tthermophila VAMP7A TTHERM 00051770
Otrifallax VAMP7A Contig6207.0.g60Synaptobrevin
Otrifallax VAMP7A Contig16670.0.g94Regulated-SNARE-like
Ptetraurelia VAMP7A GSPATP00030667001
Tthermophila VAMP7A TTHERM 00152090
Gtheta VAMP7A jgiGuith1160133estExtfgenesh2 pm.C 1020004
Bnatans VAMP7A jgiBigna186758estExtfgenesh1 pg.C 130184
Ptricornutum VAMP7A jgiPhatr245601estExtfgenesh1 pg.C chr 70256
Tpseudonana VAMP7A jgiThaps3263074thaps1 ua kg.chr 7000031
Bnatans Sec22 jgiBigna187078estExt fgenesh1 pg.C 160176
Otrifallax Sec22 Contig16008.0.g30Synaptobrevin
Ptetraurelia Sec22 GSPATP00009443001
Tthermophila Sec22 TTHERM 00727840
Athaliana Sec22 AT1G11890.1
Creinhardtii Sec22 Cre12.g554700.t1.2
Ccrispus Sec22 CDF36439
Gtheta Sec22 jgiGuith188111estExt Genewise1Plus.C 530022
Cparadoxa Sec22 ConsensusfromContig53264
Psojae Sec22 jgiPhyso3529447e gw1.12.1451.1
Pultimum Sec22 PYU1 T011513
Ptricornutum Sec22 jgiPhatr254066estExt Phatr1 ua kg.C chr 10159
Tpseudonana Sec22 jgiThaps330919e gw1.1.671.1
Aanophagefferens Sec22 jgiAuran124013
Esiliculosus Sec22 Ec-20 002130
Cvelia Sec22 Cvel 10754.t1-p1
Vbrassicaformis Sec22 Vbra 22497.t1-p1
Ccayetanensis Sec22 cyc 02453-t31 1-p1
Etenella Sec22 ETH 00031105-t26 1-p1
Hhammondi Sec22 HHA 270070-t26 1-p1
Tgondii-ME49 Sec22 TGME49 270070-t26 1-p1
Ncaninum Sec22 NCLIV 036540-t26 1-p1
Sneurona Sec22 SN3 02600445-mRNA-1-p1
Chominis Sec22 ChTU502y2012 305g0230-t32 1-p1
Cparvum Sec22 cgd1 1890-t26 1-p1
Cmuris Sec22 CMU 036640-t26 1-p1
Gniphandrodes Sec22 GNI 133180-t26 1-p1
Pmarinus Sec22 EER19528
Smicroadriaticum Sec22 Smic13861
Sminutum Sec22 symbB.v1.2.024294.t1scaffold2289.1size832952
Bmicroti Sec22 BmR1 04g09965-t32 1-p1
Bbigemina Sec22 BBBOND 0205010-t26 1-p1
Bbovis Sec22 BBOV II004640-t26 1-p1
Cfelis Sec22 CF001992.mRNA-p1
Tannulata Sec22 TA12350-t26 1-p1
Tparva Sec22 TP02 0290-t26 1-p1
Torientalis Sec22 TOT 020000269-t26 1-p1
Pgallinaceum Sec22 PGAL8A 00396200.1-p1
Prelictum Sec22 PRELSG 0818000.1-p1
Pfalciparum Sec22 PF3D7 0320100.1-p1
Preichenowi Sec22 PRCDC 0319400.1-p1
Pknowlesi Sec22 PKNH 0821100.1-p1
Pvivax Sec22 PVX 095230.1-p1
Pberghei Sec22 PBANKA 1218400.1-p1
Pyoelii Sec22 PYYM 1221000.1-p1
Pchabaudi Sec22 PCHAS 1219100.1-p1
Povale Sec22 PocGH01 08028400.1-p1
0.88023

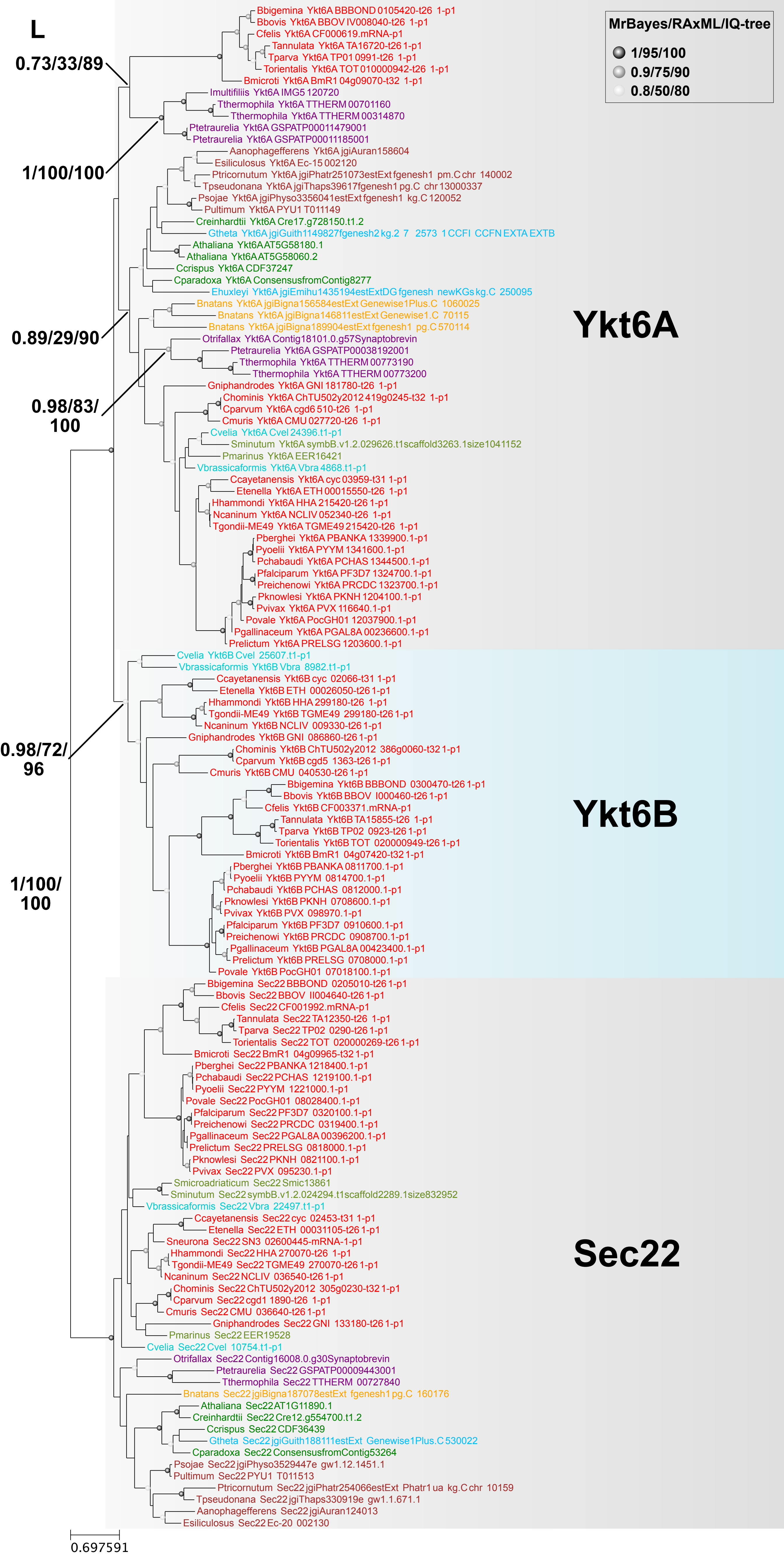
L
MrBayes/RAxML/IQ-tree
1/95/100
0.9/75/90
0.8/50/80
0.73/33/89
1/100/100
0.89/29/90
0.98/83/100
0.98/72/96
1/100/100
Ykt6A
Ykt6B
Sec22
Bbigemina Ykt6A BBBOND 0105420-t26 1-p1
Bbovis Ykt6A BBOV IV008040-t26 1-p1
Cfelis Ykt6A CF000619.mRNA-p1
Tannulata Ykt6A TA16720-t26 1-p1
Tparva Ykt6A TP01 0991-t26 1-p1
Torientalis Ykt6A TOT 010000942-t26 1-p1
Bmicroti Ykt6A BmR1 04g09070-t32 1-p1
Imultifiliis Ykt6A IMG5 120720
Tthermophila Ykt6A TTHERM 00701160
Tthermophila Ykt6A TTHERM 00314870
Ptetraurelia Ykt6A GSPATP00011479001
Ptetraurelia Ykt6A GSPATP00011185001
Aanophagefferens Ykt6A jgiAuran158604
Esiliculosus Ykt6A Ec-15 002120
Ptricornutum Ykt6A jgiPhatr251073estExt fgenesh1 pm.C chr 140002
Tpseudonana Ykt6A jgiThaps39617fgenesh1 pg.C chr 13000337
Psojae Ykt6A jgiPhyso3356041estExt fgenesh1 kg.C 120052
Pultimum Ykt6A PYU1 T011149
Creinhardtii Ykt6A Cre17.g728150.t1.2
Gtheta Ykt6A jgiGuith1149827fgenesh2 kg.2 7 2573 1 CCFI CCFN EXTA EXTB
Athaliana Ykt6A AT5G58180.1
Athaliana Ykt6A AT5G58060.2
Ccrispus Ykt6A CDF37247
Cparadoxa Ykt6A ConsensusfromContig8277
Ehuxleyi Ykt6A jgiEmihu1435194estExtDG fgenesh newKGs kg.C 250095
Bnatans Ykt6A jgiBigna156584estExt Genewise1Plus.C 1060025
Bnatans Ykt6A jgiBigna146811estExt Genewise1.C 70115
Bnatans Ykt6A jgiBigna189904estExt fgenesh1 pg.C 570114
Otrifallax Ykt6A Contig18101.0.g57Synaptobrevin
Ptetraurelia Ykt6A GSPATP00038192001
Tthermophila Ykt6A TTHERM 00773190
Tthermophila Ykt6A TTHERM 00773200
Gniphandrodes Ykt6A GNI 181780-t26 1-p1
Chominis Ykt6A ChTU502y2012 419g0245-t32 1-p1
Cparvum Ykt6A cgd6 510-t26 1-p1
Cmuris Ykt6A CMU 027720-t26 1-p1
Cvelia Ykt6A Cvel 24396.t1-p1
Sminutum Ykt6A symbB.v1.2.029626.t1scaffold3263.1size1041152
Pmarinus Ykt6A EER16421
Vbrassicaformis Ykt6A Vbra 4868.t1-p1
Ccayetanensis Ykt6A cyc 03959-t31 1-p1
Etenella Ykt6A ETH 00015550-t26 1-p1
Hhammondi Ykt6A HHA 215420-t26 1-p1
Ncaninum Ykt6A NCLIV 052340-t26 1-p1
Tgondii-ME49 Ykt6A TGME49 215420-t26 1-p1
Pberghei Ykt6A PBANKA 1339900.1-p1
Pyoelii Ykt6A PYYM 1341600.1-p1
Pchabaudi Ykt6A PCHAS 1344500.1-p1
Pfalciparum Ykt6A PF3D7 1324700.1-p1
Preichenowi Ykt6A PRCDC 1323700.1-p1
Pknowlesi Ykt6A PKNH 1204100.1-p1
Pvivax Ykt6A PVX 116640.1-p1
Povale Ykt6A PocGH01 12037900.1-p1
Pgallinaceum Ykt6A PGAL8A 00236600.1-p1
Prelictum Ykt6A PRELSG 1203600.1-p1
Cvelia Ykt6B Cvel 25607.t1-p1
Vbrassicaformis Ykt6B Vbra 8982.t1-p1
Ccayetanensis Ykt6B cyc 02066-t31 1-p1
Etenella Ykt6B ETH 00026050-t26 1-p1
Hhammondi Ykt6B HHA 299180-t26 1-p1
Tgondii-ME49 Ykt6B TGME49 299180-t26 1-p1
Ncaninum Ykt6B NCLIV 009330-t26 1-p1
Gniphandrodes Ykt6B GNI 086860-t26 1-p1
Chominis Ykt6B ChTU502y2012 386g0060-t32 1-p1
Cparvum Ykt6B cgd5 1363-t26 1-p1
Cmuris Ykt6B CMU 040530-t26 1-p1
Bbigemina Ykt6B BBBOND 0300470-t26 1-p1
Bbovis Ykt6B BBOV I000460-t26 1-p1
Cfelis Ykt6B CF003371.mRNA-p1
Tannulata Ykt6B TA15855-t26 1-p1
Tparva Ykt6B TP02 0923-t26 1-p1
Torientalis Ykt6B TOT 020000949-t26 1-p1
Bmicroti Ykt6B BmR1 04g07420-t32 1-p1
Pberghei Ykt6B PBANKA 0811700.1-p1
Pyoelii Ykt6B PYYM 0814700.1-p1
Pchabaudi Ykt6B PCHAS 0812000.1-p1
Pknowlesi Ykt6B PKNH 0708600.1-p1
Pvivax Ykt6B PVX 098970.1-p1
Pfalciparum Ykt6B PF3D7 0910600.1-p1
Preichenowi Ykt6B PRCDC 0908700.1-p1
Pgallinaceum Ykt6B PGAL8A 00423400.1-p1
Prelictum Ykt6B PRELSG 0708000.1-p1
Povale Ykt6B PocGH01 07018100.1-p1
Bbigemina Sec22 BBBOND 0205010-t26 1-p1
Bbovis Sec22 BBOV II004640-t26 1-p1
Cfelis Sec22 CF001992.mRNA-p1
Tannulata Sec22 TA12350-t26 1-p1
Tparva Sec22 TP02 0290-t26 1-p1
Torientalis Sec22 TOT 020000269-t26 1-p1
Bmicroti Sec22 BmR1 04g09965-t32 1-p1
Pberghei Sec22 PBANKA 1218400.1-p1
Pchabaudi Sec22 PCHAS 1219100.1-p1
Pyoelii Sec22 PYYM 1221000.1-p1
Povale Sec22 PocGH01 08028400.1-p1
Pfalciparum Sec22 PF3D7 0320100.1-p1
Preichenowi Sec22 PRCDC 0319400.1-p1
Pgallinaceum Sec22 PGAL8A 00396200.1-p1
Prelictum Sec22 PRELSG 0818000.1-p1
Pknowlesi Sec22 PKNH 0821100.1-p1
Pvivax Sec22 PVX 095230.1-p1
Smicroadriaticum Sec22 Smic13861
Sminutum Sec22 symbB.v1.2.024294.t1scaffold2289.1size832952
Vbrassicaformis Sec22 Vbra 22497.t1-p1
Ccayetanensis Sec22 cyc 02453-t31 1-p1
Etenella Sec22 ETH 00031105-t26 1-p1
Sneurona Sec22 SN3 02600445-mRNA-1-p1
Hhammondi Sec22 HHA 270070-t26 1-p1
Tgondii-ME49 Sec22 TGME49 270070-t26 1-p1
Ncaninum Sec22 NCLIV 036540-t26 1-p1
Chominis Sec22 ChTU502y2012 305g0230-t32 1-p1
Cparvum Sec22 cgd1 1890-t26 1-p1
Cmuris Sec22 CMU 036640-t26 1-p1
Gniphandrodes Sec22 GNI 133180-t26 1-p1
Pmarinus Sec22 EER19528
Cvelia Sec22 Cvel 10754.t1-p1
Otrifallax Sec22 Contig16008.0.g30Synaptobrevin
Ptetraurelia Sec22 GSPATP00009443001
Tthermophila Sec22 TTHERM 00727840
Bnatans Sec22 jgiBigna187078estExt fgenesh1 pg.C 160176
Athaliana Sec22 AT1G11890.1
Creinhardtii Sec22 Cre12.g554700.t1.2
Ccrispus Sec22 CDF36439
Gtheta Sec22 jgiGuith188111estExt Genewise1Plus.C 530022
Cparadoxa Sec22 ConsensusfromContig53264
Psojae Sec22 jgiPhyso3529447e gw1.12.1451.1
Pultimum Sec22 PYU1 T011513
Ptricornutum Sec22 jgiPhatr254066estExt Phatr1 ua kg.C chr 10159
Tpseudonana Sec22 jgiThaps330919e gw1.1.671.1
Aanophagefferens Sec22 jgiAuran124013
Esiliculosus Sec22 Ec-20 002130
0.697591

M

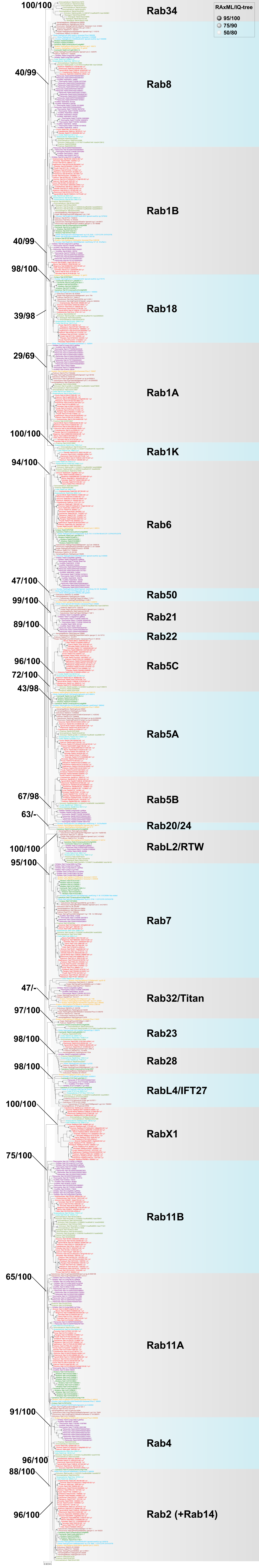
RAxML/IQ-tree
95/100
75/90
50/80
100/100
Rab34
40/99
Rab8
Rab1B
40/99
98/100
Rab18
39/98
29/69
Rab1A
100/100
Rab1K
94/100
Rab6
47/100
Rab50
99/100
Rab21
89/100
Rab22
96/100
Rab5C
72/100
43/98
Rab5A
67/98
Rab5B
63/-
Rab20/24
100/100
RabL2/RTW
95/100
Rab7
47/-
Rab32/Titan
97/100
Rab23
98/100
Rab28
98/100
RabL4/IFT27
100/100
RabX1
75/100
Rab11B
65/100
Rab11A
91/100
Rab4
96/100
88/100
96/100
Rab2 (+Rab14)

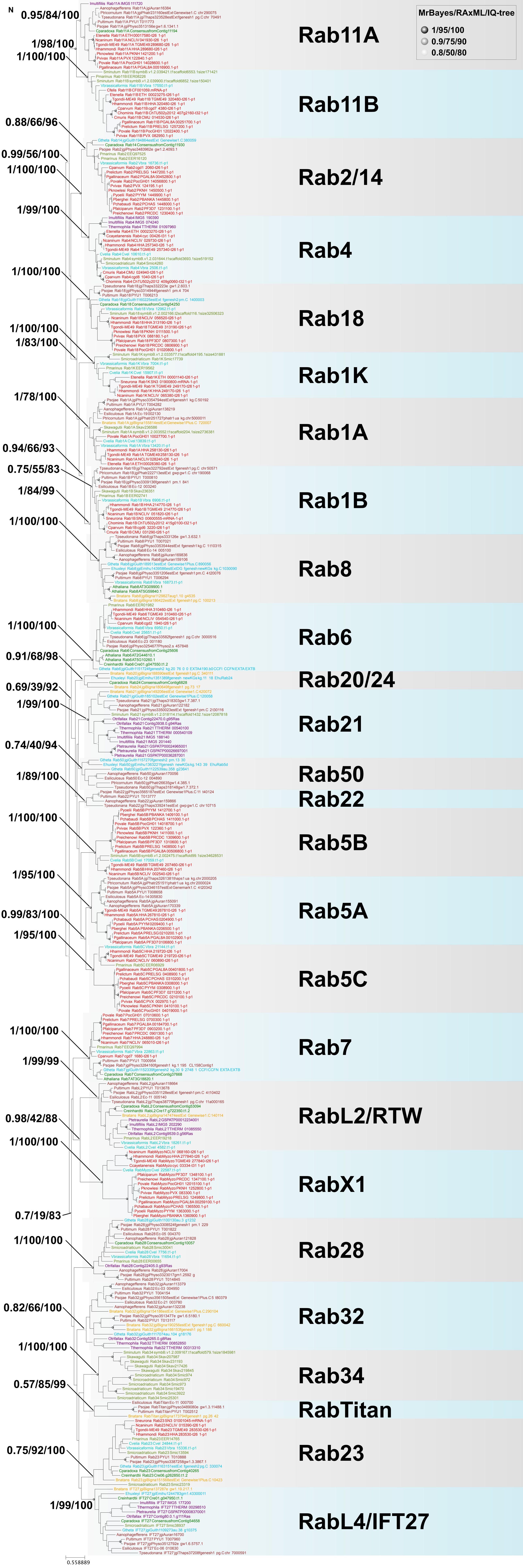
MrBayes/RAxML/IQ-tree
1/95/100
0.9/75/90
0.8/50/80
N
0.95/84/100
1/98/100
1/100/100
0.88/66/96
0.99/56/100
1/100/100
1/99/100
1/100/100
1/100/100
1/83/100
1/78/100
0.94/66/93
0.75/55/83
1/84/99
1/100/100
1/100/100
0.91/68/98
0.69/39/92
1/99/100
0.74/40/94
1/89/100
1/100/100
1/95/100
0.99/83/100
1/95/100
1/100/100
1/99/99
0.98/42/88
1/100/100
0.7/19/83
1/100/100
0.82/66/100
1/100/100
0.57/85/99
0.75/92/100
1/99/100
Rab11A
Rab11B
Rab2/14
Rab4
Rab18
Rab1K
Rab1A
Rab1B
Rab8
Rab6
Rab20/24
Rab21
Rab50
Rab22
Rab5B
Rab5A
Rab5C
Rab7
RabL2/RTW
RabX1
Rab28
Rab32
Rab34
RabTitan
Rab23
RabL4/IFT27
0.558889

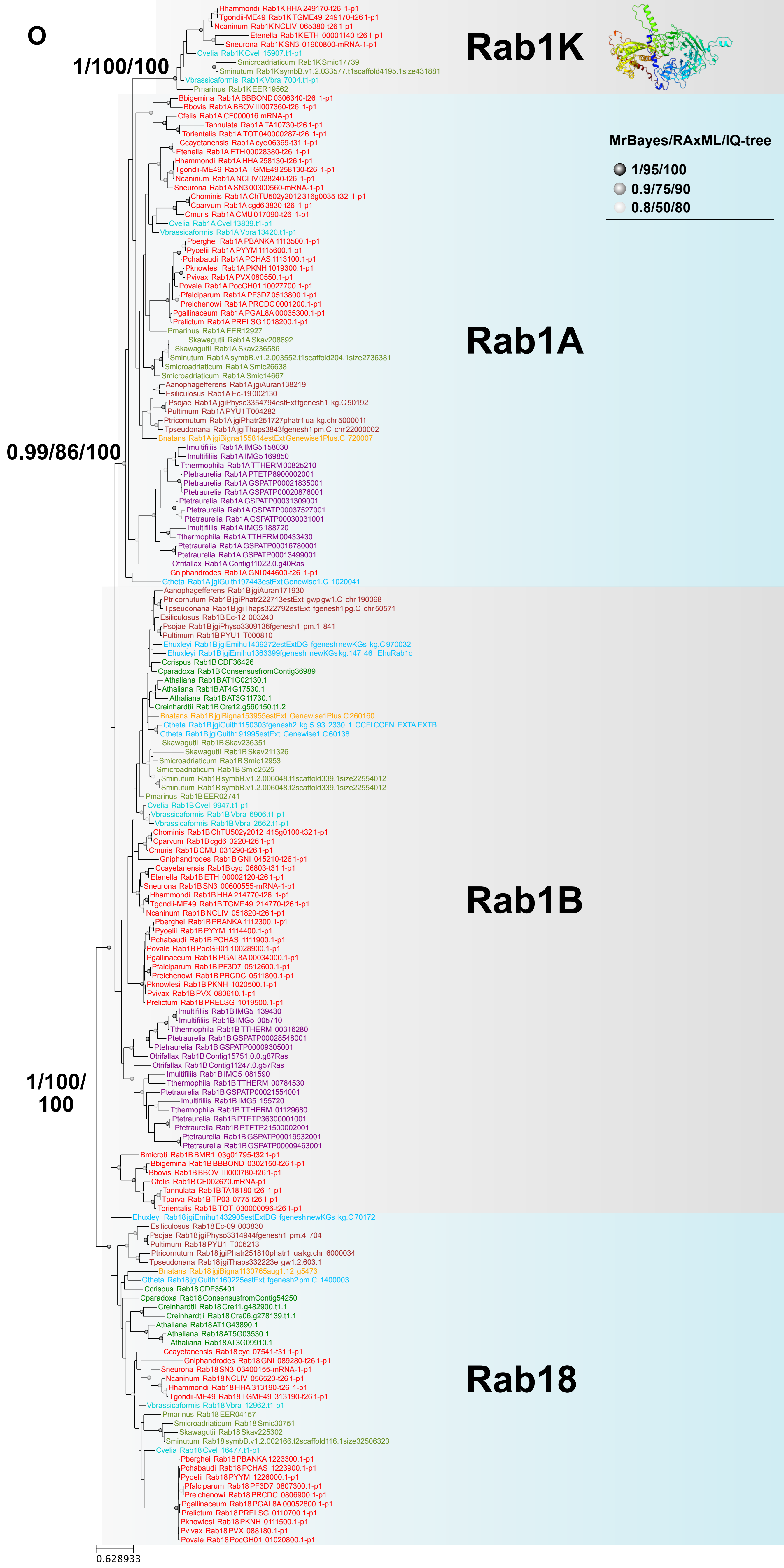
O
Rab1K
Rab1A
Rab1B
Rab18
MrBayes/RAxML/IQ-tree
1/95/100
0.9/75/90
0.8/50/80
1/100/100
0.99/86/100
1/100/
100
Hhammondi Rab1K HHA 249170-t26 1-p1
Tgondii-ME49 Rab1K TGME49 249170-t26 1-p1
Ncaninum Rab1K NCLIV 065380-t26 1-p1
Etenella Rab1K ETH 00001140-t26 1-p1
Sneurona Rab1K SN3 01900800-mRNA-1-p1
Cvelia Rab1K Cvel 15907.t1-p1
Smicroadriaticum Rab1K Smic17739
Sminutum Rab1K symbB.v1.2.033577.t1scaffold4195.1size431881
Vbrassicaformis Rab1K Vbra 7004.t1-p1
Pmarinus Rab1K EER19562
Bbigemina Rab1A BBBOND0306340-t26 1-p1
Bbovis Rab1A BBOV III007360-t26 1-p1
Cfelis Rab1A CF000016.mRNA-p1
Tannulata Rab1A TA10730-t26 1-p1
Torientalis Rab1A TOT 040000287-t26 1-p1
Ccayetanensis Rab1A cyc 06369-t31 1-p1
Etenella Rab1A ETH 00028380-t26 1-p1
Hhammondi Rab1A HHA 258130-t26 1-p1
Tgondii-ME49 Rab1A TGME49 258130-t26 1-p1
Ncaninum Rab1A NCLIV 028240-t26 1-p1
Sneurona Rab1A SN3 00300560-mRNA-1-p1
Chominis Rab1A ChTU502y2012 316g0035-t32 1-p1
Cparvum Rab1A cgd6 3830-t26 1-p1
Cmuris Rab1A CMU 017090-t26 1-p1
Cvelia Rab1A Cvel 13839.t1-p1
Vbrassicaformis Rab1A Vbra 13420.t1-p1
Pberghei Rab1A PBANKA 1113500.1-p1
Pyoelii Rab1A PYYM 1115600.1-p1
Pchabaudi Rab1A PCHAS 1113100.1-p1
Pknowlesi Rab1A PKNH 1019300.1-p1
Pvivax Rab1A PVX 080550.1-p1
Povale Rab1A PocGH01 10027700.1-p1
Pfalciparum Rab1A PF3D7 0513800.1-p1
Preichenowi Rab1A PRCDC 0001200.1-p1
Pgallinaceum Rab1A PGAL8A 00035300.1-p1
Prelictum Rab1A PRELSG 1018200.1-p1
Pmarinus Rab1A EER12927
Skawagutii Rab1A Skav208692
Skawagutii Rab1A Skav236586
Sminutum Rab1A symbB.v1.2.003552.t1scaffold204.1size2736381
Smicroadriaticum Rab1A Smic26638
Smicroadriaticum Rab1A Smic14667
Aanophagefferens Rab1A jgiAuran138219
Esiliculosus Rab1A Ec-19 002130
Psojae Rab1A jgiPhyso3354794estExt fgenesh1 kg.C 50192
Pultimum Rab1A PYU1 T004282
Ptricornutum Rab1A jgiPhatr251727phatr1 ua kg.chr 5000011
Tpseudonana Rab1A jgiThaps3843fgenesh1 pm.C chr 22000002
Bnatans Rab1A jgiBigna155814estExt Genewise1Plus.C 720007
Imultifiliis Rab1A IMG5 158030
Imultifiliis Rab1A IMG5 169850
Tthermophila Rab1A TTHERM 00825210
Ptetraurelia Rab1A PTETP8900002001
Ptetraurelia Rab1A GSPATP00021835001
Ptetraurelia Rab1A GSPATP00020876001
Ptetraurelia Rab1A GSPATP00031309001
Ptetraurelia Rab1A GSPATP00037527001
Ptetraurelia Rab1A GSPATP00030031001
Imultifiliis Rab1A IMG5 188720
Tthermophila Rab1A TTHERM 00433430
Ptetraurelia Rab1A GSPATP00016780001
Ptetraurelia Rab1A GSPATP00013499001
Otrifallax Rab1A Contig11022.0.g40Ras
Gniphandrodes Rab1A GNI 044600-t26 1-p1
Gtheta Rab1A jgiGuith197443estExt Genewise1.C 1020041
Aanophagefferens Rab1B jgiAuran171930
Ptricornutum Rab1B jgiPhatr222713estExt gwp gw1.C chr 190068
Tpseudonana Rab1B jgiThaps322792estExt fgenesh1 pg.C chr 50571
Esiliculosus Rab1B Ec-12 003240
Psojae Rab1B jgiPhyso3309136fgenesh1 pm.1 841
Pultimum Rab1B PYU1 T000810
Ehuxleyi Rab1B jgiEmihu1439272estExtDG fgenesh newKGs kg.C 970032
Ehuxleyi Rab1B jgiEmihu1363399fgenesh newKGs kg.147 46 EhuRab1c
Ccrispus Rab1B CDF36426
Cparadoxa Rab1B ConsensusfromContig36989
Athaliana Rab1B AT1G02130.1
Athaliana Rab1B AT4G17530.1
Athaliana Rab1B AT3G11730.1
Creinhardtii Rab1B Cre12.g560150.t1.2
Bnatans Rab1B jgiBigna153955estExt Genewise1Plus.C 260160
Gtheta Rab1B jgiGuith1150303fgenesh2 kg.5 93 2330 1 CCFI CCFN EXTA EXTB
Gtheta Rab1B jgiGuith191995estExt Genewise1.C 60138
Skawagutii Rab1B Skav236351
Skawagutii Rab1B Skav211326
Smicroadriaticum Rab1B Smic12953
Smicroadriaticum Rab1B Smic2525
Sminutum Rab1B symbB.v1.2.006048.t1scaffold339.1size22554012
Sminutum Rab1B symbB.v1.2.006048.t2scaffold339.1size22554012
Pmarinus Rab1B EER02741
Cvelia Rab1B Cvel 9947.t1-p1
Vbrassicaformis Rab1B Vbra 6906.t1-p1
Vbrassicaformis Rab1B Vbra 2662.t1-p1
Chominis Rab1B ChTU502y2012 415g0100-t32 1-p1
Cparvum Rab1B cgd6 3220-t26 1-p1
Cmuris Rab1B CMU 031290-t26 1-p1
Gniphandrodes Rab1B GNI 045210-t26 1-p1
Ccayetanensis Rab1B cyc 06803-t31 1-p1
Etenella Rab1B ETH 00002120-t26 1-p1
Sneurona Rab1B SN3 00600555-mRNA-1-p1
Hhammondi Rab1B HHA 214770-t26 1-p1
Tgondii-ME49 Rab1B TGME49 214770-t26 1-p1
Ncaninum Rab1B NCLIV 051820-t26 1-p1
Pberghei Rab1B PBANKA 1112300.1-p1
Pyoelii Rab1B PYYM 1114400.1-p1
Pchabaudi Rab1B PCHAS 1111900.1-p1
Povale Rab1B PocGH01 10028900.1-p1
Pgallinaceum Rab1B PGAL8A 00034000.1-p1
Pfalciparum Rab1B PF3D7 0512600.1-p1
Preichenowi Rab1B PRCDC 0511800.1-p1
Pknowlesi Rab1B PKNH 1020500.1-p1
Pvivax Rab1B PVX 080610.1-p1
Prelictum Rab1B PRELSG 1019500.1-p1
Imultifiliis Rab1B IMG5 139430
Imultifiliis Rab1B IMG5 005710
Tthermophila Rab1B TTHERM 00316280
Ptetraurelia Rab1B GSPATP00028548001
Ptetraurelia Rab1B GSPATP00009305001
Otrifallax Rab1B Contig15751.0.0.g87Ras
Otrifallax Rab1B Contig11247.0.g57Ras
Imultifiliis Rab1B IMG5 081590
Tthermophila Rab1B TTHERM 00784530
Ptetraurelia Rab1B GSPATP00021554001
Imultifiliis Rab1B IMG5 155720
Tthermophila Rab1B TTHERM 01129680
Ptetraurelia Rab1B PTETP36300001001
Ptetraurelia Rab1B PTETP21500002001
Ptetraurelia Rab1B GSPATP00019932001
Ptetraurelia Rab1B GSPATP00009463001
Bmicroti Rab1B BMR1 03g01795-t32 1-p1
Bbigemina Rab1B BBBOND 0302150-t26 1-p1
Bbovis Rab1B BBOV III000780-t26 1-p1
Cfelis Rab1B CF002670.mRNA-p1
Tannulata Rab1B TA18180-t26 1-p1
Tparva Rab1B TP03 0775-t26 1-p1
Torientalis Rab1B TOT 030000096-t26 1-p1
Ehuxleyi Rab18 jgiEmihu1432905estExtDG fgenesh newKGs kg.C 70172
Esiliculosus Rab18 Ec-09 003830
Psojae Rab18 jgiPhyso3314944fgenesh1 pm.4 704
Pultimum Rab18 PYU1 T006213
Ptricornutum Rab18 jgiPhatr251810phatr1 ua kg.chr 6000034
Tpseudonana Rab18 jgiThaps332223e gw1.2.603.1
Bnatans Rab18 jgiBigna1130765aug1.12 g5473
Gtheta Rab18 jgiGuith1160225estExt fgenesh2 pm.C 1400003
Ccrispus Rab18 CDF35401
Cparadoxa Rab18 ConsensusfromContig54250
Creinhardtii Rab18 Cre11.g482900.t1.1
Creinhardtii Rab18 Cre06.g278139.t1.1
Athaliana Rab18 AT1G43890.1
Athaliana Rab18 AT5G03530.1
Athaliana Rab18 AT3G09910.1
Ccayetanensis Rab18 cyc 07541-t31 1-p1
Gniphandrodes Rab18 GNI 089280-t26 1-p1
Sneurona Rab18 SN3 03400155-mRNA-1-p1
Ncaninum Rab18 NCLIV 056520-t26 1-p1
Hhammondi Rab18 HHA 313190-t26 1-p1
Tgondii-ME49 Rab18 TGME49 313190-t26 1-p1
Vbrassicaformis Rab18 Vbra 12962.t1-p1
Pmarinus Rab18 EER04157
Smicroadriaticum Rab18 Smic30751
Skawagutii Rab18 Skav225302
Sminutum Rab18 symbB.v1.2.002166.t2scaffold116.1size32506323
Cvelia Rab18 Cvel 16477.t1-p1
Pberghei Rab18 PBANKA 1223300.1-p1
Pchabaudi Rab18 PCHAS 1223900.1-p1
Pyoelii Rab18 PYYM 1226000.1-p1
Pfalciparum Rab18 PF3D7 0807300.1-p1
Preichenowi Rab18 PRCDC 0806900.1-p1
Pgallinaceum Rab18 PGAL8A 00052800.1-p1
Prelictum Rab18 PRELSG 0110700.1-p1
Pknowlesi Rab18 PKNH 0111500.1-p1
Pvivax Rab18 PVX 088180.1-p1
Povale Rab18 PocGH01 01020800.1-p1
0.628933

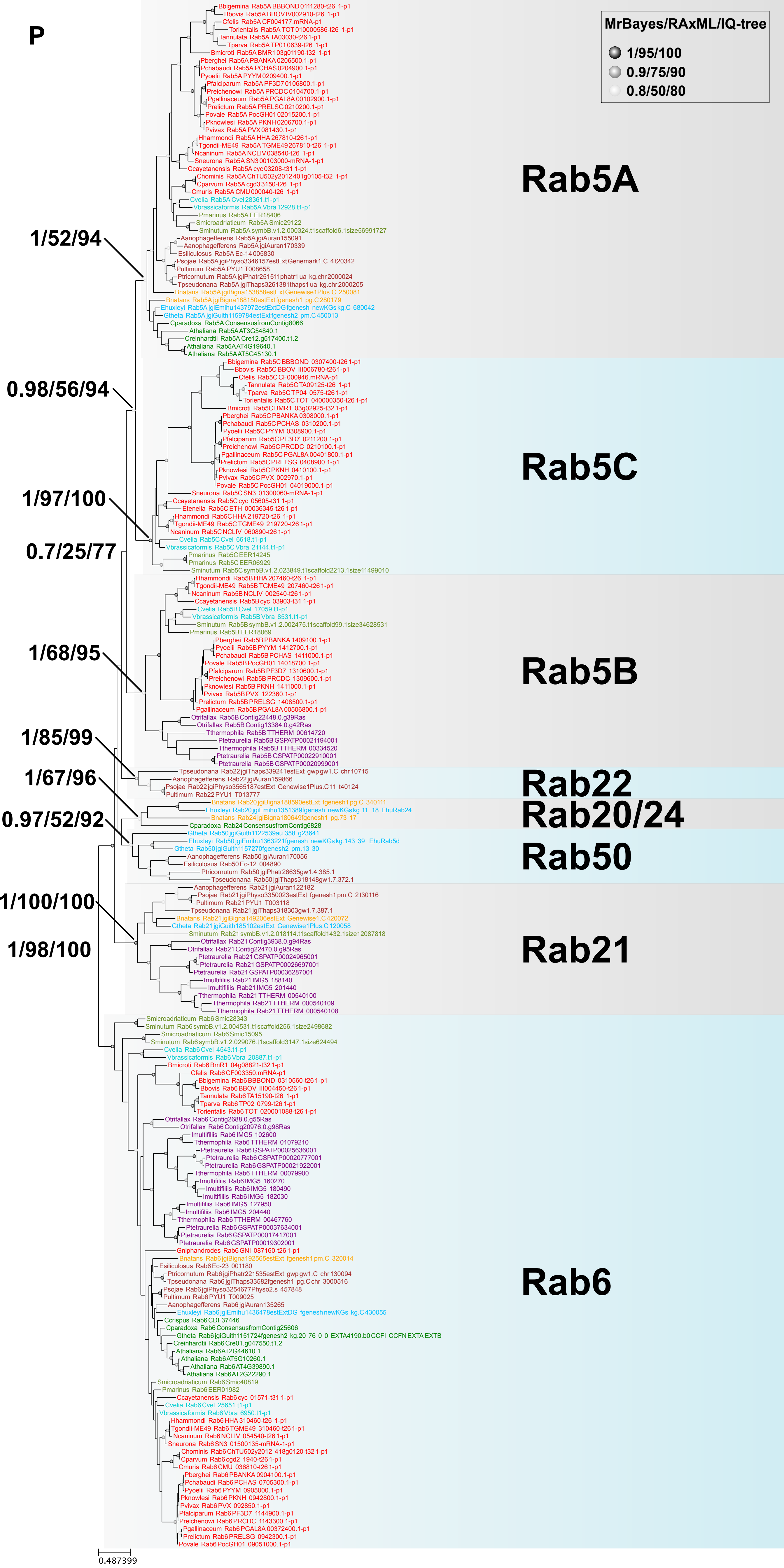

P
MrBayes/RAxML/IQ-tree
1/95/100
0.9/75/90
0.8/50/80
Rab5A
Rab5C
Rab5B
Rab22
Rab20/24
Rab50
Rab21
Rab6
1/52/94
0.98/56/94
1/97/100
0.7/25/77
1/68/95
1/85/99
1/67/96
0.97/52/92
1/100/100
1/98/100
0.487399

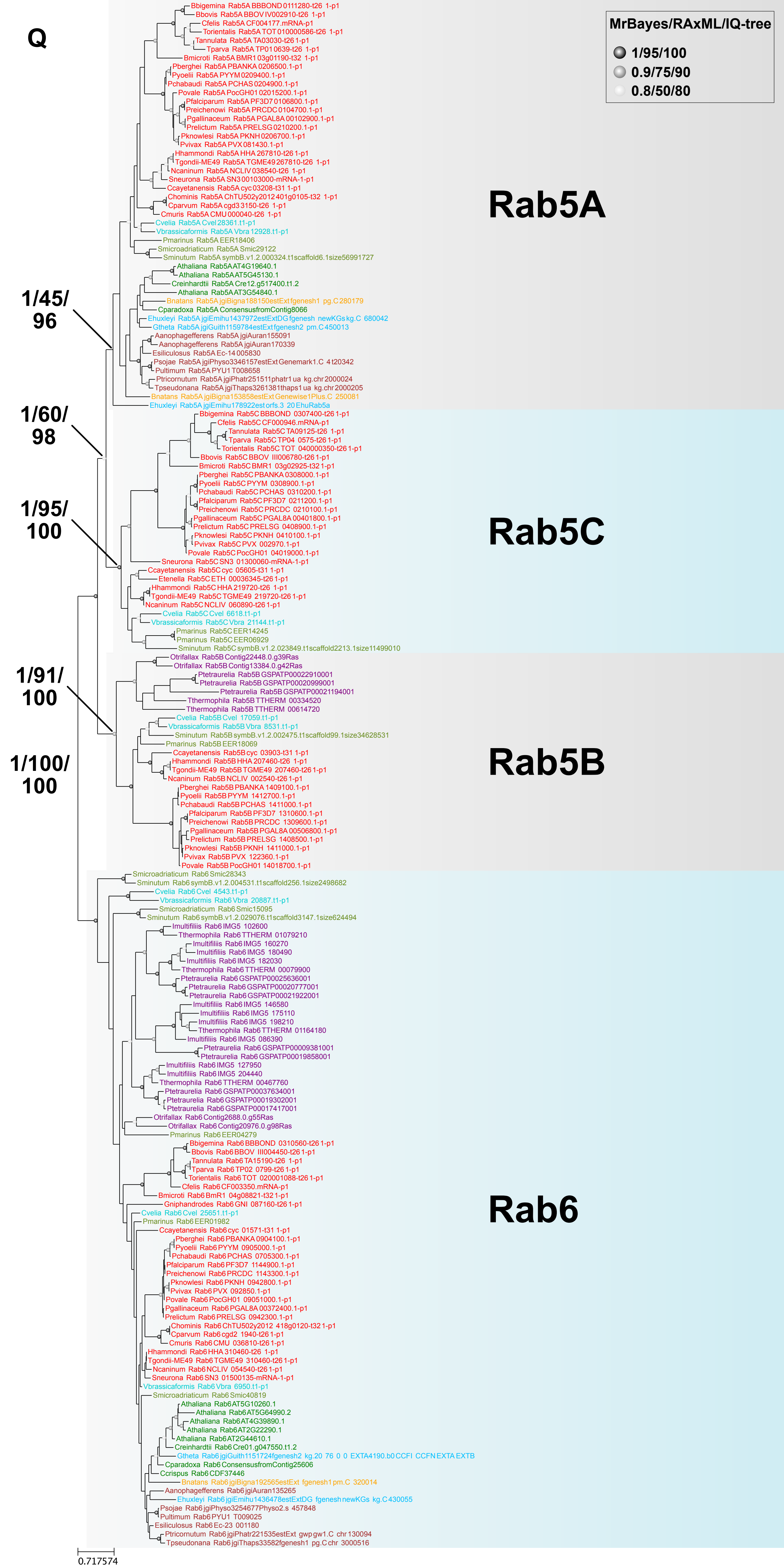
Q
MrBayes/RAxML/IQ-tree
1/95/100
0.9/75/90
0.8/50/80
Rab5A
Rab5C
Rab5B
Rab6
1/45/96
1/60/98
1/95/100
1/91/100
1/100/100
Bbigemina Rab5A BBBOND 0111280-t26 1-p1
Bbovis Rab5A BBOV IV002910-t26 1-p1
Cfelis Rab5A CF004177.mRNA-p1
Torientalis Rab5A TOT 010000586-t26 1-p1
Tannulata Rab5A TA03030-t26 1-p1
Tparva Rab5A TP01 0639-t26 1-p1
Bmicroti Rab5A BMR1 03g01190-t32 1-p1
Pberghei Rab5A PBANKA 0206500.1-p1
Pyoelii Rab5A PYYM 0209400.1-p1
Pchabaudi Rab5A PCHAS 0204900.1-p1
Povale Rab5A PocGH01 02015200.1-p1
Pfalciparum Rab5A PF3D7 0106800.1-p1
Preichenowi Rab5A PRCDC 0104700.1-p1
Pgallinaceum Rab5A PGAL8A 00102900.1-p1
Prelictum Rab5A PRELSG 0210200.1-p1
Pknowlesi Rab5A PKNH 0206700.1-p1
Pvivax Rab5A PVX 081430.1-p1
Hhammondi Rab5A HHA 267810-t26 1-p1
Tgondii-ME49 Rab5A TGME49 267810-t26 1-p1
Ncaninum Rab5A NCLIV 038540-t26 1-p1
Sneurona Rab5A SN3 00103000-mRNA-1-p1
Ccayetanensis Rab5A cyc 03208-t31 1-p1
Chominis Rab5A ChTU502y2012 401g0105-t32 1-p1
Cparvum Rab5A cgd3 3150-t26 1-p1
Cmuris Rab5A CMU 000040-t26 1-p1
Cvelia Rab5A Cvel 28361.t1-p1
Vbrassicaformis Rab5A Vbra 12928.t1-p1
Pmarinus Rab5A EER18406
Smicroadriaticum Rab5A Smic29122
Sminutum Rab5A symbB.v1.2.000324.t1scaffold6.1size56991727
Athaliana Rab5A AT4G19640.1
Athaliana Rab5A AT5G45130.1
Creinhardtii Rab5A Cre12.g517400.t1.2
Athaliana Rab5A AT3G54840.1
Bnatans Rab5A jgiBigna188150estExtfgenesh1 pg.C 280179
Cparadoxa Rab5A ConsensusfromContig8066
Ehuxleyi Rab5A jgiEmihu1437972estExtDGfgenesh newKGskg.C 680042
Gtheta Rab5A jgiGuith1159784estExtfgenesh2 pm.C 450013
Aanophagefferens Rab5A jgiAuran155091
Aanophagefferens Rab5A jgiAuran170339
Esiliculosus Rab5A Ec-14 005830
Psojae Rab5A jgiPhyso3346157estExtGenemark1.C 4 t20342
Pultimum Rab5A PYU1 T008658
Ptricornutum Rab5A jgiPhatr251511phatr1ua kg.chr 2000024
Tpseudonana Rab5A jgiThaps3261381thaps1ua kg.chr 2000205
Bnatans Rab5A jgiBigna153858estExtGenewise1Plus.C 250081
Ehuxleyi Rab5A jgiEmihu178922estorfs.3 20 EhuRab5a
Bbigemina Rab5C BBBOND 0307400-t26 1-p1
Cfelis Rab5C CF000946.mRNA-p1
Tannulata Rab5C TA09125-t26 1-p1
Tparva Rab5C TP04 0575-t26 1-p1
Torientalis Rab5C TOT 040000350-t26 1-p1
Bbovis Rab5C BBOV III006780-t26 1-p1
Bmicroti Rab5C BMR1 03g02925-t32 1-p1
Pberghei Rab5C PBANKA 0308000.1-p1
Pyoelii Rab5C PYYM 0308900.1-p1
Pchabaudi Rab5C PCHAS 0310200.1-p1
Pfalciparum Rab5C PF3D7 0211200.1-p1
Preichenowi Rab5C PRCDC 0210100.1-p1
Pgallinaceum Rab5C PGAL8A 00401800.1-p1
Prelictum Rab5C PRELSG 0408900.1-p1
Pknowlesi Rab5C PKNH 0410100.1-p1
Pvivax Rab5C PVX 002970.1-p1
Povale Rab5C PocGH01 04019000.1-p1
Sneurona Rab5C SN3 01300060-mRNA-1-p1
Ccayetanensis Rab5C cyc 05605-t31 1-p1
Etenella Rab5C ETH 00036345-t26 1-p1
Hhammondi Rab5C HHA 219720-t26 1-p1
Tgondii-ME49 Rab5C TGME49 219720-t26 1-p1
Ncaninum Rab5C NCLIV 060890-t26 1-p1
Cvelia Rab5C Cvel 6618.t1-p1
Vbrassicaformis Rab5C Vbra 21144.t1-p1
Pmarinus Rab5C EER14245
Pmarinus Rab5C EER06929
Sminutum Rab5C symbB.v1.2.023849.t1scaffold2213.1size11499010
Otrifallax Rab5B Contig22448.0.g39Ras
Otrifallax Rab5B Contig13384.0.g42Ras
Ptetraurelia Rab5B GSPATP00022910001
Ptetraurelia Rab5B GSPATP00020999001
Ptetraurelia Rab5B GSPATP00021194001
Tthermophila Rab5B TTHERM 00334520
Tthermophila Rab5B TTHERM 00614720
Cvelia Rab5B Cvel 17059.t1-p1
Vbrassicaformis Rab5B Vbra 8531.t1-p1
Sminutum Rab5B symbB.v1.2.002475.t1scaffold99.1size34628531
Pmarinus Rab5B EER18069
Ccayetanensis Rab5B cyc 03903-t31 1-p1
Hhammondi Rab5B HHA 207460-t26 1-p1
Tgondii-ME49 Rab5B TGME49 207460-t26 1-p1
Ncaninum Rab5B NCLIV 002540-t26 1-p1
Pberghei Rab5B PBANKA 1409100.1-p1
Pyoelii Rab5B PYYM 1412700.1-p1
Pchabaudi Rab5B PCHAS 1411000.1-p1
Pfalciparum Rab5B PF3D7 1310600.1-p1
Preichenowi Rab5B PRCDC 1309600.1-p1
Pgallinaceum Rab5B PGAL8A 00506800.1-p1
Prelictum Rab5B PRELSG 1408500.1-p1
Pknowlesi Rab5B PKNH 1411000.1-p1
Pvivax Rab5B PVX 122360.1-p1
Povale Rab5B PocGH01 14018700.1-p1
Smicroadriaticum Rab6 Smic28343
Sminutum Rab6 symbB.v1.2.004531.t1scaffold256.1size2498682
Cvelia Rab6 Cvel 4543.t1-p1
Vbrassicaformis Rab6 Vbra 20887.t1-p1
Smicroadriaticum Rab6 Smic15095
Sminutum Rab6 symbB.v1.2.029076.t1scaffold3147.1size624494
Imultifiliis Rab6 IMG5 102600
Tthermophila Rab6 TTHERM 01079210
Imultifiliis Rab6 IMG5 160270
Imultifiliis Rab6 IMG5 180490
Imultifiliis Rab6 IMG5 182030
Tthermophila Rab6 TTHERM 00079900
Ptetraurelia Rab6 GSPATP00025636001
Ptetraurelia Rab6 GSPATP00020777001
Ptetraurelia Rab6 GSPATP00021922001
Imultifiliis Rab6 IMG5 146580
Imultifiliis Rab6 IMG5 175110
Imultifiliis Rab6 IMG5 198210
Tthermophila Rab6 TTHERM 01164180
Imultifiliis Rab6 IMG5 086390
Ptetraurelia Rab6 GSPATP00009381001
Ptetraurelia Rab6 GSPATP00019858001
Imultifiliis Rab6 IMG5 127950
Imultifiliis Rab6 IMG5 204440
Tthermophila Rab6 TTHERM 00467760
Ptetraurelia Rab6 GSPATP00037634001
Ptetraurelia Rab6 GSPATP00019302001
Ptetraurelia Rab6 GSPATP00017417001
Otrifallax Rab6 Contig2688.0.g55Ras
Otrifallax Rab6 Contig20976.0.g98Ras
Pmarinus Rab6 EER04279
Bbigemina Rab6 BBBOND 0310560-t26 1-p1
Bbovis Rab6 BBOV III004450-t26 1-p1
Tannulata Rab6 TA15190-t26 1-p1
Tparva Rab6 TP02 0799-t26 1-p1
Torientalis Rab6 TOT 020001088-t26 1-p1
Cfelis Rab6 CF003350.mRNA-p1
Bmicroti Rab6 BmR1 04g08821-t32 1-p1
Gniphandrodes Rab6 GNI 087160-t26 1-p1
Cvelia Rab6 Cvel 25651.t1-p1
Pmarinus Rab6 EER01982
Ccayetanensis Rab6 cyc 01571-t31 1-p1
Pberghei Rab6 PBANKA 0904100.1-p1
Pyoelii Rab6 PYYM 0905000.1-p1
Pchabaudi Rab6 PCHAS 0705300.1-p1
Pfalciparum Rab6 PF3D7 1144900.1-p1
Preichenowi Rab6 PRCDC 1143300.1-p1
Pknowlesi Rab6 PKNH 0942800.1-p1
Pvivax Rab6 PVX 092850.1-p1
Povale Rab6 PocGH01 09051000.1-p1
Pgallinaceum Rab6 PGAL8A 00372400.1-p1
Prelictum Rab6 PRELSG 0942300.1-p1
Chominis Rab6 ChTU502y2012 418g0120-t32 1-p1
Cparvum Rab6 cgd2 1940-t26 1-p1
Cmuris Rab6 CMU 036810-t26 1-p1
Hhammondi Rab6 HHA 310460-t26 1-p1
Tgondii-ME49 Rab6 TGME49 310460-t26 1-p1
Ncaninum Rab6 NCLIV 054540-t26 1-p1
Sneurona Rab6 SN3 01500135-mRNA-1-p1
Vbrassicaformis Rab6 Vbra 6950.t1-p1
Smicroadriaticum Rab6 Smic40819
Athaliana Rab6 AT5G10260.1
Athaliana Rab6 AT5G64990.2
Athaliana Rab6 AT4G39890.1
Athaliana Rab6 AT2G22290.1
Athaliana Rab6 AT2G44610.1
Creinhardtii Rab6 Cre01.g047550.t1.2
Gtheta Rab6 jgiGuith1151724fgenesh2 kg.20 76 0 0 EXTA4190.b0 CCFI CCFN EXTA EXTB
Cparadoxa Rab6 ConsensusfromContig25606
Ccrispus Rab6 CDF37446
Bnatans Rab6 jgiBigna192565estExt fgenesh1 pm.C 320014
Aanophagefferens Rab6 jgiAuran135265
Ehuxleyi Rab6 jgiEmihu1436478estExtDG fgenesh newKGs kg.C 430055
Psojae Rab6 jgiPhyso3254677Physo2.s 457848
Pultimum Rab6 PYU1 T009025
Esiliculosus Rab6 Ec-23 001180
Ptricornutum Rab6 jgiPhatr221535estExt gwp gw1.C chr 130094
Tpseudonana Rab6 jgiThaps33582fgenesh1 pg.C chr 3000516
0.717574

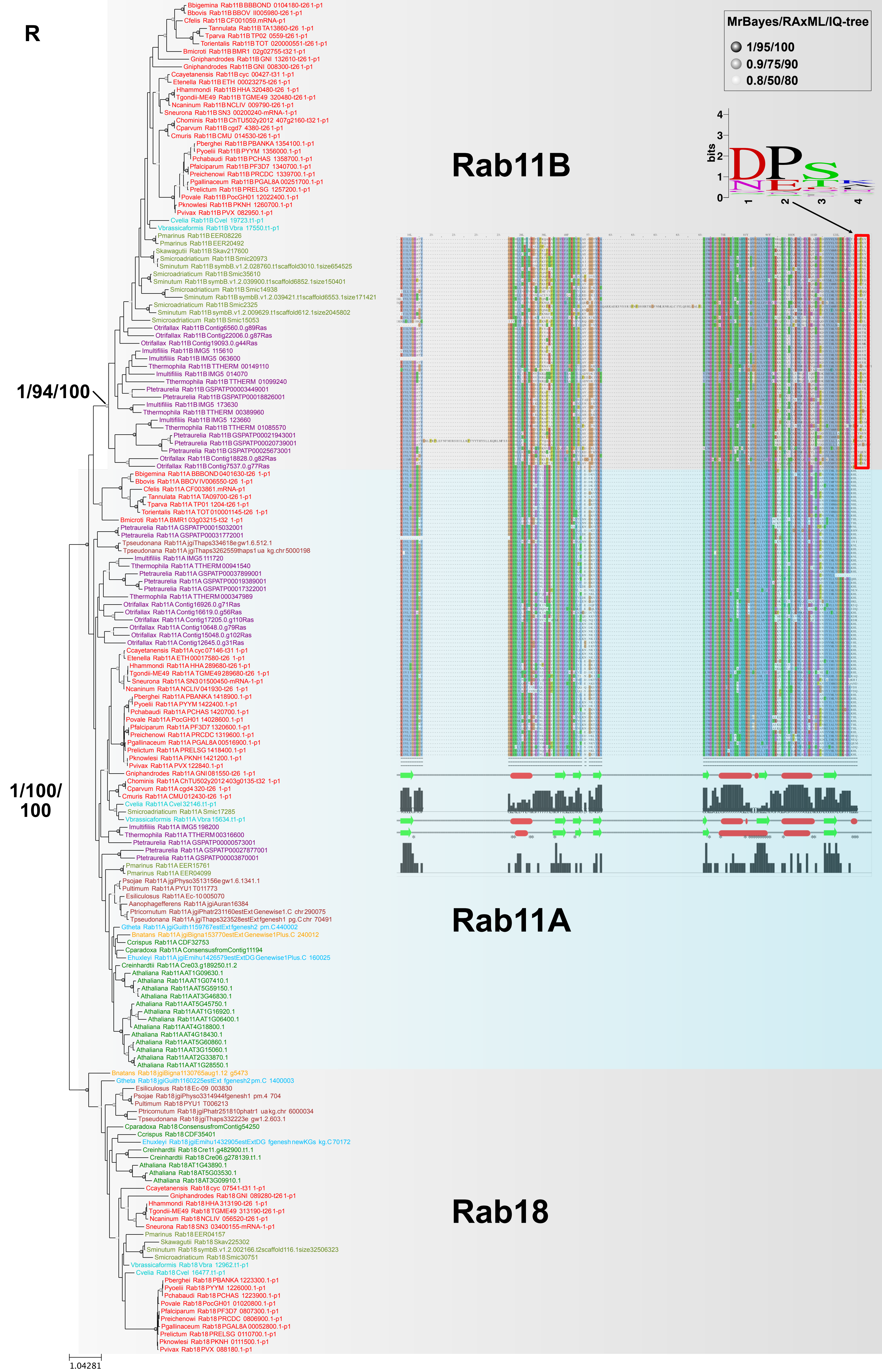

R
MrBayes/RAxML/IQ-tree
1/95/100
0.9/75/90
0.8/50/80
bits
Rab11B
Rab11A
Rab18
1/94/100
1/100/100
1.04281
Bbigemina Rab11B BBBOND 0104180-t26 1-p1
Bbovis Rab11B BBOV II005980-t26 1-p1
Cfelis Rab11B CF001059.mRNA-p1
Tannulata Rab11B TA13860-t26 1-p1
Tparva Rab11B TP02 0559-t26 1-p1
Torientalis Rab11B TOT 020000551-t26 1-p1
Bmicroti Rab11B BMR1 02g02755-t32 1-p1
Gniphandrodes Rab11B GNI 132610-t26 1-p1
Gniphandrodes Rab11B GNI 008300-t26 1-p1
Ccayetanensis Rab11B cyc 00427-t31 1-p1
Etenella Rab11B ETH 00023275-t26 1-p1
Hhammondi Rab11B HHA 320480-t26 1-p1
Tgondii-ME49 Rab11B TGME49 320480-t26 1-p1
Ncaninum Rab11B NCLIV 009790-t26 1-p1
Sneurona Rab11B SN3 00200240-mRNA-1-p1
Chominis Rab11B ChTU502y2012 407g2160-t32 1-p1
Cparvum Rab11B cgd7 4380-t26 1-p1
Cmuris Rab11B CMU 014530-t26 1-p1
Pberghei Rab11B PBANKA 1354100.1-p1
Pyoelii Rab11B PYYM 1356000.1-p1
Pchabaudi Rab11B PCHAS 1358700.1-p1
Pfalciparum Rab11B PF3D7 1340700.1-p1
Preichenowi Rab11B PRCDC 1339700.1-p1
Pgallinaceum Rab11B PGAL8A 00251700.1-p1
Prelictum Rab11B PRELSG 1257200.1-p1
Povale Rab11B PocGH01 12022400.1-p1
Pknowlesi Rab11B PKNH 1260700.1-p1
Pvivax Rab11B PVX 082950.1-p1
Cvelia Rab11B Cvel 19723.t1-p1
Vbrassicaformis Rab11B Vbra 17550.t1-p1
Pmarinus Rab11B EER08226
Pmarinus Rab11B EER20492
Skawagutii Rab11B Skav217600
Smicroadriaticum Rab11B Smic20973
Sminutum Rab11B symbB.v1.2.028760.t1scaffold3010.1size654525
Smicroadriaticum Rab11B Smic35610
Sminutum Rab11B symbB.v1.2.039900.t1scaffold6852.1size150401
Smicroadriaticum Rab11B Smic14938
Sminutum Rab11B symbB.v1.2.039421.t1scaffold6553.1size171421
Smicroadriaticum Rab11B Smic2325
Sminutum Rab11B symbB.v1.2.009629.t1scaffold612.1size2045802
Smicroadriaticum Rab11B Smic15053
Otrifallax Rab11B Contig6560.0.g89Ras
Otrifallax Rab11B Contig22006.0.g87Ras
Otrifallax Rab11B Contig19093.0.g44Ras
Imultifiliis Rab11B IMG5 115610
Imultifiliis Rab11B IMG5 063600
Tthermophila Rab11B TTHERM 00149110
Imultifiliis Rab11B IMG5 014070
Tthermophila Rab11B TTHERM 01099240
Ptetraurelia Rab11B GSPATP00003449001
Ptetraurelia Rab11B GSPATP00018826001
Imultifiliis Rab11B IMG5 173630
Tthermophila Rab11B TTHERM 00389960
Imultifiliis Rab11B IMG5 123660
Tthermophila Rab11B TTHERM 01085570
Ptetraurelia Rab11B GSPATP00021943001
Ptetraurelia Rab11B GSPATP00020739001
Ptetraurelia Rab11B GSPATP00025673001
Otrifallax Rab11B Contig18828.0.g82Ras
Otrifallax Rab11B Contig7537.0.g77Ras
Bbigemina Rab11A BBBOND 0401630-t26 1-p1
Bbovis Rab11A BBOV IV006550-t26 1-p1
Cfelis Rab11A CF003861.mRNA-p1
Tannulata Rab11A TA09700-t26 1-p1
Tparva Rab11A TP01 1204-t26 1-p1
Torientalis Rab11A TOT 010001145-t26 1-p1
Bmicroti Rab11A BMR1 03g03215-t32 1-p1
Ptetraurelia Rab11A GSPATP00015032001
Ptetraurelia Rab11A GSPATP00031772001
Tpseudonana Rab11A jgiThaps334618e gw1.6.512.1
Tpseudonana Rab11A jgiThaps3262559thaps1 ua kg.chr 5000198
Imultifiliis Rab11A IMG5 111720
Tthermophila Rab11A TTHERM 00941540
Ptetraurelia Rab11A GSPATP00037899001
Ptetraurelia Rab11A GSPATP00019389001
Ptetraurelia Rab11A GSPATP00017322001
Tthermophila Rab11A TTHERM 000347989
Otrifallax Rab11A Contig16926.0.g71Ras
Otrifallax Rab11A Contig16619.0.g56Ras
Otrifallax Rab11A Contig17205.0.g110Ras
Otrifallax Rab11A Contig10648.0.g79Ras
Otrifallax Rab11A Contig15048.0.g102Ras
Otrifallax Rab11A Contig12645.0.g31Ras
Ccayetanensis Rab11A cyc 07146-t31 1-p1
Etenella Rab11A ETH 00017580-t26 1-p1
Hhammondi Rab11A HHA 289680-t26 1-p1
Tgondii-ME49 Rab11A TGME49 289680-t26 1-p1
Sneurona Rab11A SN3 01500450-mRNA-1-p1
Ncaninum Rab11A NCLIV 041930-t26 1-p1
Pberghei Rab11A PBANKA 1418900.1-p1
Pyoelii Rab11A PYYM 1422400.1-p1
Pchabaudi Rab11A PCHAS 1420700.1-p1
Povale Rab11A PocGH01 14028600.1-p1
Pfalciparum Rab11A PF3D7 1320600.1-p1
Preichenowi Rab11A PRCDC 1319600.1-p1
Pgallinaceum Rab11A PGAL8A 00516900.1-p1
Prelictum Rab11A PRELSG 1418400.1-p1
Pknowlesi Rab11A PKNH 1421200.1-p1
Pvivax Rab11A PVX 122840.1-p1
Gniphandrodes Rab11A GNI 081550-t26 1-p1
Chominis Rab11A ChTU502y2012 403g0135-t32 1-p1
Cparvum Rab11A cgd4 320-t26 1-p1
Cmuris Rab11A CMU 012430-t26 1-p1
Cvelia Rab11A Cvel 32146.t1-p1
Smicroadriaticum Rab11A Smic17285
Vbrassicaformis Rab11A Vbra 15634.t1-p1
Imultifiliis Rab11A IMG5 198200
Tthermophila Rab11A TTHERM 00316600
Ptetraurelia Rab11A GSPATP00000573001
Ptetraurelia Rab11A GSPATP00027877001
Ptetraurelia Rab11A GSPATP00003870001
Pmarinus Rab11A EER15761
Pmarinus Rab11A EER04099
Psojae Rab11A jgiPhyso3513156e gw1.6.1341.1
Pultimum Rab11A PYU1 T011773
Esiliculosus Rab11A Ec-10 005070
Aanophagefferens Rab11A jgiAuran16384
Ptricornutum Rab11A jgiPhatr231160estExt Genewise1.C chr 290075
Tpseudonana Rab11A jgiThaps323528estExt fgenesh1 pg.C chr 70491
Gtheta Rab11A jgiGuith1159767estExt fgenesh2 pm.C 440002
Bnatans Rab11A jgiBigna153770estExt Genewise1Plus.C 240012
Ccrispus Rab11A CDF32753
Cparadoxa Rab11A ConsensusfromContig11194
Ehuxleyi Rab11A jgiEmihu1426579estExtDG Genewise1Plus.C 160025
Creinhardtii Rab11A Cre03.g189250.t1.2
Athaliana Rab11A AT1G09630.1
Athaliana Rab11A AT1G07410.1
Athaliana Rab11A AT5G59150.1
Athaliana Rab11A AT3G46830.1
Athaliana Rab11A AT5G45750.1
Athaliana Rab11A AT1G16920.1
Athaliana Rab11A AT1G06400.1
Athaliana Rab11A AT4G18800.1
Athaliana Rab11A AT4G18430.1
Athaliana Rab11A AT5G60860.1
Athaliana Rab11A AT3G15060.1
Athaliana Rab11A AT2G33870.1
Athaliana Rab11A AT1G28550.1
Bnatans Rab18 jgiBigna1130765aug1.12 g5473
Gtheta Rab18 jgiGuith1160225estExt fgenesh2 pm.C 1400003
Esiliculosus Rab18 Ec-09 003830
Psojae Rab18 jgiPhyso3314944fgenesh1 pm.4 704
Pultimum Rab18 PYU1 T006213
Ptricornutum Rab18 jgiPhatr251810phatr1 ua kg.chr 6000034
Tpseudonana Rab18 jgiThaps332223e gw1.2.603.1
Cparadoxa Rab18 ConsensusfromContig54250
Ccrispus Rab18 CDF35401
Ehuxleyi Rab18 jgiEmihu1432905estExtDG fgenesh newKGs kg.C 70172
Creinhardtii Rab18 Cre11.g482900.t1.1
Creinhardtii Rab18 Cre06.g278139.t1.1
Athaliana Rab18 AT1G43890.1
Athaliana Rab18 AT5G03530.1
Athaliana Rab18 AT3G09910.1
Ccayetanensis Rab18 cyc 07541-t31 1-p1
Gniphandrodes Rab18 GNI 089280-t26 1-p1
Hhammondi Rab18 HHA 313190-t26 1-p1
Tgondii-ME49 Rab18 TGME49 313190-t26 1-p1
Ncaninum Rab18 NCLIV 056520-t26 1-p1
Sneurona Rab18 SN3 03400155-mRNA-1-p1
Pmarinus Rab18 EER04157
Skawagutii Rab18 Skav225302
Sminutum Rab18 symbB.v1.2.002166.t2scaffold116.1size32506323
Smicroadriaticum Rab18 Smic30751
Vbrassicaformis Rab18 Vbra 12962.t1-p1
Cvelia Rab18 Cvel 16477.t1-p1
Pberghei Rab18 PBANKA 1223300.1-p1
Pyoelii Rab18 PYYM 1226000.1-p1
Pchabaudi Rab18 PCHAS 1223900.1-p1
Povale Rab18 PocGH01 01020800.1-p1
Pfalciparum Rab18 PF3D7 0807300.1-p1
Preichenowi Rab18 PRCDC 0806900.1-p1
Pgallinaceum Rab18 PGAL8A 00052800.1-p1
Prelictum Rab18 PRELSG 0110700.1-p1
Pknowlesi Rab18 PKNH 0111500.1-p1
Pvivax Rab18 PVX 088180.1-p1

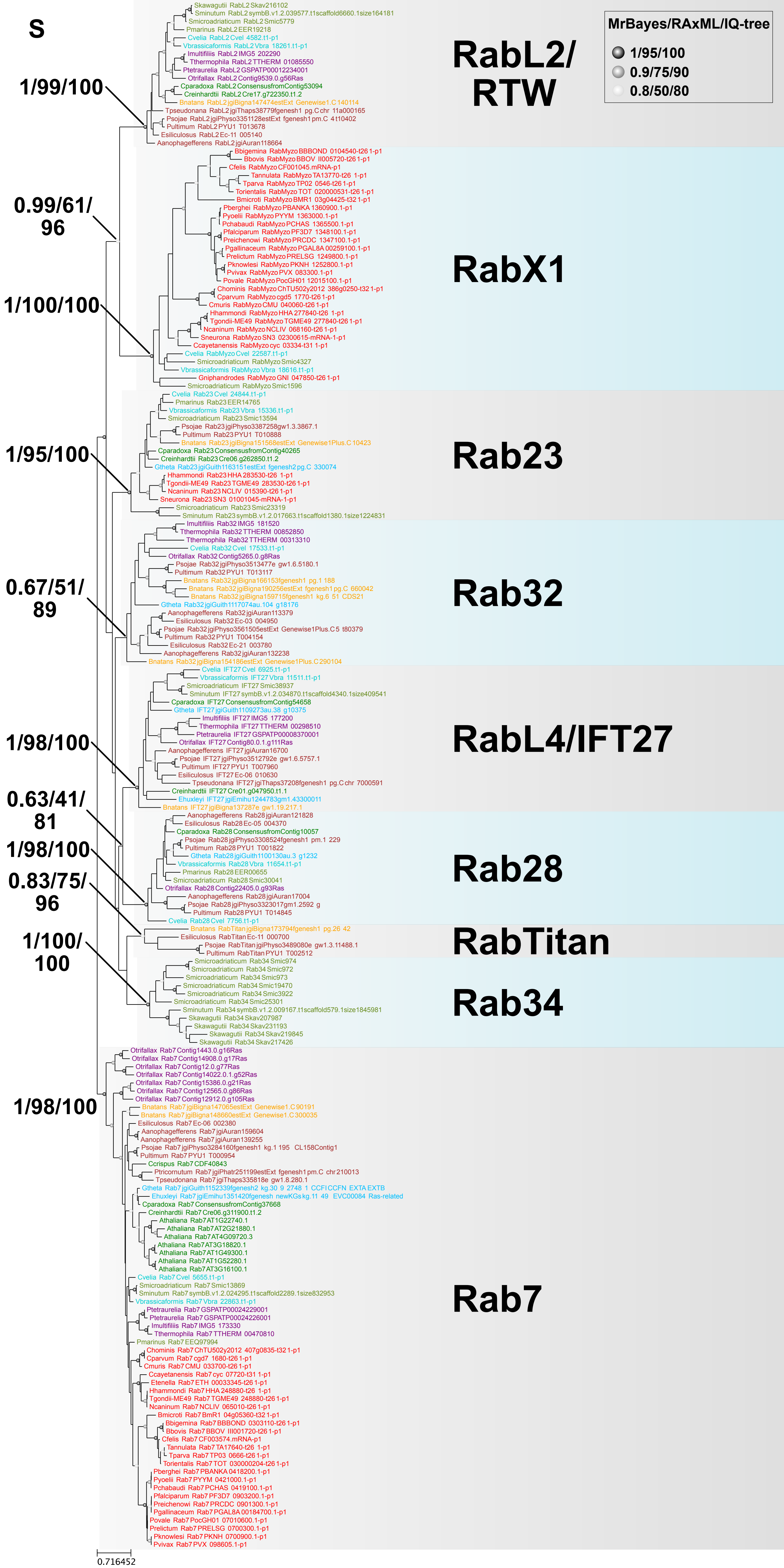
S
MrBayes/RAxML/IQ-tree
1/95/100
0.9/75/90
0.8/50/80
RabL2/RTW
RabX1
Rab23
Rab32
RabL4/IFT27
Rab28
RabTitan
Rab34
Rab7
1/99/100
0.99/61/96
1/100/100
1/95/100
0.67/51/89
1/98/100
0.63/41/81
1/98/100
0.83/75/96
1/100/100
1/98/100
0.716452
Skawagutii RabL2 Skav216102
Sminutum RabL2 symbB.v1.2.039577.t1scaffold6660.1size164181
Smicroadriaticum RabL2 Smic5779
Pmarinus RabL2 EER19218
Cvelia RabL2 Cvel 4582.t1-p1
Vbrassicaformis RabL2 Vbra 18261.t1-p1
Imultifiliis RabL2 IMG5 202290
Tthermophila RabL2 TTHERM 01085550
Ptetraurelia RabL2 GSPATP00012234001
Otrifallax RabL2 Contig9539.0.g56Ras
Cparadoxa RabL2 ConsensusfromContig53094
Creinhardtii RabL2 Cre17.g722350.t1.2
Bnatans RabL2 jgiBigna147474estExt Genewise1.C 140114
Tpseudonana RabL2 jgiThaps38779fgenesh1 pg.C chr 11a000165
Psojae RabL2 jgiPhyso3351128estExt fgenesh1 pm.C 4 t10402
Pultimum RabL2 PYU1 T013678
Esiliculosus RabL2 Ec-11 005140
Aanophagefferens RabL2 jgiAuran118664
Bbigemina RabMyzo BBBOND 0104540-t26 1-p1
Bbovis RabMyzo BBOV II005720-t26 1-p1
Cfelis RabMyzo CF001045.mRNA-p1
Tannulata RabMyzo TA13770-t26 1-p1
Tparva RabMyzo TP02 0546-t26 1-p1
Torientalis RabMyzo TOT 020000531-t26 1-p1
Bmicroti RabMyzo BMR1 03g04425-t32 1-p1
Pberghei RabMyzo PBANKA 1360900.1-p1
Pyoelii RabMyzo PYYM 1363000.1-p1
Pchabaudi RabMyzo PCHAS 1365500.1-p1
Pfalciparum RabMyzo PF3D7 1348100.1-p1
Preichenowi RabMyzo PRCDC 1347100.1-p1
Pgallinaceum RabMyzo PGAL8A 00259100.1-p1
Prelictum RabMyzo PRELSG 1249800.1-p1
Pknowlesi RabMyzo PKNH 1252800.1-p1
Pvivax RabMyzo PVX 083300.1-p1
Povale RabMyzo PocGH01 12015100.1-p1
Chominis RabMyzo ChTU502y2012 386g0250-t32 1-p1
Cparvum RabMyzo cgd5 1770-t26 1-p1
Cmuris RabMyzo CMU 040060-t26 1-p1
Hhammondi RabMyzo HHA 277840-t26 1-p1
Tgondii-ME49 RabMyzo TGME49 277840-t26 1-p1
Ncaninum RabMyzo NCLIV 068160-t26 1-p1
Sneurona RabMyzo SN3 02300615-mRNA-1-p1
Ccayetanensis RabMyzo cyc 03334-t31 1-p1
Cvelia RabMyzo Cvel 22587.t1-p1
Smicroadriaticum RabMyzo Smic4327
Vbrassicaformis RabMyzo Vbra 18616.t1-p1
Gniphandrodes RabMyzo GNI 047850-t26 1-p1
Smicroadriaticum RabMyzo Smic1596
Cvelia Rab23 Cvel 24844.t1-p1
Pmarinus Rab23 EER14765
Vbrassicaformis Rab23 Vbra 15336.t1-p1
Smicroadriaticum Rab23 Smic13594
Psojae Rab23 jgiPhyso3387258gw1.3.3867.1
Pultimum Rab23 PYU1 T010888
Bnatans Rab23 jgiBigna151568estExt Genewise1Plus.C 10423
Cparadoxa Rab23 ConsensusfromContig40265
Creinhardtii Rab23 Cre06.g262850.t1.2
Gtheta Rab23 jgiGuith1163151estExt fgenesh2 pg.C 330074
Hhammondi Rab23 HHA 283530-t26 1-p1
Tgondii-ME49 Rab23 TGME49 283530-t26 1-p1
Ncaninum Rab23 NCLIV 015390-t26 1-p1
Sneurona Rab23 SN3 01001045-mRNA-1-p1
Smicroadriaticum Rab23 Smic23319
Sminutum Rab23 symbB.v1.2.017663.t1scaffold1380.1size1224831
Imultifiliis Rab32 IMG5 181520
Tthermophila Rab32 TTHERM 00852870
Tthermophila Rab32 TTHERM 00313310
Cvelia Rab32 Cvel 17533.t1-p1
Otrifallax Rab32 Contig5265.0.g8Ras
Psojae Rab32 jgiPhyso3513477e gw1.6.5180.1
Pultimum Rab32 PYU1 T013117
Bnatans Rab32 jgiBigna166153fgenesh1 pg.1 188
Bnatans Rab32 jgiBigna190256estExt fgenesh1 pg.C 660042
Bnatans Rab32 jgiBigna159715fgenesh1 kg.6 51 CDS21
Gtheta Rab32 jgiGuith1117074au.104 g18176
Aanophagefferens Rab32 jgiAuran113379
Esiliculosus Rab32 Ec-03 004950
Psojae Rab32 jgiPhyso3561505estExt Genewise1Plus.C5 t80379
Pultimum Rab32 PYU1 T004154
Esiliculosus Rab32 Ec-21 003780
Aanophagefferens Rab32 jgiAuran132238
Bnatans Rab32 jgiBigna154186estExt Genewise1Plus.C 290104
Cvelia IFT27 Cvel 6925.t1-p1
Vbrassicaformis IFT27 Vbra 11511.t1-p1
Smicroadriaticum IFT27 Smic38937
Sminutum IFT27 symbB.v1.2.034870.t1scaffold4340.1size409541
Cparadoxa IFT27 ConsensusfromContig54658
Gtheta IFT27 jgiGuith1109273au.38 g10375
Imultifiliis IFT27 IMG5 177200
Tthermophila IFT27 TTHERM 00298510
Ptetraurelia IFT27 GSPATP00008370001
Otrifallax IFT27 Contig80.0.1.g111Ras
Aanophagefferens IFT27 jgiAuran16700
Psojae IFT27 jgiPhyso3512792e gw1.6.5757.1
Pultimum IFT27 PYU1 T007960
Esiliculosus IFT27 Ec-06 010630
Tpseudonana IFT27 jgiThaps37208fgenesh1 pg.C chr 7000591
Creinhardtii IFT27 Cre01.g047950.t1.1
Ehuxleyi IFT27 jgiEmihu1244783gm1.43300011
Bnatans IFT27 jgiBigna137287e gw1.19.217.1
Aanophagefferens Rab28 jgiAuran121828
Esiliculosus Rab28 Ec-05 004370
Cparadoxa Rab28 ConsensusfromContig10057
Psojae Rab28 jgiPhyso3308524fgenesh1 pm.1 229
Pultimum Rab28 PYU1 T001822
Gtheta Rab28 jgiGuith1100130au.3 g1232
Vbrassicaformis Rab28 Vbra 11654.t1-p1
Pmarinus Rab28 EER00655
Smicroadriaticum Rab28 Smic30041
Otrifallax Rab28 Contig22405.0.g93Ras
Aanophagefferens Rab28 jgiAuran17004
Psojae Rab28 jgiPhyso3323017gm1.2592 g
Pultimum Rab28 PYU1 T014845
Cvelia Rab28 Cvel 7756.t1-p1
Bnatans RabTitan jgiBigna173794fgenesh1 pg.26 42
Esiliculosus RabTitan Ec-11 000700
Psojae RabTitan jgiPhyso3489080e gw1.3.11488.1
Pultimum RabTitan PYU1 T002512
Smicroadriaticum Rab34 Smic974
Smicroadriaticum Rab34 Smic972
Smicroadriaticum Rab34 Smic973
Smicroadriaticum Rab34 Smic19470
Smicroadriaticum Rab34 Smic3922
Smicroadriaticum Rab34 Smic25301
Sminutum Rab34 symbB.v1.2.009167.t1scaffold579.1size1845981
Skawagutii Rab34 Skav207987
Skawagutii Rab34 Skav231193
Skawagutii Rab34 Skav219845
Skawagutii Rab34 Skav217426
Otrifallax Rab7 Contig1443.0.g16Ras
Otrifallax Rab7 Contig14908.0.g17Ras
Otrifallax Rab7 Contig12.0.g77Ras
Otrifallax Rab7 Contig14022.0.1.g52Ras
Otrifallax Rab7 Contig15386.0.g21Ras
Otrifallax Rab7 Contig12565.0.g86Ras
Otrifallax Rab7 Contig12912.0.g105Ras
Bnatans Rab7 jgiBigna147065estExt Genewise1.C 90191
Bnatans Rab7 jgiBigna148660estExt Genewise1.C 300035
Esiliculosus Rab7 Ec-06 002380
Aanophagefferens Rab7 jgiAuran159604
Aanophagefferens Rab7 jgiAuran139255
Psojae Rab7 jgiPhyso3284160fgenesh1 kg.1 195 CL158Contig1
Pultimum Rab7 PYU1 T000954
Ccrispus Rab7 CDF40843
Ptricornutum Rab7 jgiPhatr251199estExt fgenesh1 pm.C chr 210013
Tpseudonana Rab7 jgiThaps335818e gw1.8.280.1
Gtheta Rab7 jgiGuith1152339fgenesh2 kg.30 9 2748 1 CCFICCFN EXTAEXTB
Ehuxleyi Rab7 jgiEmihu1351420fgenesh newKGs kg.11 49 EVC00084 Ras-related
Cparadoxa Rab7 ConsensusfromContig37668
Creinhardtii Rab7 Cre06.g311900.t1.2
Athaliana Rab7 AT1G22740.1
Athaliana Rab7 AT2G21880.1
Athaliana Rab7 AT4G09720.3
Athaliana Rab7 AT3G18820.1
Athaliana Rab7 AT1G49300.1
Athaliana Rab7 AT1G52280.1
Athaliana Rab7 AT3G16100.1
Cvelia Rab7 Cvel 5655.t1-p1
Smicroadriaticum Rab7 Smic13869
Sminutum Rab7 symbB.v1.2.024295.t1scaffold2289.1size832953
Vbrassicaformis Rab7 Vbra 22863.t1-p1
Ptetraurelia Rab7 GSPATP00024229001
Ptetraurelia Rab7 GSPATP00024226001
Imultifiliis Rab7 IMG5 173330
Tthermophila Rab7 TTHERM 00470810
Pmarinus Rab7 EEQ97994
Chominis Rab7 ChTU502y2012 407g0835-t32 1-p1
Cparvum Rab7 cgd7 1680-t26 1-p1
Cmuris Rab7 CMU 033700-t26 1-p1
Ccayetanensis Rab7 cyc 07720-t31 1-p1
Etenella Rab7 ETH 00033345-t26 1-p1
Hhammondi Rab7 HHA 248880-t26 1-p1
Tgondii-ME49 Rab7 TGME49 248880-t26 1-p1
Ncaninum Rab7 NCLIV 065010-t26 1-p1
Bmicroti Rab7 BmR1 04g05360-t32 1-p1
Bbigemina Rab7 BBBOND 0303110-t26 1-p1
Bbovis Rab7 BBOV III001720-t26 1-p1
Cfelis Rab7 CF003574.mRNA-p1
Tannulata Rab7 TA17640-t26 1-p1
Tparva Rab7 TP03 0666-t26 1-p1
Torientalis Rab7 TOT 030000204-t26 1-p1
Pberghei Rab7 PBANKA 0418200.1-p1
Pyoelii Rab7 PYYM 0421000.1-p1
Pchabaudi Rab7 PCHAS 0419100.1-p1
Pfalciparum Rab7 PF3D7 0903200.1-p1
Preichenowi Rab7 PRCDC 0901300.1-p1
Pgallinaceum Rab7 PGAL8A 00184700.1-p1
Povale Rab7 PocGH01 07010600.1-p1
Prelictum Rab7 PRELSG 0700300.1-p1
Pknowlesi Rab7 PKNH 0700900.1-p1
Pvivax Rab7 PVX 098605.1-p1

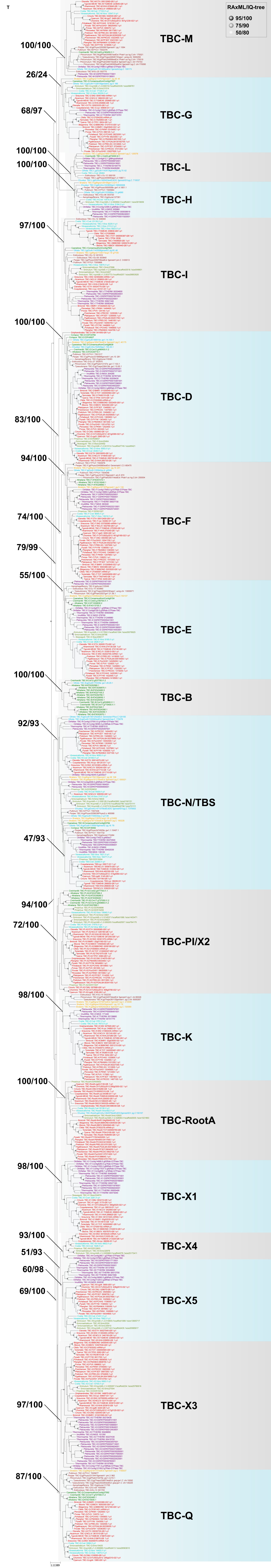

RAxML/IQ-tree
95/100
75/90
50/80
TBC-M
TBC-G
TBC-L
TBC-H
TBC-I
TBC-D
TBC-F
TBC-E
TBC-B
TBC-N/TBS
TBC-PI/X2
TBC-K
TBC-RootA
TBC-X1
TBC-X4
TBC-X5
TBC-X3
TBC-Q
100/100
26/24
68/97
100/100
100/100
97/100
100/100
83/100
94/100
74/100
79/99
55/100
100/100
92/93
47/93
94/100
72/100
98/100
100/100
98/100
93/100
51/93
60/98
69/100
97/100
87/100

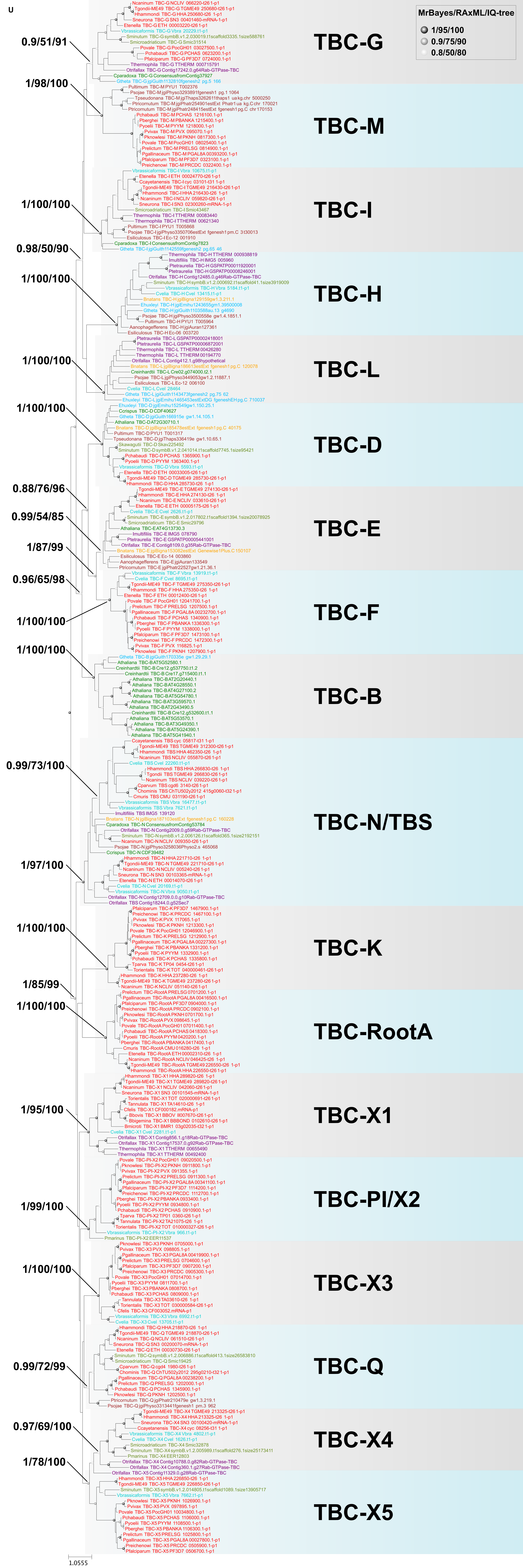
U
MrBayes/RAxML/IQ-tree
1/95/100
0.9/75/90
0.8/50/80
0.9/51/91
TBC-G
1/98/100
TBC-M
1/100/100
TBC-I
0.98/50/90
1/100/100
TBC-H
1/100/100
TBC-L
1/100/100
TBC-D
0.88/76/96
0.99/54/85
TBC-E
1/87/99
0.96/65/98
1/100/100
TBC-F
1/100/100
TBC-B
0.99/73/100
TBC-N/TBS
1/97/100
1/100/100
TBC-K
1/85/99
1/100/100
TBC-RootA
1/95/100
TBC-X1
1/99/100
TBC-PI/X2
1/100/100
TBC-X3
0.99/72/99
TBC-Q
0.97/69/100
TBC-X4
1/78/100
TBC-X5
1.0555

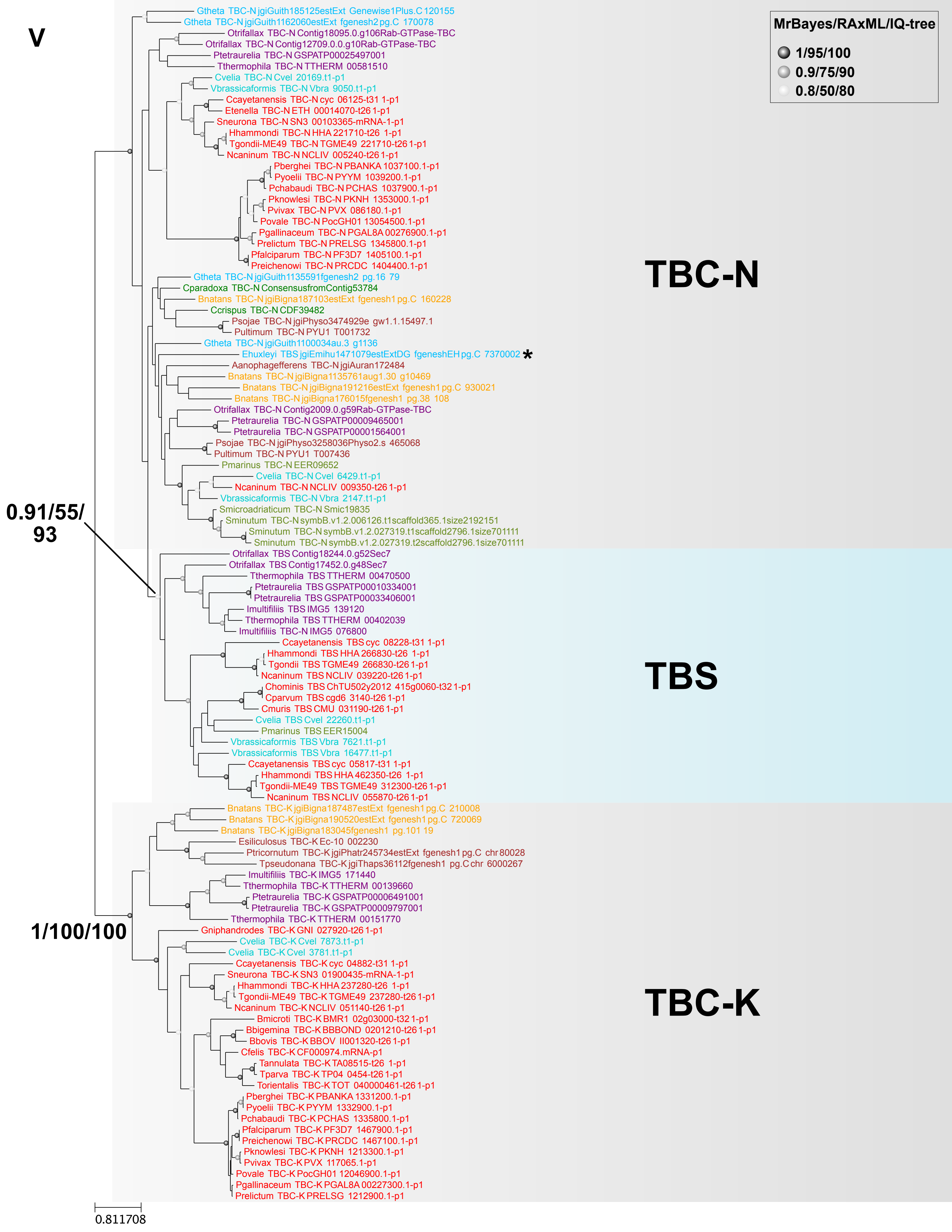
V
MrBayes/RAxML/IQ-tree
1/95/100
0.9/75/90
0.8/50/80
Gtheta TBC-N jgiGuith185125estExt Genewise1Plus.C 120155
Gtheta TBC-N jgiGuith1162060estExt fgenesh2 pg.C 170078
Otrifallax TBC-N Contig18095.0.g106Rab-GTPase-TBC
Otrifallax TBC-N Contig12709.0.0.g10Rab-GTPase-TBC
Ptetraurelia TBC-N GSPATP00025497001
Tthermophila TBC-N TTHERM 00581510
Cvelia TBC-N Cvel 20169.t1-p1
Vbrassicaformis TBC-N Vbra 9050.t1-p1
Ccayetanensis TBC-N cyc 06125-t31 1-p1
Etenella TBC-N ETH 00014070-t26 1-p1
Sneurona TBC-N SN3 00103365-mRNA-1-p1
Hhammondi TBC-N HHA 221710-t26 1-p1
Tgondii-ME49 TBC-N TGME49 221710-t26 1-p1
Ncaninum TBC-N NCLIV 005240-t26 1-p1
Pberghei TBC-N PBANKA 1037100.1-p1
Pyoelii TBC-N PYYM 1039200.1-p1
Pchabaudi TBC-N PCHAS 1037900.1-p1
Pknowlesi TBC-N PKNH 1353000.1-p1
Pvivax TBC-N PVX 086180.1-p1
Povale TBC-N PocGH01 13054500.1-p1
Pgallinaceum TBC-N PGAL8A 00276900.1-p1
Prelictum TBC-N PRELSG 1345800.1-p1
Pfalciparum TBC-N PF3D7 1405100.1-p1
Preichenowi TBC-N PRCDC 1404400.1-p1
Gtheta TBC-N jgiGuith1135591fgenesh2 pg.16 79
Cparadoxa TBC-N ConsensusfromContig53784
Bnatans TBC-N jgiBigna187103estExt fgenesh1 pg.C 160228
Ccrispus TBC-N CDF39482
Psojae TBC-N jgiPhyso3474929e gw1.1.15497.1
Pultimum TBC-N PYU1 T001732
Gtheta TBC-N jgiGuith1100034au.3 g1136
Ehuxleyi TBS jgiEmihu1471079estExtDG fgeneshEH pg.C 7370002 *
Aanophagefferens TBC-N jgiAuran172484
Bnatans TBC-N jgiBigna1135761aug1.30 g10469
Bnatans TBC-N jgiBigna191216estExt fgenesh1 pg.C 930021
Bnatans TBC-N jgiBigna176015fgenesh1 pg.38 108
Otrifallax TBC-N Contig2009.0.g59Rab-GTPase-TBC
Ptetraurelia TBC-N GSPATP00009465001
Ptetraurelia TBC-N GSPATP00001564001
Psojae TBC-N jgiPhyso3258036Physo2.s 465068
Pultimum TBC-N PYU1 T007436
Pmarinus TBC-N EER09652
Cvelia TBC-N Cvel 6429.t1-p1
Ncaninum TBC-N NCLIV 009350-t26 1-p1
Vbrassicaformis TBC-N Vbra 2147.t1-p1
Smicroadriaticum TBC-N Smic19835
Sminutum TBC-N symbB.v1.2.006126.t1scaffold365.1size2192151
Sminutum TBC-N symbB.v1.2.027319.t1scaffold2796.1size701111
Sminutum TBC-N symbB.v1.2.027319.t2scaffold2796.1size701111
TBC-N
0.91/55/
93
Otrifallax TBS Contig18244.0.g52Sec7
Otrifallax TBS Contig17452.0.g48Sec7
Tthermophila TBS TTHERM 00470500
Ptetraurelia TBS GSPATP00010334001
Ptetraurelia TBS GSPATP00033406001
Imultifiliis TBS IMG5 139120
Tthermophila TBS TTHERM 00402039
Imultifiliis TBC-N IMG5 076800
Ccayetanensis TBS cyc 08228-t31 1-p1
Hhammondi TBS HHA 266830-t26 1-p1
Tgondii TBS TGME49 266830-t26 1-p1
Ncaninum TBS NCLIV 039220-t26 1-p1
Chominis TBS ChTU502y2012 415g0060-t32 1-p1
Cparvum TBS cgd6 3140-t26 1-p1
Cmuris TBS CMU 031190-t26 1-p1
Cvelia TBS Cvel 22260.t1-p1
Pmarinus TBS EER15004
Vbrassicaformis TBS Vbra 7621.t1-p1
Vbrassicaformis TBS Vbra 16477.t1-p1
Ccayetanensis TBS cyc 05817-t31 1-p1
Hhammondi TBS HHA 462350-t26 1-p1
Tgondii-ME49 TBS TGME49 312300-t26 1-p1
Ncaninum TBS NCLIV 055870-t26 1-p1
TBS
Bnatans TBC-K jgiBigna187487estExt fgenesh1 pg.C 210008
Bnatans TBC-K jgiBigna190520estExt fgenesh1 pg.C 720069
Bnatans TBC-K jgiBigna183045fgenesh1 pg.101 19
Esiliculosus TBC-K Ec-10 002230
Ptricornutum TBC-K jgiPhatr245734estExt fgenesh1 pg.C chr80028
Tpseudonana TBC-K jgiThaps36112fgenesh1 pg.C chr 6000267
Imultifiliis TBC-K IMG5 171440
Tthermophila TBC-K TTHERM 00139660
Ptetraurelia TBC-K GSPATP00006491001
Ptetraurelia TBC-K GSPATP00009797001
Tthermophila TBC-K TTHERM 00151770
1/100/100
Gniphandrodes TBC-K GNI 027920-t26 1-p1
Cvelia TBC-K Cvel 7873.t1-p1
Cvelia TBC-K Cvel 3781.t1-p1
Ccayetanensis TBC-K cyc 04882-t31 1-p1
Sneurona TBC-K SN3 01900435-mRNA-1-p1
Hhammondi TBC-K HHA 237280-t26 1-p1
Tgondii-ME49 TBC-K TGME49 237280-t26 1-p1
Ncaninum TBC-K NCLIV 051140-t26 1-p1
Bmicroti TBC-K BMR1 02g03000-t32 1-p1
Bbigemina TBC-K BBBOND 0201210-t26 1-p1
Bbovis TBC-K BBOV II001320-t26 1-p1
Cfelis TBC-K CF000974.mRNA-p1
Tannulata TBC-K TA08515-t26 1-p1
Tparva TBC-K TP04 0454-t26 1-p1
Torientalis TBC-K TOT 040000461-t26 1-p1
Pberghei TBC-K PBANKA 1331200.1-p1
Pyoelii TBC-K PYYM 1332900.1-p1
Pchabaudi TBC-K PCHAS 1335800.1-p1
Pfalciparum TBC-K PF3D7 1467900.1-p1
Preichenowi TBC-K PRCDC 1467100.1-p1
Pknowlesi TBC-K PKNH 1213300.1-p1
Pvivax TBC-K PVX 117065.1-p1
Povale TBC-K PocGH01 12046900.1-p1
Pgallinaceum TBC-K PGAL8A 00227300.1-p1
Prelictum TBC-K PRELSG 1212900.1-p1
TBC-K
0.811708

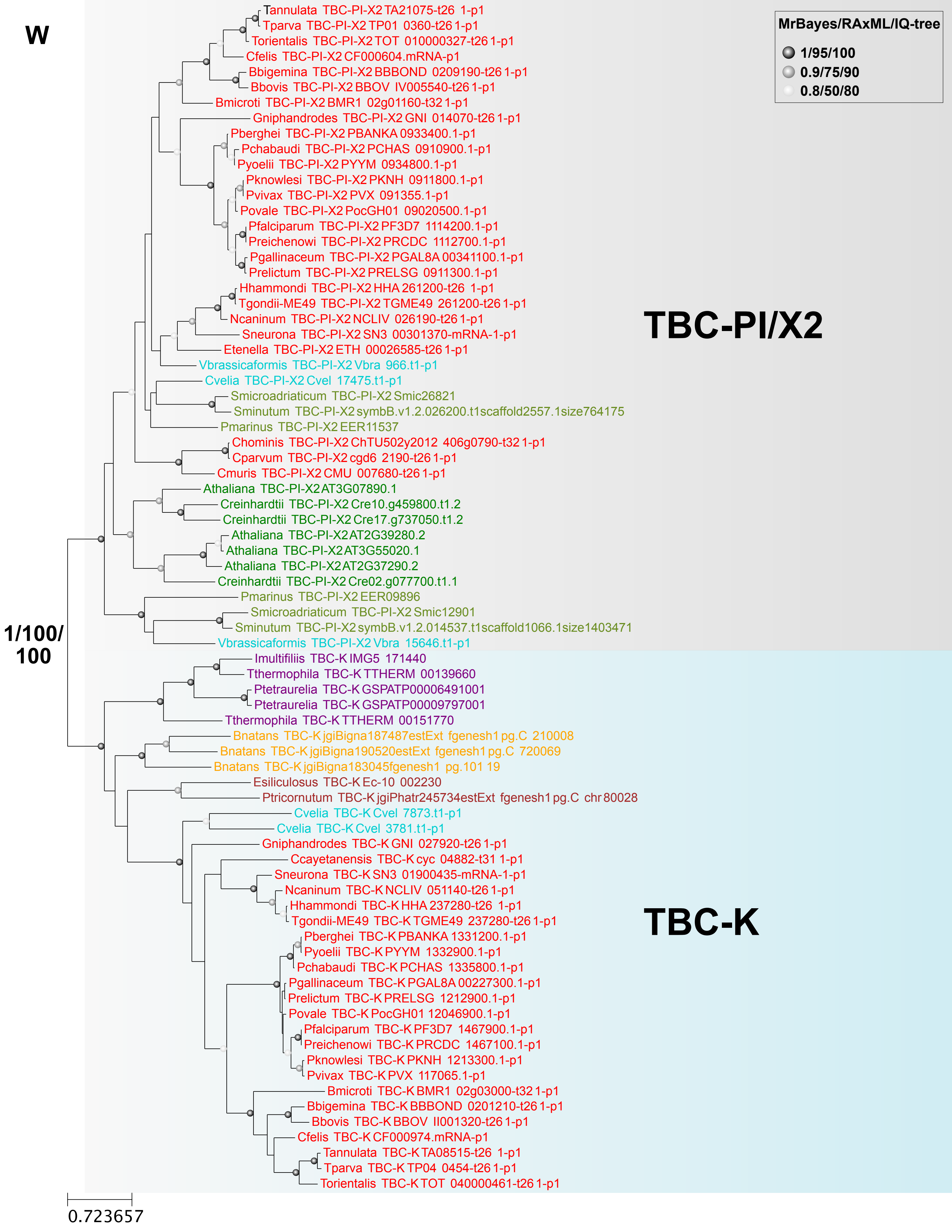
W
MrBayes/RAxML/IQ-tree
1/95/100
0.9/75/90
0.8/50/80
TBC-PI/X2
TBC-K
1/100/100
Tannulata TBC-PI-X2 TA21075-t26 1-p1
Tparva TBC-PI-X2 TP01 0360-t26 1-p1
Torientalis TBC-PI-X2 TOT 010000327-t26 1-p1
Cfelis TBC-PI-X2 CF000604.mRNA-p1
Bbigemina TBC-PI-X2 BBBOND 0209190-t26 1-p1
Bbovis TBC-PI-X2 BBOV IV005540-t26 1-p1
Bmicroti TBC-PI-X2 BMR1 02g01160-t32 1-p1
Gniphandrodes TBC-PI-X2 GNI 014070-t26 1-p1
Pberghei TBC-PI-X2 PBANKA 0933400.1-p1
Pchabaudi TBC-PI-X2 PCHAS 0910900.1-p1
Pyoelii TBC-PI-X2 PYYM 0934800.1-p1
Pknowlesi TBC-PI-X2 PKNH 0911800.1-p1
Pvivax TBC-PI-X2 PVX 091355.1-p1
Povale TBC-PI-X2 PocGH01 09020500.1-p1
Pfalciparum TBC-PI-X2 PF3D7 1114200.1-p1
Preichenowi TBC-PI-X2 PRCDC 1112700.1-p1
Pgallinaceum TBC-PI-X2 PGAL8A 00341100.1-p1
Prelictum TBC-PI-X2 PRELSG 0911300.1-p1
Hhammondi TBC-PI-X2 HHA 261200-t26 1-p1
Tgondii-ME49 TBC-PI-X2 TGME49 261200-t26 1-p1
Ncaninum TBC-PI-X2 NCLIV 026190-t26 1-p1
Sneurona TBC-PI-X2 SN3 00301370-mRNA-1-p1
Etenella TBC-PI-X2 ETH 00026585-t26 1-p1
Vbrassicaformis TBC-PI-X2 Vbra 966.t1-p1
Cvelia TBC-PI-X2 Cvel 17475.t1-p1
Smicroadriaticum TBC-PI-X2 Smic26821
Sminutum TBC-PI-X2 symbB.v1.2.026200.t1scaffold2557.1size764175
Pmarinus TBC-PI-X2 EER11537
Chominis TBC-PI-X2 ChTU502y2012 406g0790-t32 1-p1
Cparvum TBC-PI-X2 cgd6 2190-t26 1-p1
Cmuris TBC-PI-X2 CMU 007680-t26 1-p1
Athaliana TBC-PI-X2 AT3G07890.1
Creinhardtii TBC-PI-X2 Cre10.g459800.t1.2
Creinhardtii TBC-PI-X2 Cre17.g737050.t1.2
Athaliana TBC-PI-X2 AT2G39280.2
Athaliana TBC-PI-X2 AT3G55020.1
Athaliana TBC-PI-X2 AT2G37290.2
Creinhardtii TBC-PI-X2 Cre02.g077700.t1.1
Pmarinus TBC-PI-X2 EER09896
Smicroadriaticum TBC-PI-X2 Smic12901
Sminutum TBC-PI-X2 symbB.v1.2.014537.t1scaffold1066.1size1403471
Vbrassicaformis TBC-PI-X2 Vbra 15646.t1-p1
Imultifiliis TBC-K IMG5 171440
Tthermophila TBC-K TTHERM 00139660
Ptetraurelia TBC-K GSPATP00006491001
Ptetraurelia TBC-K GSPATP00009797001
Tthermophila TBC-K TTHERM 00151770
Bnatans TBC-K jgiBigna187487estExt fgenesh1 pg.C 210008
Bnatans TBC-K jgiBigna190520estExt fgenesh1 pg.C 720069
Bnatans TBC-K jgiBigna183045fgenesh1 pg.101 19
Esiliculosus TBC-K Ec-10 002230
Ptricornutum TBC-K jgiPhatr245734estExt fgenesh1 pg.C chr 80028
Cvelia TBC-K Cvel 7873.t1-p1
Cvelia TBC-K Cvel 3781.t1-p1
Gniphandrodes TBC-K GNI 027920-t26 1-p1
Ccayetanensis TBC-K cyc 04882-t31 1-p1
Sneurona TBC-K SN3 01900435-mRNA-1-p1
Ncaninum TBC-K NCLIV 051140-t26 1-p1
Hhammondi TBC-K HHA 237280-t26 1-p1
Tgondii-ME49 TBC-K TGME49 237280-t26 1-p1
Pberghei TBC-K PBANKA 1331200.1-p1
Pyoelii TBC-K PYYM 1332900.1-p1
Pchabaudi TBC-K PCHAS 1335800.1-p1
Pgallinaceum TBC-K PGAL8A 00227300.1-p1
Prelictum TBC-K PRELSG 1212900.1-p1
Povale TBC-K PocGH01 12046900.1-p1
Pfalciparum TBC-K PF3D7 1467900.1-p1
Preichenowi TBC-K PRCDC 1467100.1-p1
Pknowlesi TBC-K PKNH 1213300.1-p1
Pvivax TBC-K PVX 117065.1-p1
Bmicroti TBC-K BMR1 02g03000-t32 1-p1
Bbigemina TBC-K BBBOND 0201210-t26 1-p1
Bbovis TBC-K BBOV II001320-t26 1-p1
Cfelis TBC-K CF000974.mRNA-p1
Tannulata TBC-K TA08515-t26 1-p1
Tparva TBC-K TP04 0454-t26 1-p1
Torientalis TBC-K TOT 040000461-t26 1-p1
0.723657

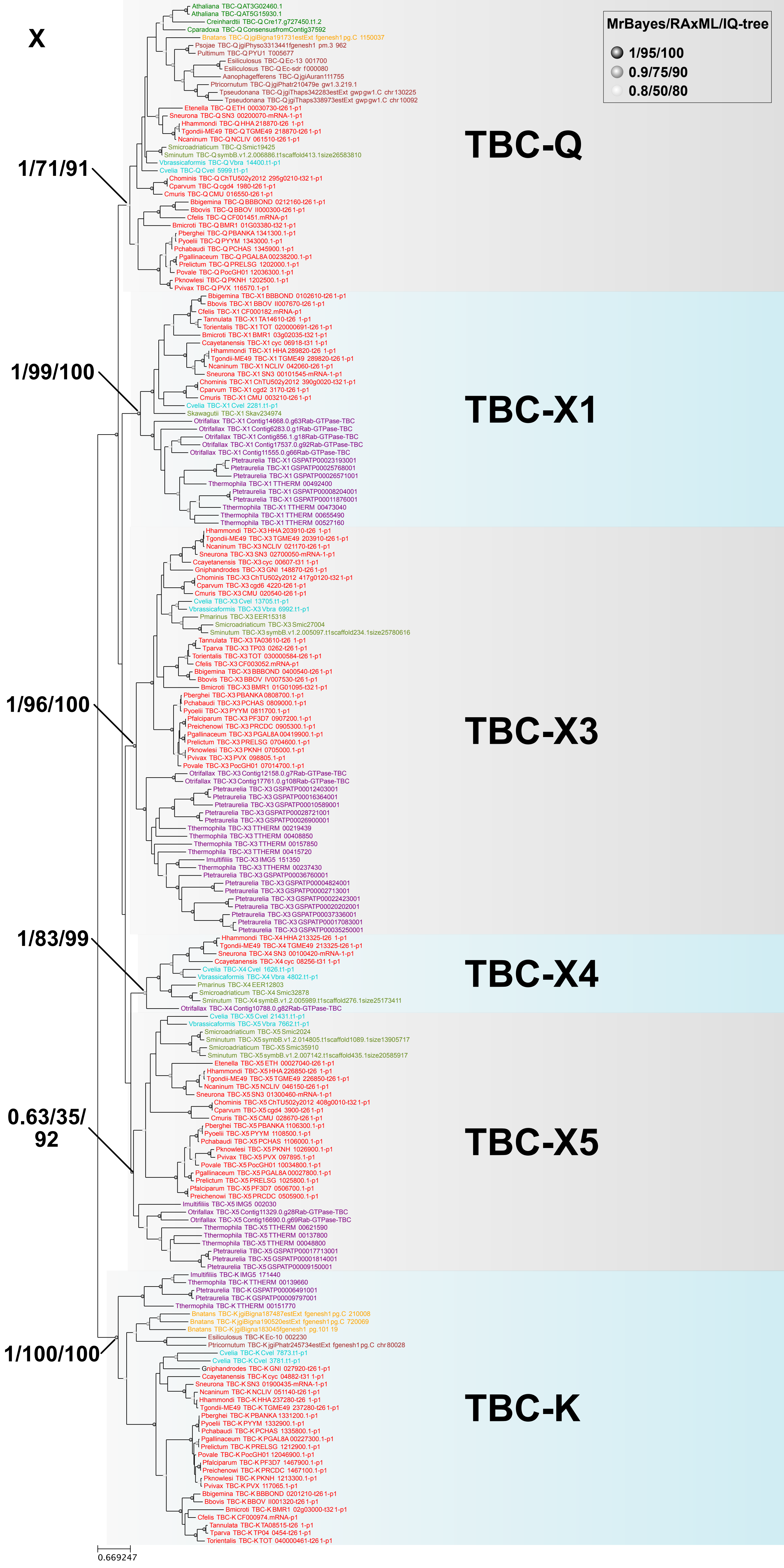
X
MrBayes/RAxML/IQ-tree
1/95/100
0.9/75/90
0.8/50/80
TBC-Q
TBC-X1
TBC-X3
TBC-X4
TBC-X5
TBC-K
1/71/91
1/99/100
1/96/100
1/83/99
0.63/35/
92
1/100/100
0.669247
Athaliana TBC-QAT3G02460.1
Athaliana TBC-QAT5G15930.1
Creinhardtii TBC-Q Cre17.g727450.t1.2
Cparadoxa TBC-Q ConsensusfromContig37592
Bnatans TBC-Q jgiBigna191731estExt fgenesh1pg.C 1150037
Psojae TBC-Q jgiPhyso3313441fgenesh1 pm.3 962
Pultimum TBC-Q PYU1 T005677
Esiliculosus TBC-Q Ec-13 001700
Esiliculosus TBC-Q Ec-sdr f000080
Aanophagefferens TBC-Q jgiAuran111755
Ptricornutum TBC-Q jgiPhatr210479e gw1.3.219.1
Tpseudonana TBC-Q jgiThaps342283estExt gwpgw1.C chr130225
Tpseudonana TBC-Q jgiThaps338973estExt gwpgw1.C chr10092
Etenella TBC-Q ETH 00030730-t26 1-p1
Sneurona TBC-Q SN3 00200070-mRNA-1-p1
Hhammondi TBC-Q HHA 218870-t26 1-p1
Tgondii-ME49 TBC-Q TGME49 218870-t26 1-p1
Ncaninum TBC-Q NCLIV 061510-t26 1-p1
Smicroadriaticum TBC-Q Smic19425
Sminutum TBC-Q symbB.v1.2.006886.t1scaffold413.1size26583810
Vbrassicaformis TBC-Q Vbra 14400.t1-p1
Cvelia TBC-Q Cvel 5999.t1-p1
Chominis TBC-Q ChTU502y2012 295g0210-t32 1-p1
Cparvum TBC-Q cgd4 1980-t26 1-p1
Cmuris TBC-Q CMU 016550-t26 1-p1
Bbigemina TBC-Q BBBOND 0212160-t26 1-p1
Bbovis TBC-Q BBOV II000300-t26 1-p1
Cfelis TBC-Q CF001451.mRNA-p1
Bmicroti TBC-Q BMR1 01G03380-t32 1-p1
Pberghei TBC-Q PBANKA 1341300.1-p1
Pyoelii TBC-Q PYYM 1343000.1-p1
Pchabaudi TBC-Q PCHAS 1345900.1-p1
Pgallinaceum TBC-Q PGAL8A 00238200.1-p1
Prelictum TBC-Q PRELSG 1202000.1-p1
Povale TBC-Q PocGH01 12036300.1-p1
Pknowlesi TBC-Q PKNH 1202500.1-p1
Pvivax TBC-Q PVX 116570.1-p1
Bbigemina TBC-X1 BBBOND 0102610-t26 1-p1
Bbovis TBC-X1 BBOV II007670-t26 1-p1
Cfelis TBC-X1 CF000182.mRNA-p1
Tannulata TBC-X1 TA14610-t26 1-p1
Torientalis TBC-X1 TOT 020000691-t26 1-p1
Bmicroti TBC-X1 BMR1 03g02035-t32 1-p1
Ccayetanensis TBC-X1 cyc 06918-t31 1-p1
Hhammondi TBC-X1 HHA 289820-t26 1-p1
Tgondii-ME49 TBC-X1 TGME49 289820-t26 1-p1
Ncaninum TBC-X1 NCLIV 042060-t26 1-p1
Sneurona TBC-X1 SN3 00101545-mRNA-1-p1
Chominis TBC-X1 ChTU502y2012 390g0020-t32 1-p1
Cparvum TBC-X1 cgd2 3170-t26 1-p1
Cmuris TBC-X1 CMU 003210-t26 1-p1
Cvelia TBC-X1 Cvel 2281.t1-p1
Skawagutii TBC-X1 Skav234974
Otrifallax TBC-X1 Contig14668.0.g63Rab-GTPase-TBC
Otrifallax TBC-X1 Contig6283.0.g1Rab-GTPase-TBC
Otrifallax TBC-X1 Contig856.1.g18Rab-GTPase-TBC
Otrifallax TBC-X1 Contig17537.0.g92Rab-GTPase-TBC
Otrifallax TBC-X1 Contig11555.0.g66Rab-GTPase-TBC
Ptetraurelia TBC-X1 GSPATP00023193001
Ptetraurelia TBC-X1 GSPATP00025768001
Ptetraurelia TBC-X1 GSPATP00026571001
Tthermophila TBC-X1 TTHERM 00492400
Ptetraurelia TBC-X1 GSPATP00008204001
Ptetraurelia TBC-X1 GSPATP00011876001
Tthermophila TBC-X1 TTHERM 00473040
Tthermophila TBC-X1 TTHERM 00655490
Tthermophila TBC-X1 TTHERM 00527160
Hhammondi TBC-X3 HHA 203910-t26 1-p1
Tgondii-ME49 TBC-X3 TGME49 203910-t26 1-p1
Ncaninum TBC-X3 NCLIV 021170-t26 1-p1
Sneurona TBC-X3 SN3 02700050-mRNA-1-p1
Ccayetanensis TBC-X3 cyc 00607-t31 1-p1
Gniphandrodes TBC-X3 GNI 148870-t26 1-p1
Chominis TBC-X3 ChTU502y2012 417g0120-t32 1-p1
Cparvum TBC-X3 cgd6 4220-t26 1-p1
Cmuris TBC-X3 CMU 020540-t26 1-p1
Cvelia TBC-X3 Cvel 13705.t1-p1
Vbrassicaformis TBC-X3 Vbra 6992.t1-p1
Pmarinus TBC-X3 EER15318
Smicroadriaticum TBC-X3 Smic27004
Sminutum TBC-X3 symbB.v1.2.005097.t1scaffold234.1size25780616
Tannulata TBC-X3 TA03610-t26 1-p1
Tparva TBC-X3 TP03 0262-t26 1-p1
Torientalis TBC-X3 TOT 030000584-t26 1-p1
Cfelis TBC-X3 CF003052.mRNA-p1
Bbigemina TBC-X3 BBBOND 0400540-t26 1-p1
Bbovis TBC-X3 BBOV IV007530-t26 1-p1
Bmicroti TBC-X3 BMR1 01G01095-t32 1-p1
Pberghei TBC-X3 PBANKA 0808700.1-p1
Pchabaudi TBC-X3 PCHAS 0809000.1-p1
Pyoelii TBC-X3 PYYM 0811700.1-p1
Pfalciparum TBC-X3 PF3D7 0907200.1-p1
Preichenowi TBC-X3 PRCDC 0905300.1-p1
Pgallinaceum TBC-X3 PGAL8A 00419900.1-p1
Prelictum TBC-X3 PRELSG 0704600.1-p1
Pknowlesi TBC-X3 PKNH 0705000.1-p1
Pvivax TBC-X3 PVX 098805.1-p1
Povale TBC-X3 PocGH01 07014700.1-p1
Otrifallax TBC-X3 Contig12158.0.g7Rab-GTPase-TBC
Otrifallax TBC-X3 Contig17761.0.g108Rab-GTPase-TBC
Ptetraurelia TBC-X3 GSPATP00012403001
Ptetraurelia TBC-X3 GSPATP00016364001
Ptetraurelia TBC-X3 GSPATP00010589001
Ptetraurelia TBC-X3 GSPATP00028721001
Ptetraurelia TBC-X3 GSPATP00026900001
Tthermophila TBC-X3 TTHERM 00219439
Tthermophila TBC-X3 TTHERM 00408850
Tthermophila TBC-X3 TTHERM 00157850
Tthermophila TBC-X3 TTHERM 00415720
Imultifiliis TBC-X3 IMG5 151350
Tthermophila TBC-X3 TTHERM 00237430
Ptetraurelia TBC-X3 GSPATP00036760001
Ptetraurelia TBC-X3 GSPATP00004824001
Ptetraurelia TBC-X3 GSPATP00002713001
Ptetraurelia TBC-X3 GSPATP00022423001
Ptetraurelia TBC-X3 GSPATP00020202001
Ptetraurelia TBC-X3 GSPATP00037336001
Ptetraurelia TBC-X3 GSPATP00017083001
Ptetraurelia TBC-X3 GSPATP00035250001
Hhammondi TBC-X4 HHA 213325-t26 1-p1
Tgondii-ME49 TBC-X4 TGME49 213325-t26 1-p1
Sneurona TBC-X4 SN3 00100420-mRNA-1-p1
Ccayetanensis TBC-X4 cyc 08256-t31 1-p1
Cvelia TBC-X4 Cvel 1626.t1-p1
Vbrassicaformis TBC-X4 Vbra 4802.t1-p1
Pmarinus TBC-X4 EER12803
Smicroadriaticum TBC-X4 Smic32878
Sminutum TBC-X4 symbB.v1.2.005989.t1scaffold276.1size25173411
Otrifallax TBC-X4 Contig10788.0.g82Rab-GTPase-TBC
Cvelia TBC-X5 Cvel 21431.t1-p1
Vbrassicaformis TBC-X5 Vbra 7662.t1-p1
Smicroadriaticum TBC-X5 Smic2024
Sminutum TBC-X5 symB.v1.2.014805.t1scaffold1089.1size13905717
Smicroadriaticum TBC-X5 Smic35910
Sminutum TBC-X5 symbB.v1.2.007142.t1scaffold435.1size20585917
Etenella TBC-X5 ETH 00027040-t26 1-p1
Hhammondi TBC-X5 HHA 226850-t26 1-p1
Tgondii-ME49 TBC-X5 TGME49 226850-t26 1-p1
Ncaninum TBC-X5 NCLIV 046150-t26 1-p1
Sneurona TBC-X5 SN3 01300460-mRNA-1-p1
Chominis TBC-X5 ChTU502y2012 408g0010-t32 1-p1
Cparvum TBC-X5 cgd4 3900-t26 1-p1
Cmuris TBC-X5 CMU 028670-t26 1-p1
Pberghei TBC-X5 PBANKA 1106300.1-p1
Pyoelii TBC-X5 PYYM 1108500.1-p1
Pchabaudi TBC-X5 PCHAS 1106000.1-p1
Pknowlesi TBC-X5 PKNH 1026900.1-p1
Pvivax TBC-X5 PVX 097895.1-p1
Povale TBC-X5 PocGH01 10034800.1-p1
Pgallinaceum TBC-X5 PGAL8A 00027800.1-p1
Prelictum TBC-X5 PRELSG 1025800.1-p1
Pfalciparum TBC-X5 PF3D7 0506700.1-p1
Preichenowi TBC-X5 PRCDC 0505900.1-p1
Imultifiliis TBC-X5 IMG5 002030
Otrifallax TBC-X5 Contig11329.0.g28Rab-GTPase-TBC
Otrifallax TBC-X5 Contig16690.0.g69Rab-GTPase-TBC
Tthermophila TBC-X5 TTHERM 00621590
Tthermophila TBC-X5 TTHERM 00137800
Tthermophila TBC-X5 TTHERM 00048800
Ptetraurelia TBC-X5 GSPATP00017713001
Ptetraurelia TBC-X5 GSPATP00001814001
Ptetraurelia TBC-X5 GSPATP00009150001
Imultifiliis TBC-K IMG5 171440
Tthermophila TBC-K TTHERM 00139660
Ptetraurelia TBC-K GSPATP00006491001
Ptetraurelia TBC-K GSPATP00009797001
Tthermophila TBC-K TTHERM 00151770
Bnatans TBC-K jgiBigna187487estExt fgenesh1pg.C 210008
Bnatans TBC-K jgiBigna190520estExt fgenesh1pg.C 720069
Bnatans TBC-K jgiBigna183045fgenesh1 pg.101 19
Esiliculosus TBC-K Ec-10 002230
Ptricornutum TBC-K jgiPhatr245734estExt fgenesh1pg.C chr80028
Cvelia TBC-K Cvel 7873.t1-p1
Cvelia TBC-K Cvel 3781.t1-p1
Gniphandrodes TBC-K GNI 027920-t26 1-p1
Ccayetanensis TBC-K cyc 04882-t31 1-p1
Sneurona TBC-K SN3 01900435-mRNA-1-p1
Ncaninum TBC-K NCLIV 051140-t26 1-p1
Hhammondi TBC-K HHA 237280-t26 1-p1
Tgondii-ME49 TBC-K TGME49 237280-t26 1-p1
Pberghei TBC-K PBANKA 1331200.1-p1
Pyoelii TBC-K PYYM 1332900.1-p1
Pchabaudi TBC-K PCHAS 1335800.1-p1
Pgallinaceum TBC-K PGAL8A 00227300.1-p1
Prelictum TBC-K PRELSG 1212900.1-p1
Povale TBC-K PocGH01 12046900.1-p1
Pfalciparum TBC-K PF3D7 1467900.1-p1
Preichenowi TBC-K PRCDC 1467100.1-p1
Pknowlesi TBC-K PKNH 1213300.1-p1
Pvivax TBC-K PVX 117065.1-p1
Bbigemina TBC-K BBBOND 0201210-t26 1-p1
Bbovis TBC-K BBOV II001320-t26 1-p1
Bmicroti TBC-K BMR1 02g03000-t32 1-p1
Cfelis TBC-K CF000974.mRNA-p1
Tannulata TBC-K TA08515-t26 1-p1
Tparva TBC-K TP04 0454-t26 1-p1
Torientalis TBC-K TOT 040000461-t26 1-p1
