## Supplementary Figure 3 for "Evolution of lineage-specific trafficking proteins and a novel post-Golgi trafficking pathway in Apicomplexa"

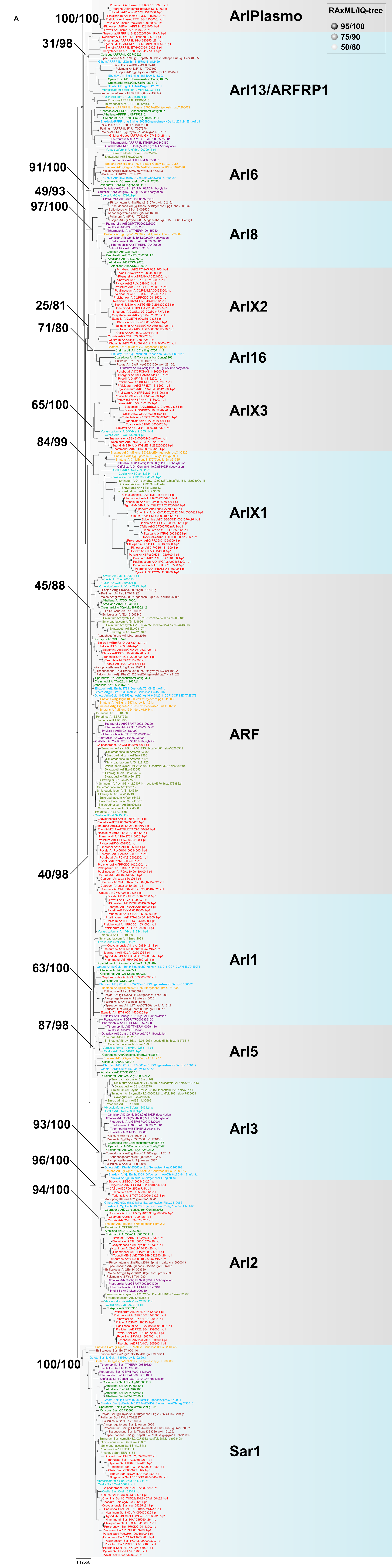
A
RAxML/IQ-tree
95/100
75/90
50/80
100/100
31/98
ArlPlasmo
Arl13/ARFRP1
91/100
49/93
97/100
Arl6
Arl8
25/81
71/80
ArlX2
Arl16
65/100
84/99
ArlX3
ArlX1
45/88
ARF
40/98
63/100
Arl1
87/98
Arl5
93/100
96/100
94/100
Arl3
Arl2
100/100
Sar1
1.12666

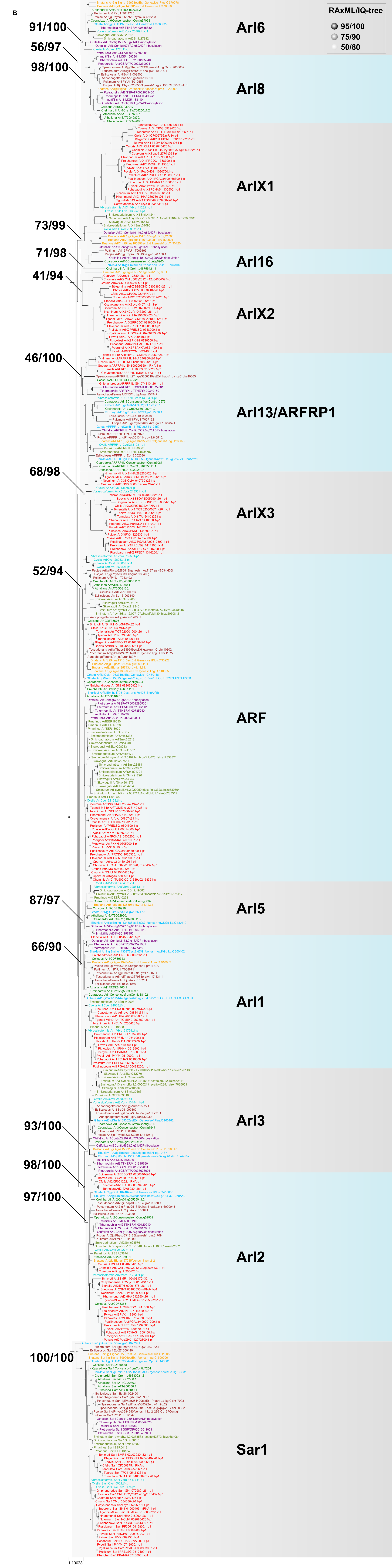

B
RAxML/IQ-tree
95/100
75/90
50/80
91/100
56/97
98/100
73/99
71/98
41/94
46/100
68/98
52/94
87/97
66/90
93/100
98/100
97/100
100/100
Arl6
Arl8
ArlX1
Arl16
ArlX2
Arl13/ARFRP1
ArlX3
ARF
Arl5
Arl1
Arl3
Arl2
Sar1

c

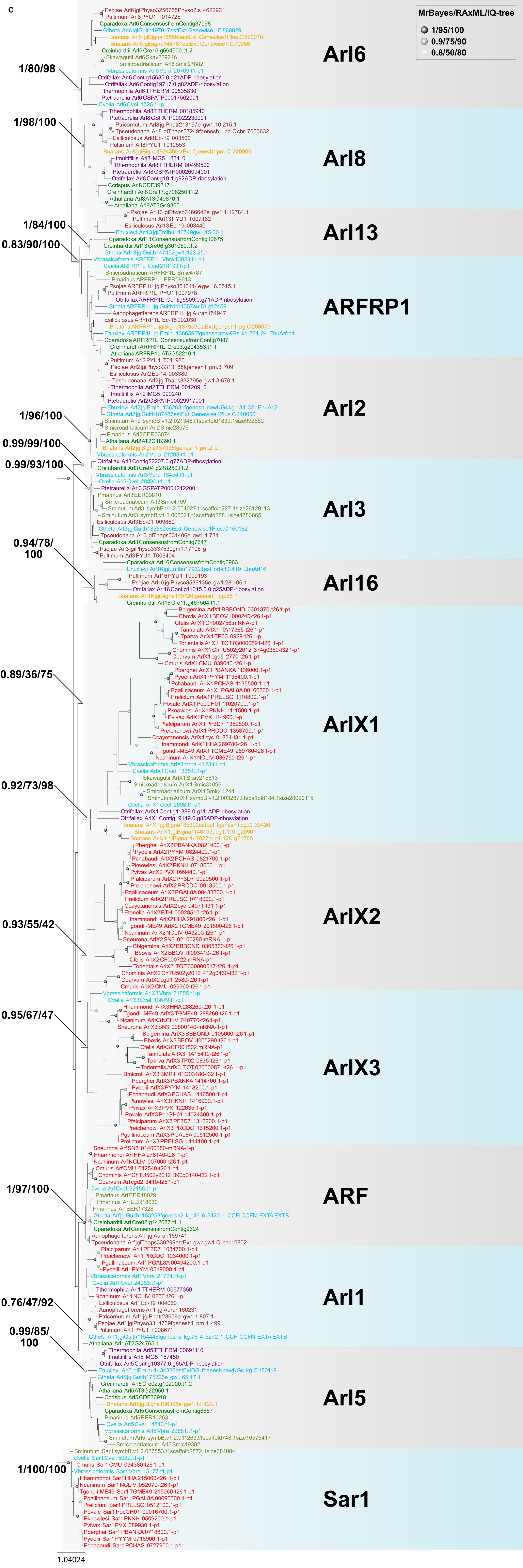
MrBayes/RAxML/IQ-tree
1/95/100
0.9/75/90
0.8/50/80
Arl6
1/80/98
Arl8
1/98/100
Arl13
1/84/100
0.83/90/100
ARFRP1
Arl2
1/96/100
0.99/99/100
0.99/93/100
Arl3
0.94/78/
100
Arl16
0.89/36/75
ArlX1
0.92/73/98
ArlX2
0.93/55/42
0.95/67/47
ArlX3
ARF
1/97/100
Arl1
0.76/47/92
0.99/85/
100
Arl5
Sar1
1/100/100
1.04024

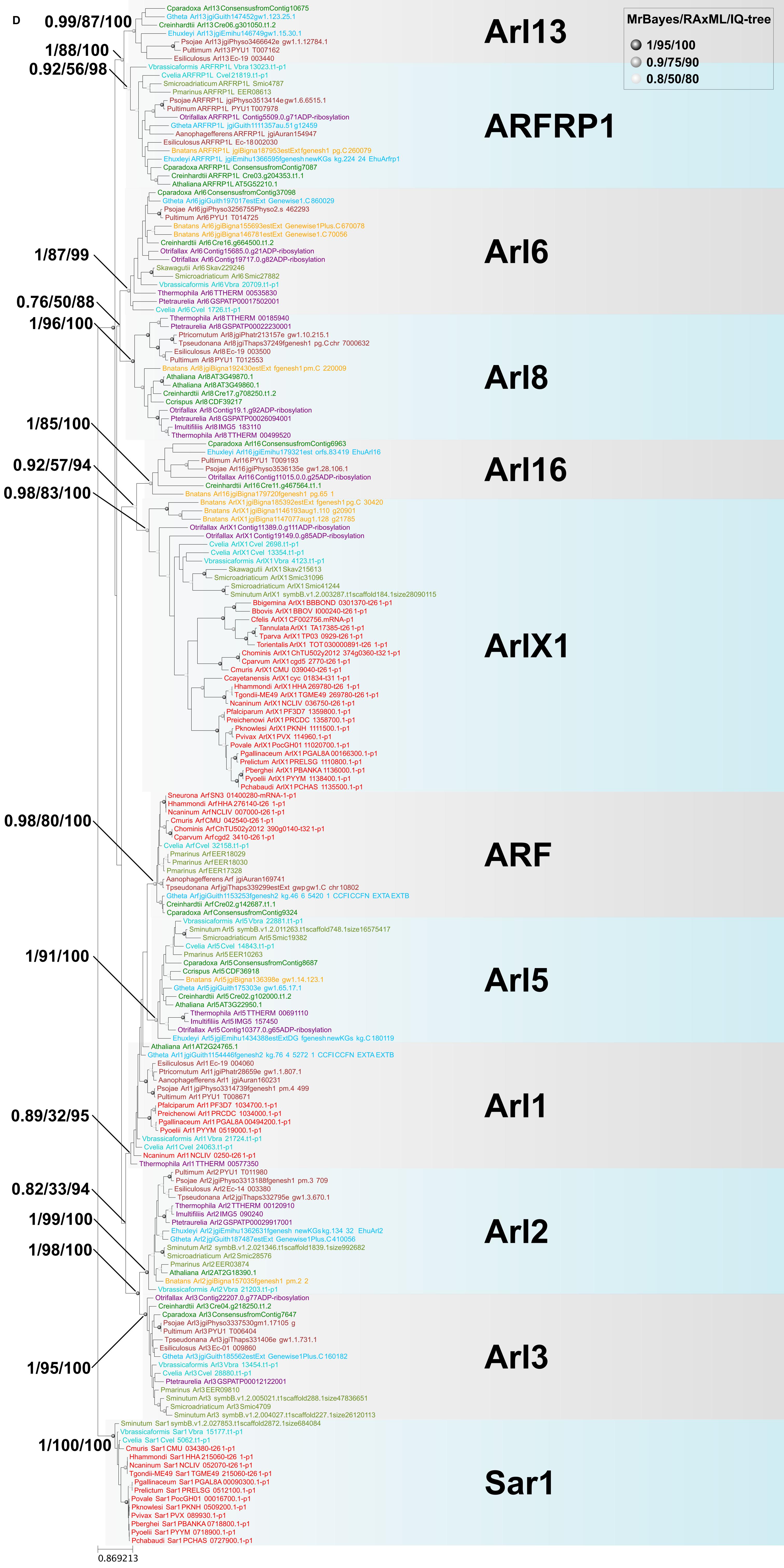
D
MrBayes/RAxML/IQ-tree
1/95/100
0.9/75/90
0.8/50/80
0.99/87/100
1/88/100
0.92/56/98
Arl13
ARFRP1
1/87/99
Arl6
0.76/50/88
1/96/100
Arl8
1/85/100
Arl16
0.92/57/94
0.98/83/100
ArlX1
0.98/80/100
ARF
1/91/100
Arl5
Arl1
0.89/32/95
0.82/33/94
1/99/100
1/98/100
Arl2
Arl3
1/95/100
1/100/100
Sar1
0.869213

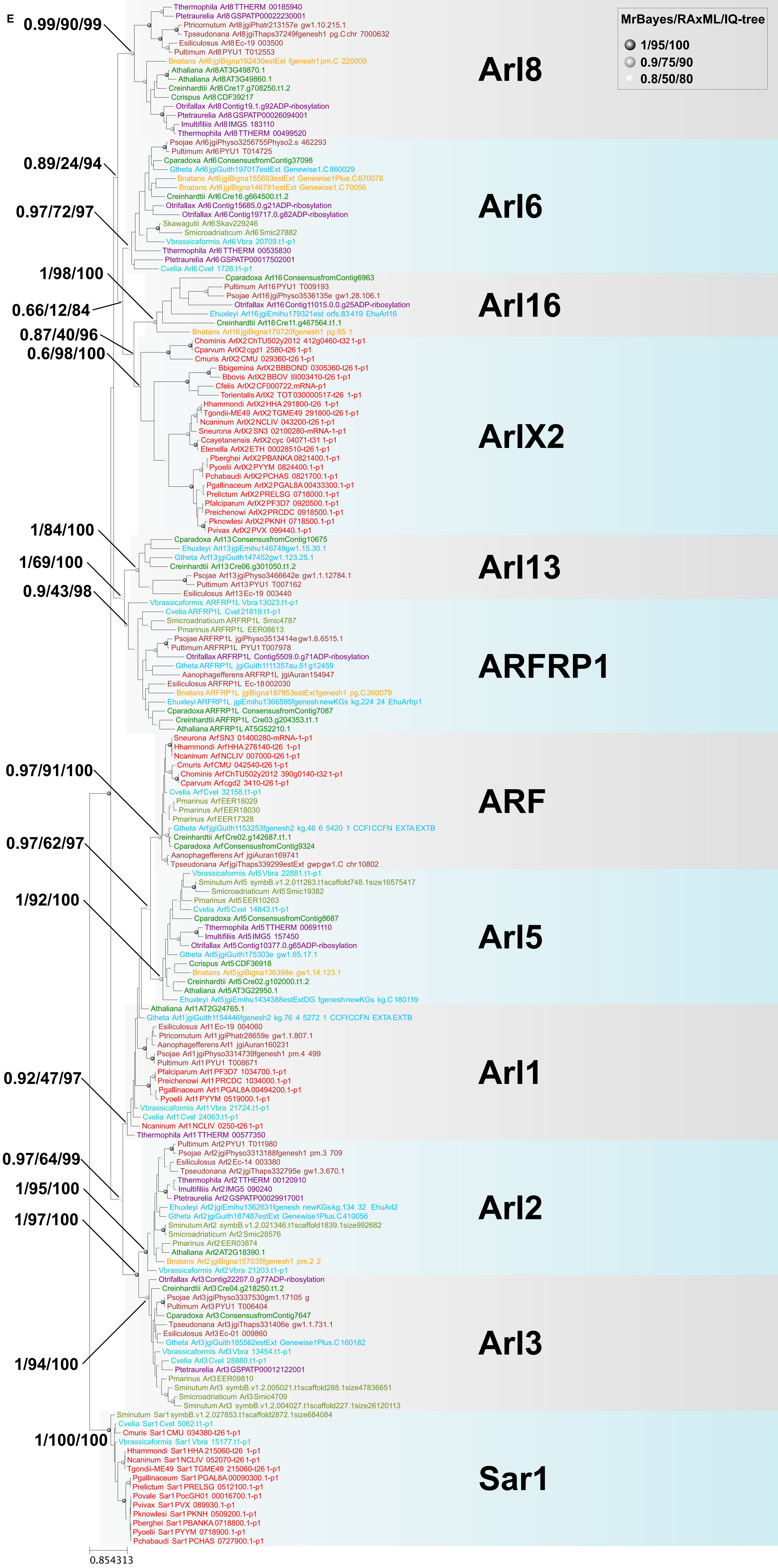
E
MrBayes/RAxML/IQ-tree
1/95/100
0.9/75/90
0.8/50/80
0.99/90/99
0.89/24/94
0.97/72/97
1/98/100
0.66/12/84
0.87/40/96
0.6/98/100
1/84/100
1/69/100
0.9/43/98
0.97/91/100
0.97/62/97
1/92/100
0.92/47/97
0.97/64/99
1/95/100
1/97/100
1/94/100
1/100/100
Arl8
Tthermophila Arl8 TTHERM 00185940
Ptetraurelia Arl8 GSPATP00022230001
Ptricornutum Arl8 jgiPhatr213157e gw1.10.215.1
Tpseudonana Arl8 jgiThaps37249fgenesh1 pg.C chr 7000632
Esiliculosus Arl8 Ec-19 003500
Pultimum Arl8 PYU1 T012553
Bnatans Arl8 jgiBigna192430estExt fgenesh1 pm.C 220009
Athaliana Arl8 AT3G49870.1
Athaliana Arl8 AT3G49860.1
Creinhardtii Arl8 Cre17.g708250.t1.2
Ccrispus Arl8 CDF39217
Otrifallax Arl8 Contig19.1.g92ADP-ribosylation
Ptetraurelia Arl8 GSPATP00026094001
Imultifiliis Arl8 IMG5 183110
Tthermophila Arl8 TTHERM 00499520
Arl6
Psojae Arl6 jgiPhyso3256755Physo2.s 462293
Pultimum Arl6 PYU1 T014725
Cparadoxa Arl6 ConsensusfromContig37098
Gtheta Arl6 jgiGuith197017estExt Genewise1.C 860029
Bnatans Arl6 jgiBigna155693estExt Genewise1Plus.C 670078
Bnatans Arl6 jgiBigna146781estExt Genewise1.C 70056
Creinhardtii Arl6 Cre16.g664500.t1.2
Otrifallax Arl6 Contig15685.0.g21ADP-ribosylation
Otrifallax Arl6 Contig19717.0.g82ADP-ribosylation
Skawagutii Arl6 Skav229246
Smicroadriaticum Arl6 Smic27882
Vbrassicaformis Arl6 Vbra 20709.t1-p1
Tthermophila Arl6 TTHERM 00535830
Ptetraurelia Arl6 GSPATP00017502001
Cvelia Arl6 Cvel 1726.t1-p1
Arl16
Cparadoxa Arl16 ConsensusfromContig6963
Pultimum Arl16 PYU1 T009193
Psojae Arl16 jgiPhyso3536135e gw1.28.106.1
Otrifallax Arl16 Contig11015.0.0.g25ADP-ribosylation
Ehuxleyi Arl16 jgiEmihu179321est orfs.83419 EhuArl16
Creinhardtii Arl16 Cre11.g467564.t1.1
Bnatans Arl16 jgiBigna179720fgenesh1 pg.65 1
ArlX2
Chominis ArlX2 ChTU502y2012 412g0460-t32 1-p1
Cparvum ArlX2 cgd1 2580-t26 1-p1
Cmuris ArlX2 CMU 029360-t26 1-p1
Bbigemina ArlX2 BBBOND 0305360-t26 1-p1
Bbovis ArlX2 BBOV III003410-t26 1-p1
Cfelis ArlX2 CF000722.mRNA-p1
Torientalis ArlX2 TOT 030000517-t26 1-p1
Hhammondi ArlX2 HHA 291800-t26 1-p1
Tgondii-ME49 ArlX2 TGME49 291800-t26 1-p1
Ncaninum ArlX2 NCLIV 043200-t26 1-p1
Sneurona ArlX2 SN3 02100280-mRNA-1-p1
Ccayetanensis ArlX2 cyc 04071-t31 1-p1
Etenella ArlX2 ETH 00028510-t26 1-p1
Pberghei ArlX2 PBANKA 0821400.1-p1
Pyoelii ArlX2 PYYM 0824400.1-p1
Pchabaudi ArlX2 PCHAS 0821700.1-p1
Pgallinaceum ArlX2 PGAL8A 00433300.1-p1
Prelictum ArlX2 PRELSG 0718000.1-p1
Pfalciparum ArlX2 PF3D7 0920500.1-p1
Preichenowi ArlX2 PRCDC 0918500.1-p1
Pknowlesi ArlX2 PKNH 0718500.1-p1
Pvivax ArlX2 PVX 099440.1-p1
Arl13
Cparadoxa Arl13 ConsensusfromContig10675
Ehuxleyi Arl13 jgiEmihu146749gw1.15.30.1
Gtheta Arl13 jgiGuith147452gw1.123.25.1
Creinhardtii Arl13 Cre06.g301050.t1.2
Psojae Arl13 jgiPhyso3466642e gw1.1.12784.1
Pultimum Arl13 PYU1 T007162
Esiliculosus Arl13 Ec-19 003440
ARFRP1
Vbrassicaformis ARFRP1L Vbra 13023.t1-p1
Cvelia ARFRP1L Cvel 21819.t1-p1
Smicroadriaticum ARFRP1L Smic4787
Pmarinus ARFRP1L EER08613
Psojae ARFRP1L jgiPhyso3513414e gw1.6.6515.1
Pultimum ARFRP1L PYU1 T007978
Otrifallax ARFRP1L Contig5509.0.g71ADP-ribosylation
Gtheta ARFRP1L jgiGuith1111357au.51 g12459
Aanophagefferens ARFRP1L jgiAuran154947
Esiliculosus ARFRP1L Ec-18 002030
Bnatans ARFRP1L jgiBigna187953estExt fgenesh1 pg.C 260079
Ehuxleyi ARFRP1L jgiEmihu1366595fgenesh newKGs kg.224 24 EhuArfrp1
Cparadoxa ARFRP1L ConsensusfromContig7087
Creinhardtii ARFRP1L Cre03.g204353.t1.1
Athaliana ARFRP1L AT5G52210.1
ARF
Sneurona Arf SN3 01400280-mRNA-1-p1
Hhammondi Arf HHA 276140-t26 1-p1
Ncaninum Arf NCLIV 007000-t26 1-p1
Cmuris Arf CMU 042540-t26 1-p1
Chominis Arf ChTU502y2012 390g0140-t32 1-p1
Cparvum Arf cgd2 3410-t26 1-p1
Cvelia Arf Cvel 32158.t1-p1
Pmarinus Arf EER18029
Pmarinus Arf EER18030
Pmarinus Arf EER17328
Gtheta Arf jgiGuith1153253fgenesh2 kg.46 6 5420 1 CCFI CCFN EXTA EXTB
Creinhardtii Arf Cre02.g142687.t1.1
Cparadoxa Arf ConsensusfromContig9324
Aanophagefferens Arf jgiAuran169741
Tpseudonana Arf jgiThaps339299estExt gwp gw1.C chr 10802
Arl5
Vbrassicaformis Arl5 Vbra 22881.t1-p1
Sminutum Arl5 symbB.v1.2.011263.t1scaffold748.1size16575417
Smicroadriaticum Arl5 Smic19382
Pmarinus Arl5 EER10263
Cvelia Arl5 Cvel 14843.t1-p1
Cparadoxa Arl5 ConsensusfromContig8687
Tthermophila Arl5 TTHERM 00691110
Imultifiliis Arl5 IMG5 157450
Otrifallax Arl5 Contig10377.0.g65ADP-ribosylation
Gtheta Arl5 jgiGuith175303e gw1.65.17.1
Ccrispus Arl5 CDF36918
Bnatans Arl5 jgiBigna136398e gw1.14.123.1
Creinhardtii Arl5 Cre02.g102000.t1.2
Athaliana Arl5 AT3G22950.1
Ehuxleyi Arl5 jgiEmihu1434388estExtDG fgenesh newKGs kg.C 180119
Arl1
Athaliana Arl1 AT2G24765.1
Gtheta Arl1 jgiGuith1154446fgenesh2 kg.76 4 5272 1 CCFI CCFN EXTA EXTB
Esiliculosus Arl1 Ec-19 004060
Ptricornutum Arl1 jgiPhatr28659e gw1.1.807.1
Aanophagefferens Arl1 jgiAuran160231
Psojae Arl1 jgiPhyso3314739fgenesh1 pm.4 499
Pultimum Arl1 PYU1 T008671
Pfalciparum Arl1 PF3D7 1034700.1-p1
Preichenowi Arl1 PRCDC 1034000.1-p1
Pgallinaceum Arl1 PGAL8A 00494200.1-p1
Pyoelii Arl1 PYYM 0519000.1-p1
Vbrassicaformis Arl1 Vbra 21724.t1-p1
Cvelia Arl1 Cvel 24063.t1-p1
Ncaninum Arl1 NCLIV 0250-t26 1-p1
Tthermophila Arl1 TTHERM 00577350
Arl2
Pultimum Arl2 PYU1 T011980
Psojae Arl2 jgiPhyso3313188fgenesh1 pm.3 709
Esiliculosus Arl2 Ec-14 003380
Tpseudonana Arl2 jgiThaps332795e gw1.3.670.1
Tthermophila Arl2 TTHERM 00120910
Imultifiliis Arl2 IMG5 090240
Ptetraurelia Arl2 GSPATP00029917001
Ehuxleyi Arl2 jgiEmihu1362631fgenesh newKGs kg.134 32 EhuArl2
Gtheta Arl2 jgiGuith187487estExt Genewise1Plus.C 410056
Sminutum Arl2 symbB.v1.2.021346.t1scaffold1839.1size992682
Smicroadriaticum Arl2 Smic28576
Pmarinus Arl2 EER03874
Athaliana Arl2 AT2G18390.1
Bnatans Arl2 jgiBigna157035fgenesh1 pm.2 2
Vbrassicaformis Arl2 Vbra 21203.t1-p1
Arl3
Otrifallax Arl3 Contig22207.0.g77ADP-ribosylation
Creinhardtii Arl3 Cre04.g218250.t1.2
Psojae Arl3 jgiPhyso3337530gm1.17105 g
Pultimum Arl3 PYU1 T006404
Cparadoxa Arl3 ConsensusfromContig7647
Tpseudonana Arl3 jgiThaps331406e gw1.1.731.1
Esiliculosus Arl3 Ec-01 009860
Gtheta Arl3 jgiGuith185562estExt Genewise1Plus.C 160182
Vbrassicaformis Arl3 Vbra 13454.t1-p1
Cvelia Arl3 Cvel 28880.t1-p1
Ptetraurelia Arl3 GSPATP00012122001
Pmarinus Arl3 EER09810
Sminutum Arl3 symbB.v1.2.005021.t1scaffold288.1size47836651
Smicroadriaticum Arl3 Smic4709
Sminutum Arl3 symbB.v1.2.004027.t1scaffold227.1size26120113
Sar1
Sminutum Sar1 symbB.v1.2.027853.t1scaffold2872.1size684084
Cvelia Sar1 Cvel 5062.t1-p1
Cmuris Sar1 CMU 034380-t26 1-p1
Vbrassicaformis Sar1 Vbra 15177.t1-p1
Hhammondi Sar1 HHA 215060-t26 1-p1
Ncaninum Sar1 NCLIV 052070-t26 1-p1
Tgondii-ME49 Sar1 TGME49 215060-t26 1-p1
Pgallinaceum Sar1 PGAL8A 00090300.1-p1
Prelictum Sar1 PRELSG 0512100.1-p1
Povale Sar1 PocGH01 00016700.1-p1
Pvivax Sar1 PVX 089930.1-p1
Pknowlesi Sar1 PKNH 0509200.1-p1
Pberghei Sar1 PBANKA 0718800.1-p1
Pyoelii Sar1 PYYM 0718900.1-p1
Pchabaudi Sar1 PCHAS 0727900.1-p1
0.854313

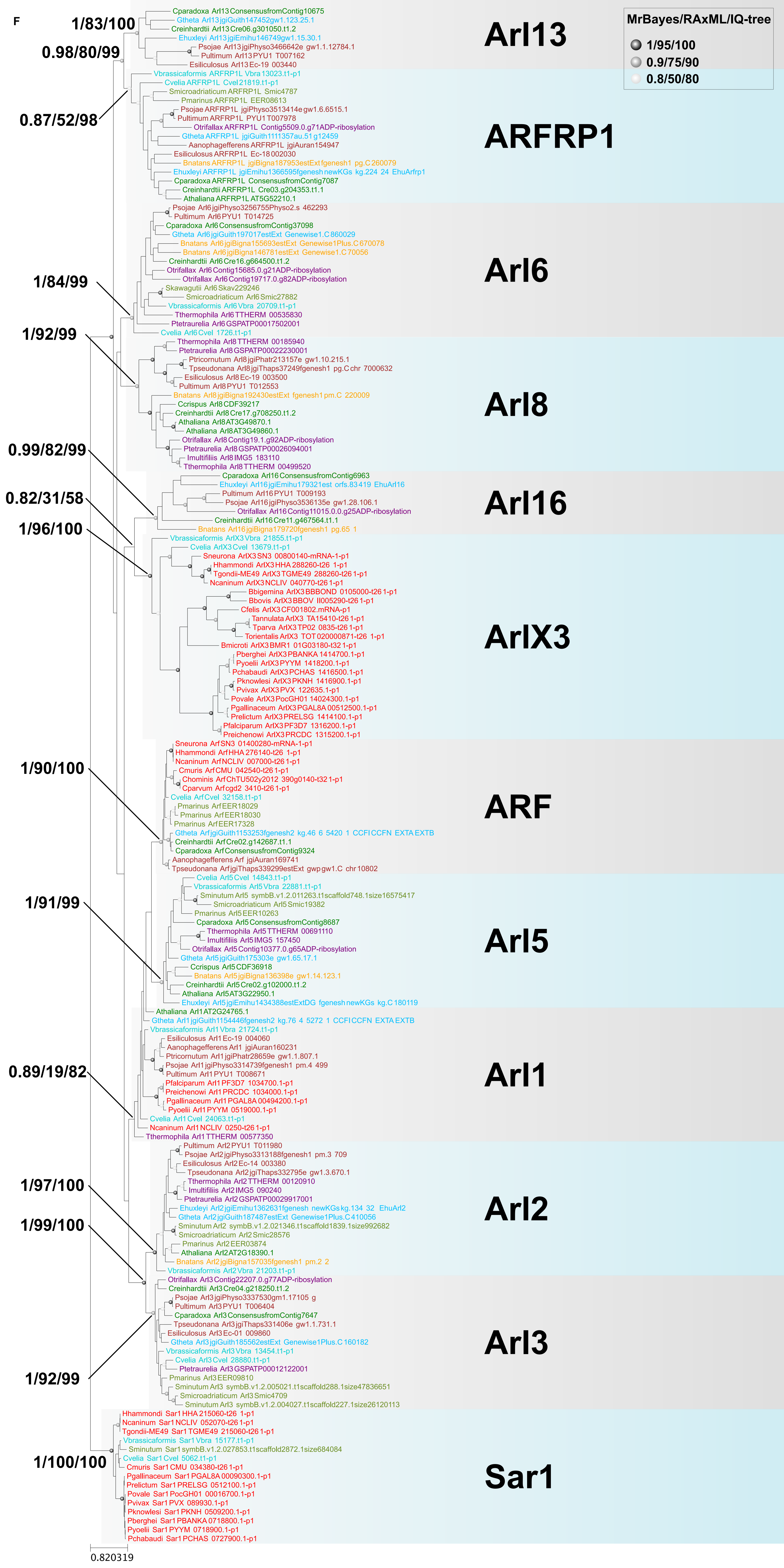
F
MrBayes/RAxML/IQ-tree
1/95/100
0.9/75/90
0.8/50/80
1/83/100
0.98/80/99
0.87/52/98
1/84/99
1/92/99
0.99/82/99
0.82/31/58
1/96/100
1/90/100
1/91/99
0.89/19/82
1/97/100
1/99/100
1/92/99
1/100/100
Arl13
Cparadoxa Arl13 ConsensusfromContig10675
Gtheta Arl13 jgiGuith147452gw1.123.25.1
Creinhardtii Arl13 Cre06.g301050.t1.2
Ehuxleyi Arl13 jgiEmihu146749gw1.15.30.1
Psojae Arl13 jgiPhyso3466642e gw1.1.12784.1
Pultimum Arl13 PYU1 T007162
Esiliculosus Arl13 Ec-19 003440
ARFRP1
Vbrassicaformis ARFRP1L Vbra 13023.t1-p1
Cvelia ARFRP1L Cvel 21819.t1-p1
Smicroadriaticum ARFRP1L Smic4787
Pmarinus ARFRP1L EER08613
Psojae ARFRP1L jgiPhyso3513414e gw1.6.6515.1
Pultimum ARFRP1L PYU1 T007978
Otrifallax ARFRP1L Contig5509.0.g71ADP-ribosylation
Gtheta ARFRP1L jgiGuith1111357au.51 g12459
Aanophagefferens ARFRP1L jgiAuran154947
Esiliculosus ARFRP1L Ec-18 002030
Bnatans ARFRP1L jgiBigna187953estExt fgenesh1 pg.C 260079
Ehuxleyi ARFRP1L jgiEmihu1366595fgenesh newKGs kg.224 24 EhuArfrp1
Cparadoxa ARFRP1L ConsensusfromContig7087
Creinhardtii ARFRP1L Cre03.g204353.t1.1
Athaliana ARFRP1L AT5G52210.1
Arl6
Psojae Arl6 jgiPhyso3256755Physo2.s 462293
Pultimum Arl6 PYU1 T014725
Cparadoxa Arl6 ConsensusfromContig37098
Gtheta Arl6 jgiGuith197017estExt Genewise1.C 860029
Bnatans Arl6 jgiBigna155693estExt Genewise1Plus.C 670078
Bnatans Arl6 jgiBigna146781estExt Genewise1.C 70056
Creinhardtii Arl6 Cre16.g664500.t1.2
Otrifallax Arl6 Contig15685.0.g21ADP-ribosylation
Otrifallax Arl6 Contig19717.0.g82ADP-ribosylation
Skawagutii Arl6 Skav229246
Smicroadriaticum Arl6 Smic27882
Vbrassicaformis Arl6 Vbra 20709.t1-p1
Tthermophila Arl6 TTHERM 00535830
Ptetraurelia Arl6 GSPATP00017502001
Cvelia Arl6 Cvel 1726.t1-p1
Arl8
Tthermophila Arl8 TTHERM 00185940
Ptetraurelia Arl8 GSPATP00022230001
Ptricornutum Arl8 jgiPhatr213157e gw1.10.215.1
Tpseudonana Arl8 jgiThaps37249fgenesh1 pg.C chr 7000632
Esiliculosus Arl8 Ec-19 003500
Pultimum Arl8 PYU1 T012553
Bnatans Arl8 jgiBigna192430estExt fgenesh1 pm.C 220009
Ccrispus Arl8 CDF39217
Creinhardtii Arl8 Cre17.g708250.t1.2
Athaliana Arl8 AT3G49870.1
Athaliana Arl8 AT3G49860.1
Otrifallax Arl8 Contig19.1.g92ADP-ribosylation
Ptetraurelia Arl8 GSPATP00026094001
Imultifiliis Arl8 IMG5 183110
Tthermophila Arl8 TTHERM 00499520
Arl16
Cparadoxa Arl16 ConsensusfromContig6963
Ehuxleyi Arl16 jgiEmihu179321est orfs.83419 EhuArl16
Pultimum Arl16 PYU1 T009193
Psojae Arl16 jgiPhyso3536135e gw1.28.106.1
Otrifallax Arl16 Contig11015.0.0.g25ADP-ribosylation
Creinhardtii Arl16 Cre11.g467564.t1.1
Bnatans Arl16 jgiBigna179720fgenesh1 pg.65 1
ArlX3
Vbrassicaformis ArlX3 Vbra 21855.t1-p1
Cvelia ArlX3 Cvel 13679.t1-p1
Sneurona ArlX3 SN3 00800140-mRNA-1-p1
Hhammondi ArlX3 HHA 288260-t26 1-p1
Tgondii-ME49 ArlX3 TGME49 288260-t26 1-p1
Ncaninum ArlX3 NCLIV 040770-t26 1-p1
Bbigemina ArlX3 BBBOND 0105000-t26 1-p1
Bbovis ArlX3 BBOV II005290-t26 1-p1
Cfelis ArlX3 CF001802.mRNA-p1
Tannulata ArlX3 TA15410-t26 1-p1
Tparva ArlX3 TP02 0835-t26 1-p1
Torientalis ArlX3 TOT020000871-t26 1-p1
Bmicroti ArlX3 BMR1 01G03180-t32 1-p1
Pberghei ArlX3 PBANKA 1414700.1-p1
Pyoelii ArlX3 PYYM 1418200.1-p1
Pchabaudi ArlX3 PCHAS 1416500.1-p1
Pknowlesi ArlX3 PKNH 1416900.1-p1
Pvivax ArlX3 PVX 122635.1-p1
Povale ArlX3 PocGH01 14024300.1-p1
Pgallinaceum ArlX3 PGAL8A 00512500.1-p1
Prelictum ArlX3 PRELSG 1414100.1-p1
Pfalciparum ArlX3 PF3D7 1316200.1-p1
Preichenowi ArlX3 PRCDC 1315200.1-p1
ARF
Sneurona Arf SN3 01400280-mRNA-1-p1
Hhammondi Arf HHA 276140-t26 1-p1
Ncaninum Arf NCLIV 007000-t26 1-p1
Cmuris Arf CMU 042540-t26 1-p1
Chominis Arf ChTU502y2012 390g0140-t32 1-p1
Cparvum Arf cgd2 3410-t26 1-p1
Cvelia Arf Cvel 32158.t1-p1
Pmarinus Arf EER18029
Pmarinus Arf EER18030
Pmarinus Arf EER17328
Gtheta Arf jgiGuith1153253fgenesh2 kg.46 6 5420 1 CCFICCFN EXTA EXTB
Creinhardtii Arf Cre02.g142687.t1.1
Cparadoxa Arf ConsensusfromContig9324
Aanophagefferens Arf jgiAuran169741
Tpseudonana Arf jgiThaps339299estExt gwp gw1.C chr 10802
Arl5
Cvelia Arl5 Cvel 14843.t1-p1
Vbrassicaformis Arl5 Vbra 22881.t1-p1
Sminutum Arl5 symbB.v1.2.011263.t1scaffold748.1size16575417
Smicroadriaticum Arl5 Smic19382
Pmarinus Arl5 EER10263
Cparadoxa Arl5 ConsensusfromContig8687
Tthermophila Arl5 TTHERM 00691110
Imultifiliis Arl5 IMG5 157450
Otrifallax Arl5 Contig10377.0.g65ADP-ribosylation
Gtheta Arl5 jgiGuith175303e gw1.65.17.1
Ccrispus Arl5 CDF36918
Bnatans Arl5 jgiBigna136398e gw1.14.123.1
Creinhardtii Arl5 Cre02.g102000.t1.2
Athaliana Arl5 AT3G22950.1
Ehuxleyi Arl5 jgiEmihu1434388estExtDG fgenesh newKGs kg.C 180119
Arl1
Athaliana Arl1 AT2G24765.1
Gtheta Arl1 jgiGuith1154446fgenesh2 kg.76 4 5272 1 CCFICCFN EXTA EXTB
Vbrassicaformis Arl1 Vbra 21724.t1-p1
Esiliculosus Arl1 Ec-19 004060
Aanophagefferens Arl1 jgiAuran160231
Ptricornutum Arl1 jgiPhatr28659e gw1.1.807.1
Psojae Arl1 jgiPhyso3314739fgenesh1 pm.4 499
Pultimum Arl1 PYU1 T008671
Pfalciparum Arl1 PF3D7 1034700.1-p1
Preichenowi Arl1 PRCDC 1034000.1-p1
Pgallinaceum Arl1 PGAL8A 00494200.1-p1
Pyoelii Arl1 PYYM 0519000.1-p1
Cvelia Arl1 Cvel 24063.t1-p1
Ncaninum Arl1 NCLIV 0250-t26 1-p1
Tthermophila Arl1 TTHERM 00577350
Arl2
Pultimum Arl2 PYU1 T011980
Psojae Arl2 jgiPhyso3313188fgenesh1 pm.3 709
Esiliculosus Arl2 Ec-14 003380
Tpseudonana Arl2 jgiThaps332795e gw1.3.670.1
Tthermophila Arl2 TTHERM 00120910
Imultifiliis Arl2 IMG5 090240
Ptetraurelia Arl2 GSPATP00029917001
Ehuxleyi Arl2 jgiEmihu1362631fgenesh newKGs kg.134 32 EhuArl2
Gtheta Arl2 jgiGuith187487estExt Genewise1Plus.C 410056
Sminutum Arl2 symbB.v1.2.021346.t1scaffold1839.1size992682
Smicroadriaticum Arl2 Smic28576
Pmarinus Arl2 EER03874
Athaliana Arl2 AT2G18390.1
Bnatans Arl2 jgiBigna157035fgenesh1 pm.2 2
Vbrassicaformis Arl2 Vbra 21203.t1-p1
Arl3
Otrifallax Arl3 Contig22207.0.g77ADP-ribosylation
Creinhardtii Arl3 Cre04.g218250.t1.2
Psojae Arl3 jgiPhyso3337530gm1.17105 g
Pultimum Arl3 PYU1 T006404
Cparadoxa Arl3 ConsensusfromContig7647
Tpseudonana Arl3 jgiThaps331406e gw1.1.731.1
Esiliculosus Arl3 Ec-01 009860
Gtheta Arl3 jgiGuith185562estExt Genewise1Plus.C 160182
Vbrassicaformis Arl3 Vbra 13454.t1-p1
Cvelia Arl3 Cvel 28880.t1-p1
Ptetraurelia Arl3 GSPATP00012122001
Pmarinus Arl3 EER09810
Sminutum Arl3 symbB.v1.2.005021.t1scaffold288.1size47836651
Smicroadriaticum Arl3 Smic4709
Sminutum Arl3 symbB.v1.2.004027.t1scaffold227.1size26120113
Sar1
Hhammondi Sar1 HHA 215060-t26 1-p1
Ncaninum Sar1 NCLIV 052070-t26 1-p1
Tgondii-ME49 Sar1 TGME49 215060-t26 1-p1
Vbrassicaformis Sar1 Vbra 15177.t1-p1
Sminutum Sar1 symbB.v1.2.027853.t1scaffold2872.1size684084
Cvelia Sar1 Cvel 5062.t1-p1
Cmuris Sar1 CMU 034380-t26 1-p1
Pgallinaceum Sar1 PGAL8A 00090300.1-p1
Prelictum Sar1 PRELSG 0512100.1-p1
Povale Sar1 PocGH01 00016700.1-p1
Pvivax Sar1 PVX 089930.1-p1
Pknowlesi Sar1 PKNH 0509200.1-p1
Pberghei Sar1 PBANKA 0718800.1-p1
Pyoelii Sar1 PYYM 0718900.1-p1
Pchabaudi Sar1 PCHAS 0727900.1-p1
0.820319

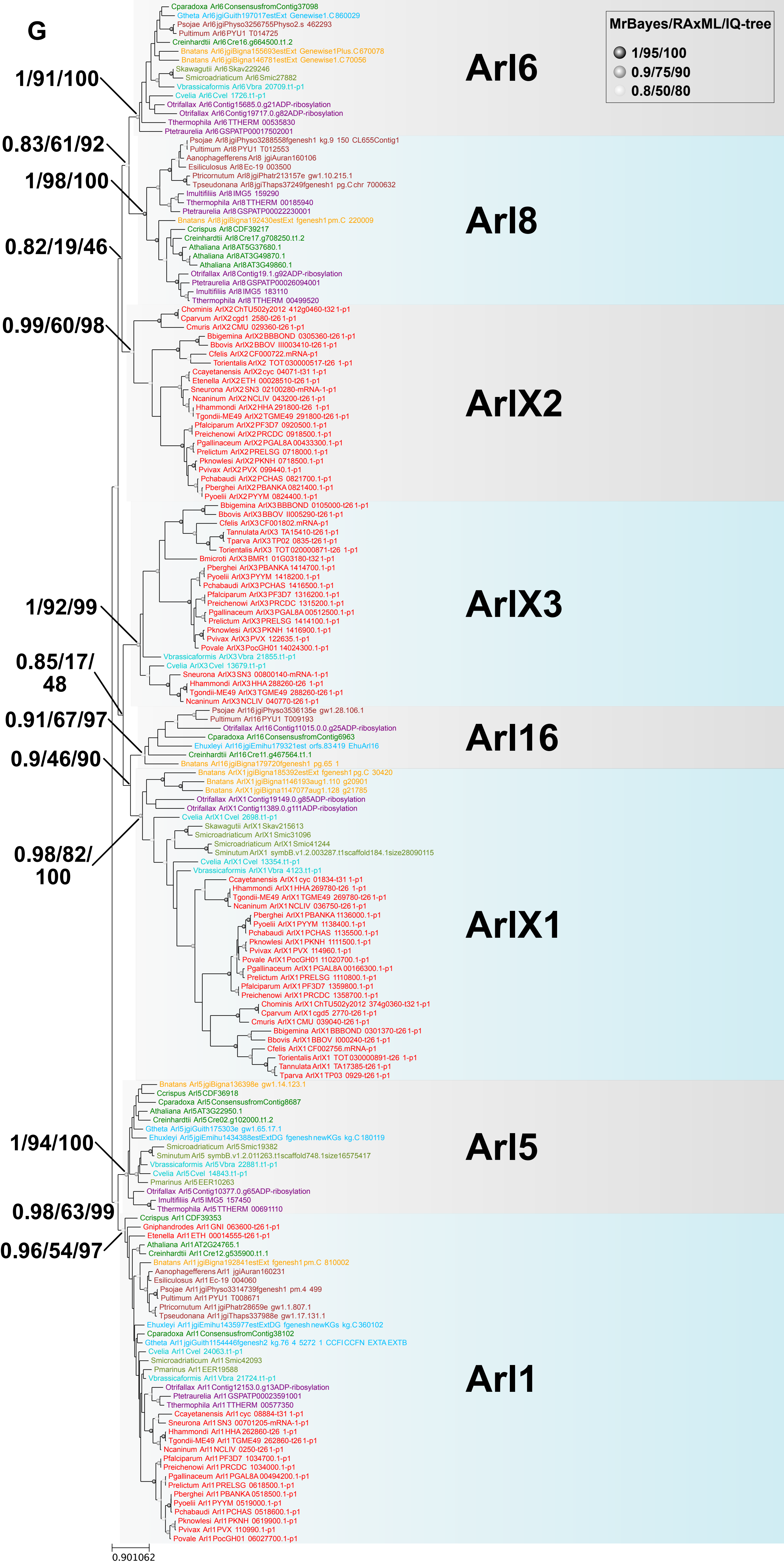

G
MrBayes/RAxML/IQ-tree
1/95/100
0.9/75/90
0.8/50/80
Arl6
Arl8
ArlX2
ArlX3
Arl16
ArlX1
Arl5
Arl1
1/91/100
0.83/61/92
1/98/100
0.82/19/46
0.99/60/98
1/92/99
0.85/17/48
0.91/67/97
0.9/46/90
0.98/82/100
1/94/100
0.98/63/99
0.96/54/97
0.901062
Cparadoxa Arl6 ConsensusfromContig37098
Gtheta Arl6 jgiGuith197017estExt Genewise1.C 860029
Psojae Arl6 jgiPhyso3256755Physo2.s 462293
Pultimum Arl6 PYU1 T014725
Creinhardtii Arl6 Cre16.g664500.t1.2
Bnatans Arl6 jgiBigna155693estExt Genewise1Plus.C 670078
Bnatans Arl6 jgiBigna146781estExt Genewise1.C 70056
Skawagutii Arl6 Skav229246
Smicroadriaticum Arl6 Smic27882
Vbrassicaformis Arl6 Vbra 20709.t1-p1
Cvelia Arl6 Cvel 1726.t1-p1
Otrifallax Arl6 Contig15685.0.g21ADP-ribosylation
Otrifallax Arl6 Contig19717.0.g82ADP-ribosylation
Tthermophila Arl6 TTHERM 00535830
Ptetraurelia Arl6 GSPATP00017502001
Psojae Arl8 jgiPhyso3288558fgenesh1 kg.9 150 CL655Contig1
Pultimum Arl8 PYU1 T012553
Aanophagefferens Arl8 jgiAuran160106
Esiliculosus Arl8 Ec-19 003500
Ptricornutum Arl8 jgiPhatr213157e gw1.10.215.1
Tpseudonana Arl8 jgiThaps37249fgenesh1 pg.C chr 7000632
Imultifiliis Arl8 IMG5 159290
Tthermophila Arl8 TTHERM 00185940
Ptetraurelia Arl8 GSPATP00022230001
Bnatans Arl8 jgiBigna192430estExt fgenesh1 pm.C 220009
Ccrispus Arl8 CDF39217
Creinhardtii Arl8 Cre17.g708250.t1.2
Athaliana Arl8 AT5G37680.1
Athaliana Arl8 AT3G49870.1
Athaliana Arl8 AT3G49860.1
Otrifallax Arl8 Contig19.1.g92ADP-ribosylation
Ptetraurelia Arl8 GSPATP00026094001
Imultifiliis Arl8 IMG5 183110
Tthermophila Arl8 TTHERM 00499520
Chominis ArlX2 ChTU502y2012 412g0460-t32 1-p1
Cparvum ArlX2 cgd1 2580-t26 1-p1
Cmuris ArlX2 CMU 029360-t26 1-p1
Bbigemina ArlX2 BBBOND 0305360-t26 1-p1
Bbovis ArlX2 BBOV III003410-t26 1-p1
Cfelis ArlX2 CF000722.mRNA-p1
Torientalis ArlX2 TOT 030000517-t26 1-p1
Ccayetanensis ArlX2 cyc 04071-t31 1-p1
Etenella ArlX2 ETH 00028510-t26 1-p1
Sneurona ArlX2 SN3 02100280-mRNA-1-p1
Ncaninum ArlX2 NCLIV 043200-t26 1-p1
Hhammondi ArlX2 HHA 291800-t26 1-p1
Tgondii-ME49 ArlX2 TGME49 291800-t26 1-p1
Pfalciparum ArlX2 PF3D7 0920500.1-p1
Preichenowi ArlX2 PRCDC 0918500.1-p1
Pgallinaceum ArlX2 PGAL8A 00433300.1-p1
Prelictum ArlX2 PRELSG 0718000.1-p1
Pknowlesi ArlX2 PKNH 0718500.1-p1
Pvivax ArlX2 PVX 099440.1-p1
Pchabaudi ArlX2 PCHAS 0821700.1-p1
Pberghei ArlX2 PBANKA 0821400.1-p1
Pyoelii ArlX2 PYYM 0824400.1-p1
Bbigemina ArlX3 BBBOND 0105000-t26 1-p1
Bbovis ArlX3 BBOV II005290-t26 1-p1
Cfelis ArlX3 CF001802.mRNA-p1
Tannulata ArlX3 TA15410-t26 1-p1
Tparva ArlX3 TP02 0835-t26 1-p1
Torientalis ArlX3 TOT 020000871-t26 1-p1
Bmicroti ArlX3 BMR1 01G03180-t32 1-p1
Pberghei ArlX3 PBANKA 1414700.1-p1
Pyoelii ArlX3 PYYM 1418200.1-p1
Pchabaudi ArlX3 PCHAS 1416500.1-p1
Pfalciparum ArlX3 PF3D7 1316200.1-p1
Preichenowi ArlX3 PRCDC 1315200.1-p1
Pgallinaceum ArlX3 PGAL8A 00512500.1-p1
Prelictum ArlX3 PRELSG 1414100.1-p1
Pknowlesi ArlX3 PKNH 1416900.1-p1
Pvivax ArlX3 PVX 122635.1-p1
Povale ArlX3 PocGH01 14024300.1-p1
Vbrassicaformis ArlX3 Vbra 21855.t1-p1
Cvelia ArlX3 Cvel 13679.t1-p1
Sneurona ArlX3 SN3 00800140-mRNA-1-p1
Hhammondi ArlX3 HHA 288260-t26 1-p1
Tgondii-ME49 ArlX3 TGME49 288260-t26 1-p1
Ncaninum ArlX3 NCLIV 040770-t26 1-p1
Psojae Arl16 jgiPhyso3536135e gw1.28.106.1
Pultimum Arl16 PYU1 T009193
Otrifallax Arl16 Contig11015.0.0.g25ADP-ribosylation
Cparadoxa Arl16 ConsensusfromContig6963
Ehuxleyi Arl16 jgiEmihu179321est orfs.83419 EhuArl16
Creinhardtii Arl16 Cre11.g467564.t1.1
Bnatans Arl16 jgiBigna179720fgenesh1 pg.65 1
Bnatans ArlX1 jgiBigna185392estExt fgenesh1 pg.C 30420
Bnatans ArlX1 jgiBigna1146193aug1.110 g20901
Bnatans ArlX1 jgiBigna1147077aug1.128 g21785
Otrifallax ArlX1 Contig19149.0.g85ADP-ribosylation
Otrifallax ArlX1 Contig11389.0.g111ADP-ribosylation
Cvelia ArlX1 Cvel 2698.t1-p1
Skawagutii ArlX1 Skav215613
Smicroadriaticum ArlX1 Smic31096
Smicroadriaticum ArlX1 Smic41244
Sminutum ArlX1 symbB.v1.2.003287.t1scaffold184.1size28090115
Cvelia ArlX1 Cvel 13354.t1-p1
Vbrassicaformis ArlX1 Vbra 4123.t1-p1
Ccayetanensis ArlX1 cyc 01834-t31 1-p1
Hhammondi ArlX1 HHA 269780-t26 1-p1
Tgondii-ME49 ArlX1 TGME49 269780-t26 1-p1
Ncaninum ArlX1 NCLIV 036750-t26 1-p1
Pberghei ArlX1 PBANKA 1136000.1-p1
Pyoelii ArlX1 PYYM 1138400.1-p1
Pchabaudi ArlX1 PCHAS 1135500.1-p1
Pknowlesi ArlX1 PKNH 1111500.1-p1
Pvivax ArlX1 PVX 114960.1-p1
Povale ArlX1 PocGH01 11020700.1-p1
Pgallinaceum ArlX1 PGAL8A 00166300.1-p1
Prelictum ArlX1 PRELSG 1110800.1-p1
Pfalciparum ArlX1 PF3D7 1359800.1-p1
Preichenowi ArlX1 PRCDC 1358700.1-p1
Chominis ArlX1 ChTU502y2012 374g0360-t32 1-p1
Cparvum ArlX1 cgd5 2770-t26 1-p1
Cmuris ArlX1 CMU 039040-t26 1-p1
Bbigemina ArlX1 BBBOND 0301370-t26 1-p1
Bbovis ArlX1 BBOV I000240-t26 1-p1
Cfelis ArlX1 CF002756.mRNA-p1
Torientalis ArlX1 TOT 030000891-t26 1-p1
Tannulata ArlX1 TA17385-t26 1-p1
Tparva ArlX1 TP03 0929-t26 1-p1
Bnatans Arl5 jgiBigna136398e gw1.14.123.1
Ccrispus Arl5 CDF36918
Cparadoxa Arl5 ConsensusfromContig8687
Athaliana Arl5 AT3G22950.1
Creinhardtii Arl5 Cre02.g102000.t1.2
Gtheta Arl5 jgiGuith175303e gw1.65.17.1
Ehuxleyi Arl5 jgiEmihu1434388estExtDG fgeneshnewKGs kg.C 180119
Smicroadriaticum Arl5 Smic19382
Sminutum Arl5 symbB.v1.2.011263.t1scaffold748.1size16575417
Vbrassicaformis Arl5 Vbra 22881.t1-p1
Cvelia Arl5 Cvel 14843.t1-p1
Pmarinus Arl5 EER10263
Otrifallax Arl5 Contig10377.0.g65ADP-ribosylation
Imultifiliis Arl5 IMG5 157450
Tthermophila Arl5 TTHERM 00691110
Ccrispus Arl1 CDF39353
Gniphandrodes Arl1 GNI 063600-t26 1-p1
Etenella Arl1 ETH 00014555-t26 1-p1
Athaliana Arl1 AT2G24765.1
Creinhardtii Arl1 Cre12.g535900.t1.1
Bnatans Arl1 jgiBigna192841estExt fgenesh1 pm.C 810002
Aanophagefferens Arl1 jgiAuran160231
Esiliculosus Arl1 Ec-19 004060
Psojae Arl1 jgiPhyso3314739fgenesh1 pm.4 499
Pultimum Arl1 PYU1 T008671
Ptricornutum Arl1 jgiPhatr28659e gw1.1.807.1
Tpseudonana Arl1 jgiThaps337988e gw1.17.131.1
Ehuxleyi Arl1 jgiEmihu1435977estExtDG fgeneshnewKGs kg.C 360102
Cparadoxa Arl1 ConsensusfromContig38102
Gtheta Arl1 jgiGuith1154446fgenesh2 kg.76 4 5272 1 CCFICCFN EXTA EXTB
Cvelia Arl1 Cvel 24063.t1-p1
Smicroadriaticum Arl1 Smic42093
Pmarinus Arl1 EER19588
Vbrassicaformis Arl1 Vbra 21724.t1-p1
Otrifallax Arl1 Contig12153.0.g13ADP-ribosylation
Ptetraurelia Arl1 GSPATP00023591001
Tthermophila Arl1 TTHERM 00577350
Ccayetanensis Arl1 cyc 08884-t31 1-p1
Sneurona Arl1 SN3 00701205-mRNA-1-p1
Hhammondi Arl1 HHA 262860-t26 1-p1
Tgondii-ME49 Arl1 TGME49 262860-t26 1-p1
Ncaninum Arl1 NCLIV 0250-t26 1-p1
Pfalciparum Arl1 PF3D7 1034700.1-p1
Preichenowi Arl1 PRCDC 1034000.1-p1
Pgallinaceum Arl1 PGAL8A 00494200.1-p1
Prelictum Arl1 PRELSG 0618500.1-p1
Pberghei Arl1 PBANKA 0518500.1-p1
Pyoelii Arl1 PYYM 0519000.1-p1
Pchabaudi Arl1 PCHAS 0518600.1-p1
Pknowlesi Arl1 PKNH 0619900.1-p1
Pvivax Arl1 PVX 110990.1-p1
Povale Arl1 PocGH01 06027700.1-p1

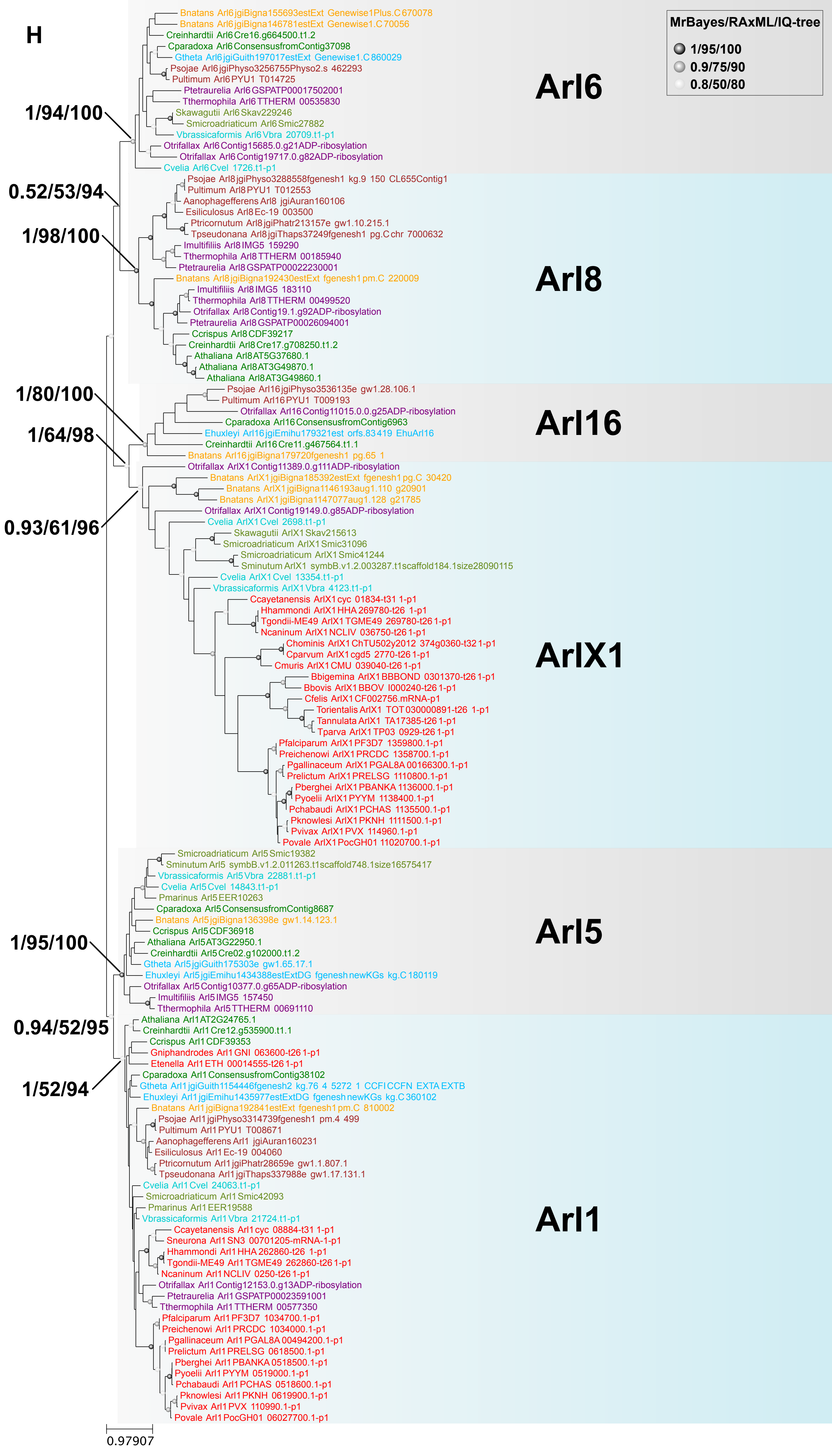
H
MrBayes/RAxML/IQ-tree
1/95/100
0.9/75/90
0.8/50/80
Arl6
Arl8
Arl16
ArlX1
Arl5
Arl1
1/94/100
0.52/53/94
1/98/100
1/80/100
1/64/98
0.93/61/96
1/95/100
0.94/52/95
1/52/94
Bnatans Arl6 jgiBigna155693estExt Genewise1Plus.C 670078
Bnatans Arl6 jgiBigna146781estExt Genewise1.C 70056
Creinhardtii Arl6 Cre16.g664500.t1.2
Cparadoxa Arl6 ConsensusfromContig37098
Gtheta Arl6 jgiGuith197017estExt Genewise1.C 860029
Psojae Arl6 jgiPhyso3256755Physo2.s 462293
Pultimum Arl6 PYU1 T014725
Ptetraurelia Arl6 GSPATP00017502001
Tthermophila Arl6 TTHERM 00535830
Skawagutii Arl6 Skav229246
Smicroadriaticum Arl6 Smic27882
Vbrassicaformis Arl6 Vbra 20709.t1-p1
Otrifallax Arl6 Contig15685.0.g21ADP-ribosylation
Otrifallax Arl6 Contig19717.0.g82ADP-ribosylation
Cvelia Arl6 Cvel 1726.t1-p1
Psojae Arl8 jgiPhyso3288558fgenesh1 kg.9 150 CL655Contig1
Pultimum Arl8 PYU1 T012553
Aanophagefferens Arl8 jgiAuran160106
Esiliculosus Arl8 Ec-19 003500
Ptricornutum Arl8 jgiPhatr213157e gw1.10.215.1
Tpseudonana Arl8 jgiThaps37249fgenesh1 pg.C chr 7000632
Imultifiliis Arl8 IMG5 159290
Tthermophila Arl8 TTHERM 00185940
Ptetraurelia Arl8 GSPATP00022230001
Bnatans Arl8 jgiBigna192430estExt fgenesh1 pm.C 220009
Imultifiliis Arl8 IMG5 183110
Tthermophila Arl8 TTHERM 00499520
Otrifallax Arl8 Contig19.1.g92ADP-ribosylation
Ptetraurelia Arl8 GSPATP00026094001
Ccrispus Arl8 CDF39217
Creinhardtii Arl8 Cre17.g708250.t1.2
Athaliana Arl8 AT5G37680.1
Athaliana Arl8 AT3G49870.1
Athaliana Arl8 AT3G49860.1
Psojae Arl16 jgiPhyso3536135e gw1.28.106.1
Pultimum Arl16 PYU1 T009193
Otrifallax Arl16 Contig11015.0.0.g25ADP-ribosylation
Cparadoxa Arl16 ConsensusfromContig6963
Ehuxleyi Arl16 jgiEmihu179321est orfs.83419 EhuArl16
Creinhardtii Arl16 Cre11.g467564.t1.1
Bnatans Arl16 jgiBigna179720fgenesh1 pg.65 1
Otrifallax ArlX1 Contig11389.0.g111ADP-ribosylation
Bnatans ArlX1 jgiBigna185392estExt fgenesh1 pg.C 30420
Bnatans ArlX1 jgiBigna1146193aug1.110 g20901
Bnatans ArlX1 jgiBigna1147077aug1.128 g21785
Otrifallax ArlX1 Contig19149.0.g85ADP-ribosylation
Cvelia ArlX1 Cvel 2698.t1-p1
Skawagutii ArlX1 Skav215613
Smicroadriaticum ArlX1 Smic31096
Smicroadriaticum ArlX1 Smic41244
Sminutum ArlX1 symbB.v1.2.003287.t1scaffold184.1size28090115
Cvelia ArlX1 Cvel 13354.t1-p1
Vbrassicaformis ArlX1 Vbra 4123.t1-p1
Ccayetanensis ArlX1 cyc 01834-t31 1-p1
Hhammondi ArlX1 HHA 269780-t26 1-p1
Tgondii-ME49 ArlX1 TGME49 269780-t26 1-p1
Ncaninum ArlX1 NCLIV 036750-t26 1-p1
Chominis ArlX1 ChTU502y2012 374g0360-t32 1-p1
Cparvum ArlX1 cgd5 2770-t26 1-p1
Cmuris ArlX1 CMU 039040-t26 1-p1
Bbigemina ArlX1 BBBOND 0301370-t26 1-p1
Bbovis ArlX1 BBOV I000240-t26 1-p1
Cfelis ArlX1 CF002756.mRNA-p1
Torientalis ArlX1 TOT030000891-t26 1-p1
Tannulata ArlX1 TA17385-t26 1-p1
Tparva ArlX1 TP03 0929-t26 1-p1
Pfalciparum ArlX1 PF3D7 1359800.1-p1
Preichenowi ArlX1 PRCDC 1358700.1-p1
Pgallinaceum ArlX1 PGAL8A 00166300.1-p1
Prelictum ArlX1 PRELSG 1110800.1-p1
Pberghei ArlX1 PBANKA 1136000.1-p1
Pyoelii ArlX1 PYYM 1138400.1-p1
Pchabaudi ArlX1 PCHAS 1135500.1-p1
Pknowlesi ArlX1 PKNH 1111500.1-p1
Pvivax ArlX1 PVX 114960.1-p1
Povale ArlX1 PocGH01 11020700.1-p1
Smicroadriaticum Arl5 Smic19382
Sminutum Arl5 symbB.v1.2.011263.t1scaffold748.1size16575417
Vbrassicaformis Arl5 Vbra 22881.t1-p1
Cvelia Arl5 Cvel 14843.t1-p1
Pmarinus Arl5 EER10263
Cparadoxa Arl5 ConsensusfromContig8687
Bnatans Arl5 jgiBigna136398e gw1.14.123.1
Ccrispus Arl5 CDF36918
Athaliana Arl5 AT3G22950.1
Creinhardtii Arl5 Cre02.g102000.t1.2
Gtheta Arl5 jgiGuith175303e gw1.65.17.1
Ehuxleyi Arl5 jgiEmihu1434388estExtDG fgenesh newKGs kg.C 180119
Otrifallax Arl5 Contig10377.0.g65ADP-ribosylation
Imultifiliis Arl5 IMG5 157450
Tthermophila Arl5 TTHERM 00691110
Athaliana Arl1 AT2G24765.1
Creinhardtii Arl1 Cre12.g535900.t1.1
Ccrispus Arl1 CDF39353
Gniphandrodes Arl1 GNI 063600-t26 1-p1
Etenella Arl1 ETH 00014555-t26 1-p1
Cparadoxa Arl1 ConsensusfromContig38102
Gtheta Arl1 jgiGuith1154446fgenesh2 kg.76 4 5272 1 CCFI CCFN EXTA EXTB
Ehuxleyi Arl1 jgiEmihu1435977estExtDG fgenesh newKGs kg.C 360102
Bnatans Arl1 jgiBigna192841estExt fgenesh1 pm.C 810002
Psojae Arl1 jgiPhyso3314739fgenesh1 pm.4 499
Pultimum Arl1 PYU1 T008671
Aanophagefferens Arl1 jgiAuran160231
Esiliculosus Arl1 Ec-19 004060
Ptricornutum Arl1 jgiPhatr28659e gw1.807.1
Tpseudonana Arl1 jgiThaps337988e gw1.17.131.1
Cvelia Arl1 Cvel 24063.t1-p1
Smicroadriaticum Arl1 Smic42093
Pmarinus Arl1 EER19588
Vbrassicaformis Arl1 Vbra 21724.t1-p1
Ccayetanensis Arl1 cyc 08884-t31 1-p1
Sneurona Arl1 SN3 00701205-mRNA-1-p1
Hhammondi Arl1 HHA 262860-t26 1-p1
Tgondii-ME49 Arl1 TGME49 262860-t26 1-p1
Ncaninum Arl1 NCLIV 0250-t26 1-p1
Otrifallax Arl1 Contig12153.0.g13ADP-ribosylation
Ptetraurelia Arl1 GSPATP00023591001
Tthermophila Arl1 TTHERM 00577350
Pfalciparum Arl1 PF3D7 1034700.1-p1
Preichenowi Arl1 PRCDC 1034000.1-p1
Pgallinaceum Arl1 PGAL8A 00494200.1-p1
Prelictum Arl1 PRELSG 0618500.1-p1
Pberghei Arl1 PBANKA 0518500.1-p1
Pyoelii Arl1 PYYM 0519000.1-p1
Pchabaudi Arl1 PCHAS 0518600.1-p1
Pknowlesi Arl1 PKNH 0619900.1-p1
Pvivax Arl1 PVX 110990.1-p1
Povale Arl1 PocGH01 06027700.1-p1
0.97907

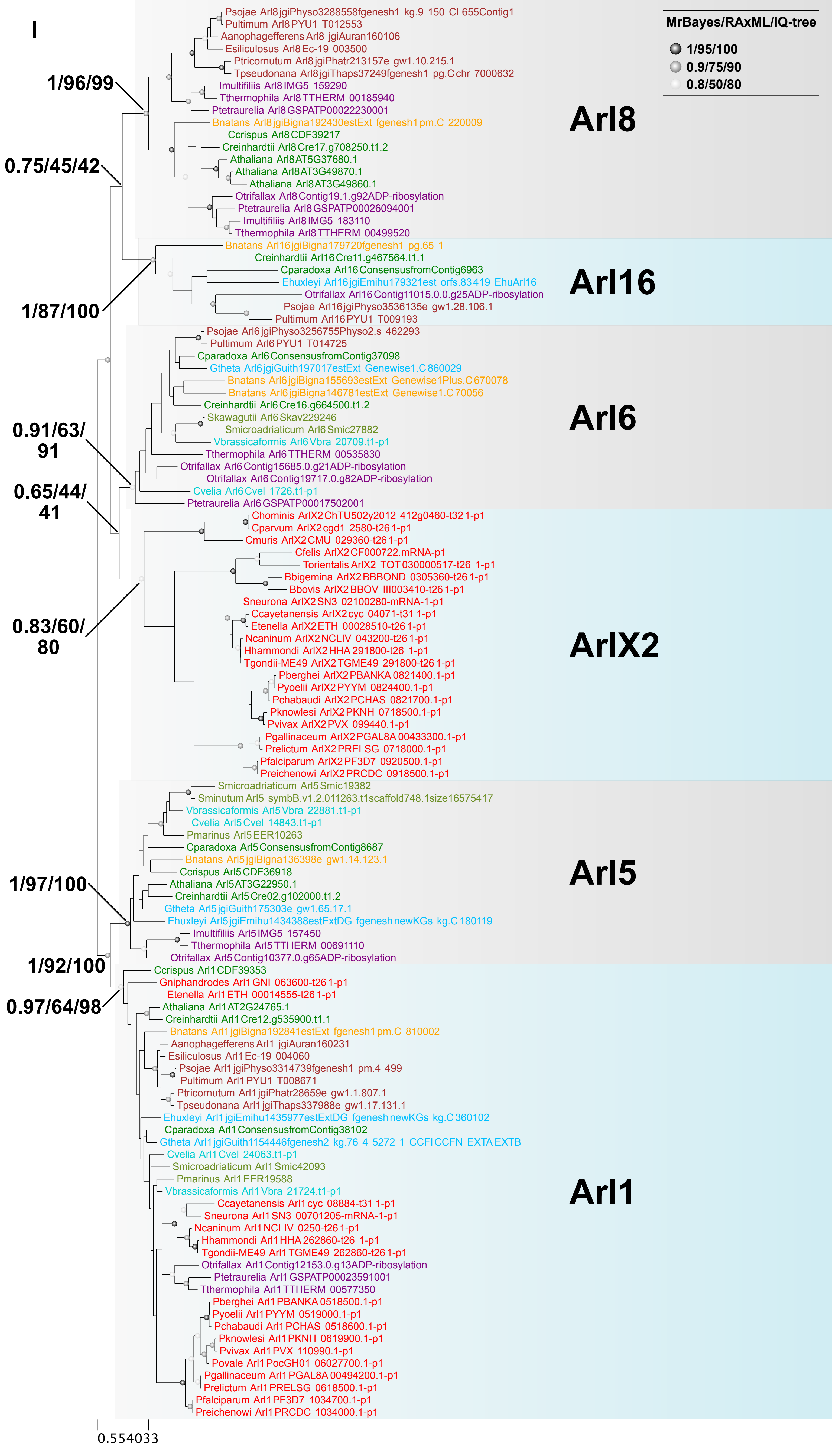

I
MrBayes/RAxML/IQ-tree
1/95/100
0.9/75/90
0.8/50/80
Psojae Arl8 jgiPhyso3288558fgenesh1 kg.9 150 CL655Contig1
Pultimum Arl8 PYU1 T012553
Aanophagefferens Arl8 jgiAuran160106
Esiliculosus Arl8 Ec-19 003500
Ptricornutum Arl8 jgiPhatr213157e gw1.10.215.1
Tpseudonana Arl8 jgiThaps37249fgenesh1 pg.C chr 7000632
Imultifiliis Arl8 IMG5 159290
Tthermophila Arl8 TTHERM 00185940
Ptetraurelia Arl8 GSPATP00022230001
Bnatans Arl8 jgiBigna192430estExt fgenesh1 pm.C 220009
Ccrispus Arl8 CDF39217
Creinhardtii Arl8 Cre17.g708250.t1.2
Athaliana Arl8 AT5G37680.1
Athaliana Arl8 AT3G49870.1
Athaliana Arl8 AT3G49860.1
Otrifallax Arl8 Contig19.1.g92ADP-ribosylation
Ptetraurelia Arl8 GSPATP00026094001
Imultifiliis Arl8 IMG5 183110
Tthermophila Arl8 TTHERM 00499520
Arl8
1/96/99
0.75/45/42
Bnatans Arl16 jgiBigna179720fgenesh1 pg.65 1
Creinhardtii Arl16 Cre11.g467564.t1.1
Cparadoxa Arl16 ConsensusfromContig6963
Ehuxleyi Arl16 jgiEmihu179321est orfs.83419 EhuArl16
Otrifallax Arl16 Contig11015.0.0.g25ADP-ribosylation
Psojae Arl16 jgiPhyso3536135e gw1.28.106.1
Pultimum Arl16 PYU1 T009193
Arl16
1/87/100
Psojae Arl6 jgiPhyso3256755Physo2.s 462293
Pultimum Arl6 PYU1 T014725
Cparadoxa Arl6 ConsensusfromContig37098
Gtheta Arl6 jgiGuith197017estExt Genewise1.C 860029
Bnatans Arl6 jgiBigna155693estExt Genewise1Plus.C 670078
Bnatans Arl6 jgiBigna146781estExt Genewise1.C 70056
Creinhardtii Arl6 Cre16.g664500.t1.2
Skawagutii Arl6 Skav229246
Smicroadriaticum Arl6 Smic27882
Vbrassicaformis Arl6 Vbra 20709.t1-p1
Tthermophila Arl6 TTHERM 00535830
Otrifallax Arl6 Contig15685.0.g21ADP-ribosylation
Otrifallax Arl6 Contig19717.0.g82ADP-ribosylation
Cvelia Arl6 Cvel 1726.t1-p1
Ptetraurelia Arl6 GSPATP00017502001
Arl6
0.91/63/91
0.65/44/41
Chominis ArlX2 ChTU502y2012 412g0460-t32 1-p1
Cparvum ArlX2 cgd1 2580-t26 1-p1
Cmuris ArlX2 CMU 029360-t26 1-p1
Cfelis ArlX2 CF000722.mRNA-p1
Torientalis ArlX2 TOT 030000517-t26 1-p1
Bbigemina ArlX2 BBBOND 0305360-t26 1-p1
Bbovis ArlX2 BBOV III003410-t26 1-p1
Sneurona ArlX2 SN3 02100280-mRNA-1-p1
Ccayetanensis ArlX2 cyc 04071-t31 1-p1
Etenella ArlX2 ETH 00028510-t26 1-p1
Ncaninum ArlX2 NCLIV 043200-t26 1-p1
Hhammondi ArlX2 HHA 291800-t26 1-p1
Tgondii-ME49 ArlX2 TGME49 291800-t26 1-p1
Pberghei ArlX2 PBANKA 0821400.1-p1
Pyoelii ArlX2 PYYM 0824400.1-p1
Pchabaudi ArlX2 PCHAS 0821700.1-p1
Pknowlesi ArlX2 PKNH 0718500.1-p1
Pvivax ArlX2 PVX 099440.1-p1
Pgallinaceum ArlX2 PGAL8A 00433300.1-p1
Prelictum ArlX2 PRELSG 0718000.1-p1
Pfalciparum ArlX2 PF3D7 0920500.1-p1
Preichenowi ArlX2 PRCDC 0918500.1-p1
ArlX2
0.83/60/80
Smicroadriaticum Arl5 Smic19382
Sminutum Arl5 symbB.v1.2.011263.t1scaffold748.1size16575417
Vbrassicaformis Arl5 Vbra 22881.t1-p1
Cvelia Arl5 Cvel 14843.t1-p1
Pmarinus Arl5 EER10263
Cparadoxa Arl5 ConsensusfromContig8687
Bnatans Arl5 jgiBigna136398e gw1.14.123.1
Ccrispus Arl5 CDF36918
Athaliana Arl5 AT3G22950.1
Creinhardtii Arl5 Cre02.g102000.t1.2
Gtheta Arl5 jgiGuith175303e gw1.65.17.1
Ehuxleyi Arl5 jgiEmihu1434388estExtDG fgenesh newKGs kg.C 180119
Imultifiliis Arl5 IMG5 157450
Tthermophila Arl5 TTHERM 00691110
Otrifallax Arl5 Contig10377.0.g65ADP-ribosylation
Arl5
1/97/100
1/92/100
0.97/64/98
Ccrispus Arl1 CDF39353
Gniphandrodes Arl1 GNI 063600-t26 1-p1
Etenella Arl1 ETH 00014555-t26 1-p1
Athaliana Arl1 AT2G24765.1
Creinhardtii Arl1 Cre12.g535900.t1.1
Bnatans Arl1 jgiBigna192841estExt fgenesh1 pm.C 810002
Aanophagefferens Arl1 jgiAuran160231
Esiliculosus Arl1 Ec-19 004060
Psojae Arl1 jgiPhyso3314739fgenesh1 pm.4 499
Pultimum Arl1 PYU1 T008671
Ptricornutum Arl1 jgiPhatr28659e gw1.1.807.1
Tpseudonana Arl1 jgiThaps337988e gw1.17.131.1
Ehuxleyi Arl1 jgiEmihu1435977estExtDG fgenesh newKGs kg.C 360102
Cparadoxa Arl1 ConsensusfromContig38102
Gtheta Arl1 jgiGuith1154446fgenesh2 kg.76 4 5272 1 CCFI CCFN EXTA EXTB
Cvelia Arl1 Cvel 24063.t1-p1
Smicroadriaticum Arl1 Smic42093
Pmarinus Arl1 EER19588
Vbrassicaformis Arl1 Vbra 21724.t1-p1
Ccayetanensis Arl1 cyc 08884-t31 1-p1
Sneurona Arl1 SN3 00701205-mRNA-1-p1
Ncaninum Arl1 NCLIV 0250-t26 1-p1
Hhammondi Arl1 HHA 262860-t26 1-p1
Tgondii-ME49 Arl1 TGME49 262860-t26 1-p1
Otrifallax Arl1 Contig12153.0.g13ADP-ribosylation
Ptetraurelia Arl1 GSPATP00023591001
Tthermophila Arl1 TTHERM 00577350
Pberghei Arl1 PBANKA 0518500.1-p1
Pyoelii Arl1 PYYM 0519000.1-p1
Pchabaudi Arl1 PCHAS 0518600.1-p1
Pknowlesi Arl1 PKNH 0619900.1-p1
Pvivax Arl1 PVX 110990.1-p1
Povale Arl1 PocGH01 06027700.1-p1
Pgallinaceum Arl1 PGAL8A 00494200.1-p1
Prelictum Arl1 PRELSG 0618500.1-p1
Pfalciparum Arl1 PF3D7 1034700.1-p1
Preichenowi Arl1 PRCDC 1034000.1-p1
Arl1
0.554033

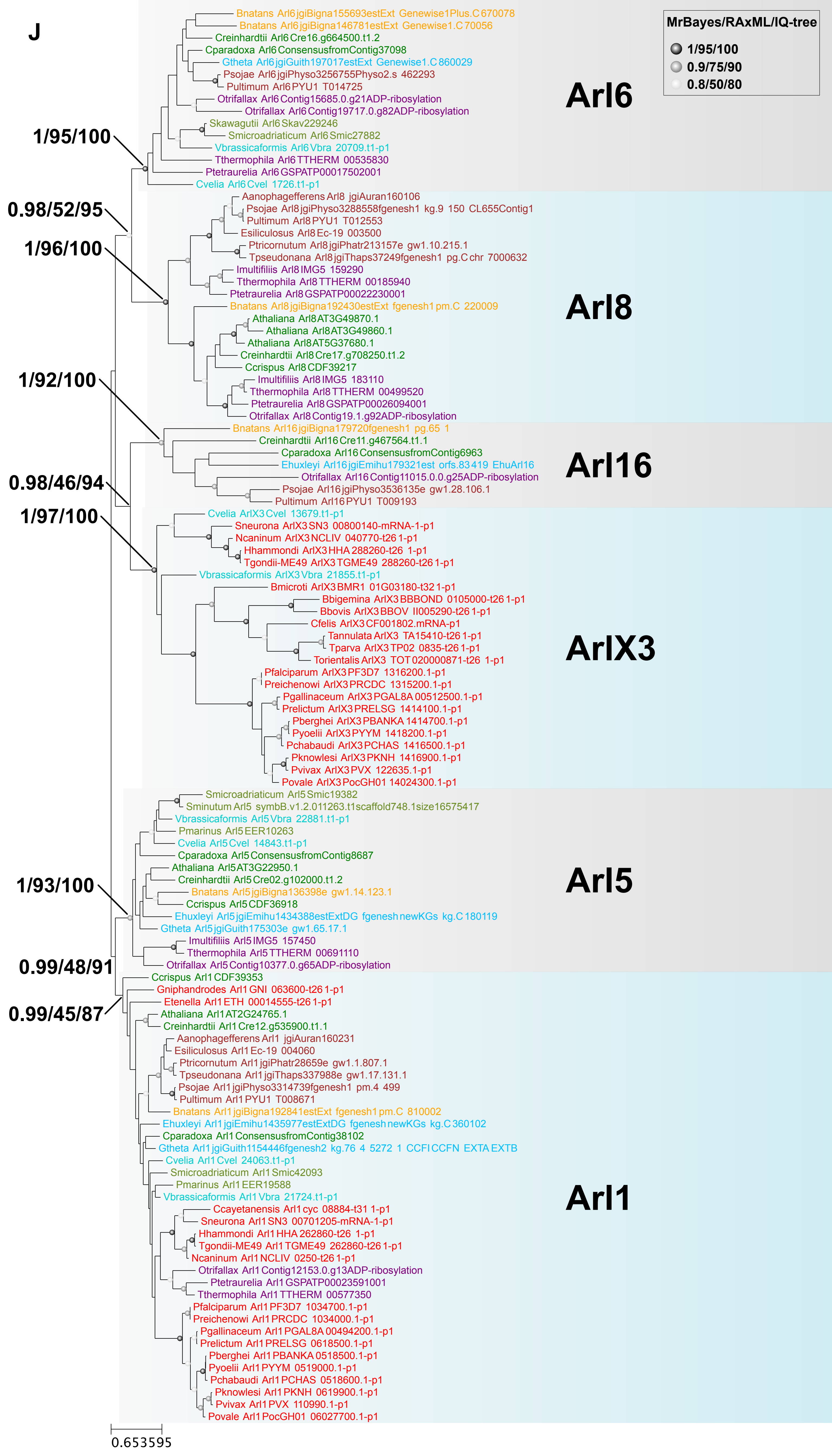

J
MrBayes/RAxML/IQ-tree
1/95/100
0.9/75/90
0.8/50/80
Arl6
Bnatans Arl6 jgiBigna155693estExt Genewise1Plus.C 670078
Bnatans Arl6 jgiBigna146781estExt Genewise1.C 70056
Creinhardtii Arl6 Cre16.g664500.t1.2
Cparadoxa Arl6 ConsensusfromContig37098
Gtheta Arl6 jgiGuith197017estExt Genewise1.C 860029
Psojae Arl6 jgiPhyso3256755Physo2.s 462293
Pultimum Arl6 PYU1 T014725
Otrifallax Arl6 Contig15685.0.g21ADP-ribosylation
Otrifallax Arl6 Contig19717.0.g82ADP-ribosylation
Skawagutii Arl6 Skav229246
Smicroadriaticum Arl6 Smic27882
Vbrassicaformis Arl6 Vbra 20709.t1-p1
Tthermophila Arl6 TTHERM 00535830
Ptetraurelia Arl6 GSPATP00017502001
Cvelia Arl6 Cvel 1726.t1-p1
1/95/100
0.98/52/95
1/96/100
Arl8
Aanophagefferens Arl8 jgiAuran160106
Psojae Arl8 jgiPhyso3288558fgenesh1 kg.9 150 CL655Contig1
Pultimum Arl8 PYU1 T012553
Esiliculosus Arl8 Ec-19 003500
Ptricornutum Arl8 jgiPhatr213157e gw1.10.215.1
Tpseudonana Arl8 jgiThaps37249fgenesh1 pg.C chr 7000632
Imultifiliis Arl8 IMG5 159290
Tthermophila Arl8 TTHERM 00185940
Ptetraurelia Arl8 GSPATP00022230001
Bnatans Arl8 jgiBigna192430estExt fgenesh1 pm.C 220009
Athaliana Arl8 AT3G49870.1
Athaliana Arl8 AT3G49860.1
Athaliana Arl8 AT5G37680.1
Creinhardtii Arl8 Cre17.g708250.t1.2
Ccrispus Arl8 CDF39217
Imultifiliis Arl8 IMG5 183110
Tthermophila Arl8 TTHERM 00499520
Ptetraurelia Arl8 GSPATP00026094001
Otrifallax Arl8 Contig19.1.g92ADP-ribosylation
1/92/100
Arl16
Bnatans Arl16 jgiBigna179720fgenesh1 pg.65 1
Creinhardtii Arl16 Cre11.g467564.t1.1
Cparadoxa Arl16 ConsensusfromContig6963
Ehuxleyi Arl16 jgiEmihu179321est orfs.83419 EhuArl16
Otrifallax Arl16 Contig11015.0.0.g25ADP-ribosylation
Psojae Arl16 jgiPhyso3536135e gw1.28.106.1
Pultimum Arl16 PYU1 T009193
0.98/46/94
1/97/100
ArlX3
Cvelia ArlX3 Cvel 13679.t1-p1
Sneurona ArlX3 SN3 00800140-mRNA-1-p1
Ncaninum ArlX3 NCLIV 040770-t26 1-p1
Hhammondi ArlX3 HHA 288260-t26 1-p1
Tgondii-ME49 ArlX3 TGME49 288260-t26 1-p1
Vbrassicaformis ArlX3 Vbra 21855.t1-p1
Bmicroti ArlX3 BMR1 01G03180-t32 1-p1
Bbigemina ArlX3 BBBOND 0105000-t26 1-p1
Bbovis ArlX3 BBOV II005290-t26 1-p1
Cfelis ArlX3 CF001802.mRNA-p1
Tannulata ArlX3 TA15410-t26 1-p1
Tparva ArlX3 TP02 0835-t26 1-p1
Torientalis ArlX3 TOT 020000871-t26 1-p1
Pfalciparum ArlX3 PF3D7 1316200.1-p1
Preichenowi ArlX3 PRCDC 1315200.1-p1
Pgallinaceum ArlX3 PGAL8A 00512500.1-p1
Prelictum ArlX3 PRELSG 1414100.1-p1
Pberghei ArlX3 PBANKA 1414700.1-p1
Pyoelii ArlX3 PYYM 1418200.1-p1
Pchabaudi ArlX3 PCHAS 1416500.1-p1
Pknowlesi ArlX3 PKNH 1416900.1-p1
Pvivax ArlX3 PVX 122635.1-p1
Povale ArlX3 PocGH01 14024300.1-p1
Arl5
Smicroadriaticum Arl5 Smic19382
Sminutum Arl5 symbB.v1.2.011263.t1scaffold748.1size16575417
Vbrassicaformis Arl5 Vbra 22881.t1-p1
Pmarinus Arl5 EER10263
Cvelia Arl5 Cvel 14843.t1-p1
Cparadoxa Arl5 ConsensusfromContig8687
Athaliana Arl5 AT3G22950.1
Creinhardtii Arl5 Cre02.g102000.t1.2
Bnatans Arl5 jgiBigna136398e gw1.14.123.1
Ccrispus Arl5 CDF36918
Ehuxleyi Arl5 jgiEmihu1434388estExtDG fgenesh newKGs kg.C 180119
Gtheta Arl5 jgiGuith175303e gw1.65.17.1
Imultifiliis Arl5 IMG5 157450
Tthermophila Arl5 TTHERM 00691110
Otrifallax Arl5 Contig10377.0.g65ADP-ribosylation
1/93/100
0.99/48/91
0.99/45/87
Arl1
Ccrispus Arl1 CDF39353
Gniphandrodes Arl1 GNI 063600-t26 1-p1
Etenella Arl1 ETH 00014555-t26 1-p1
Athaliana Arl1 AT2G24765.1
Creinhardtii Arl1 Cre12.g535900.t1.1
Aanophagefferens Arl1 jgiAuran160231
Esiliculosus Arl1 Ec-19 004060
Ptricornutum Arl1 jgiPhatr28659e gw1.1.807.1
Tpseudonana Arl1 jgiThaps337988e gw1.17.131.1
Psojae Arl1 jgiPhyso3314739fgenesh1 pm.4 499
Pultimum Arl1 PYU1 T008671
Bnatans Arl1 jgiBigna192841estExt fgenesh1 pm.C 810002
Ehuxleyi Arl1 jgiEmihu1435977estExtDG fgenesh newKGs kg.C 360102
Cparadoxa Arl1 ConsensusfromContig38102
Gtheta Arl1 jgiGuith1154446fgenesh2 kg.76 4 5272 1 CCFI CCFN EXTA EXTB
Cvelia Arl1 Cvel 24063.t1-p1
Smicroadriaticum Arl1 Smic42093
Pmarinus Arl1 EER19588
Vbrassicaformis Arl1 Vbra 21724.t1-p1
Ccayetanensis Arl1 cyc 08884-t31 1-p1
Sneurona Arl1 SN3 00701205-mRNA-1-p1
Hhammondi Arl1 HHA 262860-t26 1-p1
Tgondii-ME49 Arl1 TGME49 262860-t26 1-p1
Ncaninum Arl1 NCLIV 0250-t26 1-p1
Otrifallax Arl1 Contig12153.0.g13ADP-ribosylation
Ptetraurelia Arl1 GSPATP00023591001
Tthermophila Arl1 TTHERM 00577350
Pfalciparum Arl1 PF3D7 1034700.1-p1
Preichenowi Arl1 PRCDC 1034000.1-p1
Pgallinaceum Arl1 PGAL8A 00494200.1-p1
Prelictum Arl1 PRELSG 0618500.1-p1
Pberghei Arl1 PBANKA 0518500.1-p1
Pyoelii Arl1 PYYM 0519000.1-p1
Pchabaudi Arl1 PCHAS 0518600.1-p1
Pknowlesi Arl1 PKNH 0619900.1-p1
Pvivax Arl1 PVX 110990.1-p1
Povale Arl1 PocGH01 06027700.1-p1
0.653595

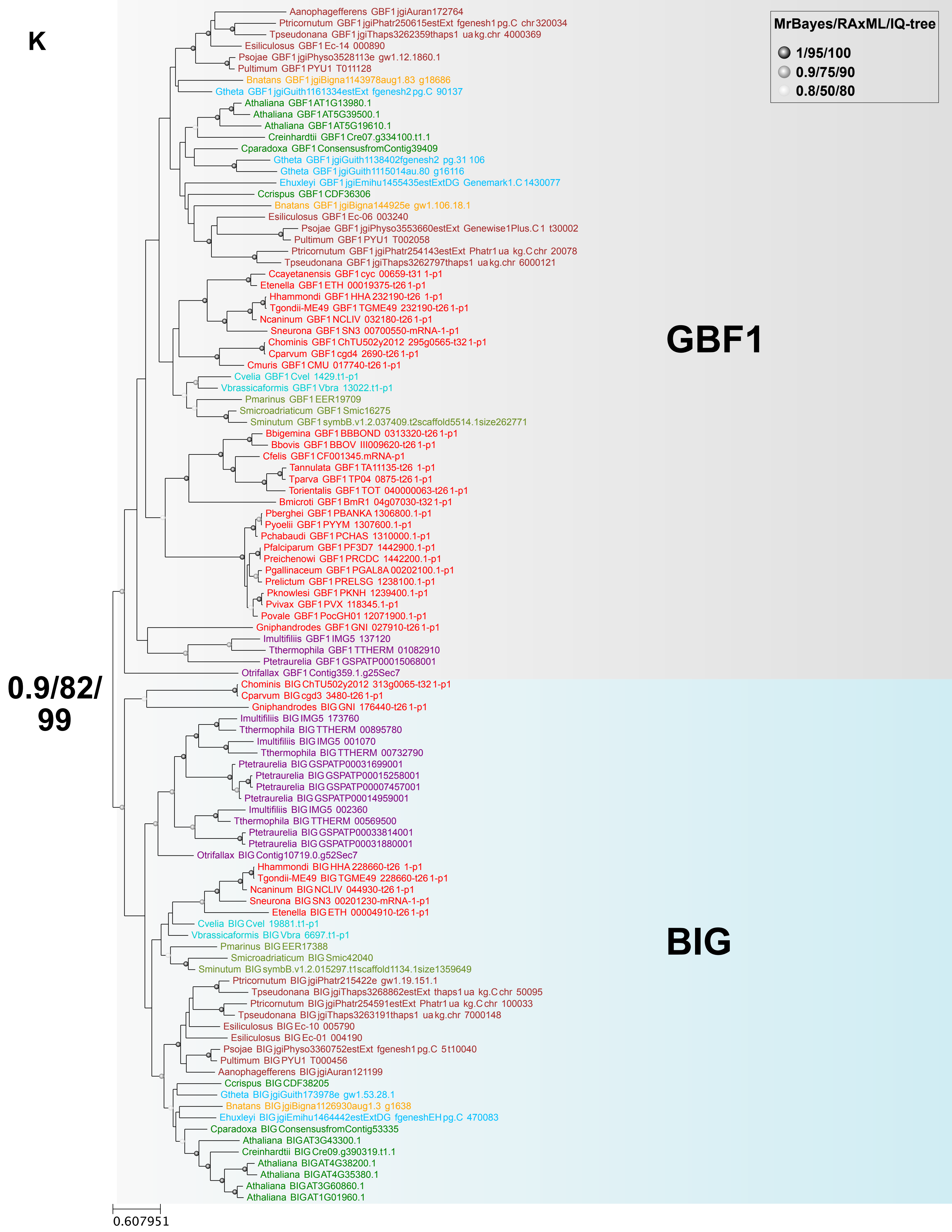
K
MrBayes/RAxML/IQ-tree
1/95/100
0.9/75/90
0.8/50/80
GBF1
BIG
0.9/82/
99
0.607951
Aanophagefferens GBF1 jgiAuran172764
Ptricornutum GBF1 jgiPhatr250615estExt fgenesh1 pg.C chr 320034
Tpseudonana GBF1 jgiThaps3262359thaps1 ua kg.chr 4000369
Esiliculosus GBF1 Ec-14 000890
Psojae GBF1 jgiPhyso3528113e gw1.12.1860.1
Pultimum GBF1 PYU1 T011128
Bnatans GBF1 jgiBigna1143978aug1.83 g18686
Gtheta GBF1 jgiGuith1161334estExt fgenesh2 pg.C 90137
Athaliana GBF1 AT1G13980.1
Athaliana GBF1 AT5G39500.1
Athaliana GBF1 AT5G19610.1
Creinhardtii GBF1 Cre07.g334100.t1.1
Cparadoxa GBF1 ConsensusfromContig39409
Gtheta GBF1 jgiGuith1138402fgenesh2 pg.31 106
Gtheta GBF1 jgiGuith1115014au.80 g16116
Ehuxleyi GBF1 jgiEmihu1455435estExtDG Genemark1.C 1430077
Ccrispus GBF1 CDF36306
Bnatans GBF1 jgiBigna144925e gw1.106.18.1
Esiliculosus GBF1 Ec-06 003240
Psojae GBF1 jgiPhyso3553660estExt Genewise1Plus.C 1 t30002
Pultimum GBF1 PYU1 T002058
Ptricornutum GBF1 jgiPhatr254143estExt Phatr1 ua kg.C chr 20078
Tpseudonana GBF1 jgiThaps3262797thaps1 ua kg.chr 6000121
Ccayetanensis GBF1 cyc 00659-t31 1-p1
Etenella GBF1 ETH 00019375-t26 1-p1
Hhammondi GBF1 HHA 232190-t26 1-p1
Tgondii-ME49 GBF1 TGME49 232190-t26 1-p1
Ncaninum GBF1 NCLIV 032180-t26 1-p1
Sneurona GBF1 SN3 00700550-mRNA-1-p1
Chominis GBF1 ChTU502y2012 295g0565-t32 1-p1
Cparvum GBF1 cgd4 2690-t26 1-p1
Cmuris GBF1 CMU 017740-t26 1-p1
Cvelia GBF1 Cvel 1429.t1-p1
Vbrassicaformis GBF1 Vbra 13022.t1-p1
Pmarinus GBF1 EER19709
Smicroadriaticum GBF1 Smic16275
Sminutum GBF1 symbB.v1.2.037409.t2scaffold5514.1size262771
Bbigemina GBF1 BBBOND 0313320-t26 1-p1
Bbovis GBF1 BBOV III009620-t26 1-p1
Cfelis GBF1 CF001345.mRNA-p1
Tannulata GBF1 TA11135-t26 1-p1
Tparva GBF1 TP04 0875-t26 1-p1
Torientalis GBF1 TOT 040000063-t26 1-p1
Bmicroti GBF1 BmR1 04g07030-t32 1-p1
Pberghei GBF1 PBANKA 1306800.1-p1
Pyoelii GBF1 PYYM 1307600.1-p1
Pchabaudi GBF1 PCHAS 1310000.1-p1
Pfalciparum GBF1 PF3D7 1442900.1-p1
Preichenowi GBF1 PRCDC 1442200.1-p1
Pgallinaceum GBF1 PGAL8A 00202100.1-p1
Prelictum GBF1 PRELSG 1238100.1-p1
Pknowlesi GBF1 PKNH 1239400.1-p1
Pvivax GBF1 PVX 118345.1-p1
Povale GBF1 PocGH01 12071900.1-p1
Gniphandrodes GBF1 GNI 027910-t26 1-p1
Imultifiliis GBF1 IMG5 137120
Tthermophila GBF1 TTHERM 01082910
Ptetraurelia GBF1 GSPATP00015068001
Otrifallax GBF1 Contig359.1.g25Sec7
Chominis BIG ChTU502y2012 313g0065-t32 1-p1
Cparvum BIG cgd3 3480-t26 1-p1
Gniphandrodes BIG GNI 176440-t26 1-p1
Imultifiliis BIG IMG5 173760
Tthermophila BIG TTHERM 00895780
Imultifiliis BIG IMG5 001070
Tthermophila BIG TTHERM 00732790
Ptetraurelia BIG GSPATP00031699001
Ptetraurelia BIG GSPATP00015258001
Ptetraurelia BIG GSPATP00007457001
Ptetraurelia BIG GSPATP00014959001
Imultifiliis BIG IMG5 002360
Tthermophila BIG TTHERM 00569500
Ptetraurelia BIG GSPATP00033814001
Ptetraurelia BIG GSPATP00031880001
Otrifallax BIG Contig10719.0.g52Sec7
Hhammondi BIG HHA 228660-t26 1-p1
Tgondii-ME49 BIG TGME49 228660-t26 1-p1
Ncaninum BIG NCLIV 044930-t26 1-p1
Sneurona BIG SN3 00201230-mRNA-1-p1
Etenella BIG ETH 00004910-t26 1-p1
Cvelia BIG Cvel 19881.t1-p1
Vbrassicaformis BIG Vbra 6697.t1-p1
Pmarinus BIG EER17388
Smicroadriaticum BIG Smic42040
Sminutum BIG symbB.v1.2.015297.t1scaffold1134.1size1359649
Ptricornutum BIG jgiPhatr215422e gw1.19.151.1
Tpseudonana BIG jgiThaps3268862estExt thaps1 ua kg.C chr 50095
Ptricornutum BIG jgiPhatr254591estExt Phatr1 ua kg.C chr 100033
Tpseudonana BIG jgiThaps3263191thaps1 ua kg.chr 7000148
Esiliculosus BIG Ec-10 005790
Esiliculosus BIG Ec-01 004190
Psojae BIG jgiPhyso3360752estExt fgenesh1 pg.C 5t10040
Pultimum BIG PYU1 T000456
Aanophagefferens BIG jgiAuran121199
Ccrispus BIG CDF38205
Gtheta BIG jgiGuith173978e gw1.53.28.1
Bnatans BIG jgiBigna1126930aug1.3 g1638
Ehuxleyi BIG jgiEmihu1464442estExtDG fgeneshEH pg.C 470083
Cparadoxa BIG ConsensusfromContig53335
Athaliana BIG AT3G43300.1
Creinhardtii BIG Cre09.g390319.t1.1
Athaliana BIG AT4G38200.1
Athaliana BIG AT4G35380.1
Athaliana BIG AT3G60860.1
Athaliana BIG AT1G01960.1

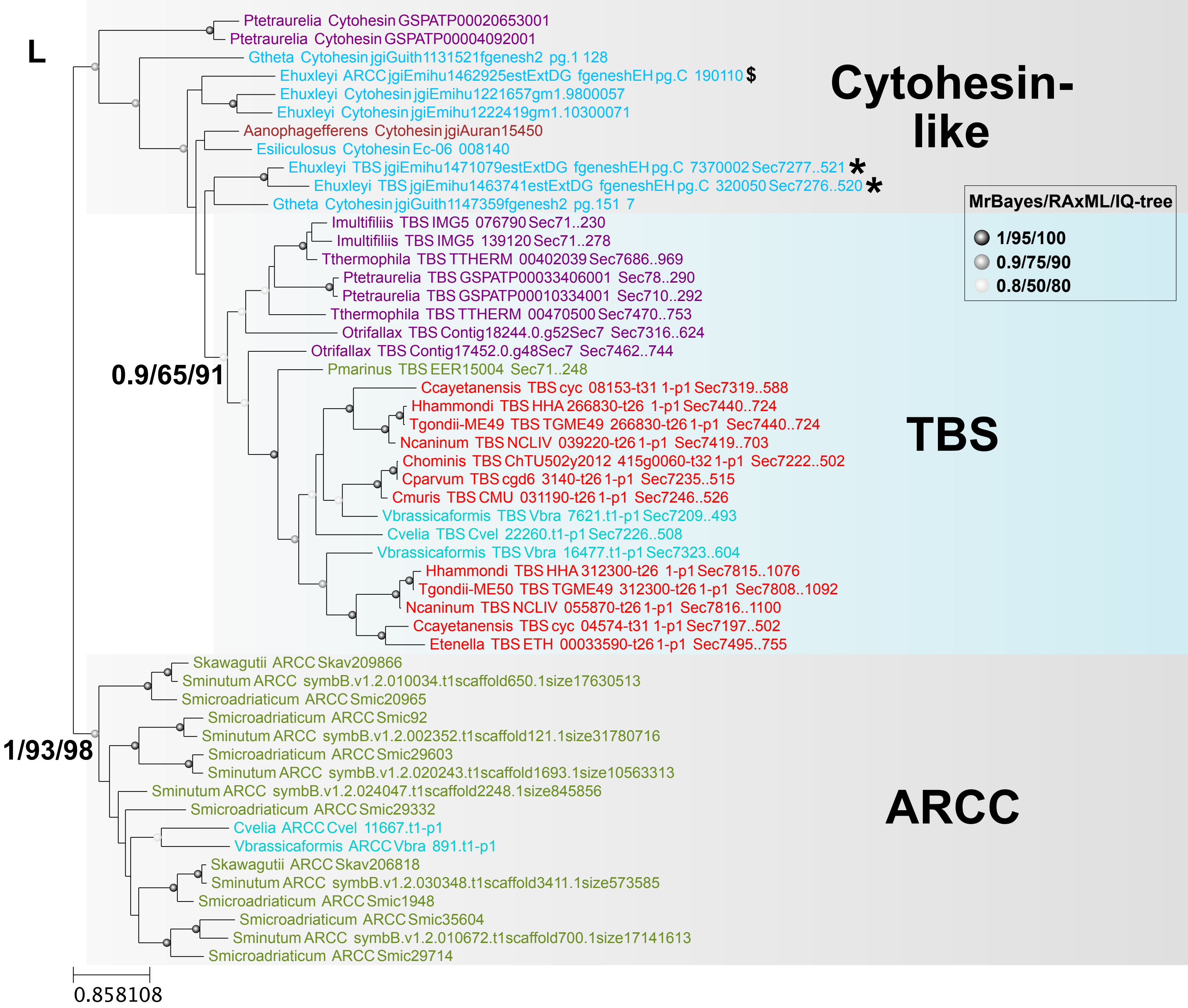
L
Ptetraurelia Cytohesin GSPATP00020653001
Ptetraurelia Cytohesin GSPATP00004092001
Gtheta Cytohesin jgiGuith1131521fgenesh2 pg.1 128
Ehuxleyi ARCC jgiEmihu1462925estExtDG fgeneshEH pg.C 190110 $
Ehuxleyi Cytohesin jgiEmihu1221657gm1.9800057
Ehuxleyi Cytohesin jgiEmihu1222419gm1.10300071
Aanophagefferens Cytohesin jgiAuran15450
Esiliculosus Cytohesin Ec-06 008140
Ehuxleyi TBS jgiEmihu1471079estExtDG fgeneshEH pg.C 7370002 Sec7277..521 *
Ehuxleyi TBS jgiEmihu1463741estExtDG fgeneshEH pg.C 320050 Sec7276..520 *
Gtheta Cytohesin jgiGuith1147359fgenesh2 pg.151 7
Imultifiliis TBS IMG5 076790 Sec71..230
Imultifiliis TBS IMG5 139120 Sec71..278
Tthermophila TBS TTHERM 00402039 Sec7686..969
Ptetraurelia TBS GSPATP00033406001 Sec78..290
Ptetraurelia TBS GSPATP00010334001 Sec710..292
Tthermophila TBS TTHERM 00470500 Sec7470..753
Otrifallax TBS Contig18244.0.g52Sec7 Sec7316..624
Otrifallax TBS Contig17452.0.g48Sec7 Sec7462..744
0.9/65/91
Pmarinus TBS EER15004 Sec71..248
Ccayetanensis TBS cyc 08153-t31 1-p1 Sec7319..588
Hhammondi TBS HHA 266830-t26 1-p1 Sec7440..724
Tgondii-ME49 TBS TGME49 266830-t26 1-p1 Sec7440..724
Ncaninum TBS NCLIV 039220-t26 1-p1 Sec7419..703
Chominis TBS ChTU502y2012 415g0060-t32 1-p1 Sec7222..502
Cparvum TBS cgd6 3140-t26 1-p1 Sec7235..515
Cmuris TBS CMU 031190-t26 1-p1 Sec7246..526
Vbrassicaformis TBS Vbra 7621.t1-p1 Sec7209..493
Cvelia TBS Cvel 22260.t1-p1 Sec7226..508
Vbrassicaformis TBS Vbra 16477.t1-p1 Sec7323..604
Hhammondi TBS HHA 312300-t26 1-p1 Sec7815..1076
Tgondii-ME50 TBS TGME49 312300-t26 1-p1 Sec7808..1092
Ncaninum TBS NCLIV 055870-t26 1-p1 Sec7816..1100
Ccayetanensis TBS cyc 04574-t31 1-p1 Sec7197..502
Etenella TBS ETH 00033590-t26 1-p1 Sec7495..755
Skawagutii ARCC Skav209866
Sminutum ARCC symbB.v1.2.010034.t1scaffold650.1size17630513
Smicroadriaticum ARCC Smic20965
Smicroadriaticum ARCC Smic92
Sminutum ARCC symbB.v1.2.002352.t1scaffold121.1size31780716
Smicroadriaticum ARCC Smic29603
Sminutum ARCC symbB.v1.2.020243.t1scaffold1693.1size10563313
1/93/98
Sminutum ARCC symbB.v1.2.024047.t1scaffold2248.1size845856
Smicroadriaticum ARCC Smic29332
Cvelia ARCC Cvel 11667.t1-p1
Vbrassicaformis ARCC Vbra 891.t1-p1
Skawagutii ARCC Skav206818
Sminutum ARCC symbB.v1.2.030348.t1scaffold3411.1size573585
Smicroadriaticum ARCC Smic1948
Smicroadriaticum ARCC Smic35604
Sminutum ARCC symbB.v1.2.010672.t1scaffold700.1size17141613
Smicroadriaticum ARCC Smic29714
Cytohesin-like
TBS
ARCC
MrBayes/RAxML/IQ-tree
1/95/100
0.9/75/90
0.8/50/80
0.858108

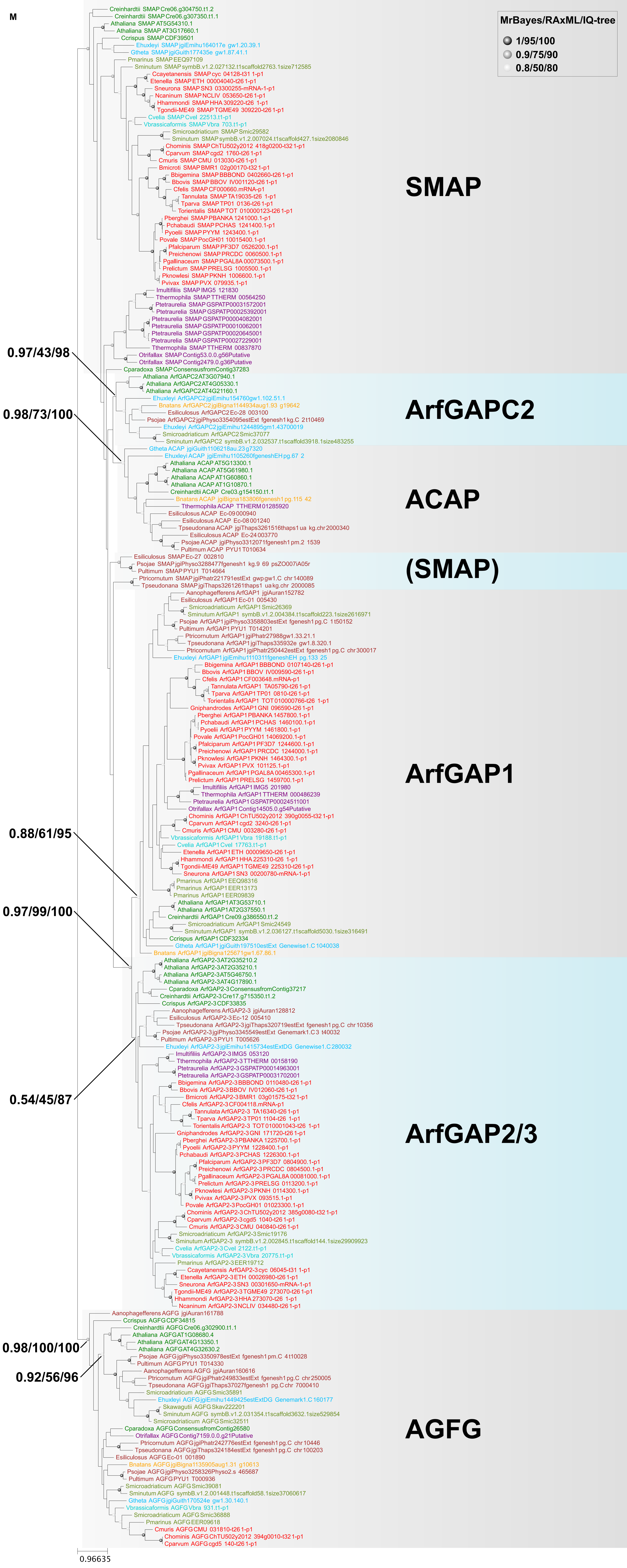
M
MrBayes/RAxML/IQ-tree
1/95/100
0.9/75/90
0.8/50/80
SMAP
ArfGAPC2
ACAP
(SMAP)
ArfGAP1
ArfGAP2/3
AGFG
0.97/43/98
0.98/73/100
0.88/61/95
0.97/99/100
0.54/45/87
0.98/100/100
0.92/56/96
0.96635
